# Convergent evolution during host range expansion and virulence increase in a *Salmonella* bacteriophage

**DOI:** 10.1101/2024.03.13.584857

**Authors:** Amandine Maurin, Marie Vasse, Cécile Breyton, Carlos Zarate-Chaves, Sarah Bouzidi, Juliette Hayer, Jacques Dainat, Margaux Mesleard-Roux, François-Xavier Weill, Ignacio G. Bravo, Alexandre Feugier, Rémy Froissart

## Abstract

Viral host range expansion is predicted to evolve at the cost of reduced mean fitness. We investigated the adaptive walks of a virulent phage (*Tequintavirus*) in a spatially variable environment composed of four susceptible bacterial isolates and four resistant ones (*Salmonella enterica* serotype Tennessee, sequence types ST5018 and ST319 respectively). Starting from a single ancestral phage, we evolved multiple independent populations through serial passages on non-coevolving bacteria, following the Appelmans protocol. The phage populations evolved an expanded host range and increased virulence. Whole-genome sequencing revealed recurrent parallel mutations across populations (i.e. convergent evolution), particularly in genes encoding exo- and endo-nucleases, dUTPase, and caudal proteins. Notably, two parallel mutations in the gene coding for the *Long Tail Fibre* became fixed early in the evolutionary trajectories. Reverse-genetics experiments introducing these mutations into the ancestral genome expanded the host range but yielded only marginal increases in virulence, highlighting the effect of compensatory mutations.

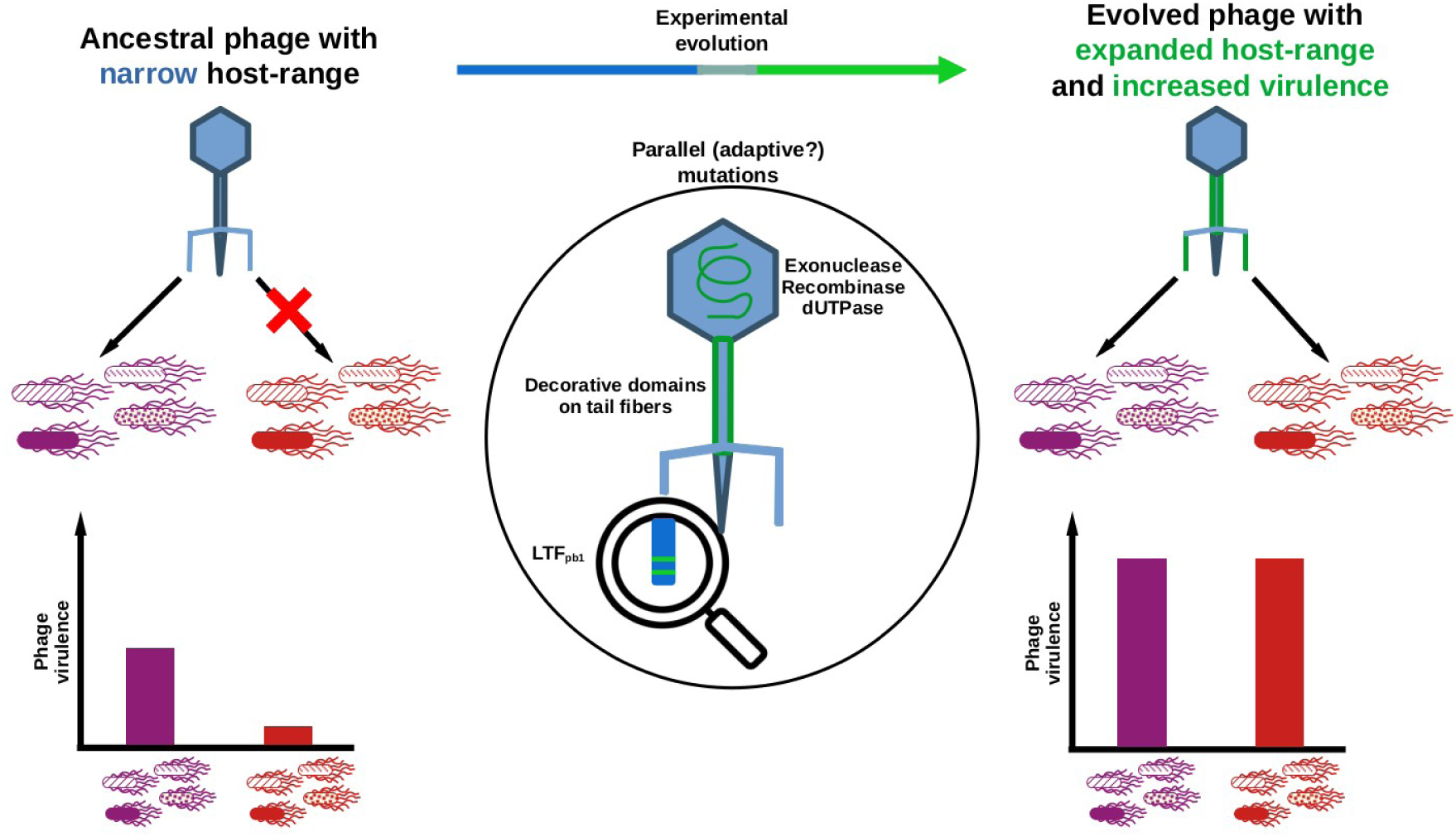

**Highlights:**

• A phage (*Tequintavirus*) was evolved on susceptible and resistant *Salmonella enterica* strains
• Experimentally evolved phage populations displayed expanded host range and increased virulence
• Convergent evolution revealed adaptive mutations modifying receptor recognition in caudal proteins
• Reverse-genetic showed implication of two Long Tail Fibre mutations in host range expansion

**In Brief:** Generalism is traditionally predicted to evolve at the cost of lower mean fitness. Contrary to this textbook view, we demonstrate that generalist phages with expanded host range and increased virulence can readily evolve *in vitro* and be purposely optimized for phage therapy applications.

## Introduction

Viruses infecting bacteria, known as bacteriophages or phages, have an enormous impact on bacterial population dynamics across diverse environments ^1,2^ including oceans ^3,4^, soils ^5^ and the human gut ^6,7,8^. Most bacterial communities consist of multiple species and genotypes, and one could thus expect that the best strategy would be that phages infect and replicate in a broad range of hosts ^9,10^. Contrary to this expectation, most phages display a narrow host spectrum, typically infecting only a few bacterial species ^11,12,13,14^ or genotypes within a single species ^15,16^. The prevalence of such specialist phages remains an open question ^17^ tied to long-standing debates about the evolution of specialisation and the potential costs of generalism ^18,19^. One hypothesis among others to explain this cost is the antagonistic pleiotropy of adaptive mutations. It suggests that mutations are advantageous in one environment but deleterious in another ^20^, which limits the evolution of broad host ranges in phages.

Several studies on phage-bacteria interactions have evaluated the impact of phage mutations on host range expansion ^21^. A strong negative relationship has been reported between replication rates and host range of different members of *Caudoviricetes* infecting genotypes of *Escherichia coli* ^22^ and as well as *Klebsiella* ^23^. Similarly, experimental evolution of the phage ɸX174 (*Microviridae*) on *Salmonella enterica* resulted in a reduced ability to infect *E. coli* ^24^. Moreover, most mutations conferring an expanded host range of ɸ6 phage (*Cystoviridae*) also impose significant fitness costs on the original host, presumably due to antagonistic pleiotropy ^25,26^. These findings suggest that mutations expanding phage host ranges seem to carry selective costs, potentially explaining the rarity of generalist phages.

To bridge fundamental research seeking to understand the dynamics of selection favouring specialist *versus* generalist organisms with a practical application of such research, we adapted a phage to infect multiple genotypes of *Salmonella enterica*.

Nontyphoidal *Salmonella* infections are a significant public health concern, representing the second most common cause of bacterial outbreaks in humans (6%) with the leading cause of enteric illness in humans being direct contact with animals (46%) ^27^. In Europe, *Salmonella* was the second most reported zoonosis, with 60,050 confirmed cases in 2021 ^28^. Unfortunately, current cleaning methods, sanitation procedures and antimicrobial strategies are insufficient to eliminate pathogenic *Salmonella* contaminations, particularly in the food industry, due to the development of resistant phenotypes, such as those associated with biofilm formation and efflux pump expression ^29,30^. Additional methods are thus needed to control this major bacterial pathogen, and phages present a promising alternative. Phages have demonstrated efficacy against *Salmonella* both as standalone biocontrol agents ^31,32^ and in combination with chemical disinfectants ^33^, offering potential solutions for addressing these health challenges.

The aim of our study was to trace the adaptive walk of a phage exposed to an experimental evolution procedure designed to foster changes in host range and virulence. To evaluate phage virulence, we compared bacterial growth in the presence or absence of either “ancestral” or “evolved” phages. The underlying rationale was that changes in life history traits of a phage (i.e. modification of burst size, latency periods and/or adsorption rates) will modify bacterial population dynamics ^34,35^. Using serial passages of a virulent phage (*Tequintavirus*) with a modular genome ^36^ following the Appelmans protocol ^37^, we generated several evolved phage populations and compared these to the ancestral phage population (i.e. from before the experimental evolution process). The evolved populations showed expanded host range and increased virulence compared to the ancestral phage population. Through reverse-genetics, we directly demonstrated the role of two adaptive mutations in host range expansion and highlighted the need of compensatory mutations to achieve higher virulence. Our results pave the way for future development of phage biocontrol on pathogenic bacteria. By rapidly generating phages with high efficacy across multiple host genotypes, this approach has the potential to enhance biocontrol efficiency while reducing the likelihood of bacteria developing resistance, a major factor contributing to biocontrol failure.

## Materials and Methods

### Bacterial and Phage Strains

Thirty-one isolates of *S. enterica* were collected by swabbing a food processing factory in Poland, over a two-year period (2017-2019). They belong to the Tennessee serotype, as determined by Eurofins (Aix en Provence, France). These isolates are referred to hereafter with the prefix “SeeT” followed by a number from one to thirty-one. Bacteria were routinely cultured in lysogeny broth (LB Lennox, Athena Enzyme Systems; Baltimore, MD, USA) or LB agar (1,2%). Eight out of the 31 SeeT isolates were used in the evolution experiment described below. The isolates belonged to two Sequence Types (ST), ST5018 and ST319, as determined by the French National Reference Center (CNR) for *Escherichia coli*, *Shigella* and *Salmonella* at the Institut Pasteur (Paris, France).

To prepare stock bacterial solutions, isolated colonies grown on LB agar plates were transferred to 7 mL of LB in soda glass cell culture test tubes (16 mL nominal capacity; #11823370; Fisher Scientific, Illkirch, France) and incubated overnight at 35°C with shaking (150 rpm; MaxQ 4000, Thermo Fisher). The following day, 500 µL of overnight bacterial culture was aliquoted into each well of a 96-deepwell polypropylene plate (1 mL nominal capacity; #135702, Dutscher, Bernolsheim, France), mixed with 250 µL of 60% (v/v) glycerol (#G6279, BioXtra, ≥99%; Sigma-Aldrich, St. Louis, MO, USA), and stored at -70 °C (CryoCube F750h, Eppendorf). This frozen plate is referred to as the stock deepwell plate. For experiments, new 96-deepwell plates were filled with 500 µL of LB, inoculated from the stock deepwell plate using a 96-pin replicator (#140500, Boekel Scientific; Feasterville, PA, USA) and grown overnight at 35°C with shaking (450 rpm; Aqualytic ventilated incubator with Titramax 101, Heidolph). After overnight growth, each well contained approximately 10^8^ CFU/mL ± 1x10⁸ CFU/mL (estimation from easySpiral plater; #412 000, Interscience, Saint-Nom-la-Bretèche, France). These plates, stored at 4°C, are referred to as the inoculation deepwell plates.

The phage used in this study was isolated from Marseille’s wastewater (November 2017), filtered through a 0.22 µm Minisart polyethersulfone (PES) filter (#16541--K; Sartorius, Göttingen, Germany), and stored in glass bottles. To amplify potential phages, 500 µL of LB in each well of a 96-deepwell plate was inoculated with 2 µL of an overnight culture of SeeT2 using a 96-pin replicator, followed by the addition of 50 µL of filtered wastewater. The plate was incubated overnight at 37°C with shaking (450 rpm, 1.5 mm orbital; Titramax 101 #544-11300-00; Heidolph Instruments, Schwabach, Germany). The following day, 50 µL of chloroform was added to each well and the plate was incubated at 4°C for at least four hours. From the supernatant, 2 µL were transferred to a new 96-well polystyrene plate (PS, #82.1581001; Sarstedt, Nümbrecht, Germany), containing 200 µL of LB supplemented in 10mM CaCl_2_ (Sigma-Aldrich, #C3881) and inoculated with 2 µL of an overnight culture of SeeT2. Bacterial growth was monitored by evaluating turbidity through the measure of Optical Density at 600 nm wave length (OD_600nm_) over 16 to 20 hours at 37°C with shaking (300 rpm, spectrophotometer FLUOstar Omega, BMG Labtech, Ortenberg, Germany). Solutions present in wells that showed delayed bacterial growth were transferred to polypropylene tubes (Eppendorf). Residual bacteria were cleared by adding 10% chloroform and centrifuged (10 min at 15,871 Relative Centrifugal Force or rcf, Eppendorf 5415 R) to keep only the phages.

To purify a phage, we used the double-layer method ^38^. This involved mixing 100 µL of the appropriate dilution of the phage solution with top LB agar (6g/L; #LF611001 Liofilchem, Italy) previously mixed with 100 µL of SeeT2 overnight culture. After an overnight incubation at 35°C, we collected one isolated lysis plaque from the top agar into 200 µL SM buffer (NaCl 100 mM, MgSO_4_ 10 mM, Tris-HCl 50 mM, pH = 7.4), and stored the tube at 4°C for over an hour. The phage was then purified through five consecutive rounds of the double-layer method, picking one isolated lysis plaque at each round. In the last round, we collected the full top LB agar layer in SM buffer, centrifuged (10 min, 3,000 rcf, Eppendorf Centrifuge 5702R) and filtered it through a 0.22 µm PES filter, then stored it at 4°C in polypropylen 15 mL tubes (#352096; Falcon, Corning, Mexico). We thus isolated one phage, that we named *Salmonella phage Tennessee Salten*, and hereafter called Salten.

### Experimental Evolution

To allow for the host range expansion and increased virulence of Salten, we first separately exposed high concentrations of the phage to several SeeT bacterial isolates. The ones able to grow in the presence of phages were considered resistant, and the ones showing no growth were considered susceptible. We then picked four susceptible SeeT isolates (SeeT2, SeeT4, SeeT7 and SeeT17 from ST5018) and four resistant SeeT isolates (SeeT1, SeeT3, SeeT5 and SeeT6 from ST319). Following the Appelmans protocol ^37^, the Salten phage was separately exposed to each of these eight bacterial isolates. Briefly, at each passage, 200 µL of LB with 10 mM CaCl_2_ was dispensed into each well of a 96-well PS plate (referred to as the test plate). The wells were then each inoculated with 2 µL of the corresponding bacteria stock from the inoculation deepwell plate, as well as with 2 µL of phages at different concentrations in each well - in both cases using a 96-pin replicator. Phage dilutions were prepared in a reusable 96-well polypropylene plate (#290-8353-03R; EVCORP, USA) by serial ten-fold dilutions in 180 µL of SM buffer with 20 µL of filtrated phage solution. Note that the reusable plates were prepared with bleach decontamination, washed in a dishwasher, rinsed with ionized water, dried at room temperature and then autoclaved individually. The particularity of the Appelmans protocol is that each bacterial isolate is submitted to different concentrations of phages at each passage, in such a way that ratios of bacteria and phage vary between high to low multiplicities of infection (MOI) depending on the well. Positive controls (LB with 10 mM CaCl_2_ inoculated with bacteria) and negative controls (LB with 10 mM CaCl_2_) were included in test wells. Bacterial growth kinetics were monitored at 37°C with shaking (300 rpm) and OD_600nm_ (spectrophotometer FLUOstar Omega) for at least 16 hours. Wells presenting delayed to complete inhibition of bacterial growth compared to positive controls were harvested, pooled into a single 15 mL polypropylene tube, cleared with 500 μL chloroform, mixed, centrifuged (10 min at 3,000 rcf) and filtered through a 0.22 µm PES filter. The resulting phage solution was stored at 4°C. Phage concentrations between passages were not measured. We performed five independent replicates - or lineages - of the full experimental protocol. The first lineage involved six serial passages of the Appelmans protocol (called “SaltenE1”, E for “evolved”). Three additional independent lineages (SaltenE2, SaltenE3 and SaltenE4) underwent seven serial passages several months later. Finally, after an 18-month pause to ensure no residual Salten phages were present in the laboratory, we conducted an additional evolution experiment (SaltenE5) with eight serial passages.

### Estimation of Host Range and Virulence

We assessed the host range and virulence of the ancestral Salten and the evolved SaltenE populations in liquid conditions by following bacterial growth in 96-well PS plate. Phage concentrations were estimated by spot-assays ^38^ on top-agar containing SeeT17. Test plates were prepared as described for experimental evolution, with wells inoculated with either Salten or an evolved SaltenE population at five multiplicities of infection (MOIs) ranging from 1 to 0.0001 (ten-fold dilutions). Each experiment was replicated three times on different days.

Host range was further assessed in solid culture using the spot-assay method on 18 additional SeeT isolates beyond those used for evolution. Presence/absence of plaques was visually evaluated the next day (examples in Fig. S5). Each experiment was tested three times on different days.

### Phage Morphology

Salten solution (15 mL at 10^11^ PFU/mL) was concentrated into 1.5 mL tubes by two one-hour rounds of centrifugation at 16,000 rcf, 4°C (Eppendorf Centrifuge 5415R). The pellet was resuspended in 600 µL 100 mM ammonium acetate (Sigma-Aldrich) and filtered through a 0.22 µm PES filter. Phages were then adsorbed onto a Formvar/carbon 300 grid (# CU 50/BX 9012.90.0000; Electron Microscopy Sciences, Hatfield, PA, USA), contrasted with 2% uranyl acetate, and visualized via transmission electronic microscopy (TEM, JEM-1400Plus, JEOL, Akishima, Tokyo, Japan).

### DNA Sequencing

Whole-genome sequencing was performed for the eight bacterial isolates used in the evolution experiment (SeeT1, 2, 3, 4, 5, 6, 7 and 17) by the reference center CNR (Paris, France). Bacterial DNA was purified using the Maxwell 16-cell DNA purification kit (Promega) and sequenced by Illumina NovaSeq technology (paired-end). For identity verification, a region of the *cpn60* gene ^39^ (Table S1) was amplified and sequenced via Sanger sequencing for all 31 SeeT isolates (PCR from boiled colonies and using Roche Taq DNA polymerase [#11146165001; Merck KGaA, Darmstadt, Germany]).

Whole-genome sequencing was conducted on the ancestral phage Salten and on four lineages of evolved populations, SaltenE1, E2, E3 and E4 (SaltenE5 phage population was not sequenced because it was obtained after the DNA sequencing experiment of previous lineages). DNA extraction was done according to a protocol ^40^ adapted by Nicolat Ginet (Bacterial chemistry Laboratory, Marseille, France) after amplification of each phage population on the most susceptible isolate, SeeT17. Briefly, genetic material of bacterial origin potentially surrounding the phages was eliminated by adding 10 µL DNAse I (1 U/µL; #D5307; Sigma-Aldrich), 5 µL RNAse A (10mg/mL; #EN0531; Thermo Fisher) and 2 µL Dpn I (10 U/µL; #ER1702; Thermo Fisher). Phage DNA was extracted using phenol-chloroform-isoamyl acid 25/24/1 (#77617; Sigma-Aldrich). After DNA quantification with NanodropOne (Thermo Fischer) and Qubit 4 Fluoremeter (Invitrogen, Thermo Fisher), we multiplexed the different DNA extractions using the NebNext Ultra II FS Library Prep Kit (#E7805, #E6440 and #E6177; NEB). Phage DNA was fragmented with a transposase enzyme and prepared for multiplexing according to the recommendations of the supplier for inputs over 100 ng. Fragmented end-prepared DNA was ligated to Illumina adaptors and then sorted with beads to select fragment sizes between 150-250 bp, for a final fragment size between 270-370 bp. Adaptor-ligated DNA was enriched by four cycles of PCR. These amplicons were then cleaned and ligated with a unique pair of primers. The final library concentration was evaluated through Qubit 4 and fragment sizes were checked by migration using QIAxcel Advanced Instrument (QIAgen, Hilden, Germany). Fragments ready for sequencing were sized between 280 bp and 320 bp. DNA libraries were sequenced in paired-ends using an Illumina in-lab sequencer (iSeq100 instrument). Raw reads were deposited in the European Nucleotide Archive (https://www.ebi.ac.uk/ena/browser/support) with the accession numbers ERR13191102 (Salten), ERR13191103 (SaltenE1), ERR13191104 (SaltenE2), ERR13191105 (SaltenE3) and ERR13191106 (SaltenE4).

To determine the order of occurrence of mutations across serial passages, we designed primers flanking the mutations of interest in the viral ORFs *pb1*, *pb2* and the recombination related exonuclease encoding gene (Table S1). Amplicons were obtained using Q5 high-fidelity DNA polymerase (#M0491; New England Biolabs NEB) and sent for Sanger sequencing to Eurofins Genomics France (Nantes, France).

### Bioinformatic Analysis and Annotation

Reads of the eight SeeT isolates used in our experiment were assembled by the CNR (Paris, France), using SPAdes v3.15.2 ^41^. SeeT genomes were virtually genotyped using the Salmonella multilocus sequence typing scheme based on seven housekeeping genes (MLST7) and the core genome MLST (cgMLST) scheme based on the analysis of 3002 genes ^42^. Both genomes have been deposited in EnteroBase (https://enterobase.warwick.ac.uk/; barcodes SAL-QB8964AA - named Sten1 for SeeT1, SAL-QB8962AA - named Sten2 for SeeT2, SAL-QB8963AA - named Sten3 for SeeT3, SAL-QB8959AA - named Sten4 for SeeT4, SAL-QB8958AA -named Sten5 for SeeT5, SAL-QB8957AA - named Sten6 for SeeT6, SAL-QB8960AA - named Sten7 for SeeT7 and SAL-QB8961AA - named Sten 17 for SeeT17). Baargin workflow ^43^ was used for genomic assembly and annotation. PanExplorer online tool ^44^ was used for core-genome visualisation and genetic diversity analysis between the eight SeeT isolates, while DefenseFinder ^45^ was employed to identify defense mechanisms against phages. Phaster ^46,47^ was used to detect prophages within the eight SeeT genomes. Phylogenetic relationships among the eight SeeT isolates were inferred using the nucleotide sequences of the *cpn60* gene, aligned with Muscle ^48^, checked at the codon level using AliView v.1.28 ^49^ and then manually curated. Jmodeltest ^50^ identifed JC69 as the most suitable nucleotide substitution model. Maximum-likelihood phylogenetic inference was performed with PHYML ^51^ using 1,000 bootstrap cycles.

After Illumina sequencing, phage read quality was assessed using FastQC v.0.12.1 ^52^. Primers were trimmed with Fastp v.0.22.0 ^53^, using default parameters. Phage genome contigs were prepared following the workflow recommended in Turner et al. ^54^. *De novo* phage genome assembly was carried out with SPAdes v.3.14.1 ^55,56^ with default parameters. The Salten complete genome sequence, including the two terminal repeated sequences, was generated with PhageTerm Virome v.4.3 ^57^. For the ancestral Salten phage, we obtained a single contig of 110,076 nt with high coverage (average 400 reads depth), and a number of short contigs (below 1200 nt) with low coverage (average 10-20 reads depth).

The ancestral Salten contig as well as the contigs of each evolved phage population were polished with Pilon v.1.24 ^58^, using their respective reads. The new contig of 109,999 nt was deposited in the European Nucleotide Archive (https://www.ebi.ac.uk/ena/browser/support) with Accession Number OZ075147. It was annotated thanks to the Genome Annotation online tool from the Bacterial and Viral Bioinformatics Resource Center ^59^ (BV-BRC, https://www.bv-brc.org/), based on Tequintavirus annotation (Taxonomy ID = 187218). Structural genes were annotated manually based on Zivanovic et al (2014) ^60^, Linares et al (2023) ^61^, using blastn or blastp on the NCBI platform (Blast® services, available from: https://www.ncbi.nlm.nih.gov/Blast.cgi). The gggenes package v.0.5.0 ^62^ was used to generate the genome mapping of Salten. breseq v.0.38.1 ^63^ was used as a pipeline for calling of single nucleotide polymorphism (SNPs) and small insertions/detections (indels). The associated gdtools COMPARE tool was used to visualize the nature, locations and frequencies (percentage of reads containing a mutation at a position of interest) detected by breseq.

### Phage Mutagenesis by Reverse-Genetics

The two parallel mutations identified in the Long Tail Fibre *pb1* ORF of evolved SaltenE phages were introduced into the ancestral Salten *pb1* ORF by directed mutagenesis. Mutations were confirmed in the *pb1* ORF in SaltenE populations by Sanger sequencing (Eurofins, Nantes, France). The amplicons were obtained by PCR using Q5 high-fidelity DNA polymerase (#M0491; New England Biolabs NEB) and the primers Salten-pb1-3183-F / Salten-pb1-4136-R primers (Table S1). We then amplified by PCR the surrounding region of these mutations (Salten-pb1-3374-F and Salten-pb1-3557-Rv), using both evolved and ancestral Salten (negative control) DNA as templates. PCR products were then introduced into pBBR1-MCS2 plasmid at *Xho*I and *Hind*III sites (#R0146 and #R0104; New England Biolabs NEB). We heat-shock transformed *E. coli* DH10β made competent in the laboratory with the plasmid containing the amplicon, and let them grow on LB agar supplemented with 50 µg/mL kanamycin, 50 µg/mL Xgal and 10 µM IPTG. The transformation yielded clones harbouring a plasmid with an insert presenting *pb1* partial sequence with or without the two potential adaptive mutations (called pBBR1-E or pBBR1-A, respectively). To control for any plasmid effect, we also transformed *E. coli* DH10β with the empty vector (called pBBR1-EV). White colonies were selected and amplified, DNA extracted (Monarch Plasmid Miniprep kit, #T1010; NEB), the *pb1* target amplified (primers pBBR1-MSC2-F and Salten-3470-F; Table S1) and the products Sanger-sequenced (Eurofins, Nantes, France) to select appropriate bacterial clones. The bacterial SeeT17 isolate was then transformed with either pBBR1-A, pBBR1-E or pBBR1-EV by electroporation (∼100 ng/µL) using a MicroPulser electroporator (Bio-Rad, Hercules, CA, USA) on the Ecoli1 program and then plated on LB agar supplemented with 50 µg/mL kanamycin. Resistant colonies were selected and colony-PCR amplicons (primers pBBR1-MSC2-F and Salten-pb1-3470-F; Table S1) were sent for sequencing. Transformed SeeT17 with the appropriate inserts were grown in glass tubes containing LB supplemented with 50 µg/mL kanamycin. When the culture reached OD_600nm_ = 0.2, we added the ancestral phage, Salten (MOI = 0.001), in order to allow for natural recombination between the phage and the insert present on the plasmid. After an overnight co-culture, phages were recovered by removing bacteria with centrifugation (30 min at 3,000 rcf) and filtration of the supernatant (0.22 µm; PES #16541-K, Sartorius). Phage populations coming from overnight co-culture with SeeT17 transformed with pBBR1-E, pBBR1-A or pBBR1-EV were respectively named rE_Salten, rA_Salten and rEV_Salten.

Phage populations (rE_Salten, rA_Salten, and rEV_Salten) were tested on resistant (SeeT1, SeeT3, SeeT5, and SeeT6 from ST319) and susceptible (SeeT17 from ST5018) bacterial isolates using the double-layer method. In cases where plaques appeared, isolated plaques were picked and resuspended into 200 µL SM buffer and cleared with 20 µL chloroform. One microlitre was used as DNA matrix to produce PCR amplicons with Phusion High Fidelity Taq Polymerase (#M0530; NEB) and the new primers Salten-pb1-3183-F and Salten-pb1-4136-R (Table S1). To avoid any plasmid amplification, these latter primers targeted a *pb1* phage region outside the sequence inserted into the pBBR1 plasmid. We also verified the presence of the two parallel mutations within an isolated rE_Salten lysis plaque by sequencing amplicons targeting a large region of the gene *pb1* (larger than the plasmid insert). Additionally, to rule out any accidental contaminations with an evolved phage, we sequenced a region of the *pb2* ORF harbouring a specific sequence signature of either the ancestral or evolved phages (using primers Salten-pb2-8720-F and Salten-pb2-9129-R; Table S1). This verification confirmed that rE_Salten harboured the ancestral *pb2* sequence.

### Statistical Analyses

Data were analysed using R software (2023-04-21, R Core Team, v.4.3.0 ^64^) in Rstudio (2020-04-01, RStudio team, v.1.2.5042 ^65^). We estimated the impact of phages on bacterial growth in liquid cultures using bacterial kinetic data of OD_600nm_ over at least 16h (Fig. S3). For each MOI, a virulence index (Vi) was obtained following the formula ^66^

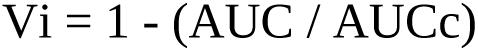

where AUC corresponds to the area under the curve of bacterial growth in the presence of phages and AUCc the area under the curve of the bacterial growth control (i.e. without phages). The AUCs were computed with the linear option of the MESS package (v.0.5.9 ^67^). For phage phenotypic characterisation, we analysed the Vi data at MOI = 0.01, using a linear mixed-effect model with identity of phages (Salten *versus* each of the five evolved SaltenE populations; n = 5), identity of bacterial isolates (n = 8) nested into ST (n = 2) and their interactions as fixed effects, and the experimental plates (three replicates per phage, n = 15) as a random effect. Reverse-genetics experiments were analysed similarly, with the replicates (n = 3) treated as a random effect.

After an assessment of data normality, statistical models were followed by type III ANOVA (package car v.3.1-2 ^68^) and contrasts on marginal means with the Tukey method (package emmeans v.1.8.7 ^69^). When needed, a one-sample t-test against 0 with correction for multiple testing (Benjamini & Hochberg correction) was done. ggplot2 package v3.4.2 ^70^ was used to construct plots. All R scripts and datasets are available on GitLab at https://src.koda.cnrs.fr/MAURINAmandine/Salten.

## Results

### Characterization of Bacterial Isolates

Over a two-year period, 31 isolates of *S. enterica* subspecies *enterica* serotype Tennessee (SeeT) were collected from a food processing factory. Eight of these isolates were selected for a detailed study of phage-bacteria interactions. Whole-genome sequencing and *in silico* serotyping confirmed that all eight isolates belonged to serotype Tennessee (antigenic formula 6,7:z29). Multilocus sequence typing on seven housekeeping genes (MLST7) identified two sequence types (ST): ST319 (SeeT1, SeeT3, SeeT5 and SeeT6) and ST5018 (SeeT2, SeeT4, SeeT7 and SeeT17). Furthermore, cgMLST was used to analyse differences between 3,002 genes of the core genome in our eight isolates. We identified phylogenetic divergence between ST319 and ST5018 (differences up to 900 alleles of the 3,002) and classified them in two clusters, HC900_139 and HC900_131799 respectively. Approximately ninety one percent (90.7%) of the complete genome of the eight genotypes corresponded to the core genome (PanExplorer ^44^). Clustering based on presence/absence of genes in the accessory-genome confirmed the separation into two STs (Fig. 1A), with 300 to 354 present/absent genes differing between the two STs (Fig. 1B).

**Figure 1.**
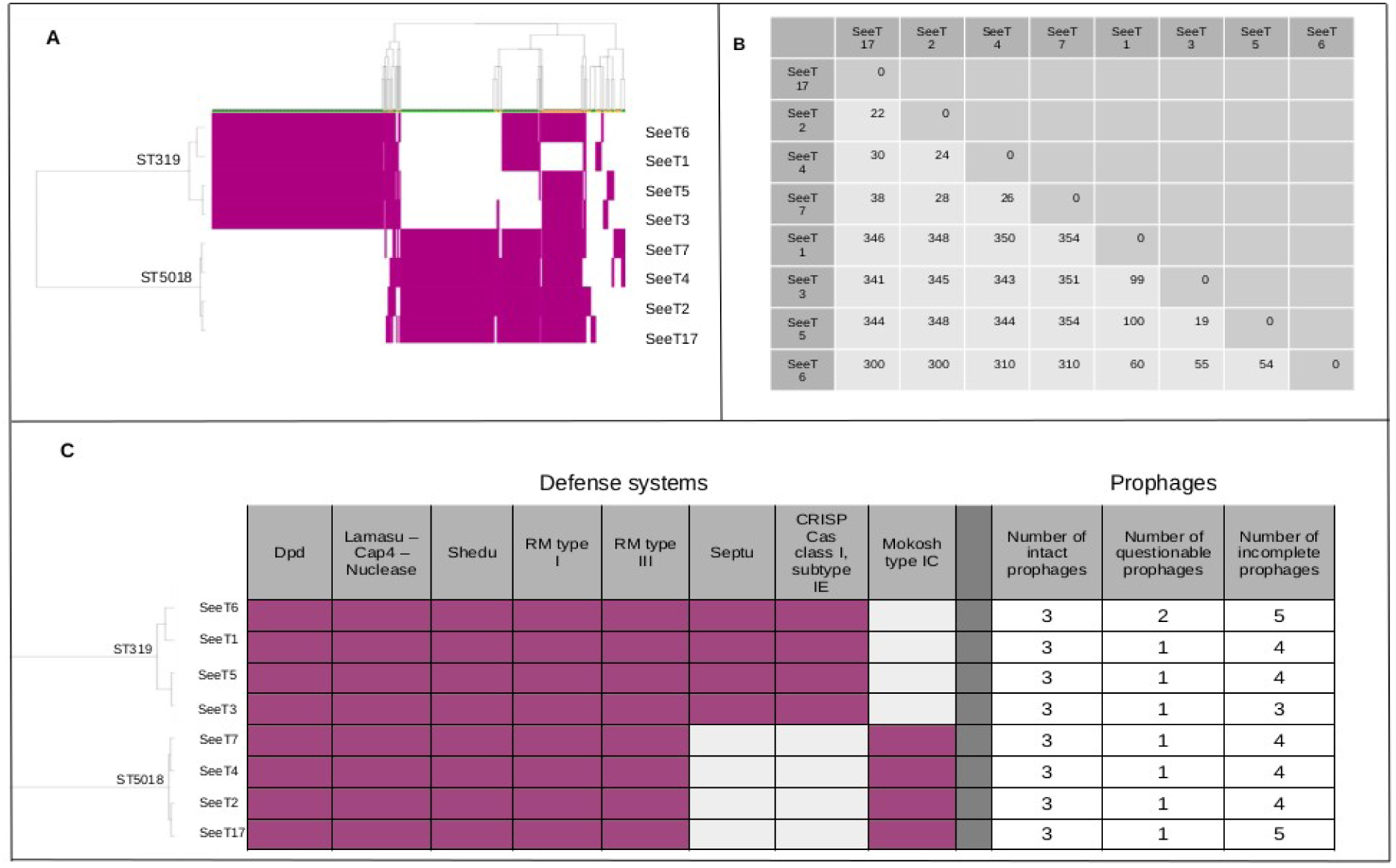
Genetic diversity in the accessory-genome of the eight SeeTs used for the evolutionary training. **A** Absent/present (respectively white/purple) genes clustering heatmap generated by PanExplorer on the accessory-genome. Each column of the dendrogram (on the top) represents a gene, and these are hierarchically clustered (each cluster is coloured green and yellow alternately). The dendrogram derived from the heatmap (on the left) illustrates the genetic relationships among the SeeTs. **B** Matrix summary of Fig. 1A, based on the presence/absence of genes in the accessory-genome, between SeeTs. The SeeT17 genome was used as a reference genome. SeeTs are classified according to their STs. Data should be read as follows: there are 22 genes that are differentially present/absent when comparing the genomes of just SeeT2 and SeeT17. **C** Defense systems that are differentially absent/present in different isolates and the numbers of prophages that are inferred to be present in each of the bacterial isolates.

Some of the differently present genes are likely involved in defense mechanisms. DefenseFinder detected defense systems common to both ST (Dpd, Lamassu-Cap4-nuclease, Shedu and RM type I and III) as well as some ST-specific mechanisms: the Septu defense system was exclusive to ST319, while the Mokosh type I C defense system was unique to ST5018 (Fig. 1C).

The ST5018 isolates were more genetically homogeneous than the ST319 isolates with ST5018 differing from one another by between 22 and 38 differentially present genes, and the ST319 isolates differing from one another by 19 to 100 differentially present genes.

We then focused on genes coding for potential phage receptors such as those involved in the lipopolysaccharide (LPS) synthesis. Based on the different genes encoding assembly-involved and structural proteins of LPS listed in Adler et al (2021) ^71^, we retrieved the protein sequences of each of these genes (annotated using Bakta tool ^72^ integrated into the baargin workflow) for each of the eight isolates and aligned these using Uniprot ^73^ (https://www.uniprot.org/align). We thus identified three genes encoding amino acid polymorphisms that differentiated between ST5018 isolates and ST319 isolates (where isolates from the same ST had 100% identical amino acid sequences for these proteins; Fig. S1A,B). These differences were: (i) V159A (valine in ST5018 and alanine in ST319) within the fepE protein, which is known to regulates LPS O-antigen chain length; (ii) D181N (aspartic acid in ST5018 and asparagine in ST319) and R222K (arginine in ST5018 and lysine in ST319) within waaK which is a a1,2-N-acetylglucosaminyltransferase; (iii) M463T (methionine in ST5018 and threonine in ST319) within wzxC which is a translocase involved in colanic acid synthesis. Analysis of FhuA, another potential phage receptor, identified two amino acid differences between ST5018 and ST319, both located in the inner bacterial membrane, indicating that the amino acids involved are unlikely to interact directly with phages (Fig. S1C).

Finally, prophage analysis with Phaster ^46^ identified three intact prophages in each isolate, along with three to five incomplete or questionable prophages (Fig. 1C), most classified as *Enterobacteriaceae* phages (Salmophages, Coliphages, Enterophages, see Fig. S2). Functional temperate phages were confirmed in the ST5018 isolate SeeT2 and the ST319 isolates SeeT3 and SeeT5, as indicated by the occasional appearance of turbid plaques on their bacterial lawns.

### Phage characterization

Transmission electron microscopy (TEM) of the ancestral phage which we named Salten, revealed a T5-like Siphophage morphology, with a long non-contractile flexible tail approximately 180 nm in length, attached to an icosahedral head with diameter of 60 nm (Fig. 2A).

**Figure 2.**
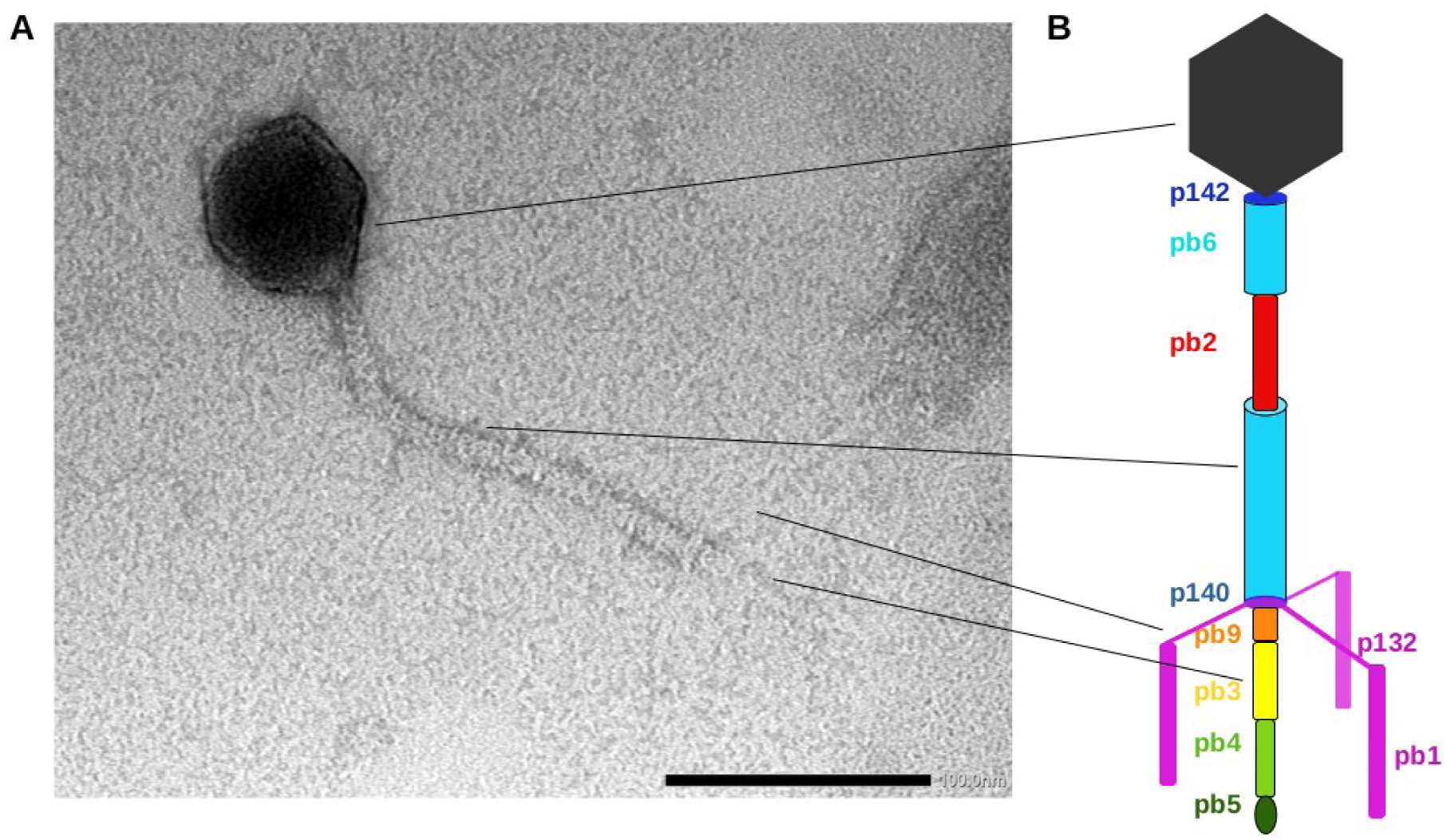
Morphology of the *Tequintavirus* Salten. **A** Transmission electron microscopy (TEM). **B** Schematic structure of T5 adapted from Zivanovic et al. ^60^ with pb6: cyan; p142: blue; p132 and pb1: pink; pb9: orange; pb3: yellow; pb4: green, pb5: dark green. The scale bar represents 100 nm.

The isolated phage Salten has a genome containing 109,999 base pairs (coverage min = 295x and max = 1171x; after assembly by Spades, polish by Pilon and sequence consensus created by breseq) with approximately 39% GC content. Based on an analysis of 94 phage genomes from the same species (ANI >80%), PanExplorer ^44^ identified a core genome consisting of 29 genes. The Salten genome contains all 29 of these core genes. The closest described phage is the *Escherichia* phage HildyBeyeler strain Bas33 (GenBank accession: MZ501074.1). Salten covers 85% of the total Bas33 genome, with 97% nucleotide identity within this portion after a blastn analysis. Concerning the core genome shared with Bas33, Salten exhibits 97% protein identity and 93% nucleotide identity. Salten thus represents a member from a new species within the *Tequintavirus* genus ^74^.

Given that the structure of the T5 tail is available ^61^ as a model, we described by analogy the caudal structure of Salten (Fig. 2B). Specifically, the T5 tail of Salten is composed of a tube formed by the Tail Tube Protein pb6 (TTP_pb6_, cyan) buried under the collar which serves as an anchor for the three lateral Long Tail fibres formed by pb1 (LTF_pb1_, purple). At the extremity of the central fibre is located the Receptor Binding Protein pb5 (RBP_pb5_, dark green). The length of the tube is determined by the Tape Measure Protein pb2 (TMP_pb2_, red), which is located in the lumen of the tail as a long coiled-coil. The extremity of the Long Tail Fibres have been shown to reversibly bind to the polysaccharide moiety of LPS ^75,76^, promoting host recognition, whereas RBP_pb5_ irreversibly binds to an outer membrane transporter, such as FhuA, FepA or BtuB, to trigger infection ^77^.

Salten’s host range was established against the eight SeeT bacterial isolates in liquid culture (96-well plate, Fig. S3). We calculated a virulence index (Vi) to quantitatively evaluate the ability of Salten to infect each bacterial isolate (black crosses in Fig. 3A), defined as follows: Vi∼ 0 indicates that bacterial growth was not affected by the presence of phage while Vi∼ 1 indicates a complete inhibition of bacterial growth in presence of the phage.

**Figure 3.**
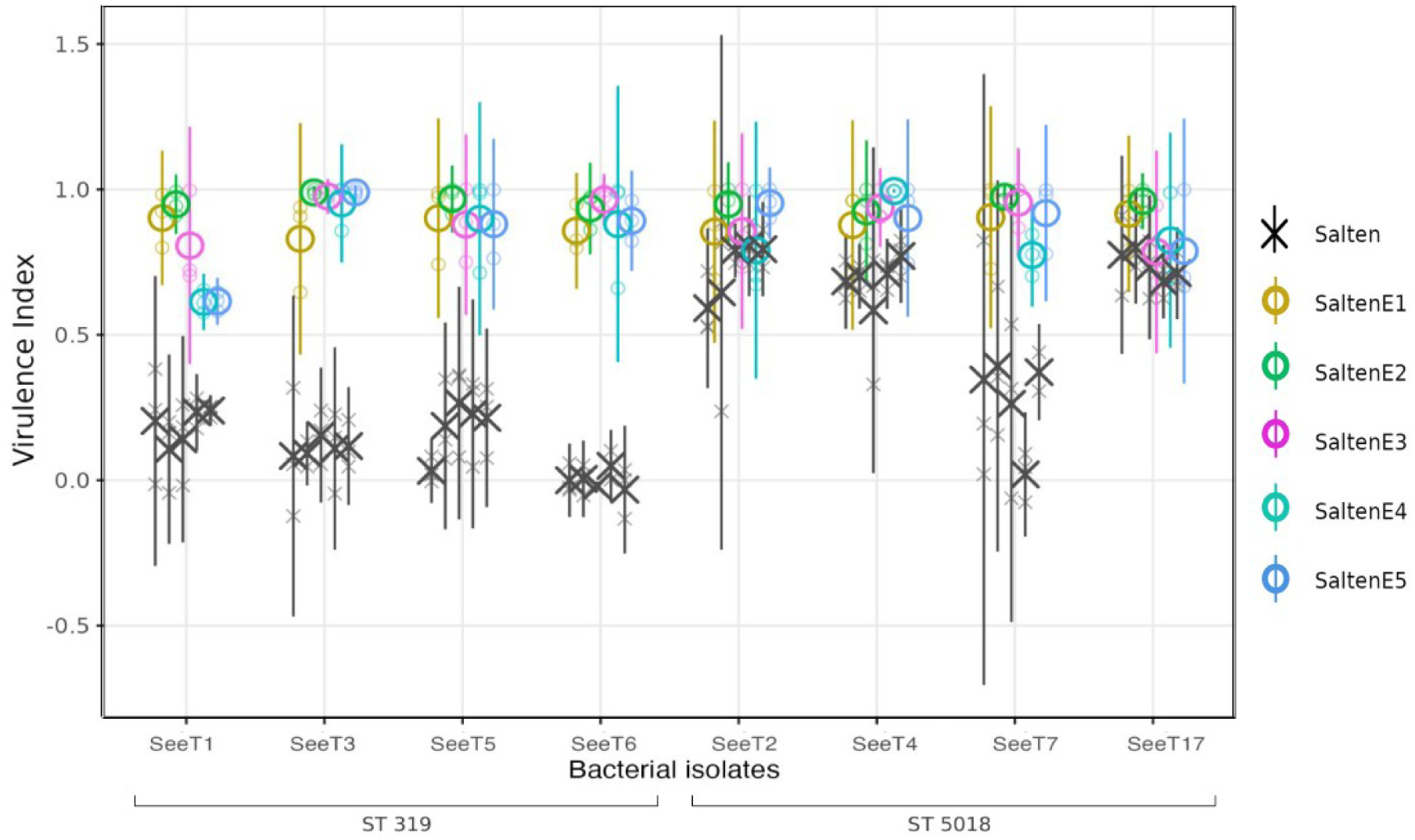
Phenotypic characterisation (host range and virulence) of the ancestral Salten and the evolved SaltenE phage populations on the eight *S. enterica* serotype Tennessee (SeeT) isolates. SeeT isolates are arranged according to their sequence type with ST319 isolates on the left and ST5018 isolates on the right. Phage virulence indexes were evaluated in liquid cultures with MOI = 0.01 at inoculation time, in the presence of ancestral (black cross) or experimentally evolved (coloured circles) phage populations.

We first verified that the factor “plate” (one plate per day) had no effect on Vi estimates (Chisq_14_ = 11.213, p.value = 0.669; type II ANOVA). We then analysed the combined data using a type III ANOVA and concluded that Salten was more virulent on ST5018 isolates than on ST319 isolates (post-hoc comparison by bacteria nested in ST; between ST319 and ST5018, t_98_ = -18.300, p.value < 0.001), with respective marginal means of 0.121 (emmeans package; confidence interval CI 95% [0.083 – 0.158]) and 0.609 (CI 95% [0.571 – 0.647]).

For isolates within each of the ST (ST319 or ST5018), Salten overall had similar virulence on each strain, as virulence indexes were not significantly different between strains (p.value > 0.05; post-hoc pairwise comparison on marginal means between bacteria and by ST). However, SeeT6 (ST319) and SeeT7 (ST5018) were each an exception in their ST, since the ancestral phage Salten was not infectious on SeeT6 (values of virulence index not different from 0, t_112_ = 0.010, p.value = 0.992, post-hoc comparison using Kenward & Roger method with Benjamini & Hochberg correction), which made SeeT6 virulence index significantly different from that of SeeT1 (t_98_ = 3.459, p.value = 0.036) and SeeT5 (t_98_ = 3.468, p.value = 0.035). SeeT7 was unpredictably sensitive or completely resistant across replicates, regardless of whether the experiments were done in liquid or in solid cultures. We thus observed significant differences between SeeT7 and all the other SeeT isolates belonging to ST5018 (SeeT2: t_98_ = 8.379, p.value < 0.001; SeeT4: t_98_ = 7.713, p.value < 0.001; SeeT17: t_98_ = 8.655, p.value < 0.001).

### Phenotypic Characterization of Evolved Phage Populations

Phage host range expansion depends on the ability of phages to reproduce and generate adaptive mutations. We first tried to evolve the Salten ancestral phage under liquid conditions on each bacterial genotype, separately. While Salten was able to inhibit growth of each bacterial genotype belonging to ST5018, we did not observe any effect, even after five serial passages, on the growth of each of the ST319 bacterial isolates (data not shown).

To expand Salten host range, we then applied the Appelmans protocol ^78,37^, a protocol known to enable host-range expansion ^79,80^. This protocol involved repeated growth-selection-mixing cycles on eight non co-evolving SeeT isolates (four ST5018, susceptible isolates; and four ST319, resistant isolates). Experimental evolution was independently repeated five times over a 30-month period. Each experimental evolution replicate lasted six to eight passages (one day per passage).

We then compared the bacterial growth inhibition potential of each of the evolved phage populations (named SaltenE1 to SaltenE5) to that of the Salten ancestral phage (i.e. phage used to initiate each experimental evolution replicate), in 96-well plates. This comparison was done in an absolute manner by adding one or the other phage populations in respective wells of each plate. We repeated each comparison three times, on different days. First, we tested for any potential effects of running plates over several days (one 96-well plate per day) and found no significant effect (linear mixed-effect model with plate as main effect, bacterial identity and phage identity as random factors, Chisq_14_ = 5.175, p.value = 0.983).

The evolved phages displayed an overall higher virulence than the ancestral Salten (respectively emmean = 0.888, CI 95% [0.863 – 0.913] and emmean = 0.365, IC 95% [0.339 – 0.390]; post-hoc contrast between the ancestral phage and the evolved phages, averaged over bacterial isolates; t_210_ = - 30.628, p.value < 0.001; Fig. 3A and Fig. S4). When comparing evolved SaltenE populations (pairwise comparison), each SaltenE population had a similar virulence on each bacterial isolate. Thus SaltenE populations displayed similar values between ST (t_70_ = -0.450, p.value = 0.654; post-hoc comparison over the levels of phages and bacteria, following a type III ANOVA on a dataset keeping only virulence indexes of SaltenE populations and a linear mixed-effect model in which bacterial isolates were nested in ST). Moreover, virulence of SaltenE populations was comparable between bacterial isolates within each ST (p.values between 0.778 and 1.000, post-hoc comparison of phage by ST over some or all levels of bacteria).

The virulence of Salten and SaltenE phage populations were also tested in solid culture using a spot-assay, against an additional 18 isolates of SeeT belonging to either ST319 or ST5018. The ancestral Salten was able to infect 11 out of 26 isolates (i.e. it yielded clear spots in the spot-assay), while the five evolved SaltenE populations were able to infect 25 out of 26 of the bacterial isolates. The SaltenE populations induced only turbid spots on SeeT18. After sending this isolate to the reference center CNR (Paris, France), it was attributed by MLST7 to another serotype and sequence type, Mbandaka ST413 (Fig. S7).

### Genomic Characterization of Evolved Phage Populations

To elucidate which exact mutations underlie host range expansion and increased virulence of the evolved populations, we sequenced with Illumina technology the genome of both the ancestral Salten and those of the four evolved populations (SaltenE1, E2, E3 and E4). We thus confirmed first that the observed SaltenE genotypes were derived from the ancestral Salten genotype, in order to rule out contamination or prophage excision from the bacterial hosts ^81^. Between 95.7% and 97.5% of SaltenE reads mapped against the ancestral Salten genome, after alignment using bwamem2 v.2.2.1 ^83^, confirming that the vast majority of the evolved phage sequences were descended from the ancestral Salten sequence.

We then compared the prevalence of mutations detected along the sequenced genotypes for each of the four SaltenE populations (Fig. S6) by performing a variant calling using Breseq (see Table S2 for breseq output). Overall, 2% of mutated positions corresponded to indels and 98% to SNPs, with a hot spot of mutations detected in ORFs upstream or within the coding structural region (in between positions 75 kb to 100 kb). Identical mutations detected at the same *loci* and shared by two, three or four SaltenE populations (hereafter referred to as parallel mutations) were identified in structural and non-structural genes (Table 1). Notably, prevalent parallel mutations were detected in genes coding for DNA modification, recombination, repair and replication enzymes. Interestingly, both SaltenE2 and SaltenE4 harboured similar mutations in the phage-associated recombinase (Table 1 and Table S2), with a within-population frequency exceeding 50%. Additionally, several parallel mutations accumulated in the ORFs encoding tail-structure proteins, which are known to influence host range (detailed in Fig. S6, non-synonymous mutations detailed in Fig. 5A).

**Table 1.** Number of similar mutations shared by several evolved phage populations (SaltenE).

| Gene | ORF length | Number of parallel mutations | Non-synonymous mutations | Number of lineages sharing these mutations |
| --- | --- | --- | --- | --- |
| Gene coding for a ATP-dependent helicase (ortholog to uvsW protein) | 1353 | 55 | 6 | 2 |
| Gene coding for an ortholog to gp32 of phage T4 | 774 | 21 | 4 | 2 |
| Gene coding for a phage-associated recombinase | 978 | 59 | 17 | 2 |
| Gene coding for a phage recombination related exonuclease | 1839 | 61 | 11 | 2 |
|  |  | 45 | 5 | 3 |
| Gene coding for an ortholog to a type II restriction endonuclease | 483 | 28 | 7 | 3 |
|  |  | 13 | 6 | 2 |
| Gene coding for endonuclease | 876 | 62 | 5 | 3 |
|  |  | 5 | 0 | 2 |
| Gene coding for deoxyuridine 5'triphosphate nucleotidohydrolase (dUTPase) | 447 | 27 | 3 | 4 |
|  |  | 7 | 1 del / 1 insert | 3 |
| *pb6* (Tail Tube Protein) | 1392 | 119 | 21 | 3 |
| *pb4* (Central Fibre Protein) | 2103 | 92 | 14 | 4 |
| *pb3* (Baseplate Hub Protein) | 2850 | 107 | 22 | 4 |
|  | 2850 | 21 | 0 | 3 |
| *p138* | 369 | 20 | 1 | 3 |
| *p139* | 405 | 27 | 1 | 3 |
| *p140* (Baseplate Tail Tube Protein) | 897 | 38 | 16 | 3 |
| *p132* (Collar Protein) | 423 | 11 | 4 | 4 |
| *pb2* (Tape Measure Protein) | 3708 | 101 | 11 | 4 |
|  |  | 89 | 27 | 3 |
| *pb1* (lateral Long Tail Fibre) | 4152 | 2 | 2 | 4 |

By analogy to the published T5 tail structure ^61^, we determined the positions of the parallel mutations on the protein structure (Fig. 5C,E). Synonymous substitutions are expected to induce no or little change in substrate binding specificity, so we only took into account from then non-synonymous substitutions with dissimilar or weakly similar properties.

At the protein level of pb6, the mutation frequency (i.e. the number of non-synonymous mutations detected over the total number of amino-acids concerned) in the tube domain was 3.3%, whereas it went up to 20.7% in the Ig-like domain. Mutations in the Ig-like domain were mainly located in the variable loops, known to be responsible for ligand binding (Fig. 5C,D).

Mutations in *pb4* ORF were present at high frequencies in all of the SaltenE populations (Fig. S6; above 50% within SaltenE1 and SaltenE2, and between 25% and 75% within SaltenE3 and SaltenE4). Interestingly, non-synonymous substitutions in the pb4 protein were mostly concentrated in the FNIII domains and in the decorative domains of the spike (Fig. 5C,D), domains that are also suggested to contribute to host binding ^61^.

For the *pb9* ORF, read coverage (evolved reads coming from each of the four evolved phage lineages and respectively aligned against the ancestral genome) was <10x and therefore too low to conclusively evaluate the frequencies of mutations.

Only 4% of all the caudal protein mutations were in p140 protein, which is consistent with the fact that it is completely surrounded by the p132 collar and therefore does not interact with the host surface.

Mutations in p132 protein were also concentrated in its Ig-like domain (12.1% of the total mutations in this gene), and, as for pb6, mutations were mainly located in loops, which are known to interact with specific ligands (Fig. 5C,D).

The gene in which we observed the highest numbers of non-synonymous parallel mutations was *pb2.* The impact of such non-synonymous mutations on the structure of the protein pb2 is difficult to predict because the pb2 protein structure has not been completely solved, and residues 420 to 891 appear to be highly divergent among known T5-pb2 proteins available on NCBI database. However, when compared to other ORFs encoding caudal proteins, the *pb2* ORF harboured one of the highest variability among our evolved SaltenE nucleotide sequences, with 101 parallel mutations shared by all four SaltenE lineages and 89 more shared by three of the SaltenE lineages (Fig. 4A).

**Figure 4.**
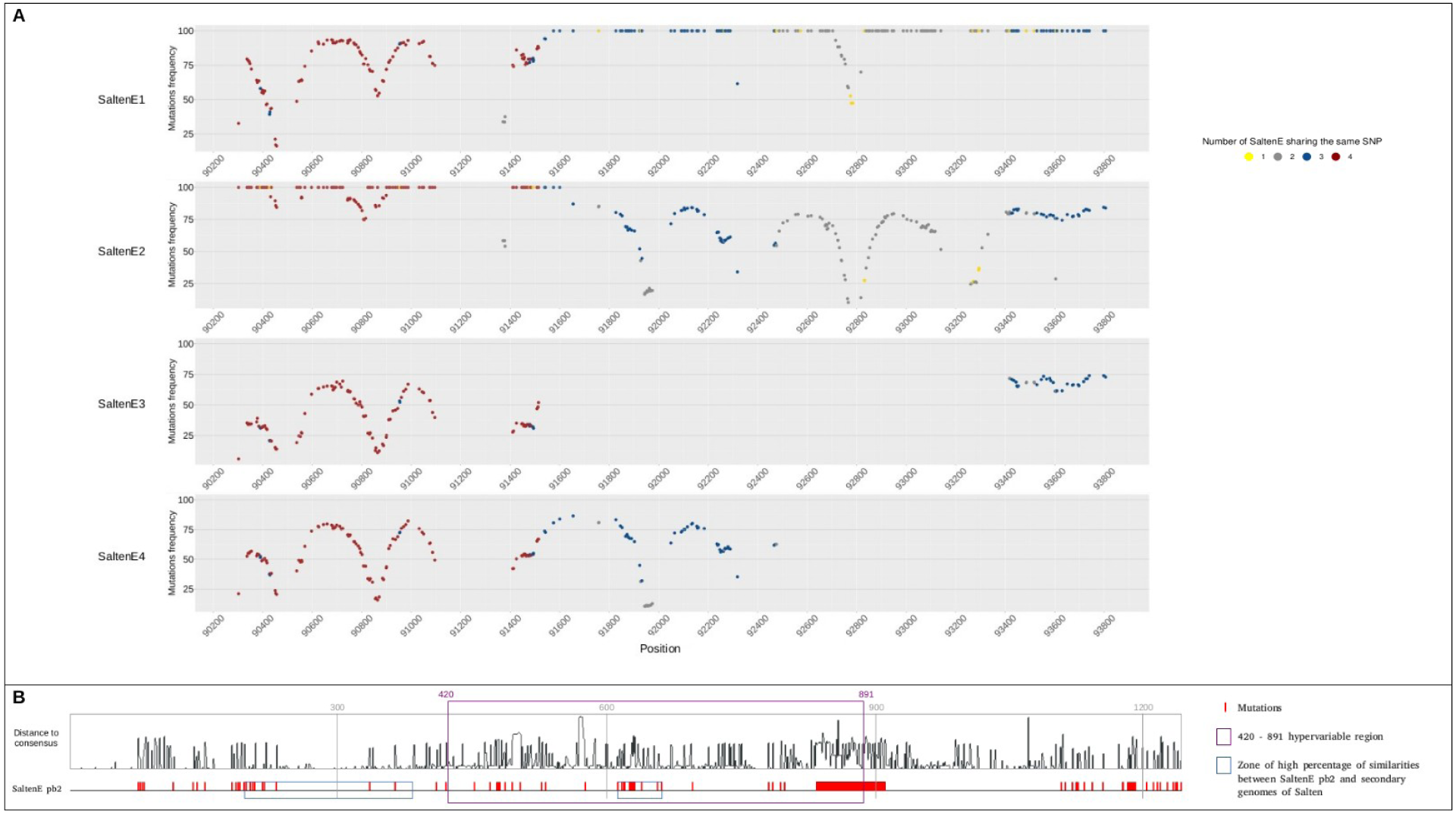
Frequency of mutations in Salten *pb2* ORF. **A** Frequency of mutations accumulated in *pb2* ORFs in four independent evolved populations (SaltenE1, E2, E3, E4). colours of each mutation correspond to its presence over populations: yellow mutations are present in only one evolved population, gray in two, blue in three and dark red in four evolved phage populations. **B** Standing diversity among 100 pb2 proteins from NCBI compared to Salten. The black histogram represents the distance to Salten consensus. The lower red panel represents positions at which we observed mutations in three or four SaltenE evolved populations (pb2 consensus protein sequences) compared to ancestral Salten pb2 protein. The dark purple rectangle bounds the 420 - 891 hypervariable region. Blue rectangles represent SaltenE pb2 regions with higher degrees of sequence identity to minority small contigs (assembled following ancestral Salten population sequencing) than to the consensus Salten sequence.

Intrigued by these multiple parallel mutations in *pb2* ORF, we looked at their order of appearance by Sanger sequencing PCR amplicons at passages one, three and five of the experimental evolution procedure. Most of the mutations appeared to become fixed in the evolved populations between passages four and five in SaltenE1 and SaltenE2, and passages six and seven in SaltenE3 and SaltenE4. Strikingly, we observed small differences in mutational profiles according to each evolved population, such as mutations appearing fixed at passage five within SaltenE3 whereas secondary peaks were observed at this *locus* in SaltenE1, E2 and E4, showing a transient co-existence of alleles over time rising to fixation.

In order to verify whether such mutations were part of the standing genetic variation or of the ancestral Salten population at the onset of the experiment, or whether they had arisen as *de novo* mutations, we analysed the Salten ancestral population sequencing data. We especially focused on the small contigs (< 1,200bp) present within Illumina outputs of Salten genome assemblies. Originally, these small contigs were removed because of their very low coverage (less than five reads of coverage) which suggested that they were artifacts or contaminants. We aligned these small minor contigs with the highly mutated regions present in evolved populations, so as to determine whether, under selection, the mutations observed in these contigs might have substantially contributed to evolved SaltenE populations. For two different regions (base pairs 553 to 1116 and 1794 to 2026) of the *pb*2 ORF, we found >99% identity of some of these minor contigs with the SaltenE population *pb2* consensus nucleotide sequences but only 85% identity with Salten consensus ancestral *pb2* sequence (Fig. S8). In order to verify whether the observed accumulation of mutations within the evolved consensus sequence of pb2 protein tended to occur in “hypervariable” zones, we aligned 100 sequences of pb2 proteins retrieved from NCBI GenBank as well as the Salten and SaltenE pb2 consensus sequences. Consistent with our hypothesis, the multiple parallel mutations retrieved between consensus pb2 protein sequences from the four evolved SaltenE populations and from the ancestral Salten population (Fig. 4B; red rectangles) were located in hypervariable zones, including the 420 - 891 region mentioned above which seemed to be hypervariable (Fig. 4B; dark purple rectangle). The two regions matching with minor contigs obtained by sequencing the ancestral Salten population were also located in these hypervariable zones on the protein (Fig. 4B; blue rectangles).

Notably, minor contigs present in the initial Salten inoculum showed >98% identity with other regions present in the evolved tail structural protein encoding sequences. For instance, the evolved *p132* ORF consensus sequence from the SaltenE populations presented 100% identity with a minor Salten contig, but only 73% with Salten consensus *p132* ORF (Fig. S8). We also found three regions in *pb3* evolved ORF consensus sequence (base pairs 1 to 854, 1272 to 1746 and 2253 to 2701) presenting 99% identity with some minor Salten contigs, but only 76% with Salten consensus *pb3* ORF (Fig. S8). Two regions on *pb6* evolved ORF consensus sequence (base pairs 462 to 761 and 1277 to 1407) presented >98% identity with minor Salten contigs *versus* 81% and 76% identity with Salten consensus *pb6* ORF, respectively (Fig. S8). Finally, region 205 - 957 bp of *pb4* evolved ORF consensus sequence presented >99% identity with minor Salten contigs *versus* 74% identity with Salten consensus *pb4* ORF sequence (Fig. S8). These results suggested that some phage regions are hypervariable and one or another sequence might be selected and increase in frequency over time (and serial passages) according to selection pressures. Our results also suggest that even after five first severe bottlenecks meant to purify a phage isolated from the environment and obtain a homogeneous solution of viral particles, within-population variation still remains or can be quickly generated at least at some *loci* that can be qualified as hypervariable.

#### Focus on Variation within the pb1 Gene (Long Tail Fibre)

The last considered gene involved in caudal structure is *pb1*. After experimental evolution, we detected only two parallel mutations, which became fixed in all four evolved SaltenE populations: C80453T (alanine into valine on the protein; A1157V) and A80495G (asparagine into aspartic acid on the protein; N1178D). They were also fixed in the four evolved SaltenE populations (Fig. 5F), despite the varying coverage across replicates (SaltenE3 and E4: ∼120X and ∼200X of coverage, respectively; SaltenE1 and E2: <6X coverage around this particular locus position). To evaluate fixation dynamics of both mutations, we Sanger-sequenced PCR amplicons (obtained through primers flanking positions 80453 and 80495 of the Salten genome) at passages one, three and five of our experimental evolution. Concerning the four evolved populations, the A80495G parallel mutation was fixed at passage three and the C80453T parallel mutation was fixed at the fifth passage of Salten’s experimental evolution.

**Figure 5.**
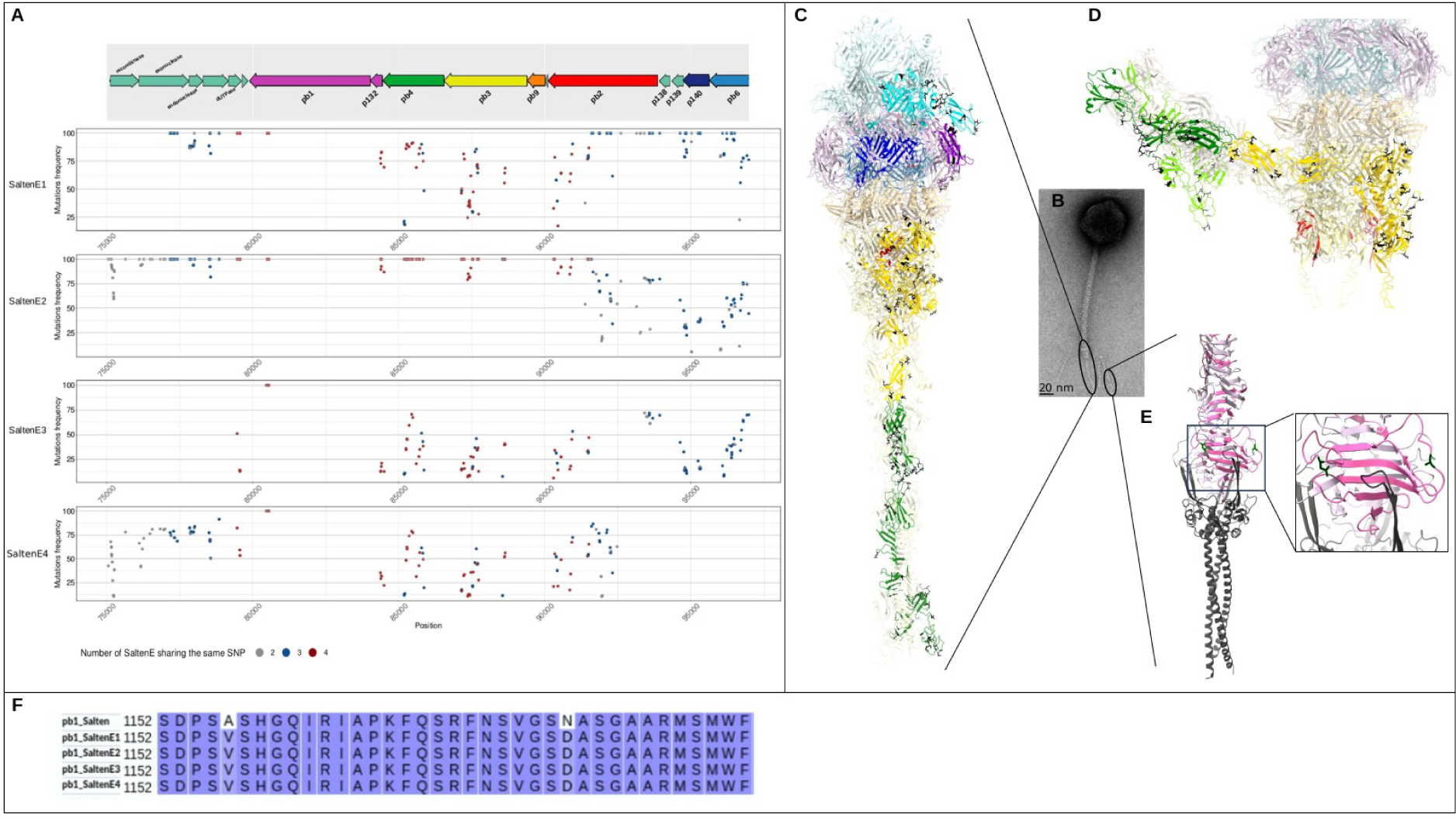
Non-synonymous parallel mutations present in caudal genes of evolved phage populations SaltenE1, E2, E3, E4. **A** Frequency of non-synonymous parallel mutations according to their genomic position. Degree of parallelism of each mutation is indicated with colours as in Fig. 4A. **B** Negative stain micrograph of phage T5. **C,D,E** Mutations (indicated in black) on the different proteins contributing to the T5 tail tip structure: pb6: cyan; p140: blue; p132: purple; pb9: orange; pb3: gold; pb4: green (note that the three FNIII domains and the spike are in different shades of green); pb1: pink (note the chaperone domain in dark grey). **C** Ribbon representation of T5 tail tip structure. **D** Ribbon representation of T5 tail tip structure when injecting phage dsDNA after being adsorbed on the bacterial membrane. **E** Ribbon representation of the C-terminal tip of pb1 with its chaperone domain (4UW8 ^75^). **F** Alignment of Salten and the four SaltenE of pb1 amino acids 1152 to 1216, corresponding to the C-terminal part of pb1.

Interestingly, phage populations undergoing experimental evolution were able to infect the eight bacterial isolates from the third passage. Even though the growth of the ST319 bacterial genotypes was not yet entirely inhibited at this passage, it was almost completely inhibited by at the fifth passage.

The C-terminal domain of T5-pb1 protein is known to point towards the curved groove at the subunit interface, a region suggested to be the poly-mannose binding site of the protein and a chaperone domain ^75^. Because the full-length Salten-pb1 was not easily alignable to T5-pb1, the structure of its C-terminal domains was predicted with AlphaFold2 ^82^ with very high confidence (plDDT higher than 90%; Fig. S9). The predicted structure had a C-terminal chaperone domain with a long loop, which, similarly to T5-pb1, could serve as the poly-mannose binding site. Most pertinent in the context of the evolution experiment, the two parallel C80453T and A80495G mutations resulted in two amino acid changes identified within this predicted binding groove (Fig. 5E).

#### Experimental Evaluation of pb1 Point-Mutations on the Phenotype

Given the importance of the *pb1* ORF in host range specificity ^84,75^, we aimed at experimentally testing the adaptive potential of the two parallel mutations. To do so, we carried out a reverse-genetics experiment by taking advantage of the naturally high recombination rate of Tequintaviruses to acquire the two parallel mutations previously introduced within a plasmid ^85^. We thus created three plasmids that we introduced into the most susceptible bacterial isolate, SeeT17, either without any phage sequence (hereafter called “rEV” for “reverse empty vector”), or with the ancestral phage region encompassing positions C80453 and A80495 (hereafter called “rA” as “reverse ancestral”), or with the evolved sequence region encompassing T80453 and G80495 (hereafter called “rE” as “reverse evolved”). After infection of each of the three transformed SeeT17 with the ancestral phage Salten population, we recovered the phage progeny. PCR amplicons targeting the appropriate region in *pb1* ORF were Sanger-sequenced in order to confirm the presence of evolved *pb1* mutations in phages having recombined with rE plasmid (hereafter called rE_Salten) or rEV or rA plasmids (hereafter called rEV_Salten and rA_Salten respectively). We then tested their phenotypes on ST319 bacterial lawn. As expected, ancestral Salten population as well as rEV_Salten and rA_Salten were not able to create clear lysis plaques on ST319 isolates (SeeT1, SeeT3, SeeT5 and SeeT6; Fig. 6A). The inability or highly reduced ability of these phages to infect SeeT3, SeeT5 and SeeT6 was obvious (no plaque observed or light turbid plaques) while Salten, rEV_Salten and rA_Salten induced some turbid lysis plaques on SeeT1 lawns. As expected, Salten, rEV_Salten and rA_Salten were also able to create clear lysis on SeeT17, the positive control from ST5018. Confirming our prediction, rE_Salten produced clear lysis plaques on all lawns produced by ST319 isolates, in addition to SeeT17 bacterial lawn (Fig. 6A last column).

**Figure 6.**
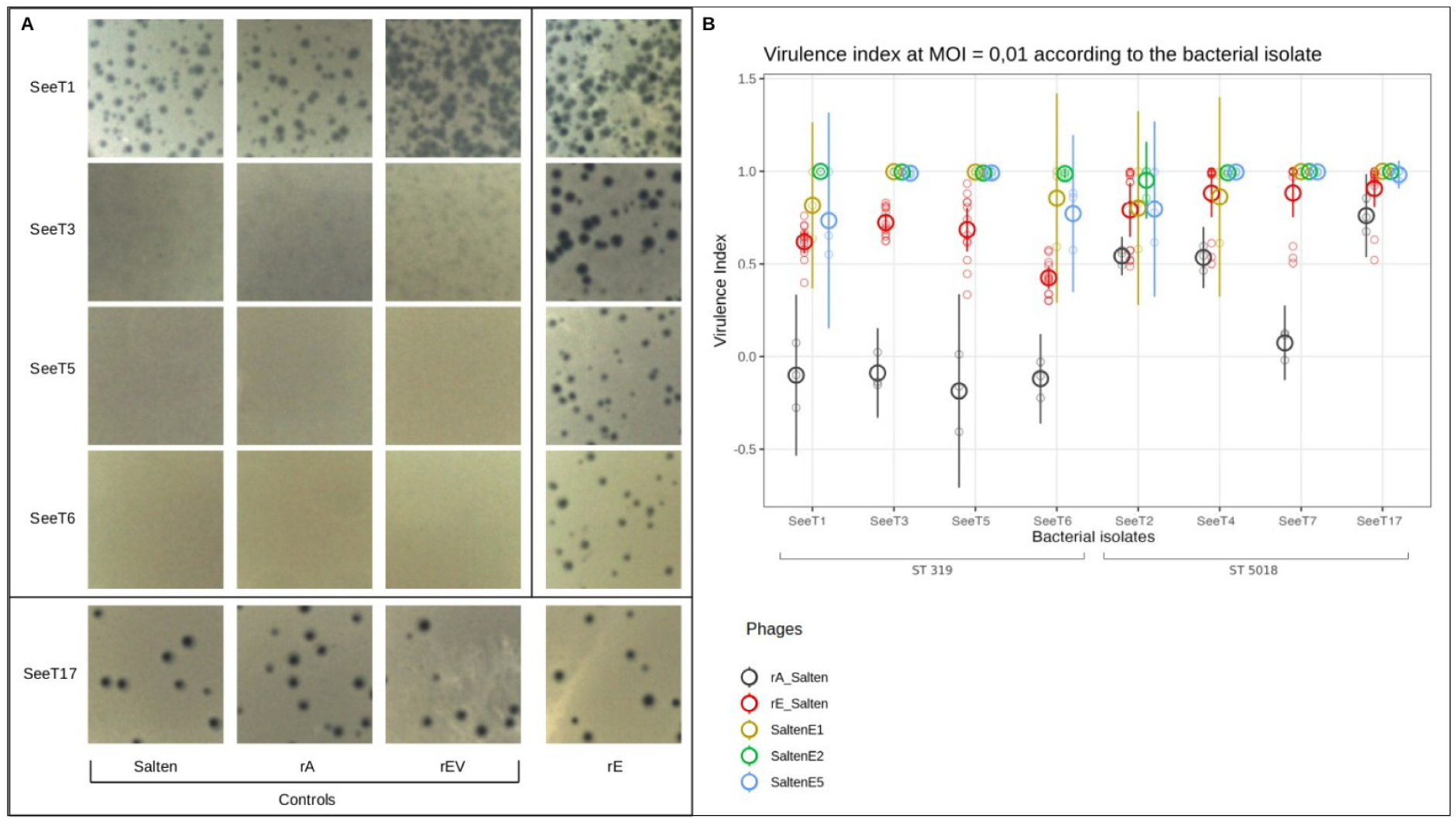
host range and virulence evaluation of rE_Salten. **A** Visualisation of the reverse-genetics double-layer assay. Bacteria are represented in rows and phages in columns. SeeT17 served as positive control as it is susceptible to the ancestral Salten phage infection. rEV_Salten, rA_Salten and rE_Salten correspond to the ancestral phage having potentially recombined with a plasmid harbouring, respectively, no insert (no recombination allowed), an insert with the ancestral sequence of the *pb1* gene, and finally the insert with the evolved sequence of the *pb1* gene. **B** Virulence index (Vi) of rA_Salten (dark grey), rE_Salten (red), SaltenE1 (yellow), SaltenE2 (light green) and SaltenE5 (dark blue), according to *S. enterica* isolates and ST, at MOI = 0.01.

In order to evaluate whether the two parallel C80453T and A80495G mutations also enhanced infection in liquid cultures, we compared the growth curves of the eight bacterial isolates in presence of either rA_Salten or rE_Salten, in 96-well plates (Fig. 6B). Confirming the data obtained in solid conditions, on the one hand, rA_Salten (emmean = -0.123, CI 95% [-0.210 -0.036]) was significantly less capable than rE_Salten (emmean = 0.613, CI 95% [0.556 – 0.671]) to inhibit growth of ST319 bacterial isolates (post-hoc contrast by pairs of phages; t_164_ = -16.088, p.value < 0.001). But rA_Salten was also significantly less capable than rE_Salten to inhibit bacterial growth of two isolates from ST5018 (post-hoc comparison by pairs of phage by bacteria; SeeT4: t_164_ = -3.786, p.value = 0.038 and SeeT7: t_164_ = -8.833, p.value < 0.001). Overall, rE_Salten showed an expanded host range as well as a higher degree of virulence than rA_Salten.

To evaluate whether the two parallel mutations in *pb1* ORF alone were sufficient to fully account for the increased virulence of the SaltenE populations, we directly compared the capacity of rE_Salten and SaltenE populations to inhibit bacterial growth in 96-well plates. On ST319 bacterial isolates, SaltenE populations were significantly more virulent than rE_Salten (post-hoc contrast by ST; t_164_ = 10.034, p.value < 0.001; emmean = 0.850, CI 95% [0.752 - 0.948] and 0.620, CI 95% [0.533 - 0.706], respectively) except for SeeT1 in which SaltenE populations were marginally more virulent than rE_Salten (post-hoc contrast by bacteria; t_164_ = 3.683, p.value = 0.053). On ST5018 bacterial isolates, SaltenE populations were also significantly more virulent (but to a lesser degree than on ST318) than rE_Salten (t_164_ = 2.624, p.value = 0.026).

## Discussion

The question of whether generalism (i.e. the ability to successfully reproduce in different environments) evolves at the cost of a lower mean fitness across environments remains an open fundamental question with applied dimensions ^17,9^. In the field of bacterial control using phages, there is no consensus yet favouring the use of one generalist phage or a cocktail of several specialist phages ^86,87^. On the one hand, if generalist phages replicate less efficiently than specialist ones, resistant bacterial genotypes might appear with higher probability thanks to bigger bacterial population sizes ^88,89,90,91^. On the other hand, using several specialist phages could modify phage-bacteria networks due to phage-phage (antagonist) interactions, such as agglutination among phages resulting in lower effective phage concentrations ^92,93^.

If possible, it would then be of great interest to generate generalist phages adapted to a broad host range with high virulence. One aim of our study was to generate such a highly efficient generalist phage by adapting it to several bacterial genotypes. Viral adaptation to hosts resides in optimizing the several life cycle steps of adsorption and entry on the target cell, bypassing of cell defense, followed by gene expression, genome replication, virion assembly and release ^94^. In phages, the determinants of host range often consist of proteins allowing for adsorption on the bacterial surface, such as capsid or spike proteins in microvirid phages ^95,96,24,97^, receptor binding proteins in the RNA virus Φ6 ^25,26^ or surface polysaccharide-related traits (K-serotype, LPS outer core or O-antigen serotype) in *Caudoviricetes* ^98^ such as observed in fibres of phages T3 ^99^, T4 ^100^, T7 ^9^, T2 ^101^ and T5 ^75^. Our study showed that in the *Tequintavirus* Salten, parallel mutations (revealing convergent evolution) notably accumulated in *pb1* ORF, coding for the Long Tail Fibre, a caudal protein involved in host recognition through LPS interaction. Interestingly, we demonstrated (by reverse-genetics) the adaptive function of these parallel mutations and showed that they appeared early in our experimental evolution assay, confirming that host entry is one of the primordial steps of adaptation.

Specifically, we showed that mutations in the C-terminal region of the pb1 protein were necessary for host-recognition. We did not observe any other adaptive mutations in the rest of the protein. Although, these results are not consistent with the finding that for BD13 (another recent study on a *Tequintavirus*), the domain responsible for host interaction is in the N-terminal region of pb1 protein ^84^. This discrepancy between results might be attributable to the predicted BD13-pb1 protein (as determined by AlphaFold2) which harboured a longer coiled-coil domain (connecting the fibre to the tail tube) and a shorter fibre domain than the Salten-pb1 protein. The predicted fibre structure of the Salten-pb1 protein aligned well with the central domain of the BD13-pb1 protein (DALI ^102^ z-score of 9.6 over 302 residues), but the alignment also revealed that the BD13-pb1 protein lacked the C-terminal saccharide binding domain found in Salten (Fig. S9).

Besides those in pb1, multiple parallel mutations were observed in evolved phage populations at variable within-population frequencies in other phage caudal proteins: pb6, p140, pb3, pb4 and p132. These structural tail proteins present accessory/decoration or structural domains with protein or oligosaccharide binding folds. These features have been proposed to increase phage infectivity through specific binding to protein and/or saccharides at the surface of bacteria (e.g. Ig-like and FNII domains, or Oligosaccharide-binding domain ^103^). Notably, a large proportion of the parallel mutations observed in our study were located in these decorative domains, and more specifically in the loops responsible for binding with bacterial ligands. Such great variability of phage proteins recognizing LPS as a first step of infection is thus in line with the great variability of *Salmonella* O-antigens (more than 2500 registered serotypes). Indeed, we showed on the one hand that host range adaptive mutations accumulated on long tail fibre involved in recognizing LPS and O-Ag. On the other hand, the polymorphisms in genes involved in O-Ag processing were only present at three codon sites in two genes (*waaK* and/or *wzxC*). Further studies will be needed to determine (i) on the phage side, if the mutations detected in *pb1* ORF are sufficient to expand the phage host range to other serotypes and sequence types harbouring the same profiles on these two genes; and (ii) on the bacterial side, if mutants resistant to Salten will involve genetic variations (mutations / deletions) in the *loci* present in genes that encoded waaK and/or wzxC or other regions as it has already been described in other gram-negative bacteria ^104,105^.

Interestingly, we did not observe any parallel mutation in phage receptor binding protein pb5, which is known to irreversibly bind the outer-membrane transporter FhuA ^61^. When looking carefully at bacterial genomic polymorphisms in the two sequence types involved in our study (ST319 and ST5018), we noted that FhuA differed between our ST5018 and ST319 isolates by only two amino acids: one located in a periplasmic loop and the other in the plug (Fig. S1C). Neither of these amino acid sites are in extracellular loops that mediate the interaction with the phage RBP ^77^. Therefore, it was with no surprise that we did not observe any adaptive mutations accumulating within the phage RBP gene.

One striking result of our study is the high number of mutations observed in caudal and non-structural proteins. Interestingly, similar patterns have been reported in other experimental evolution studies, such as in phage T7 ^106^, where hundreds of mutations – some of them parallel – were detected. Several factors may explain this high mutational diversity. First, as observed in the gene coding for pb2, some parallel mutations might have been present at the beginning of the experiment but at such low frequencies that they went undetected. Their subsequent increase across serial passages could result from direct selection due to fitness advantages or indirect selection via hitchhiking. These hyper-variable regions could arise spontaneously, particularly in proteins with immunoglobulin-like structures as is frequently observed in tailed ds-DNA *Caudoviridetes* genomes ^107,108^. Second, the abundance of mutations may reflect the selection of mutator genotypes, a phenomenon suggested to be transiently adaptive in fluctuating environments ^109,110,111^. Supporting this idea, some of the parallel mutations in evolved Salten occur in genes encoding endonucleases and recombination exonucleases, proteins known to modulate mutation rate, as shown in myophage T4 ^109^, podophage T7 ^112^ and SARS-CoV-2 ^113^. Moreover, Salten’s recombinase and exonuclease share structural similarity with the Mre11 – Rad50 complex in eukaryotes and its bacterial and phage orthologs, including SbcCD (prokaryotes), gp46/gp47 (phage T4) and gpD13/gpD12 (phage T5) ^114^. Structural modelling of SaltenE parallel mutations further places them near the DNA processing site ^115^, which supports our hypothesis that these mutations may drive the selection of mutators. Further studies will be needed to directly test the effects of these parallel mutations in the SaltenE endo- and exonuclease on mutation rates and transient adaptation.

Other parallel mutations were observed within phage dUTPase, a protein primarily responsible for removing excess of dUTP mistakenly incorporated into DNA by DNA polymerases. Mutations in this protein may be adaptive because it is absolutely vital for the phage to supplement *E. coli* dUTPase activity which is, on its own, insufficient to exclude uracil from progeny DNA ^116^. dUTPase seems also to be involved in mutation rate variation ^117^ and in potentially additional functions that have not yet been characterised ^118^.

Further, we observed parallel mutations within proteins involved in recombination-dependent replication (RDR) and DNA-repair, including an ortholog of the T4 ATP-dependent helicase uvsW. These proteins are key contributors to phage counter-defense systems. Notably, phage mutants in RDR and helicase genes have been suggested by Loeff et al. (2023) ^119^ to contribute to the evasion of the bacterial defense system Shedu, a protein complex targeting free-end ssDNA and detected in our bacteria. Moreover, Wu et al. (2021)^120^ found that the phage T4’s recombinase, uvsX, is essential to escape CRISPR bacterial defenses by removing the targeted nucleotides. To deepen our molecular understanding of both phage adaptation to bacterial hosts, and bacterial resistance mechanisms against phages, further studies will be necessary to experimentally test these hypotheses.

Finally, one of the most promising results of our study resides in our ability to rapidly evolve a phage with both an expanded host range and a significantly increased degree of virulence. Indeed, the evolved phage populations inhibited bacterial growth with significantly more efficiency than the ancestral phage on both the ST319 and ST5018 host genotypes (Fig. 3). We subsequently showed that adaptive mutations present in the *pb1* gene were not sufficient on their own to explain the significant increase in virulence (Fig. 6B). We then suggested that increased virulence was due to mutations allowing evolved phages to escape from the bacterial host defense system.

It is important to point out that our results are inconsistent with those of other experimental evolution studies. We suggest three possible explanations for this. First, contrary to other studies, we evolved Salten in a spatially variable environment thanks to the Appelmans protocol ^37^, while other experimental evolution studies have evolved their phages in temporally variable environments. More explicitly, we argue that the nature of adaptive mutations and their interactions may vary depending on the environment in which individuals evolve (temporal *versus* spatial variable environments). In cases where individuals alternate between different environments (i.e. temporal variability), all individuals experience simultaneously the same conditions (peaks or valleys in their fitness landscape). Each genotype can thus accumulate mutations with deleterious effects in one or the other environment because all competing individuals experience these costs simultaneously, maintaining comparable relative fitness. Such a process might explain why alternation between *S. enterica* and *E. coli* was accompanied by a reduced ability of ΦX174 to infect *E. coli* ^24^. On the contrary, in cases where each individual of a population is confronted with one or the other of two different environments (spatial variability), all individuals experience different conditions at a given time. Natural selection might then select for mutations with the lowest selective costs, due to antagonistic pleiotropy imposed by adaptation to one or the other environment. Moreover, natural selection might also select for other mutations compensating the selective cost(s) imposed by adaptive mutations conferring a selective advantage that is exclusive to one or the other environment. In this regard, the Appelmans protocol creates a spatially variable environment. Each well of a 96-well plate consists of a single sub-population of phage-bacteria with a unique condition, due to different population sizes, multiplicities of infection (MOI) as well as bacterial genotype present in each well (one genotype out of eight in our conditions). At each transfer, all phages are thus mixed and randomly dispersed into new sub-populations, each phage competing with other genotypes that previously experienced different conditions.

Second, and contrary to other experimental evolution assays such as that described in Sant’s study ^9^, our ability to rapidly select for a phage with generalist behavior and high virulence might also come from another peculiarity of the Appelmans protocol ^37^, in which some sub-populations are at high MOI while other are at low MOI. Thus, in wells with high MOI, both large population sizes and population processes, such as amphimixy or complementation, likely promote the retention of high mutational diversity within phage populations ^121,36^. Conversely, in solutions with a low MOI, strong selection favors mutants with an expanded host range and those carrying compensatory mutations, eliminating the possibility of phenotypic masking of deleterious mutations ^122,123^.

Third, we evolved Salten through several replication cycles (six to seven serial transfers with approximately 18 hours of bacterial growth at each passage), whereas other experiments evaluated virulence after as few generations as possible, without letting compensatory mutations accumulate. For example, Duffy et al., (2006) ^25^ and Ferris et al., (2007) ^26^ were interested in measuring the fitness cost induced by host range mutations. They thus measured Φ6 relative fitness after just one replication cycle. If our hypothesis holds, we would expect these expanded host range Φ6 mutants to accumulate compensatory mutations if they were evolved for longer in spatially variable environments composed of different bacterial hosts.

## Conclusion

We have isolated a novel species of *Tequintavirus* from an environment sample. Initially, this phage was able to infect only one *Salmonella enterica* sequence type. After seven serial experimental evolution passages exposing the phage to different hosts at different MOI, a phage population was evolved displaying an expanded host range and increased virulence towards new *Salmonella enterica* sequence types. Our results highlight the power of the Appelmans experimental evolution protocol to exploit the high evolvability potential of phages.

We include in our results a warning to supporters of synthetic biology ^124,125^ who suggest the generation of synthetic phages with expanded host ranges via the modification of genes involved in host-recognition ^99^. Specifically, we have shown that host range mutations are not sufficient to confer efficient inhibition of bacterial growth. In fact, other mutations seem to be needed to increase virulence, such as those subverting host antiviral defense mechanisms, which are advantageous to the phage. A situation where engineered phages would have expanded host range but low virulence may encourage the evolution of new bacterial genotypes that are resistant to the phages.

Finally, the question of why we do not isolate more generalist phages in natural conditions still remains. To answer this question, we would like to highlight the impact of environmental heterogeneity on competition among phage genotypes. Under our experimental conditions, phages were pooled within the same spatially variable environment at each passage despite high degrees of hosts diversity (eight bacterial genotypes belonging to two sequence types). In natural conditions, bacterial host populations are mostly segregated into separate patches in which competition happens and selective benefit to a generalist phage may be enhanced or reduced depending on the frequency of one or the other permissive host. Generalist phages may then be favored when the hosts occur only in mixed patches (with specialized phages being eliminated when environments are composed of separated patches of host genotypes), as is predicted by analytical and simulation models based on optimal foraging theory ^126,127^.

## Supporting information

Table S2. Breseq output.

## Author contribution

Conceptualisation, R.F. and A.M.; Methodology & Investigation, R.F. and A.M. with the help of C.Z.-C. and M.M.-R. for the reverse-genetics part and F.-X.W for bacterial sequencing; Results analysis, R.F., A.M., M.V., C.Z.-C., C.B., S.B., J.H., J.D., F.-X.W. and I.B; Writing, R.F. and A.M.; Revision, R.F., A.M., M.V., C.Z.-C., C.B., S.B., J.H., J.D., F.-X.W., I.B. and A.F.

## Acknowledgments

We thank M. Ansaldi (LCB, Marseille) for providing wastewater from which we isolated our phage, M-S. Vernerey (PHIM, Montpellier) for the electron microscopy, C. Mariac (DIADE, Montpellier) for the library quality check before sequencing, A. Talman (MIVEGEC, Montpellier) for the help on the iSeq use, O. Rossier (I2BC, Montpellier) for helpful discussion on reverse-genetics, C. Mariac (PHIM, Montpellier) for QiAxcel use, P. Agnew for his help on statistics, Y. Anciaux (ISEM, Montpellier) for the Vi calculation, J. Garneau for PhageTerm help, M. Monot (I. Pasteur, Paris) and A. Dereeper (PHIM, Montpellier) for bioinformatics advice, C. Torres-Barceló (PHIM, Montpellier), C. Whittington (CED, Bordeaux) and D. Martin (UCT, South Africa) for helpful commentaries and discussion of the manuscript. The authors thanks UMR MIVEGEC for providing an efficient working environment and acknowledge the ISO 9001 certified IRD itrop HPC (member of the South Green Platform) and the whole bioinformatics team support in Montpellier for providing HPC resources that have contributed to the research results reported within this paper. URL: https://bioinfo.ird.fr/ - http://www.southgreen.fr This work was funded by Royal Canin through the support of the CNRS (contract 245420).

## DATA AND SCRIPT AVAILABILITY

The data and scripts that support the findings of this study are openly available at https://src.koda.cnrs.fr/MAURINAmandine/salten.

## Supplemental Online Materials

**Figure S1.**
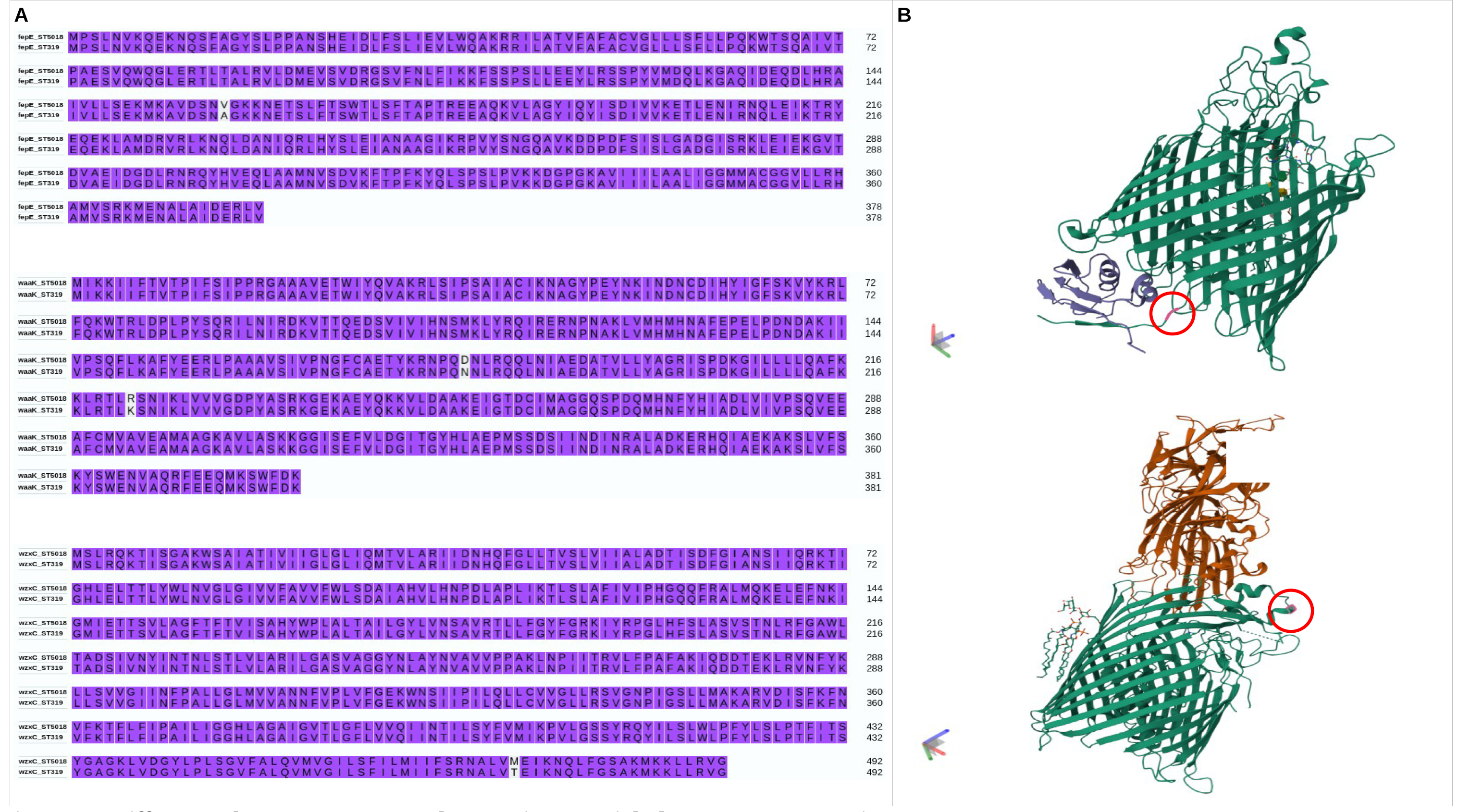
Differences between ST5018 and ST319 in potential phage receptor proteins. **A-**Amino-acid sequence alignment of fepE, waaK and wzxC between ST5018 and ST319, three proteins involved il LPS synthesis. **B-**FhuA protein structure and the two amino-acids differing between ST5018 and ST319, located in a periplasmic loop and in the plug.

**Figure S2.**
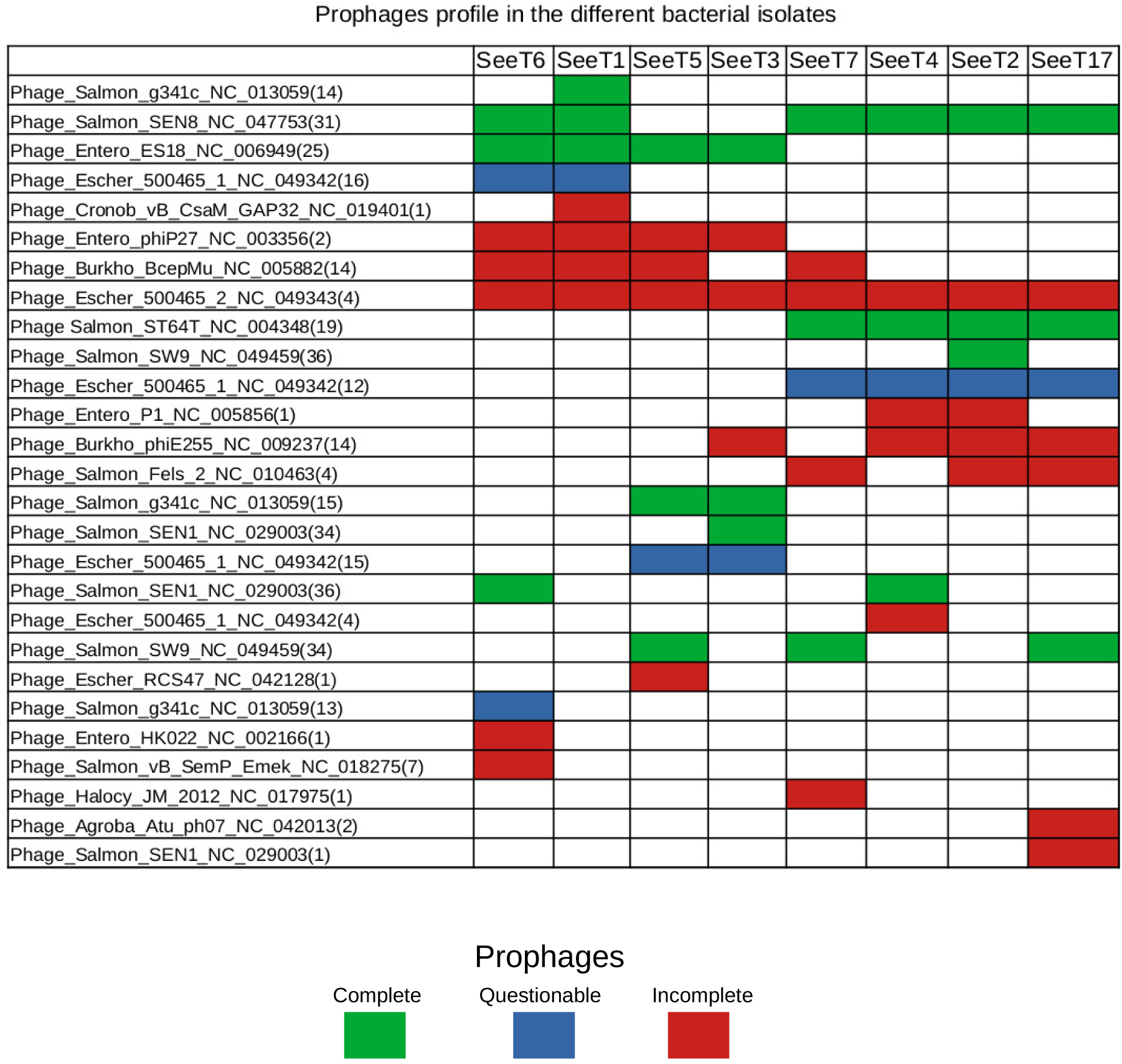
List of prophages present in each bacterial isolate. Prophages detected by Phaster (Arndt et al., 2019). Complete prophages are in green (score >90), questionable in blue (score 70 - 90) and incomplete prophages in red (score <70).

**Figure S3.**
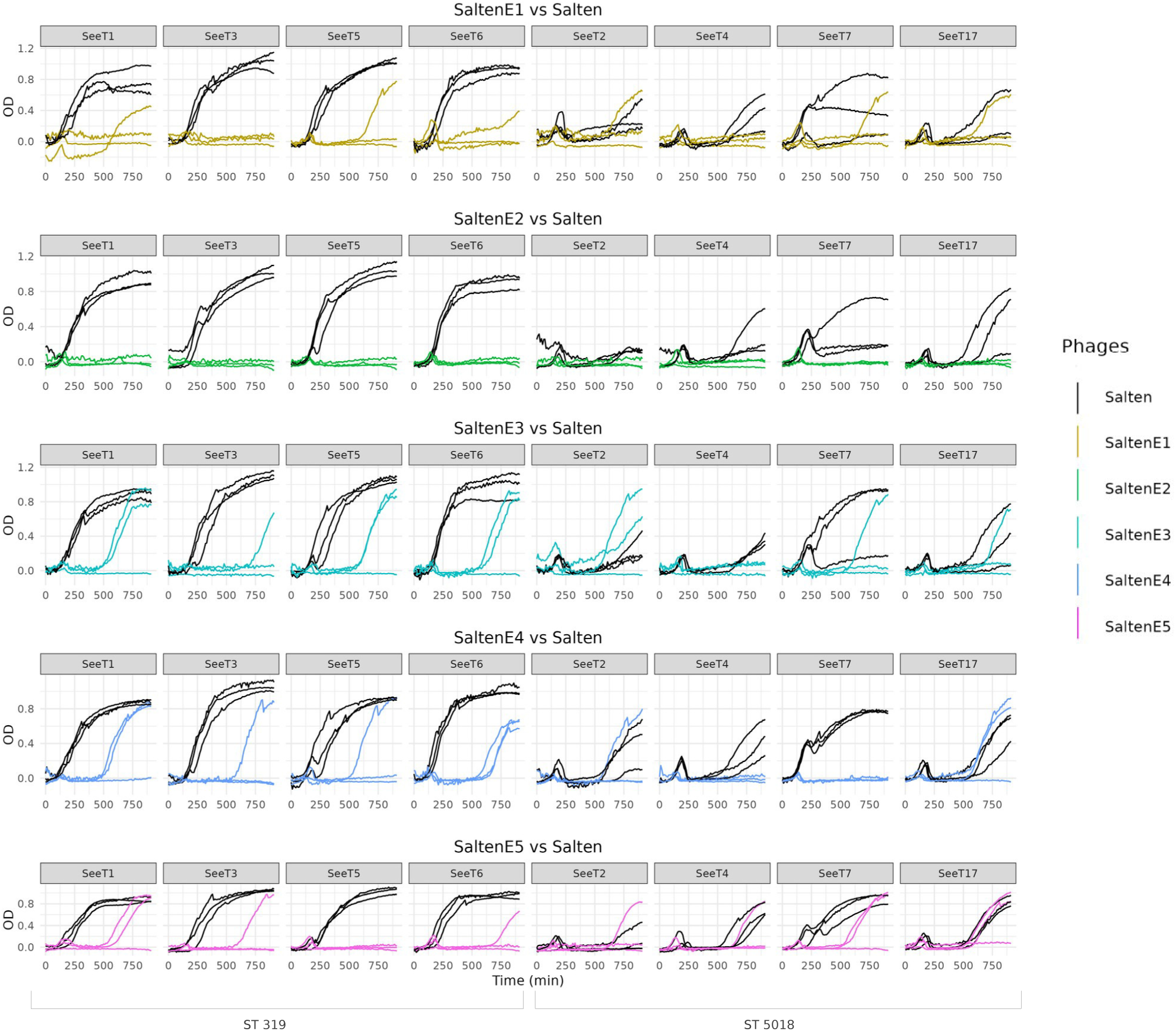
Bacterial kinetics monitored during 16 h at OD_600nm_. Ancestral phage Salten kinetics compared to each evolved populations SaltenE, on each bacterial isolates according to their sequence type (ST).

**Figure S4.**
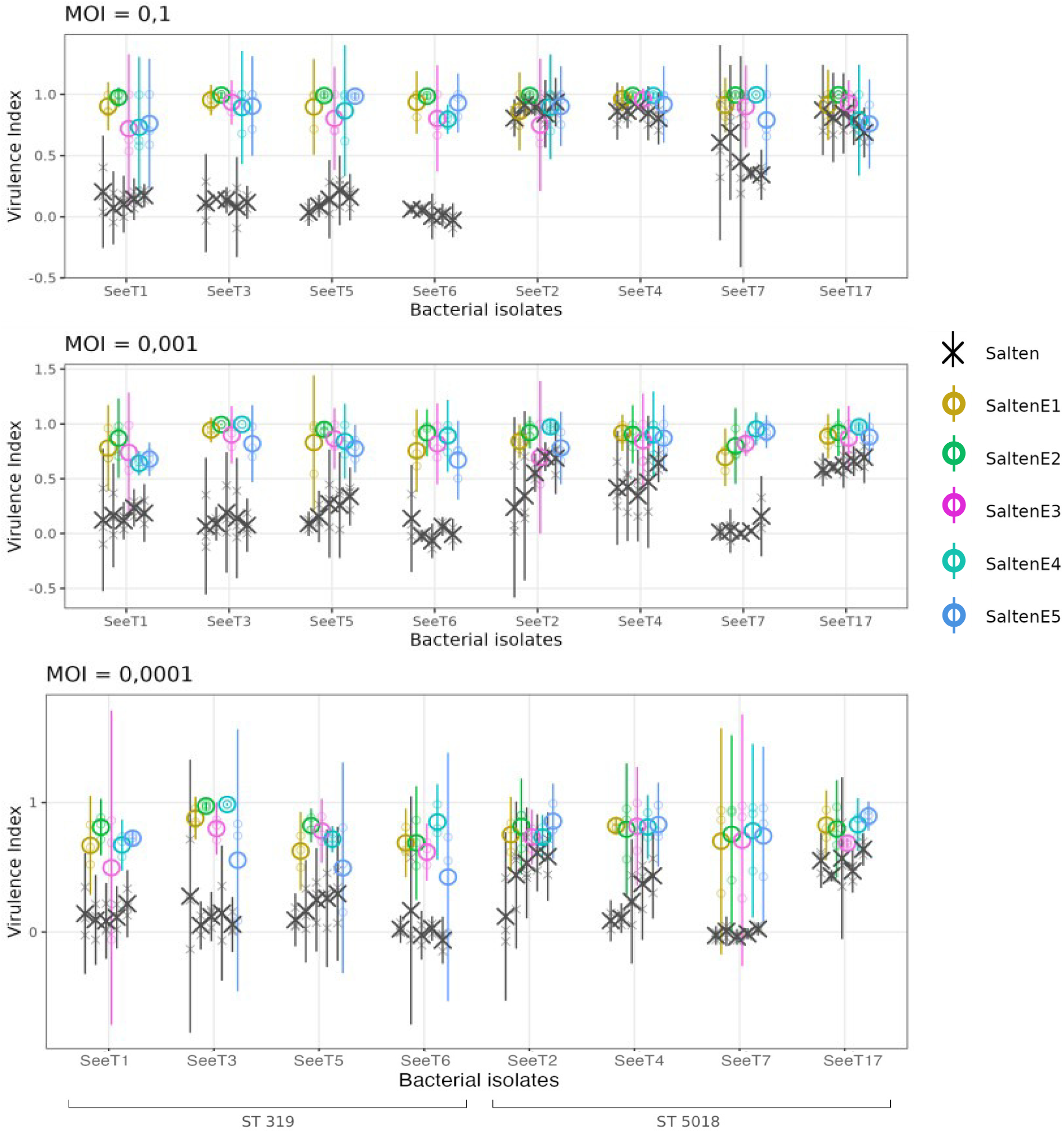
Phenotypic characterization (host-range and virulence) of the ancestral Salten and the evolved SaltenE phage populations on the eight *S. enterica* serotype Tennessee (SeeT) isolates, at different Multiplicity Of Infection (MOI). SeeT isolates are displayed according to their sequence type (ST319 or ST5018). Evaluation of phage virulence index was made in liquid, in presence of ancestral (black cross) or experimentally evolved (colored circles) phage populations.

**Figure S5.**
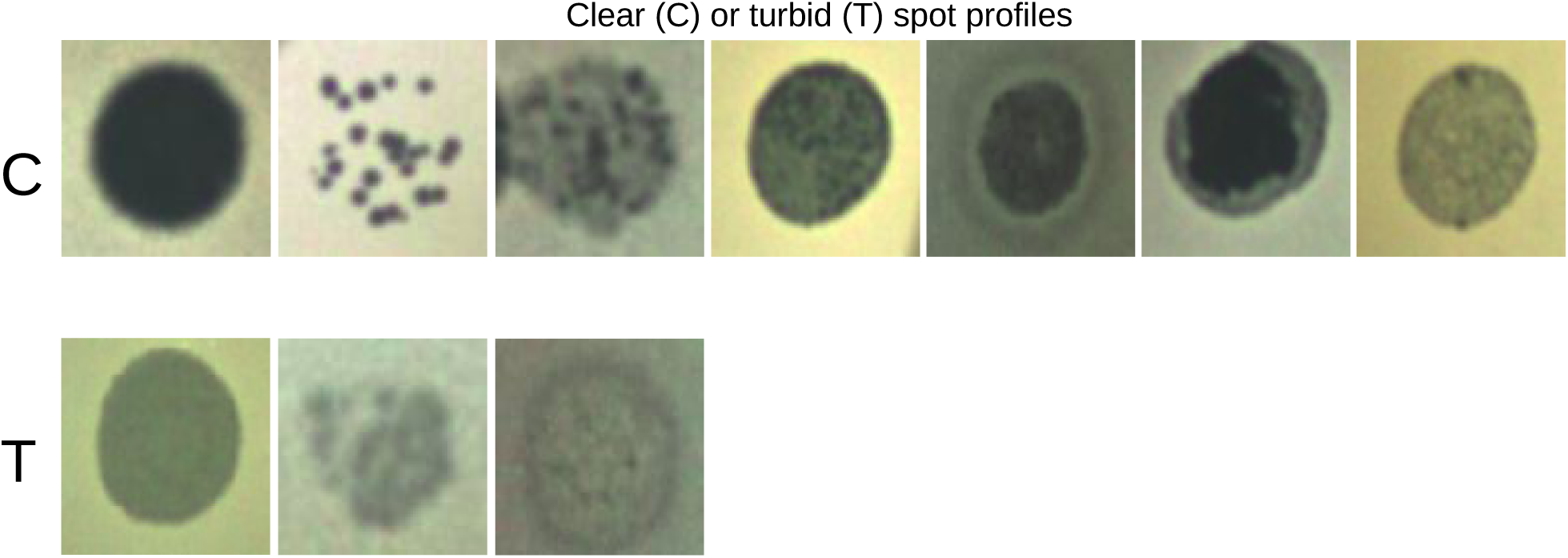
Visual evaluation of clear or turbid plaques. Each image represents an example of a visual assessment assigned to clear (C) or turbid (T).

**Figure S6.**
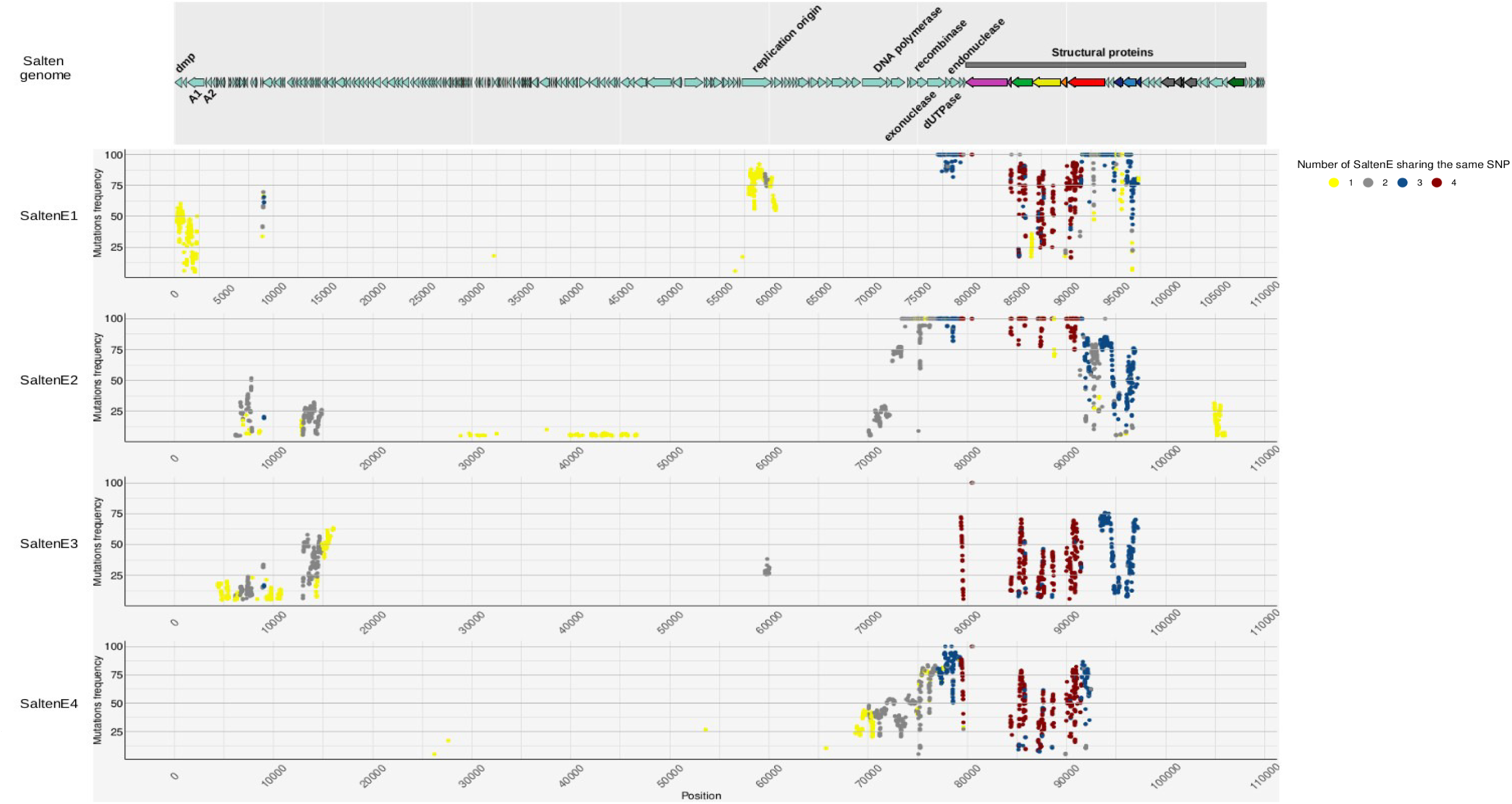
Frequency of parallel mutations according to their genomic position. Frequency of mutations accumulated in four independent evolved populations (SaltenE1, E2, E3, E4). Colors of each mutation correspond to its presence over populations: yellow mutations are present in only one evolved population, gray in two, blue in three and dark red in four evolved phage populations.

**Figure S7.**
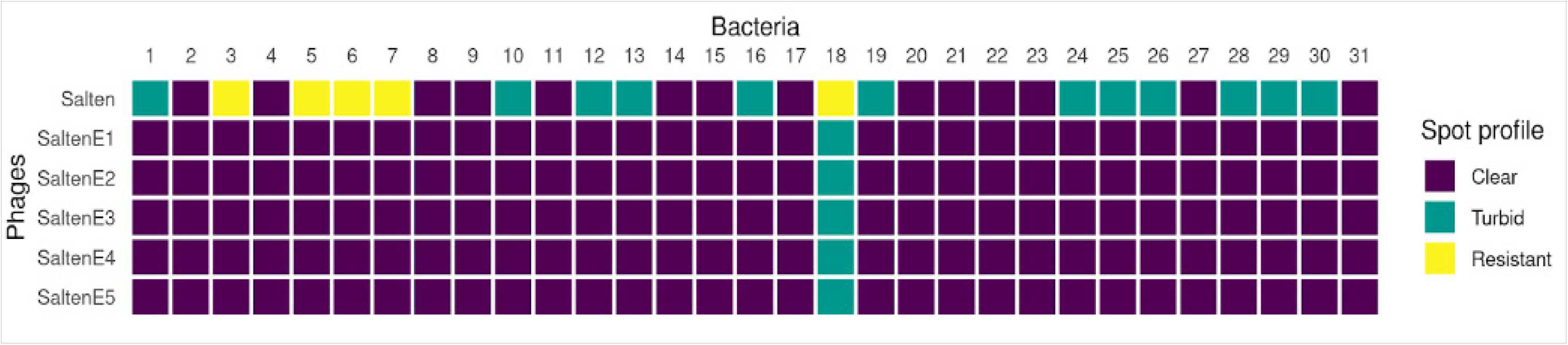
Virulence of the ancestral phage Salten and the five independent evolved populations SaltenE in solid condition against the 31 isolates of *S. enterica* serotype Tennessee (SeeT). Plaques visual evaluation were assessed following a spot-assay. Clear plaques are represented in dark blue, turbid plaques in blue-green and no-plaque observed (resistant bacterial isolate to the phage) are represented in yellow.

**Figure S8.**
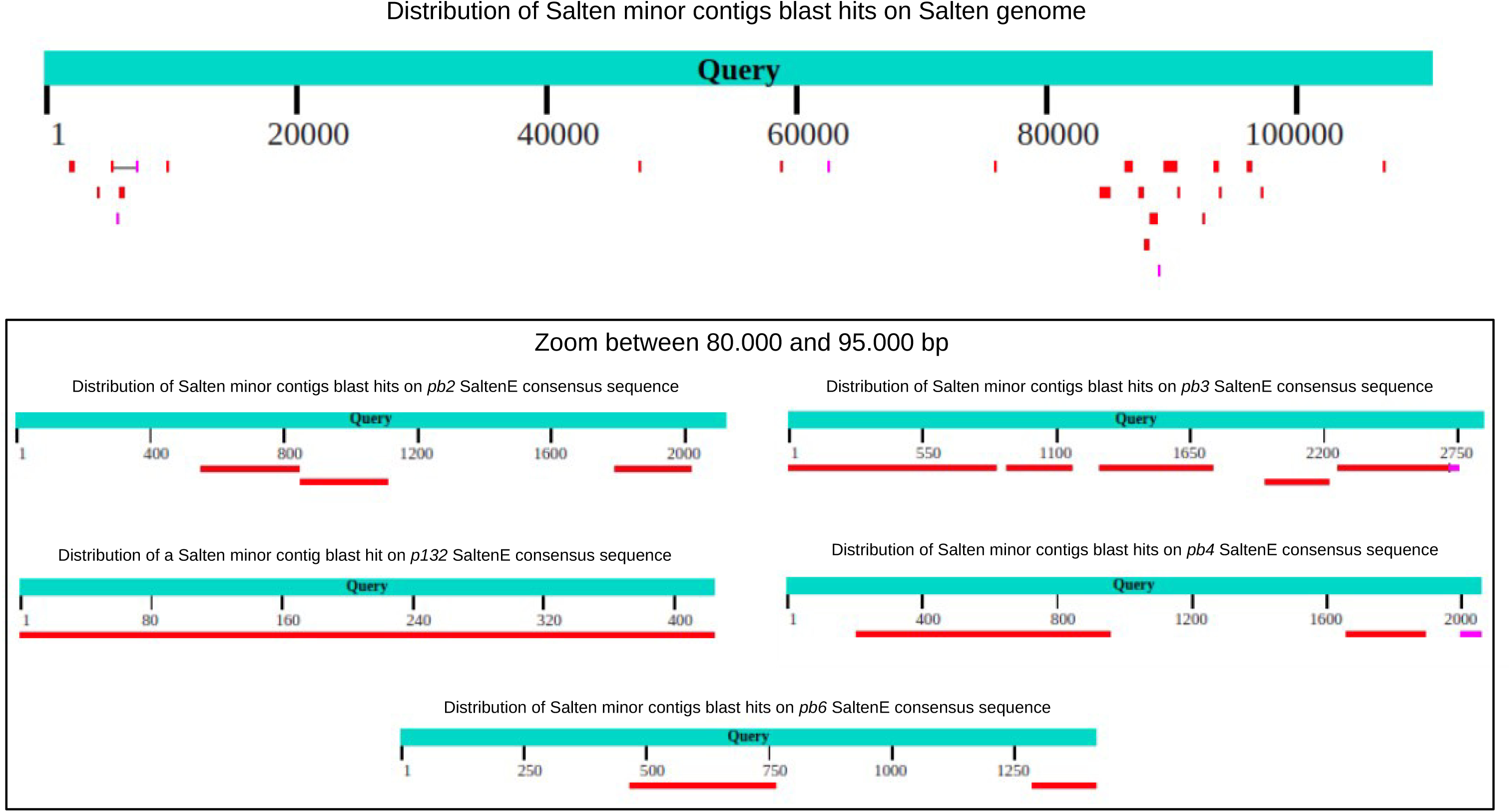
Distribution of minor Salten contigs along the Salten genome and evolved consensus structural genes. Minor Salten contigs were aligned using blastn in NCBI platform (Blast® services, available from: https://www.ncbi.nlm.nih.gov/Blast.cgi).

**Figure S9.**
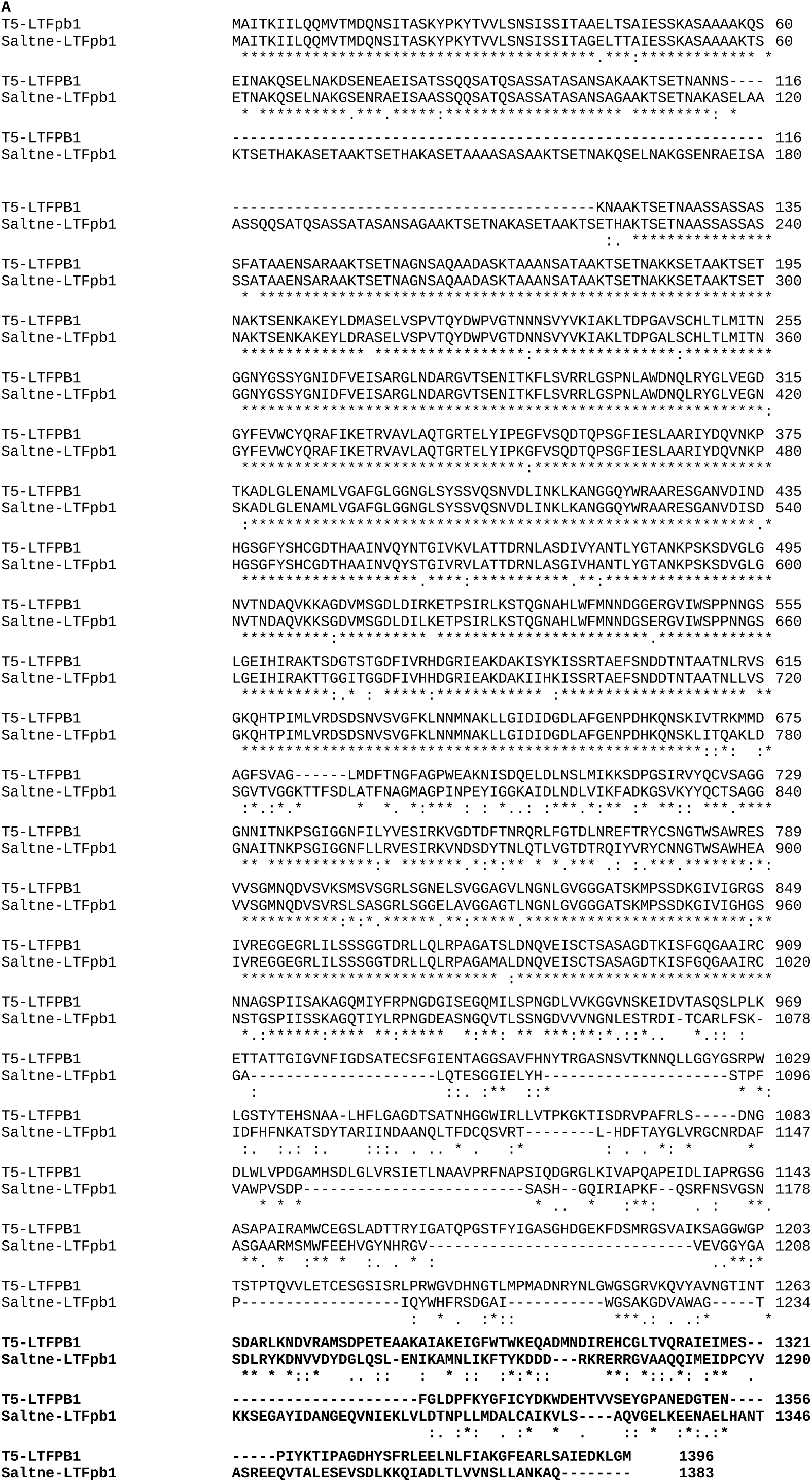
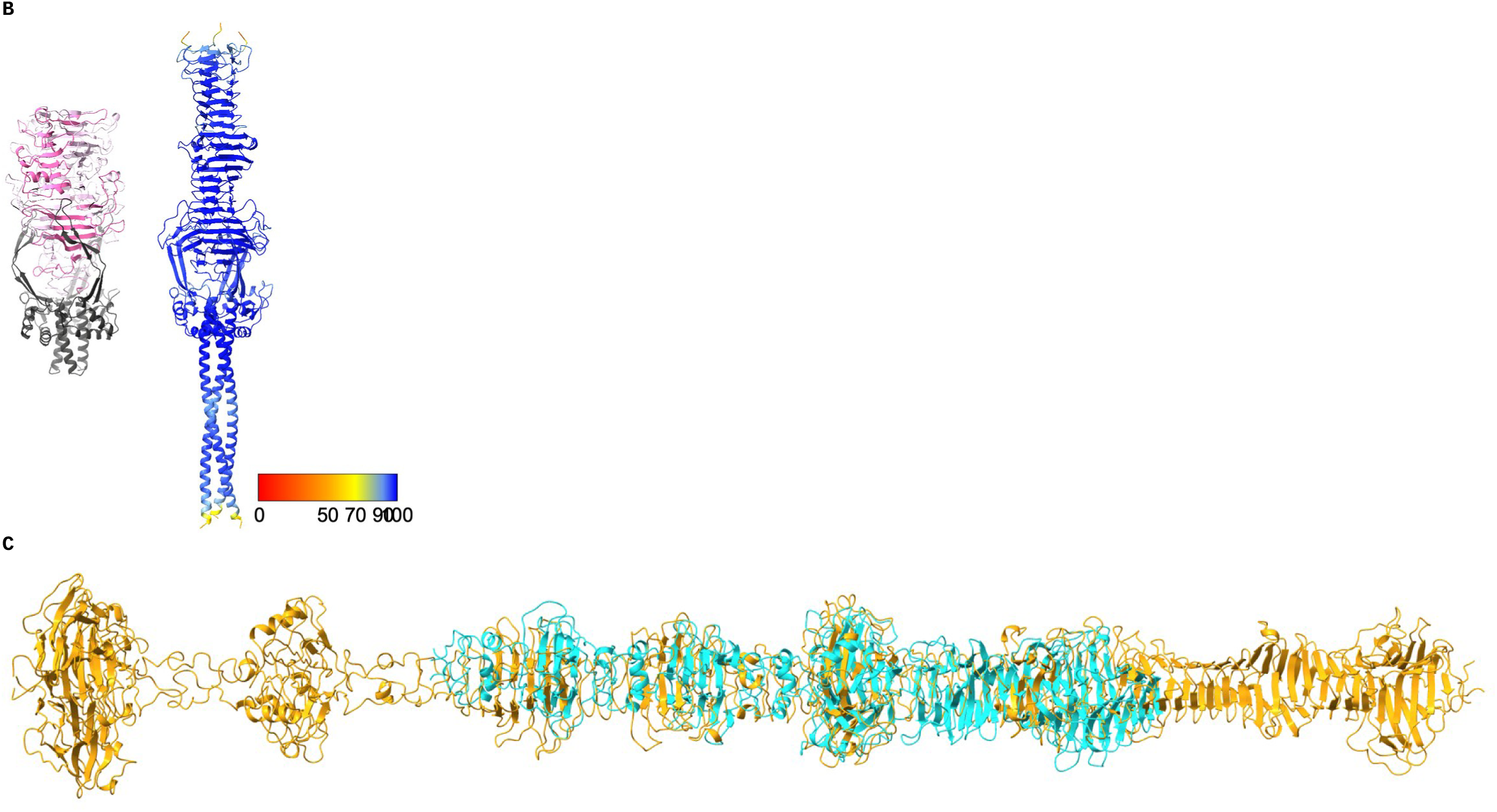
Analysis of LTFpb1. **A-**Sequence alignment of T5- and Salten-LTFpb1. The chaperone domain is in bold. **B-**Structure of the C-terminal domains of T5-LTFpb1 (left, the chaperone domain is in dark grey, the polymannose binding domain in shades of pink) and AlphaFold2 predicted structure of Salten-LTFpb1 C-terminal domains (residues 1022 to 1386), coloured according to the confidence factor plDDT. **C-**DALI alignment of the AlphaFold2 predicted structure from the fibre domain of Salten-LTFpb1 (orange) and of BD13-LTFpb1 (cyan), the chaperone domain has been removed in both predicted structure for clarity.

**Table S1.**
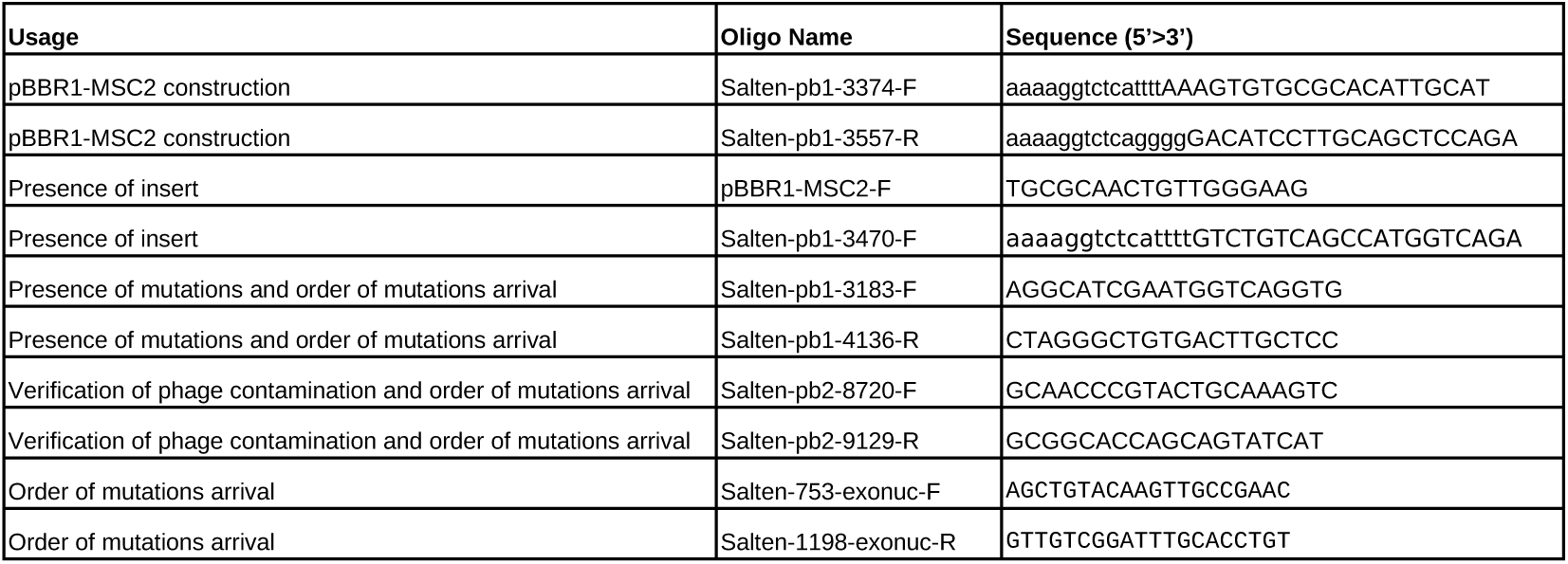
Primer list.

**Table S2. Breseq output.** List of mutations, their position on the ancestral Salten genome, their frequency along evolved reads in each evolved phage populations SaltenE and the name of the protein where they are located.

