## Supplementary material for "Convergent evolution during host range expansion and virulence increase in a *Salmonella* bacteriophage": Table S2. Breseq output.

List of mutations, their position on the ancestral Salten genome, their frequency along evolved reads in each evolved phage populations SaltenE and the name of the protein where they are located.

| | | | Frequency | | | | |
|---|---|---|---|---|---|---|---|
| Position | Mutation (nucleotide) | Mutation (amino-acid) | SaltenE1 | SaltenE2 | SaltenE3 | SaltenE4 | Protein |
| 301 | T→C | G145G (GGA→GGG) | 47.5% | | | | hypothetical protein |
| 302 | C→T | G145E (GGA→GAA) | 46.1% | | | | hypothetical protein |
| 307 | A→G | Y143Y (TAT→TAC) | 46.3% | | | | hypothetical protein |
| 310 | G→A | Y142Y (TAC→TAT) | 45.0% | | | | hypothetical protein |
| 312 | A→G | Y142H (TAC→CAC) | 46.6% | | | | hypothetical protein |
| 313 | A→G | S141S (AGT→AGC) | 47.2% | | | | hypothetical protein |
| 324 | C→T | V138I (GTT→ATT) | 49.1% | | | | hypothetical protein |
| 325 | T→A | A137A (GCA→GCT) | 49.6% | | | | hypothetical protein |
| 334 | G→A | S134S (AGC→AGT) | 51.6% | | | | hypothetical protein |
| 346 | T→C | E130E (GAA→GAG) | 56.8% | | | | hypothetical protein |
| 364 | T→C | K124K (AAA→AAG) | 53.9% | | | | hypothetical protein |
| 366 | T→C | K124E (AAA→GAA) | 53.9% | | | | hypothetical protein |
| 376 | A→G | H120H (CAT→CAC) | 53.4% | | | | hypothetical protein |
| 379 | C→T | L119L (CTG→CTA) | 53.1% | | | | hypothetical protein |
| 409 | A→G | S109S (AGT→AGC) | 48.4% | | | | hypothetical protein |
| 415 | T→A | G107G (GGA→GGT) | 47.4% | | | | hypothetical protein |
| 416 | C→T | G107E (GGA→GAA) | 48.0% | | | | hypothetical protein |
| 421 | C→A | G105G (GGG→GGT) | 49.2% | | | | hypothetical protein |
| 424 | T→G | R104R (CGA→CGC) | 47.5% | | | | hypothetical protein |
| 430 | T→G | G102G (GGA→GGC) | 48.1% | | | | hypothetical protein |
| 457 | T→C | S93S (TCA→TCG) | 44.9% | | | | hypothetical protein |
| 460 | T→C | K92K (AAA→AAG) | 45.0% | | | | hypothetical protein |
| 469 | T→C | L89L (CTA→CTG) | 42.5% | | | | hypothetical protein |
| 471 | G→A | L89L (CTA→TTA) | 42.1% | | | | hypothetical protein |
| 481 | A→G | C85C (TGT→TGC) | 38.3% | | | | hypothetical protein |
| 484 | A→G | I84I (ATT→ATC) | 37.2% | | | | hypothetical protein |
| 490 | G→A | N82N (AAC→AAT) | 36.8% | | | | hypothetical protein |
| 496 | A→C | T80T (ACT→ACG) | 29.5% | | | | hypothetical protein |
| 499 | A→G | N79N (AAT→AAC) | 29.1% | | | | hypothetical protein |
| 502 | T→A | A78A (GCA→GCT) | 29.7% | | | | hypothetical protein |
| 503 | G→A | A78V (GCA→GTA) | 30.2% | | | | hypothetical protein |
| 505 | A→G | D77D (GAT→GAC) | 31.5% | | | | hypothetical protein |
| 514 | A→G | C74C (TGT→TGC) | 39.5% | | | | hypothetical protein |
| 517 | T→C | Q73Q (CAA→CAG) | 40.2% | | | | hypothetical protein |
| 530 | T→G | E69A (GAA→GCA) | 39.9% | | | | hypothetical protein |
| 531 | C→T | E69K (GAA→AAA) | 40.2% | | | | hypothetical protein |
| 535 | T→A | L67F (TTA→TTT) | 39.8% | | | | hypothetical protein |
| 537 | A→G | L67L (TTA→CTA) | 41.0% | | | | hypothetical protein |
| 538 | T→C | P66P (CCA→CCG) | 41.3% | | | | hypothetical protein |
| 546 | A→G | L64L (TTA→CTA) | 46.9% | | | | hypothetical protein |
| 547 | G→T | T63T (ACC→ACA) | 46.1% | | | | hypothetical protein |
| 567 | G→C | H57D (CAT→GAT) | 54.7% | | | | hypothetical protein |
| 616 | A→G | N40N (AAT→AAC) | 56.5% | | | | hypothetical protein |
| 617 | T→C | N40S (AAT→AGT) | 57.7% | | | | hypothetical protein |
| 622 | G→A | F38F (TTC→TTT) | 60.3% | | | | hypothetical protein |

| | | | | |
|---|---|---|---|---|
| 646 | A→C | S30S (TCT→TCG) | 54.2% | hypothetical protein |
| 654 | C→T | V28I (GTT→ATT) | 52.0% | hypothetical protein |
| 667 | T→C | L23L (TTA→TTG) | 48.3% | hypothetical protein |
| 669 | A→G | L23L (TTA→CTA) | 48.9% | hypothetical protein |
| 697 | T→C | P13P (CCA→CCG) | 51.5% | hypothetical protein |
| 711 | A→T | S9T (TCC→ACC) | 46.9% | hypothetical protein |
| 714 | G→T | L8I (CTT→ATT) | 48.7% | hypothetical protein |
| 719 | A→G | M6T (ATG→ACG) | 44.5% | hypothetical protein |
| 721 | G→T | I5I (ATC→ATA) | 44.5% | hypothetical protein |
| 722 | A→T | I5N (ATC→AAC) | 44.5% | hypothetical protein |
| 725 | G→A | A4V (GCT→GTT) | 44.9% | hypothetical protein |
| 729 | T→G | K3Q (AAA→CAA) | 45.5% | hypothetical protein |
| 809 | C→T | intergenic (-74/+6) | 54.3% | Phage protein |
| 827 | A→C | A127A (GCT→GCG) | 52.3% | Phage protein |
| 842 | G→A | I122I (ATC→ATT) | 49.4% | Phage protein |
| 857 | C→T | E117E (GAG→GAA) | 48.9% | Phage protein |
| 863 | G→A | S115S (TCC→TCT) | 48.7% | Phage protein |
| 870 | T→C | Y113C (TAT→TGT) | 46.1% | Phage protein |
| 871 | A→C | Y113D (TAT→GAT) | 46.0% | Phage protein |
| 884 | T→C | A108A (GCA→GCG) | 50.8% | Phage protein |
| 899 | T→C | K103K (AAA→AAG) | 48.0% | Phage protein |
| 908 | C→G | M100I (ATG→ATC) | 46.6% | Phage protein |
| 909 | A→G | M100T (ATG→ACG) | 47.3% | Phage protein |
| 911 | T→C | L99L (TTA→TTG) | 45.9% | Phage protein |
| 914 | C→T | Q98Q (CAG→CAA) | 45.3% | Phage protein |
| 956 | G→C | A84A (GCC→GCG) | 27.5% | Phage protein |
| 965 | G→T | S81S (TCC→TCA) | 20.8% | Phage protein |
| 974 | T→A | A78A (GCA→GCT) | 12.0% | Phage protein |
| 980 | G→A | I76I (ATC→ATT) | 5.8% | Phage protein |
| 1,246 | T→G | intergenic (-39/+28) | 10.5% | Phage protein |
| 1,259 | A→C | intergenic (-52/+15) | 14.5% | Phage protein |
| 1,263 | T→G | intergenic (-56/+11) | 14.8% | Phage protein |
| 1,265:1 | +T | intergenic (-58/+9) | 13.4% | Phage protein |
| 1,266 | C→A | intergenic (-59/+8) | 13.2% | Phage protein |
| 1,277 | T→G | A556A (GCA→GCC) | 25.6% | hypothetical protein |
| 1,289 | A→G | G552G (GGT→GGC) | 32.6% | hypothetical protein |
| 1,298 | A→G | F549F (TTT→TTC) | 31.7% | hypothetical protein |
| 1,301 | G→A | D548D (GAC→GAT) | 30.7% | hypothetical protein |
| 1,313 | G→A | R544R (CGC→CGT) | 36.6% | hypothetical protein |
| 1,316 | C→T | Q543Q (CAG→CAA) | 36.5% | hypothetical protein |
| 1,319 | A→C | G542G (GGT→GGG) | 36.2% | hypothetical protein |
| 1,367 | A→C | V526V (GTT→GTG) | 41.4% | hypothetical protein |
| 1,376 | A→G | G523G (GGT→GGC) | 38.1% | hypothetical protein |
| 1,382 | G→A | D521D (GAC→GAT) | 31.4% | hypothetical protein |
| 1,385 | C→A | L520L (CTG→CTT) | 32.1% | hypothetical protein |
| 1,388 | A→G | S519S (AGT→AGC) | 33.0% | hypothetical protein |
| 1,389 | C→G | S519T (AGT→ACT) | 33.1% | hypothetical protein |
| 1,39 | T→A | S519C (AGT→TGT) | 32.9% | hypothetical protein |
| 1,43 | T→A | A505A (GCA→GCT) | 37.0% | hypothetical protein |
| 1,436 | A→G | H503H (CAT→CAC) | 37.8% | hypothetical protein |
| 1,442 | A→G | H501H (CAT→CAC) | 37.7% | hypothetical protein |
| 1,457 | C→G | K496N (AAG→AAC) | 28.9% | hypothetical protein |
| 1,458 | T→G | K496T (AAG→ACG) | 29.3% | hypothetical protein |

| | | | | |
|---|---|---|---|---|
| 1,459 | T→C | K496E (AAG→GAG) | 28.7% | hypothetical protein |
| 1,460:1 | +T | coding (1485/1671 nt) | 29.7% | hypothetical protein |
| 1,466 | T→C | K493K (AAA→AAG) | 34.5% | hypothetical protein |
| 1,478 | T→C | K489K (AAA→AAG) | 39.9% | hypothetical protein |
| 1,493 | G→A | S484S (AGC→AGT) | 43.3% | hypothetical protein |
| 1,508 | G→A | N479N (AAC→AAT) | 44.1% | hypothetical protein |
| 1,517 | G→A | H476H (CAC→CAT) | 45.3% | hypothetical protein |
| 1,529 | T→C | E472E (GAA→GAG) | 47.1% | hypothetical protein |
| 1,535 | G→T | G470G (GGC→GGA) | 46.6% | hypothetical protein |
| 1,538 | C→T | A469A (GCG→GCA) | 47.4% | hypothetical protein |
| 1,562 | G→A | T461T (ACC→ACT) | 47.7% | hypothetical protein |
| 1,574 | G→A | A457A (GCC→GCT) | 46.2% | hypothetical protein |
| 1,583 | C→T | E454E (GAG→GAA) | 42.7% | hypothetical protein |
| 1,595 | G→A | V450V (GTC→GTT) | 39.0% | hypothetical protein |
| 1,601 | A→T | R448R (CGT→CGA) | 36.5% | hypothetical protein |
| 1,603 | G→T | R448S (CGT→AGT) | 35.9% | hypothetical protein |
| 1,604 | C→A | L447L (CTG→CTT) | 36.1% | hypothetical protein |
| 1,616 | T→C | L443L (CTA→CTG) | 33.6% | hypothetical protein |
| 1,618 | G→A | L443L (CTA→TTA) | 33.5% | hypothetical protein |
| 1,619 | C→T | V442V (GTG→GTA) | 33.2% | hypothetical protein |
| 1,625 | G→A | F440F (TTC→TTT) | 35.1% | hypothetical protein |
| 1,628 | A→G | T439T (ACT→ACC) | 35.3% | hypothetical protein |
| 1,637 | C→T | Q436Q (CAG→CAA) | 37.5% | hypothetical protein |
| 1,649 | A→T | A432A (GCT→GCA) | 40.4% | hypothetical protein |
| 1,652 | T→G | A431A (GCA→GCC) | 40.9% | hypothetical protein |
| 1,655 | A→G | Y430Y (TAT→TAC) | 43.1% | hypothetical protein |
| 1,658 | G→A | I429I (ATC→ATT) | 43.0% | hypothetical protein |
| 1,661 | A→C | A428A (GCT→GCG) | 43.2% | hypothetical protein |
| 1,664 | G→A | A427A (GCC→GCT) | 42.1% | hypothetical protein |
| 1,688 | C→G | A419A (GCG→GCC) | 40.8% | hypothetical protein |
| 1,703 | G→A | D414D (GAC→GAT) | 37.0% | hypothetical protein |
| 1,706 | T→C | K413K (AAA→AAG) | 38.6% | hypothetical protein |
| 1,709 | G→A | I412I (ATC→ATT) | 37.6% | hypothetical protein |
| 1,718 | A→G | N409N (AAT→AAC) | 39.0% | hypothetical protein |
| 1,73 | C→A | L405F (TTG→TTT) | 37.0% | hypothetical protein |
| 1,732 | A→G | L405L (TTG→CTG) | 37.4% | hypothetical protein |
| 1,736 | A→G | R403R (CGT→CGC) | 37.1% | hypothetical protein |
| 1,739 | G→T | S402S (TCC→TCA) | 33.6% | hypothetical protein |
| 1,742 | T→A | L401F (TTA→TTT) | 33.8% | hypothetical protein |
| 1,744 | A→G | L401L (TTA→CTA) | 33.7% | hypothetical protein |
| 1,75 | G→A | L399L (CTA→TTA) | 34.0% | hypothetical protein |
| 1,754 | G→A | H397H (CAC→CAT) | 32.9% | hypothetical protein |
| 1,76 | A→T | S395S (TCT→TCA) | 33.5% | hypothetical protein |
| 1,781 | C→A | S388S (TCG→TCT) | 27.0% | hypothetical protein |
| 1,79 | G→A | R385R (CGC→CGT) | 21.0% | hypothetical protein |
| 1,793 | T→C | E384E (GAA→GAG) | 20.8% | hypothetical protein |
| 1,799 | T→A | S382S (TCA→TCT) | 16.6% | hypothetical protein |
| 1,802 | T→C | E381E (GAA→GAG) | 16.8% | hypothetical protein |
| 1,805 | C→T | L380L (CTG→CTA) | 17.0% | hypothetical protein |
| 1,807 | G→A | L380L (CTG→TTG) | 16.6% | hypothetical protein |
| 1,808 | C→A | V379V (GTG→GTT) | 17.2% | hypothetical protein |
| 1,814 | C→T | G377G (GGG→GGA) | 17.9% | hypothetical protein |
| 1,82 | A→G | D375D (GAT→GAC) | 18.5% | hypothetical protein |

| | | | | |
|---|---|---|---|---|
| 1,826 | T→A | L373L (CTA→CTT) | 17.8% | hypothetical protein |
| 1,835 | C→T | L370L (CTG→CTA) | 14.3% | hypothetical protein |
| 1,837 | G→A | L370L (CTG→TTG) | 13.9% | hypothetical protein |
| 1,841 | T→C | K368K (AAA→AAG) | 14.3% | hypothetical protein |
| 1,844 | A→G | D367D (GAT→GAC) | 15.0% | hypothetical protein |
| 1,856 | T→G | A363A (GCA→GCC) | 12.5% | hypothetical protein |
| 1,859 | A→G | Y362Y (TAT→TAC) | 11.9% | hypothetical protein |
| 1,868 | C→A | A359A (GCG→GCT) | 8.2% | hypothetical protein |
| 1,871 | T→A | L358L (CTA→CTT) | 7.8% | hypothetical protein |
| 2,024 | A→G | A307A (GCT→GCC) | 5.6% | hypothetical protein |
| 2,036 | A→G | G303G (GGT→GGC) | 5.4% | hypothetical protein |
| 2,053 | A→G | L298L (TTA→CTA) | 5.5% | hypothetical protein |
| 2,06 | C→A | G295G (GGG→GGT) | 5.4% | hypothetical protein |
| 2,063 | C→G | T294T (ACG→ACC) | 6.0% | hypothetical protein |
| 2,065 | T→A | T294S (ACG→TCG) | 7.1% | hypothetical protein |
| 2,066 | T→C | T293T (ACA→ACG) | 6.0% | hypothetical protein |
| 2,073 | T→A | Y291F (TAT→TTT) | 5.2% | hypothetical protein |
| 2,075 | G→A | C290C (TGC→TGT) | 5.2% | hypothetical protein |
| 2,078 | C→T | G289G (GGG→GGA) | 5.5% | hypothetical protein |
| 2,081 | T→G | A288A (GCA→GCC) | 5.6% | hypothetical protein |
| 2,083 | C→T | A288T (GCA→ACA) | 5.1% | hypothetical protein |
| 2,084 | G→A | P287P (CCC→CCT) | 5.1% | hypothetical protein |
| 2,087 | A→C | T286T (ACT→ACG) | 5.6% | hypothetical protein |
| 2,088 | G→A | T286I (ACT→ATT) | 5.0% | hypothetical protein |
| 2,09 | G→A | A285A (GCC→GCT) | 5.4% | hypothetical protein |
| 2,099 | G→A | N282N (AAC→AAT) | 15.9% | hypothetical protein |
| 2,104 | G→A | L281L (CTG→TTG) | 19.2% | hypothetical protein |
| 2,111 | A→G | F278F (TTT→TTC) | 17.9% | hypothetical protein |
| 2,116 | C→T | V277M (GTG→ATG) | 17.9% | hypothetical protein |
| 2,117 | C→T | G276G (GGG→GGA) | 18.1% | hypothetical protein |
| 2,12 | A→G | S275S (AGT→AGC) | 17.2% | hypothetical protein |
| 2,147 | A→G | V266V (GTT→GTC) | 17.9% | hypothetical protein |
| 2,148 | Δ1 bp | coding (797/1671 nt) | 17.8% | hypothetical protein |
| 2,151:1 | +T | coding (794/1671 nt) | 17.8% | hypothetical protein |
| 2,159 | C→T | G262G (GGG→GGA) | 18.3% | hypothetical protein |
| 2,165 | C→G | E260D (GAG→GAC) | 18.7% | hypothetical protein |
| 2,177 | A→C | V256V (GTT→GTG) | 16.2% | hypothetical protein |
| 2,18 | A→C | I255M (ATT→ATG) | 16.3% | hypothetical protein |
| 2,182 | T→G | I255L (ATT→CTT) | 15.6% | hypothetical protein |
| 2,185 | G→A | L254L (CTG→TTG) | 16.4% | hypothetical protein |
| 2,186 | C→A | A253A (GCG→GCT) | 16.4% | hypothetical protein |
| 2,189 | A→G | G252G (GGT→GGC) | 16.8% | hypothetical protein |
| 2,193 | A→T | F251Y (TTC→TAC) | 15.4% | hypothetical protein |
| 2,201 | C→T | L248L (TTG→TTA) | 17.9% | hypothetical protein |
| 2,204 | C→A | A247A (GCG→GCT) | 19.4% | hypothetical protein |
| 2,207 | T→C | E246E (GAA→GAG) | 19.7% | hypothetical protein |
| 2,21 | C→A | A245A (GCG→GCT) | 19.6% | hypothetical protein |
| 2,219 | C→A | G242G (GGG→GGT) | 28.2% | hypothetical protein |
| 2,228 | T→C | Q239Q (CAA→CAG) | 28.2% | hypothetical protein |
| 2,24 | A→G | N235N (AAT→AAC) | 29.1% | hypothetical protein |
| 2,246 | T→A | L233L (CTA→CTT) | 27.6% | hypothetical protein |
| 2,252 | T→A | A231A (GCA→GCT) | 37.5% | hypothetical protein |
| 2,258 | G→T | G229G (GGC→GGA) | 37.6% | hypothetical protein |

| | | | | | |
|---|---|---|---|---|---|
| 2,264 | C→T | S227S (TCG→TCA) | 37.8% | | hypothetical protein |
| 2,267 | G→A | Y226Y (TAC→TAT) | 37.2% | | hypothetical protein |
| 2,279 | T→C | V222V (GTA→GTG) | 50.0% | | hypothetical protein |
| 4,397 | T→C | Y15Y (TAT→TAC) | | 18.3% | hypothetical protein |
| 4,401 | T→G | L17V (TTA→GTA) | | 16.9% | hypothetical protein |
| 4,403 | A→G | L17L (TTA→TTG) | | 16.6% | hypothetical protein |
| 4,422 | T→C | F24L (TTT→CTT) | | 17.6% | hypothetical protein |
| 4,434 | A→G | I28V (ATC→GTC) | | 16.4% | hypothetical protein |
| 4,442 | A→G | K30K (AAA→AAG) | | 18.4% | hypothetical protein |
| 4,452 | C→T | L34L (CTG→TTG) | | 16.9% | hypothetical protein |
| 4,46 | A→T | S36S (TCA→TCT) | | 16.7% | hypothetical protein |
| 4,465 | A→C | D38A (GAC→GCC) | | 15.2% | hypothetical protein |
| 4,47 | A→C | I40L (ATA→CTA) | | 16.6% | hypothetical protein |
| 4,679 | A→G | intergenic (-20/+36) | | 18.6% | hypothetical protein |
| 4,68 | T→A | intergenic (-21/+35) | | 17.6% | hypothetical protein |
| 4,71 | G→A | intergenic (-51/+5) | | 11.4% | hypothetical protein |
| 4,712 | T→A | intergenic (-53/+3) | | 11.5% | hypothetical protein |
| 4,716 | Δ1 bp | coding (221/222 nt) | | 12.3% | hypothetical protein |
| 4,718 | G→A | N73N (AAC→AAT) | | 11.5% | hypothetical protein |
| 4,731 | A→G | M69T (ATG→ACG) | | 10.9% | hypothetical protein |
| 4,749 | A→G | V63A (GTG→GCG) | | 9.8% | hypothetical protein |
| 4,77 | G→T | A56E (GCG→GAG) | | 8.7% | hypothetical protein |
| 4,778 | T→A | S53S (TCA→TCT) | | 9.0% | hypothetical protein |
| 4,791 | G→C | S49C (TCC→TGC) | | 5.8% | hypothetical protein |
| 4,793 | Δ1 bp | coding (144/222 nt) | | 5.5% | hypothetical protein |
| 4,798 | C→G | V47L (GTA→CTA) | | 5.6% | hypothetical protein |
| 4,805 | G→T | P44P (CCC→CCA) | | 5.1% | hypothetical protein |
| 4,813 | A→C | F42V (TTT→GTT) | | 5.4% | hypothetical protein |
| 4,814 | T→C | I41M (ATA→ATG) | | 5.5% | hypothetical protein |
| 4,815 | A→T | I41K (ATA→AAA) | | 5.2% | hypothetical protein |
| 5,219 | C→T | intergenic (+117/-236) | | 6.9% | hypothetical protein |
| 5,237 | T→G | intergenic (+135/-218) | | 8.2% | hypothetical protein |
| 5,238 | G→C | intergenic (+136/-217) | | 7.8% | hypothetical protein |
| 5,241 | T→G | intergenic (+139/-214) | | 8.3% | hypothetical protein |
| 5,242 | T→A | intergenic (+140/-213) | | 8.1% | hypothetical protein |
| 5,244 | T→C | intergenic (+142/-211) | | 7.8% | hypothetical protein |
| 5,245 | C→T | intergenic (+143/-210) | | 7.9% | hypothetical protein |
| 5,265 | A→C | intergenic (+163/-190) | | 14.4% | hypothetical protein |
| 5,274 | C→T | intergenic (+172/-181) | | 15.4% | hypothetical protein |
| 5,275 | T→C | intergenic (+173/-180) | | 15.4% | hypothetical protein |
| 5,298 | A→G | intergenic (+196/-157) | | 19.1% | hypothetical protein |
| 5,299 | T→A | intergenic (+197/-156) | | 19.0% | hypothetical protein |
| 5,312 | A→T | intergenic (+210/-143) | | 20.1% | hypothetical protein |
| 5,336 | 5 bp→37 bp | intergenic (+234/-115) | | 18.4% | hypothetical protein |
| 5,34 | G→A | intergenic (+238/-115) | | 5.8% | hypothetical protein |
| 5,352 | G→T | intergenic (+250/-103) | | 15.6% | hypothetical protein |
| 5,353 | G→C | intergenic (+251/-102) | | 16.0% | hypothetical protein |
| 5,379 | C→G | intergenic (+277/-76) | | 15.9% | hypothetical protein |
| 5,386 | C→A | intergenic (+284/-69) | | 15.6% | hypothetical protein |
| 5,39 | A→T | intergenic (+288/-65) | | 13.5% | hypothetical protein |
| 5,396 | C→T | intergenic (+294/-59) | | 12.6% | hypothetical protein |
| 5,397 | C→A | intergenic (+295/-58) | | 13.6% | hypothetical protein |
| 5,41 | A→G | intergenic (+308/-45) | | 11.8% | hypothetical protein |

| | | | | | |
|---|---|---|---|---|---|
| 5,424 | A→G | intergenic (+322/-31) | | 9.2% | hypothetical protein |
| 5,428 | A→G | intergenic (+326/-27) | | 8.2% | hypothetical protein |
| 5,431 | T→A | intergenic (+329/-24) | | 8.0% | hypothetical protein |
| 5,433 | T→A | intergenic (+331/-22) | | 7.5% | hypothetical protein |
| 5,435 | G→A | intergenic (+333/-20) | | 7.0% | hypothetical protein |
| 5,436 | G→A | intergenic (+334/-19) | | 7.2% | hypothetical protein |
| 5,437 | C→G | intergenic (+335/-18) | | 7.2% | hypothetical protein |
| 5,457 | G→C | M1M (ATG→ATC) † | | 6.3% | hypothetical protein |
| 5,458 | T→C | Y2H (TAC→CAC) | | 6.4% | hypothetical protein |
| 5,462 | G→T | R3M (AGG→ATG) | | 6.9% | hypothetical protein |
| 6,091 | T→G | intergenic (+4/-43) | | 5.0% | Phage protein |
| 6,149 | T→G | S6A (TCT→GCT) | | 7.5% | Phage protein |
| 6,152 | G→A | V7I (GTT→ATT) | | 7.3% | Phage protein |
| 6,157 | A→G | L8L (CTA→CTG) | | 8.2% | Phage protein |
| 6,184 | C→T | S17S (TCC→TCT) | 5.3% | 8.6% | Phage protein |
| 6,185 | C→T | L18F (CTT→TTT) | 5.4% | 8.0% | Phage protein |
| 6,187 | T→G | L18L (CTT→CTG) | 5.8% | 9.1% | Phage protein |
| 6,193 | C→A | A20A (GCC→GCA) | 6.1% | 8.0% | Phage protein |
| 6,204 | G→A | G24D (GGC→GAC) | 5.1% | 7.2% | Phage protein |
| 6,213 | T→C | V27A (GTG→GCG) | 6.2% | 8.7% | Phage protein |
| 6,226 | C→T | G31G (GGC→GGT) | | 6.5% | Phage protein |
| 6,251 | T→C | L40L (TTG→CTG) | | 6.6% | Phage protein |
| 6,312 | C→A | S10S (TCC→TCA) | | 5.6% | Phage protein |
| 6,318 | T→C | G12G (GGT→GGC) | | 6.3% | Phage protein |
| 6,333 | T→C | D17D (GAT→GAC) | | 8.0% | Phage protein |
| 6,336 | C→T | V18V (GTC→GTT) | | 8.1% | Phage protein |
| 6,345 | T→C | F21F (TTT→TTC) | 5.0% | 7.9% | Phage protein |
| 6,351 | T→G | R23R (CGT→CGG) | | 7.6% | Phage protein |
| 6,352 | C→G | P24A (CCA→GCA) | | 7.6% | Phage protein |
| 6,353 | C→A | P24Q (CCA→CAA) | | 7.3% | Phage protein |
| 6,378 | A→G | A32A (GCA→GCG) | 5.2% | 8.5% | Phage protein |
| 6,381 | C→A | R33R (CGC→CGA) | 5.1% | 9.4% | Phage protein |
| 6,384 | C→T | N34N (AAC→AAT) | | 8.8% | Phage protein |
| 6,393 | G→C | A37A (GCG→GCC) | | 10.9% | Phage protein |
| 6,396 | T→C | F38F (TTT→TTC) | | 10.2% | Phage protein |
| 6,405 | A→T | T41T (ACA→ACT) | | 10.9% | Phage protein |
| 6,408 | A→G | E42E (GAA→GAG) | | 10.2% | Phage protein |
| 6,417 | T→C | F45F (TTT→TTC) | | 9.2% | Phage protein |
| 6,426 | T→C | R48R (CGT→CGC) | 5.0% | 9.2% | Phage protein |
| 6,432 | T→C | N50N (AAT→AAC) | | 9.2% | Phage protein |
| 6,433 | T→G | L51V (TTG→GTG) | | 8.7% | Phage protein |
| 6,436 | G→A | V52I (GTA→ATA) | | 8.4% | Phage protein |
| 6,438 | A→C | V52V (GTA→GTC) | | 8.6% | Phage protein |
| 6,441 | C→T | A53A (GCC→GCT) | | 8.4% | Phage protein |
| 6,489 | A→G | V69V (GTA→GTG) | | 9.5% | Phage protein |
| 6,607 | A→T | *38L (TAA→TTA)<br>V2V (GTA→GTT) | 5.0% | 9.5% | Phage protein |
| 6,622 | C→T | N7N (AAC→AAT) | | 8.6% | Phage protein |
| 6,625 | A→T | R8R (CGA→CGT) | | 8.5% | Phage protein |
| 6,631 | A→T | V10V (GTA→GTT) | | 8.7% | Phage protein |
| 6,632 | A→G | T11A (ACT→GCT) | 5.2% | 8.9% | Phage protein |
| 6,667 | A→G | L22L (TTA→TTG) | 32.3% | 15.4% | Phage protein |
| 6,67 | C→T | N23N (AAC→AAT) | 31.6% | 15.1% | Phage protein |

| | | | | | |
|---|---|---|---|---|---|
| 6,679 | C→A | S26S (TCC→TCA) | 32.2% | 15.7% | Phage protein |
| 6,718 | A→G | K39K (AAA→AAG) | 29.3% | 13.9% | Phage protein |
| 6,721 | A→G | A40A (GCA→GCG) | 29.1% | 14.4% | Phage protein |
| 6,733 | T→C | I44I (ATT→ATC) | 23.6% | 11.1% | Phage protein |
| 6,734 | T→A | S45T (TCT→ACT) | 23.3% | 11.0% | Phage protein |
| 6,735 | C→A | S45Y (TCT→TAT) | 23.5% | 11.1% | Phage protein |
| 6,739 | C→G | A46A (GCC→GCG) | 23.7% | 11.5% | Phage protein |
| 6,742 | T→C | V47V (GTT→GTC) | 23.4% | 11.2% | Phage protein |
| 6,745 | C→T | V48V (GTC→GTT) | 23.4% | 11.2% | Phage protein |
| 6,755 | T→C | L52L (TTA→CTA) | 28.4% | 15.9% | Phage protein |
| 6,76 | A→G | A53A (GCA→GCG) | 28.8% | 15.9% | Phage protein |
| 6,856 | A→G | K85K (AAA→AAG) | 23.1% | 12.6% | Phage protein |
| 6,857 | G→A | V86I (GTA→ATA) | 22.5% | 11.9% | Phage protein |
| 6,862 | T→C | C87C (TGT→TGC) | 23.8% | 13.2% | Phage protein |
| 6,865 | A→C | T88T (ACA→ACC) | 24.2% | 14.0% | Phage protein |
| 6,877 | T→A | I92I (ATT→ATA) | 23.6% | 13.6% | Phage protein |
| 6,898 | G→A | Q99Q (CAG→CAA) | 17.7% | 5.4% | Phage protein |
| 6,901 | A→C | S100S (TCA→TCC) | 19.0% | 6.0% | Phage protein |
| 6,91 | A→T | E103D (GAA→GAT) | 17.6% | | Phage protein |
| 6,922 | G→A | T107T (ACG→ACA) | 13.9% | | Phage protein |
| 7,248 | T→A | G21G (GGA→GGT) | 7.1% | | hypothetical protein |
| 7,249 | C→T | G21E (GGA→GAA) | 6.7% | | hypothetical protein |
| 7,255 | A→T | I19N (ATT→AAT) | 21.8% | | hypothetical protein |
| 7,279 | T→G | H11P (CAT→CCT) | 25.8% | 6.3% | hypothetical protein |
| 7,29 | A→G | A7A (GCT→GCC) | 26.6% | 9.4% | hypothetical protein |
| 7,291 | G→C | A7G (GCT→GGT) | 26.4% | 9.5% | hypothetical protein |
| 7,302 | G→A | N3N (AAC→AAT) | 30.8% | 10.5% | hypothetical protein |
| 7,33 | A→T | intergenic (-20/+304) | 32.8% | 12.2% | hypothetical protein |
| 7,331 | A→T | intergenic (-21/+303) | 34.0% | 13.1% | hypothetical protein |
| 7,336 | T→C | intergenic (-26/+298) | 34.3% | 14.3% | hypothetical protein |
| 7,342 | A→G | intergenic (-32/+292) | 34.5% | 15.4% | hypothetical protein |
| 7,347 | C→G | intergenic (-37/+287) | 34.4% | 15.8% | hypothetical protein |
| 7,352 | T→G | intergenic (-42/+282) | 34.9% | 16.8% | hypothetical protein |
| 7,365 | G→A | intergenic (-55/+269) | 35.6% | 16.4% | hypothetical protein |
| 7,381 | C→G | intergenic (-71/+253) | 37.2% | 20.3% | hypothetical protein |
| 7,391 | T→C | intergenic (-81/+243) | 37.9% | 19.6% | hypothetical protein |
| 7,398 | G→A | intergenic (-88/+236) | 38.0% | 21.6% | hypothetical protein |
| 7,419 | A→G | intergenic (-109/+215) | 37.6% | 23.2% | hypothetical protein |
| 7,42 | C→T | intergenic (-110/+214) | 36.8% | 22.6% | hypothetical protein |
| 7,431 | A→C | intergenic (-121/+203) | 38.9% | 24.2% | hypothetical protein |
| 7,439 | G→A | intergenic (-129/+195) | 35.9% | 23.0% | hypothetical protein |
| 7,456 | G→C | intergenic (-146/+178) | 31.2% | 18.5% | hypothetical protein |
| 7,457 | C→A | intergenic (-147/+177) | 31.2% | 18.3% | hypothetical protein |
| 7,46 | T→C | intergenic (-150/+174) | 31.6% | 19.2% | hypothetical protein |
| 7,463 | G→A | intergenic (-153/+171) | 30.7% | 18.2% | hypothetical protein |
| 7,468 | C→A | intergenic (-158/+166) | 31.0% | 17.9% | hypothetical protein |
| 7,469 | A→G | intergenic (-159/+165) | 31.1% | 17.9% | hypothetical protein |
| 7,477 | C→A | intergenic (-167/+157) | 31.4% | 18.9% | hypothetical protein |
| 7,489 | T→C | intergenic (-179/+145) | 28.1% | 15.5% | hypothetical protein |
| 7,512 | C→T | intergenic (-202/+122) | 17.6% | 10.0% | hypothetical protein |
| 7,514 | Δ1 bp | intergenic (-204/+120) | 17.5% | 9.6% | hypothetical protein |
| 7,518:1 | +C | intergenic (-208/+116) | 17.2% | 9.5% | hypothetical protein |
| 7,544 | A→G | intergenic (-234/+90) | 10.1% | | hypothetical protein |

| | | | | | | |
|---|---|---|---|---|---|---|
| 7,578 | G→A | intergenic (-268/+56) | | 6.6% | | hypothetical protein |
| 7,622 | 25 bp→AC | | | 16.9% | 8.1% | hypothetical protein |
| 7,677 | T→G | Y63S (TAT→TCT) | | 25.4% | 13.5% | hypothetical protein |
| 7,68 | T→A | H62L (CAT→CTT) | | 24.3% | 13.6% | hypothetical protein |
| 7,681 | G→C | H62D (CAT→GAT) | | 24.8% | 13.4% | hypothetical protein |
| 7,685 | C→A | E60D (GAG→GAT) | | 24.0% | 13.4% | hypothetical protein |
| 7,741 | Δ1 bp | coding (124/231 nt) | | 44.1% | 11.0% | hypothetical protein |
| 7,748:1 | +G | coding (117/231 nt) | | 42.6% | 10.4% | hypothetical protein |
| 7,748:2 | +T | coding (117/231 nt) | | 41.9% | 10.4% | hypothetical protein |
| 7,749 | A→G | I39T (ATC→ACC) | | 43.9% | 11.3% | hypothetical protein |
| 7,755 | C→T | R37H (CGC→CAC) | | 43.9% | 10.2% | hypothetical protein |
| 7,757 | T→C | P36P (CCA→CCG) | | 45.1% | 11.5% | hypothetical protein |
| 7,762 | A→G | C35R (TGC→CGC) | | 50.2% | 14.7% | hypothetical protein |
| 7,772 | A→G | F31F (TTT→TTC) | | 51.9% | 14.4% | hypothetical protein |
| 7,809 | G→C | S19W (TCG→TGG) | | 11.7% | 9.2% | hypothetical protein |
| 7,814 | A→T | N17K (AAT→AAA) | | 10.7% | 9.1% | hypothetical protein |
| 7,819 | T→G | K16Q (AAA→CAA) | | 8.9% | 9.4% | hypothetical protein |
| 7,82 | A→G | Y15Y (TAT→TAC) | | 9.0% | 9.7% | hypothetical protein |
| 7,824 | A→T | L14H (CTC→CAC) | | 8.6% | 8.8% | hypothetical protein |
| 7,867 | A→T | intergenic (-3/+57) | | | 23.0% | hypothetical protein |
| 8,354 | A→T | D28V (GAC→GTC) | | | 5.7% | hypothetical protein |
| 8,558 | C→A | intergenic (+197/-90) | | 7.3% | | hypothetical protein |
| 8,56 | Δ1 bp | intergenic (+199/-88) | | 7.4% | | hypothetical protein |
| 8,576 | C→T | intergenic (+215/-72) | | 7.8% | | hypothetical protein |
| 8,579 | Δ1 bp | intergenic (+218/-69) | | 8.3% | | hypothetical protein |
| 8,582 | T→A | intergenic (+221/-66) | | 8.4% | | hypothetical protein |
| 8,587 | C→A | intergenic (+226/-61) | | 8.3% | | hypothetical protein |
| 8,589 | A→G | intergenic (+228/-59) | | 8.6% | | hypothetical protein |
| 8,59 | T→G | intergenic (+229/-58) | | 8.8% | | hypothetical protein |
| 8,595 | C→A | intergenic (+234/-53) | | 8.3% | | hypothetical protein |
| 8,599 | A→T | intergenic (+238/-49) | | 8.8% | | hypothetical protein |
| 8,603 | G→C | intergenic (+242/-45) | | 8.6% | | hypothetical protein |
| 8,608 | T→A | intergenic (+247/-40) | | 8.5% | | hypothetical protein |
| 8,611 | A→T | intergenic (+250/-37) | | 9.2% | | hypothetical protein |
| 8,615 | G→A | intergenic (+254/-33) | | 8.4% | | hypothetical protein |
| 8,879 | G→A | V9I (GTT→ATT) | 33.7% | | | hypothetical protein |
| 8,9 | G→A | D16N (GAT→AAT) | 42.0% | | 15.8% | hypothetical protein |
| 8,903 | C→A | L17I (CTT→ATT) | 41.1% | | 14.8% | hypothetical protein |
| 8,907 | C→T | T18I (ACA→ATA)<br>L295L (CTG→CTA) | 40.3% | | 14.6% | Phage protein |
| 8,909 | G→A | V19I (GTA→ATA)<br>L295L (CTG→TTG) | 39.0% | | 14.6% | Phage protein |
| 8,943 | G→A | C30Y (TGT→TAT)<br>N283N (AAC→AAT) | 64.3% | | 18.9% | Phage protein |
| 8,945 | T→C | *31Q (TAG→CAG)<br>N283D (AAC→GAC) | 65.6% | | 19.0% | Phage protein |
| 8,964 | A→T | I276I (ATT→ATA) | 57.1% | | 31.3% | Phage protein |
| 8,966 | T→A | I276F (ATT→TTT) | 57.4% | | 31.5% | Phage protein |
| 8,967 | A→C | A275A (GCT→GCG) | 58.4% | | 33.0% | Phage protein |
| 8,97 | T→A | I274I (ATA→ATT) | 58.6% | | 32.1% | Phage protein |
| 8,975 | A→G | L273L (TTA→CTA) | 64.6% | | 32.7% | Phage protein |
| 8,982 | T→C | T270T (ACA→ACG) | 69.6% | | 33.8% | Phage protein |
| 8,985 | G→T | I269I (ATC→ATA) | 69.6% | | 33.3% | Phage protein |
| 8,997 | T→C | T265T (ACA→ACG) | 66.7% | | | Phage protein |

| | | | | | | |
|---|---|---|---|---|---|---|
| 9,039 | C→T | R251R (AGG→AGA) | 65.7% | 20.6% | 17.1% | Phage protein |
| 9,045 | G→T | I249I (ATC→ATA) | 61.5% | 19.2% | 16.4% | Phage protein |
| 9,048 | A→G | F248F (TTT→TTC) | 61.1% | 19.5% | 15.9% | Phage protein |
| 9,272 | C→T | V174I (GTA→ATA) | | | 21.4% | Phage protein |
| 9,303 | T→C | R163R (AGA→AGG) | | | 7.7% | Phage protein |
| 9,308 | T→C | I162V (ATT→GTT) | | | 6.8% | Phage protein |
| 9,311:1 | +T | coding (481/888 nt) | | | 7.0% | Phage protein |
| 9,315 | Δ1 bp | coding (477/888 nt) | | | 6.7% | Phage protein |
| 9,32 | A→T | C158S (TGT→AGT) | | | 5.9% | Phage protein |
| 9,322 | T→G | N157T (AAT→ACT) | | | 6.8% | Phage protein |
| 9,324 | T→A | P156P (CCA→CCT) | | | 6.9% | Phage protein |
| 9,327 | A→G | Y155Y (TAT→TAC) | | | 6.7% | Phage protein |
| 9,333 | T→C | R153R (AGA→AGG) | | | 7.8% | Phage protein |
| 9,339 | G→T | A151A (GCC→GCA) | | | 7.3% | Phage protein |
| 9,345 | A→G | H149H (CAT→CAC) | | | 7.9% | Phage protein |
| 9,351 | C→T | P147P (CCG→CCA) | | | 6.1% | Phage protein |
| 9,354 | G→A | S146S (AGC→AGT) | | | 5.9% | Phage protein |
| 9,36 | A→T | S144S (TCT→TCA) | | | 5.4% | Phage protein |
| 9,363 | A→G | Y143Y (TAT→TAC) | | | 6.8% | Phage protein |
| 9,663 | T→C | K43K (AAA→AAG) | | | 5.4% | Phage protein |
| 9,681 | G→A | T37T (ACC→ACT) | | | 8.6% | Phage protein |
| 9,683 | T→A | T37S (ACC→TCC) | | | 9.0% | Phage protein |
| 9,684 | T→C | L36L (CTA→CTG) | | | 8.5% | Phage protein |
| 9,69 | A→T | P34P (CCT→CCA) | | | 9.5% | Phage protein |
| 9,693 | G→A | F33F (TTC→TTT) | | | 9.0% | Phage protein |
| 9,696 | A→T | P32P (CCT→CCA) | | | 9.6% | Phage protein |
| 9,704 | T→C | I30V (ATA→GTA) | | | 11.6% | Phage protein |
| 9,708 | A→G | A28A (GCT→GCC) | | | 10.4% | Phage protein |
| 9,719 | T→C | I25V (ATT→GTT) | | | 13.2% | Phage protein |
| 9,726 | G→A | N22N (AAC→AAT) | | | 11.7% | Phage protein |
| 9,737 | C→G | A19P (GCT→CCT) | | | 15.2% | Phage protein |
| 9,742 | C→G | G17A (GGA→GCA) | | | 14.0% | Phage protein |
| 9,743 | C→T | G17R (GGA→AGA) | | | 13.3% | Phage protein |
| 9,762 | C→T | L10L (CTG→CTA) | | | 14.0% | Phage protein |
| 9,768 | G→A | D8D (GAC→GAT) | | | 10.4% | Phage protein |
| 9,777 | A→G | N5N (AAT→AAC) | | | 8.6% | Phage protein |
| 9,783 | Δ1 bp | coding (9/888 nt) | | | 5.7% | Phage protein |
| 9,784 | Δ1 bp | coding (8/888 nt) | | | 5.7% | Phage protein |
| 9,785 | T→C | T3A (ACA→GCA) | | | 5.5% | Phage protein |
| 9,787 | Δ1 bp | coding (5/888 nt) | | | 5.6% | Phage protein |
| 9,789 | C→T | M1M (ATG→ATA) † | | | 5.5% | Phage protein |
| 9,793 | G→A | intergenic (-2/+68) | | | 5.6% | Phage protein |
| 9,795 | G→C | intergenic (-4/+66) | | | 5.8% | Phage protein |
| 9,797 | T→C | intergenic (-6/+64) | | | 5.5% | Phage protein |
| 9,798 | T→C | intergenic (-7/+63) | | | 5.5% | Phage protein |
| 9,8 | C→G | intergenic (-9/+61) | | | 5.8% | Phage protein |
| 9,811:1 | +T | intergenic (-20/+50) | | | 5.9% | Phage protein |
| 9,812 | A→G | intergenic (-21/+49) | | | 5.6% | Phage protein |
| 9,821 | G→T | intergenic (-30/+40) | | | 6.0% | Phage protein |
| 9,834 | A→G | intergenic (-43/+27) | | | 6.1% | Phage protein |
| 10,433 | C→T | R96R (CGG→CGA) | | | 6.1% | Phage protein |
| 10,442 | C→T | V93V (GTG→GTA) | | | 6.4% | Phage protein |
| 10,445 | T→A | P92P (CCA→CCT) | | | 6.1% | Phage protein |

| | | | | | |
|---|---|---|---|---|---|
| 10,46 | A→G | N87N (AAT→AAC) | | 9.8% | Phage protein |
| 10,478 | G→A | L81L (CTC→CTT) | | 9.1% | Phage protein |
| 10,487 | T→C | G78G (GGA→GGG) | | 9.1% | Phage protein |
| 10,49 | A→T | A77A (GCT→GCA) | | 8.3% | Phage protein |
| 10,501 | G→A | L74L (CTA→TTA) | | 6.4% | Phage protein |
| 10,504 | C→T | A73T (GCA→ACA) | | 6.6% | Phage protein |
| 10,505 | A→G | N72N (AAT→AAC) | | 7.5% | Phage protein |
| 10,514 | A→G | N69N (AAT→AAC) | | 7.9% | Phage protein |
| 10,529 | A→G | R64R (CGT→CGC) | | 8.8% | Phage protein |
| 10,628 | A→G | F31F (TTT→TTC) | | 6.1% | Phage protein |
| 10,631 | C→T | Q30Q (CAG→CAA) | | 6.0% | Phage protein |
| 10,637 | T→A | T28T (ACA→ACT) | | 6.1% | Phage protein |
| 10,646 | C→T | L25L (CTG→CTA) | | 7.7% | Phage protein |
| 10,648 | G→A | L25L (CTG→TTG) | | 7.9% | Phage protein |
| 10,661 | G→A | Y20Y (TAC→TAT) | | 12.1% | Phage protein |
| 10,675 | C→T | D16N (GAC→AAC) | | 13.4% | Phage protein |
| 10,676 | G→C | L15L (CTC→CTG) | | 12.4% | Phage protein |
| 10,685 | A→G | D12D (GAT→GAC) | | 11.3% | Phage protein |
| 10,688 | T→G | A11A (GCA→GCC) | | 11.7% | Phage protein |
| 10,69 | C→G | A11P (GCA→CCA) | | 12.2% | Phage protein |
| 10,692 | A→G | V10A (GTA→GCA) | | 12.0% | Phage protein |
| 10,699 | T→G | K8Q (AAA→CAA) | | 13.6% | Phage protein |
| 10,703 | G→A | T6T (ACC→ACT) | | 13.0% | Phage protein |
| 10,705 | T→G | T6P (ACC→CCC) | | 12.9% | Phage protein |
| 10,747 | C→T | K53K (AAG→AAA) | | 11.6% | Phage protein |
| 10,751 | C→T | R52H (CGT→CAT) | | 9.6% | Phage protein |
| 10,753 | A→G | A51A (GCT→GCC) | | 9.4% | Phage protein |
| 12,827 | G→A | I104I (ATC→ATT) | 17.4% | | Phage protein |
| 12,836 | A→G | N101N (AAT→AAC) | 17.2% | | Phage protein |
| 12,845 | G→A | F98F (TTC→TTT) | 14.5% | | Phage protein |
| 12,848 | G→A | D97D (GAC→GAT) | 14.1% | | Phage protein |
| 12,851 | A→G | H96H (CAT→CAC) | 14.4% | | Phage protein |
| 12,86 | A→T | H93Q (CAT→CAA) | 11.7% | | Phage protein |
| 12,863 | T→A | R92R (CGA→CGT) | 12.1% | | Phage protein |
| 12,866 | T→A | A91A (GCA→GCT) | 12.4% | | Phage protein |
| 12,869 | A→G | D90D (GAT→GAC) | 11.8% | | Phage protein |
| 12,875 | G→A | F88F (TTC→TTT) | 11.2% | | Phage protein |
| 12,884 | G→A | F85F (TTC→TTT) | 8.7% | | Phage protein |
| 12,887 | C→T | K84K (AAG→AAA) | 9.0% | | Phage protein |
| 12,893 | T→C | A82A (GCA→GCG) | 8.9% | | Phage protein |
| 12,907 | G→A | L78L (CTG→TTG) | 5.6% | | Phage protein |
| 12,914 | A→T | G75G (GGT→GGA) | 6.2% | | Phage protein |
| 12,971 | G→T | L56L (CTC→CTA) | 5.9% | 6.3% | Phage protein |
| 12,974 | A→G | D55D (GAT→GAC) | 5.9% | 6.5% | Phage protein |
| 12,98 | A→G | I53I (ATT→ATC) | 7.1% | 8.6% | Phage protein |
| 12,989 | G→A | N50N (AAC→AAT) | 8.0% | 16.6% | Phage protein |
| 12,995 | C→T | E48E (GAG→GAA) | 8.7% | 19.8% | Phage protein |
| 13,01 | C→T | K43K (AAG→AAA) | 9.6% | 21.8% | Phage protein |
| 13,013 | C→T | T42T (ACG→ACA) | 9.7% | 22.0% | Phage protein |
| 13,019 | A→G | H40H (CAT→CAC) | 10.3% | 22.5% | Phage protein |
| 13,022 | G→A | F39F (TTC→TTT) | 10.4% | 22.4% | Phage protein |
| 13,025 | A→C | T38T (ACT→ACG) | 11.1% | 23.0% | Phage protein |
| 13,026 | G→T | T38N (ACT→AAT) | 11.0% | 22.4% | Phage protein |

| | | | | | |
|---|---|---|---|---|---|
| 13,037 | G→T | R34R (CGC→CGA) | 15.1% | 45.2% | Phage protein |
| 13,043 | A→G | N32N (AAT→AAC) | 16.5% | 44.0% | Phage protein |
| 13,046 | A→G | F31F (TTT→TTC) | 15.4% | 44.9% | Phage protein |
| 13,049 | A→G | R30R (CGT→CGC) | 15.5% | 45.9% | Phage protein |
| 13,082 | T→G | L19F (TTA→TTC) | 20.7% | 48.6% | Phage protein |
| 13,084 | A→G | L19L (TTA→CTA) | 20.7% | 48.1% | Phage protein |
| 13,098 | C→T | G14D (GGC→GAC) | 19.1% | 51.5% | Phage protein |
| 13,099 | C→T | G14S (GGC→AGC) | 19.6% | 51.8% | Phage protein |
| 13,12 | A→G | L7L (TTA→CTA) | 21.5% | 48.1% | Phage protein |
| 13,123 | T→G | R6R (AGA→CGA) | 22.1% | 47.7% | Phage protein |
| 13,127 | C→T | L4L (CTG→CTA) | 20.9% | 48.3% | Phage protein |
| 13,156 | C→T | A60A (GCG→GCA) | 26.0% | 52.1% | Phage protein |
| 13,161 | T→C | M59V (ATG→GTG) | 25.8% | 49.7% | Phage protein |
| 13,162 | A→G | I58I (ATT→ATC) | 25.8% | 50.2% | Phage protein |
| 13,191 | A→G | L49L (TTA→CTA) | 25.4% | 49.8% | Phage protein |
| 13,192 | C→T | G48G (GGG→GGA) | 25.1% | 48.8% | Phage protein |
| 13,201 | A→G | Y45Y (TAT→TAC) | 27.9% | 49.0% | Phage protein |
| 13,207 | T→G | I43I (ATA→ATC) | 27.1% | 48.6% | Phage protein |
| 13,409 | T→C | intergenic (-74/+26) | 27.0% | 50.5% | Phage protein |
| 13,411 | C→A | intergenic (-76/+24) | 27.2% | 52.0% | Phage protein |
| 13,429 | G→A | intergenic (-94/+6) | 26.6% | 57.9% | Phage protein |
| 13,477 | A→G | S95S (TCT→TCC) | 24.9% | 54.1% | Phage protein |
| 13,484 | C→T | G93D (GGT→GAT) | 22.2% | 54.2% | Phage protein |
| 13,524 | C→T | D80N (GAT→AAT) | 19.4% | 48.2% | Phage protein |
| 13,525 | A→C | A79A (GCT→GCG) | 19.9% | 48.8% | Phage protein |
| 13,561 | G→A | T67T (ACC→ACT) | 19.0% | 25.1% | Phage protein |
| 13,564 | A→T | N66K (AAT→AAA) | 19.0% | 25.8% | Phage protein |
| 13,566 | T→C | N66D (AAT→GAT) | 19.2% | 26.0% | Phage protein |
| 13,567 | G→A | I65I (ATC→ATT) | 18.6% | 27.0% | Phage protein |
| 13,581 | T→C | I61V (ATT→GTT) | 20.8% | 25.7% | Phage protein |
| 13,585 | T→C | K59K (AAA→AAG) | 20.4% | 25.0% | Phage protein |
| 13,591 | A→T | N57K (AAT→AAA) | 20.2% | 24.2% | Phage protein |
| 13,597 | T→A | P55P (CCA→CCT) | 17.2% | 20.8% | Phage protein |
| 13,6 | C→T | T54T (ACG→ACA) | 16.2% | 17.8% | Phage protein |
| 13,603 | T→C | L53L (TTA→TTG) | 16.9% | 18.8% | Phage protein |
| 13,605 | A→G | L53L (TTA→CTA) | 17.3% | 18.7% | Phage protein |
| 13,607 | A→T | V52E (GTA→GAA) | 17.4% | 18.0% | Phage protein |
| 13,612 | A→G | L50L (CTT→CTC) | 17.2% | 18.2% | Phage protein |
| 13,615 | T→C | P49P (CCA→CCG) | 16.6% | 18.2% | Phage protein |
| 13,64 | A→T | I41N (ATT→AAT) | 21.2% | 21.6% | Phage protein |
| 13,656 | T→C | K36E (AAG→GAG) | 22.6% | 23.6% | Phage protein |
| 13,741 | C→T | E7E (GAG→GAA) | 28.2% | 30.0% | Phage protein |
| 13,744 | A→G | N6N (AAT→AAC) | 29.3% | 30.4% | Phage protein |
| 13,747 | G→A | F5F (TTC→TTT) | 28.9% | 30.6% | Phage protein |
| 13,768 | T→G | I76L (ATT→CTT) | 26.9% | 31.7% | Phage protein |
| 13,816 | T→G | S60R (AGC→CGC) | 20.1% | 22.4% | Phage protein |
| 13,818 | G→T | T59K (ACG→AAG) | 20.0% | 23.2% | Phage protein |
| 13,822 | T→C | I58V (ATT→GTT) | 20.8% | 24.2% | Phage protein |
| 13,826 | G→T | T56T (ACC→ACA) | 20.4% | 24.4% | Phage protein |
| 13,829 | C→T | K55K (AAG→AAA) | 19.6% | 23.7% | Phage protein |
| 13,832 | C→T | T54T (ACG→ACA) | 19.1% | 22.7% | Phage protein |
| 13,835 | C→T | V53V (GTG→GTA) | 19.1% | 21.7% | Phage protein |
| 13,852 | A→T | L48M (TTG→ATG) | 25.6% | 31.2% | Phage protein |

| | | | | | |
|---|---|---|---|---|---|
| 13,904 | C→T | V30V (GTG→GTA) | 24.2% | 35.4% | Phage protein |
| 13,911 | G→A | A28V (GCT→GTT) | 22.9% | 32.9% | Phage protein |
| 13,949 | C→T | L15L (CTG→CTA) | 25.8% | 41.7% | Phage protein |
| 13,958 | T→C | V12V (GTA→GTG) | 25.9% | 40.5% | Phage protein |
| 13,975 | C→A | V7F (GTT→TTT) | 27.1% | 42.1% | Phage protein |
| 13,976 | C→T | A6A (GCG→GCA) | 27.1% | 42.6% | Phage protein |
| 13,994 | +ATT | coding (278/282 nt) | 25.1% | 39.4% | Phage protein |
| 14,008 | A→G | I88I (ATT→ATC) | 27.0% | 36.1% | Phage protein |
| 14,083 | T→C | K63K (AAA→AAG) | 32.3% | 47.6% | Phage protein |
| 14,095 | T→C | S59S (TCA→TCG) | 31.0% | 46.7% | Phage protein |
| 14,104 | C→T | Q56Q (CAG→CAA) | 29.8% | 44.2% | Phage protein |
| 14,107 | T→G | A55A (GCA→GCC) | 31.7% | 44.3% | Phage protein |
| 14,112 | G→T | L54I (CTA→ATA) | 29.6% | 44.5% | Phage protein |
| 14,14 | G→A | R44R (CGC→CGT) | 27.6% | 41.3% | Phage protein |
| 14,174 | T→C | N33S (AAT→AGT) | 25.2% | 43.0% | Phage protein |
| 14,202 | T→C | I24V (ATT→GTT) | 16.0% | 38.8% | Phage protein |
| 14,203 | C→T | G23G (GGG→GGA) | 16.5% | 40.2% | Phage protein |
| 14,23 | T→C | T14T (ACA→ACG) | 9.7% | 34.3% | Phage protein |
| 14,233 | C→A | M13I (ATG→ATT) | 9.3% | 33.2% | Phage protein |
| 14,234 | A→G | M13T (ATG→ACG) | 9.6% | 33.2% | Phage protein |
| 14,239 | C→A | V11V (GTG→GTT) | 9.5% | 33.5% | Phage protein |
| 14,241 | C→T | V11M (GTG→ATG) | 9.1% | 32.3% | Phage protein |
| 14,245 | C→T | L9L (TTG→TTA) | 8.9% | 31.3% | Phage protein |
| 14,248 | T→C | G8G (GGA→GGG) | 9.2% | 30.6% | Phage protein |
| 14,254 | G→A | L6L (CTC→CTT) | 9.3% | 30.9% | Phage protein |
| 14,264 | G→A | T3I (ACT→ATT) | 6.9% | 23.9% | Phage protein |
| 14,265 | T→C | T3A (ACT→GCT) | 8.0% | 25.3% | Phage protein |
| 14,266 | T→C | L2L (TTA→TTG) | 7.5% | 24.1% | Phage protein |
| 14,277 | G→A | G53G (GGC→GGT) | | 16.4% | Phage protein |
| 14,278 | C→T | G53D (GGC→GAC) | | 17.0% | Phage protein |
| 14,281 | C→T | R52K (AGG→AAG) | | 15.8% | Phage protein |
| 14,292 | G→A | N48N (AAC→AAT) | | 11.8% | Phage protein |
| 14,295 | A→G | N47N (AAT→AAC) | | 11.8% | Phage protein |
| 14,348 | C→G | V30L (GTA→CTA) | | 8.5% | Phage protein |
| 14,352 | A→T | T28T (ACT→ACA) | | 7.5% | Phage protein |
| 14,354 | T→C | T28A (ACT→GCT) | | 8.8% | Phage protein |
| 14,358 | G→A | R26R (CGC→CGT) | | 8.4% | Phage protein |
| 14,366 | T→G | N24H (AAC→CAC) | | 16.7% | Phage protein |
| 14,376 | C→T | M20I (ATG→ATA) | | 21.4% | Phage protein |
| 14,38 | T→C | N19S (AAT→AGT) | | 23.5% | Phage protein |
| 14,39 | G→A | L16L (CTA→TTA) | | 20.7% | Phage protein |
| 14,391 | T→C | T15T (ACA→ACG) | | 22.0% | Phage protein |
| 14,392 | G→A | T15I (ACA→ATA) | | 20.9% | Phage protein |
| 14,393 | T→C | T15A (ACA→GCA) | | 22.2% | Phage protein |
| 14,399 | T→A | I13L (ATA→TTA) | | 21.6% | Phage protein |
| 14,405 | A→G | F11L (TTC→CTC) | | 21.7% | Phage protein |
| 14,409 | C→T | L9L (CTG→CTA) | | 21.5% | Phage protein |
| 14,412 | T→C | I8M (ATA→ATG) | | 22.1% | Phage protein |
| 14,415 | T→A | S7S (TCA→TCT) | | 19.9% | Phage protein |
| 14,416 | G→A | S7L (TCA→TTA) | | 19.5% | Phage protein |
| 14,421 | T→G | S5S (TCA→TCC) | 6.8% | 23.8% | Phage protein |
| 14,426 | C→T | V4I (GTA→ATA) | 6.4% | 26.8% | Phage protein |
| 14,456 | T→C | K77K (AAA→AAG) | 12.1% | 36.2% | Phage protein |

| | | | | | |
|---|---|---|---|---|---|
| 14,492 | T→G | A65A (GCA→GCC) | 14.0% | 33.3% | Phage protein |
| 14,495 | C→A | V64V (GTG→GTT) | 13.9% | 33.5% | Phage protein |
| 14,496 | A→G | V64A (GTG→GCG) | 14.1% | 33.8% | Phage protein |
| 14,497 | C→T | V64M (GTG→ATG) | 13.6% | 31.6% | Phage protein |
| 14,501 | T→C | Q62Q (CAA→CAG) | 15.0% | 32.9% | Phage protein |
| 14,51 | C→T | K59K (AAG→AAA) | 16.8% | 39.7% | Phage protein |
| 14,519 | G→A | R56R (CGC→CGT) | 16.5% | 43.5% | Phage protein |
| 14,525 | C→T | E54E (GAG→GAA) | 17.3% | 45.1% | Phage protein |
| 14,537 | T→A | L50F (TTA→TTT) | 17.5% | 48.8% | Phage protein |
| 14,539 | A→G | L50L (TTA→CTA) | 18.0% | 47.8% | Phage protein |
| 14,543 | A→G | F48F (TTT→TTC) | 17.6% | 47.5% | Phage protein |
| 14,546 | G→T | A47A (GCC→GCA) | 18.2% | 47.4% | Phage protein |
| 14,555 | G→A | N44N (AAC→AAT) | 19.8% | 49.0% | Phage protein |
| 14,561 | G→A | N42N (AAC→AAT) | 17.9% | 47.1% | Phage protein |
| 14,564 | A→C | L41L (CTT→CTG) | 18.5% | 48.7% | Phage protein |
| 14,567 | A→G | S40S (TCT→TCC) | 19.1% | 50.9% | Phage protein |
| 14,573 | G→A | Y38Y (TAC→TAT) | 20.4% | 48.9% | Phage protein |
| 14,591 | T→C | P32P (CCA→CCG) | 20.7% | 52.5% | Phage protein |
| 14,618 | T→C | E23E (GAA→GAG) | 20.3% | 53.3% | Phage protein |
| 14,624 | A→T | G21G (GGT→GGA) | 21.2% | 56.4% | Phage protein |
| 14,627 | C→T | Q20Q (CAG→CAA) | 20.5% | 56.7% | Phage protein |
| 14,667 | C→T | S7N (AGT→AAT) | 20.6% | 55.0% | Phage protein |
| 14,78 | A→G | V139V (GTT→GTC) | 24.5% | 49.2% | Phage protein |
| 14,782 | C→A | V139F (GTT→TTT) | 24.7% | 48.9% | Phage protein |
| 14,786 | G→T | S137S (TCC→TCA) | 24.4% | 47.5% | Phage protein |
| 14,843 | G→A | C118C (TGC→TGT) | 24.5% | 44.2% | Phage protein |
| 14,849 | C→T | L116L (CTG→CTA) | 22.8% | 44.0% | Phage protein |
| 14,851 | G→A | L116L (CTG→TTG) | 22.3% | 42.8% | Phage protein |
| 14,852 | T→C | E115E (GAA→GAG) | 23.2% | 43.9% | Phage protein |
| 14,861 | T→C | A112A (GCA→GCG) | 26.2% | 47.0% | Phage protein |
| 14,872 | T→C | T109A (ACT→GCT) | 25.9% | 49.2% | Phage protein |
| 14,921 | A→T | P92P (CCT→CCA) | | 50.8% | Phage protein |
| 14,93 | C→G | Q89H (CAG→CAC) | | 50.0% | Phage protein |
| 14,98 | T→C | K73E (AAG→GAG) | | 48.5% | Phage protein |
| 15,04 | A→G | L53L (TTG→CTG) | | 49.4% | Phage protein |
| 15,047 | G→A | C50C (TGC→TGT) | | 46.0% | Phage protein |
| 15,11 | A→G | S29S (TCT→TCC) | | 40.8% | Phage protein |
| 15,116 | G→A | F27F (TTC→TTT) | | 39.3% | Phage protein |
| 15,127 | G→T | R24R (CGA→AGA) | | 42.6% | Phage protein |
| 15,185 | T→A | V4V (GTA→GTT) | | 46.6% | Phage protein |
| 15,201 | T→A | intergenic (-5/+48) | | 48.4% | Phage protein |
| 15,206 | C→T | intergenic (-10/+43) | | 48.2% | Phage protein |
| 15,222 | A→C | intergenic (-26/+27) | | 49.0% | Phage protein |
| 15,243 | G→A | intergenic (-47/+6) | | 48.7% | Phage protein |
| 15,281 | G→A | L31L (CTA→TTA) | | 50.5% | hypothetical protein |
| 15,288 | A→G | G28G (GGT→GGC) | | 52.0% | hypothetical protein |
| 15,293 | T→C | I27V (ATT→GTT) | | 50.6% | hypothetical protein |
| 15,323 | C→T | V17I (GTA→ATA) | | 47.3% | hypothetical protein |
| 15,324 | T→C | A16A (GCA→GCG) | | 48.7% | hypothetical protein |
| 15,327 | T→C | Q15Q (CAA→CAG) | | 48.2% | hypothetical protein |
| 15,349 | G→A | A8V (GCT→GTT) | | 46.5% | hypothetical protein |
| 15,354 | G→A | A6A (GCC→GCT) | | 47.7% | hypothetical protein |
| 15,367 | T→G | Y2S (TAT→TCT) | | 48.4% | hypothetical protein |

| | | | | | | | |
|---|---|---|---|---|---|---|---|
| | | V196V (GTA→GTC) | | | | | Phosphoesterase |
| 15,385 | T→C | L190L (TTA→TTG) | | | 49.5% | | Phosphoesterase |
| 15,4 | A→G | D185D (GAT→GAC) | | | 50.4% | | Phosphoesterase |
| 15,406 | C→T | G183G (GGG→GGA) | | | 50.4% | | Phosphoesterase |
| 15,421 | T→C | E178E (GAA→GAG) | | | 61.6% | | Phosphoesterase |
| 15,442 | T→C | V171V (GTA→GTG) | | | 59.4% | | Phosphoesterase |
| 15,451 | A→G | Y168Y (TAT→TAC) | | | 56.4% | | Phosphoesterase |
| 15,469 | T→A | S162S (TCA→TCT) | | | 58.9% | | Phosphoesterase |
| 15,472 | T→C | V161V (GTA→GTG) | | | 59.4% | | Phosphoesterase |
| 15,481 | T→C | E158E (GAA→GAG) | | | 57.9% | | Phosphoesterase |
| 15,487 | A→T | P156P (CCT→CCA) | | | 56.2% | | Phosphoesterase |
| 15,493 | A→T | P154P (CCT→CCA) | | | 56.2% | | Phosphoesterase |
| 15,5 | T→C | K152R (AAA→AGA) | | | 56.6% | | Phosphoesterase |
| 15,502 | G→A | S151S (AGC→AGT) | | | 55.0% | | Phosphoesterase |
| 15,523 | G→A | I144I (ATC→ATT) | | | 54.3% | | Phosphoesterase |
| 15,526 | G→A | Y143Y (TAC→TAT) | | | 53.6% | | Phosphoesterase |
| 15,532 | A→T | P141P (CCT→CCA) | | | 52.3% | | Phosphoesterase |
| 15,543 | T→A | I138L (ATA→TTA) | | | 51.8% | | Phosphoesterase |
| 15,552 | C→T | A135T (GCT→ACT) | | | 50.9% | | Phosphoesterase |
| 15,559 | G→A | H132H (CAC→CAT) | | | 55.8% | | Phosphoesterase |
| 15,576 | A→G | L127L (TTA→CTA) | | | 54.8% | | Phosphoesterase |
| 15,583 | C→A | R124R (CGG→CGT) | | | 53.5% | | Phosphoesterase |
| 15,589 | C→T | E122E (GAG→GAA) | | | 54.8% | | Phosphoesterase |
| 15,598 | G→A | H119H (CAC→CAT) | | | 56.7% | | Phosphoesterase |
| 15,61 | A→G | H115H (CAT→CAC) | | | 54.5% | | Phosphoesterase |
| 15,769 | T→C | K62K (AAA→AAG) | | | 56.7% | | Phosphoesterase |
| 15,95 | C→T | R2K (AGG→AAG) | | | 61.8% | | Phosphoesterase |
| 16,02 | G→T | P74P (CCC→CCA) | | | 63.5% | | hypothetical protein |
| 16,057 | A→T | F62Y (TTC→TAC) | | | 62.2% | | hypothetical protein |
| 26,229 | T→C | D81G (GAT→GGT) | | | | 5.2% | Phage protein |
| 27,635 | T→C | E49G (GAA→GGA) | | | | 17.2% | 2-ketobutyrate formate-lyase (EC 2.3.1.-) |
| 28,879 | A→G | I16I (ATT→ATC) | | 5.1% | | | Phage protein |
| 29,748 | T→A | E5D (GAA→GAT) | | 7.0% | | | hypothetical protein |
| 29,841 | C→A | intergenic (-79/+29) | | 5.6% | | | hypothetical protein |
| 29,86 | C→T | intergenic (-98/+10) | | 5.2% | | | hypothetical protein |
| 29,9 | G→A | G29G (GGC→GGT) | | 5.6% | | | hypothetical protein |
| 30,556 | A→G | intergenic (-38/+63) | | 5.6% | | | hypothetical protein |
| 30,748 | T→C | R17R (CGA→CGG) | | 5.2% | | | hypothetical protein |
| 30,799 | C→T | intergenic (-1/+16) | | 5.2% | | | hypothetical protein/Phage protein |
| 31,097 | C→A | intergenic (-19/+187) | | 5.6% | | | Phage protein |
| 31,33 | G→A | P15L (CCC→CTC) | | 5.3% | | | hypothetical protein |
| 32,231 | G→A | M54I (ATG→ATA) | 18.0% | | | | hypothetical protein |
| 32,505 | G→A | Y41Y (TAC→TAT) | | 6.8% | | | Phage protein |
| 32,513 | T→C | I39V (ATA→GTA) | | 6.7% | | | Phage protein |
| 37,549 | C→T | A74T (GCA→ACA) | | 10.1% | | | Phage protein |
| 39,904 | T→A | P110P (CCA→CCT) | | 5.7% | | | Phage protein |
| 39,955 | A→G | V93V (GTT→GTC) | | 5.2% | | | Phage protein |
| 39,956 | A→G | V93A (GTT→GCT) | | 6.3% | | | Phage protein |
| 39,973 | G→A | T87T (ACC→ACT) | | 5.8% | | | Phage protein |
| 39,997 | G→A | A79A (GCC→GCT) | | 5.8% | | | Phage protein |
| 39,999 | C→A | A79S (GCC→TCC) | | 5.6% | | | Phage protein |
| 40 | C→T | V78V (GTG→GTA) | | 5.2% | | | Phage protein |

| | | | | |
|---|---|---|---|---|
| 40,042 | A→G | S64S (TCT→TCC) | 5.1% | Phage protein |
| 40,066 | A→T | V56V (GTT→GTA) | 5.4% | Phage protein |
| 40,44 | G→A | F39F (TTC→TTT) | 5.9% | Phage protein |
| 40,476 | T→G | V27V (GTA→GTC) | 5.9% | Phage protein |
| 40,684 | A→G | intergenic (-8/-174) | 6.6% | hypothetical protein |
| 40,841 | T→C | intergenic (-165/-17) | 5.0% | hypothetical protein |
| 41,179 | A→G | R108G (AGA→GGA) | 5.5% | hypothetical protein |
| 42,053 | T→C | K198K (AAA→AAG) | 6.5% | Metallopeptidase, phage-associated |
| 42,069 | T→A | K193I (AAA→ATA) | 5.5% | Metallopeptidase, phage-associated |
| 42,089 | A→G | A186A (GCT→GCC) | 5.4% | Metallopeptidase, phage-associated |
| 42,099 | T→G | D183A (GAT→GCT) | 5.6% | Metallopeptidase, phage-associated |
| 42,101 | T→C | K182K (AAA→AAG) | 5.7% | Metallopeptidase, phage-associated |
| 42,11 | A→G | Y179Y (TAT→TAC) | 6.1% | Metallopeptidase, phage-associated |
| 42,113 | A→T | G178G (GGT→GGA) | 6.1% | Metallopeptidase, phage-associated |
| 42,152 | G→A | G165G (GGC→GGT) | 5.1% | Metallopeptidase, phage-associated |
| 42,173 | C→T | P158P (CCG→CCA) | 6.4% | Metallopeptidase, phage-associated |
| 42,239 | C→T | P136P (CCG→CCA) | 6.6% | Metallopeptidase, phage-associated |
| 42,333 | G→T | T105N (ACT→AAT) | 6.3% | Metallopeptidase, phage-associated |
| 42,407 | A→G | I80I (ATT→ATC) | 5.3% | Metallopeptidase, phage-associated |
| 42,416 | C→G | P77P (CCG→CCC) | 6.1% | Metallopeptidase, phage-associated |
| 42,535 | T→C | I38V (ATT→GTT) | 5.5% | Metallopeptidase, phage-associated |
| 42,57 | G→A | A26V (GCA→GTA) | 5.0% | Metallopeptidase, phage-associated |
| 43,091 | G→A | G126G (GGC→GGT) | 5.0% | Phage protein |
| 43,133 | C→T | K112K (AAG→AAA) | 5.8% | Phage protein |
| 43,199 | A→T | A90A (GCT→GCA) | 6.8% | Phage protein |
| 43,221 | G→A | T83I (ACA→ATA) | 5.1% | Phage protein |
| 43,222 | T→C | T83A (ACA→GCA) | 5.9% | Phage protein |
| 43,244 | A→G | Y75Y (TAT→TAC) | 5.3% | Phage protein |
| 43,273 | T→C | T66A (ACC→GCC) | 6.7% | Phage protein |
| 43,289 | A→G | N60N (AAT→AAC) | 5.7% | Phage protein |
| 43,337 | C→T | Q44Q (CAG→CAA) | 5.3% | Phage protein |
| 43,34 | T→C | A43A (GCA→GCG) | 5.7% | Phage protein |
| 43,346 | T→C | Q41Q (CAA→CAG) | 6.2% | Phage protein |
| 43,351 | C→T | A40T (GCT→ACT) | 5.4% | Phage protein |
| 43,355 | A→G | T38T (ACT→ACC) | 5.8% | Phage protein |
| 43,367 | G→A | R34R (CGC→CGT) | 5.8% | Phage protein |
| 43,394 | G→A | R25R (CGC→CGT) | 7.4% | Phage protein |
| 43,412 | C→A | P19P (CCG→CCT) | 6.9% | Phage protein |
| 43,515 | A→G | intergenic (-47/+37) | 5.5% | Phage protein/Phage protein |
| 43,516 | G→T | intergenic (-48/+36) | 5.7% | Phage protein/Phage protein |
| 43,648 | T→C | E61E (GAA→GAG) | 5.7% | Phage protein |
| 43,657 | A→G | V58V (GTT→GTC) | 5.8% | Phage protein |
| 43,658 | A→G | V58A (GTT→GCT) | 6.2% | Phage protein |
| 43,669 | A→G | R54R (CGT→CGC) | 6.6% | Phage protein |
| 44,021 | A→G | H121H (CAT→CAC) | 5.9% | Phage ribonuclease H (EC 3.1.26.4) |
| 44,074 | A→G | L104L (TTA→CTA) | 5.2% | Phage ribonuclease H (EC 3.1.26.4) |
| 44,084 | A→G | V100V (GTT→GTC) | 5.5% | Phage ribonuclease H (EC 3.1.26.4) |
| 44,178 | A→G | V69A (GTT→GCT) | 5.0% | Phage ribonuclease H (EC 3.1.26.4) |
| 45,088 | G→T | S261S (TCC→TCA) | 5.1% | Thymidylate synthase (EC 2.1.1.45) |
| 45,1 | C→T | T257T (ACG→ACA) | 5.8% | Thymidylate synthase (EC 2.1.1.45) |
| 45,196 | C→T | E225E (GAG→GAA) | 5.7% | Thymidylate synthase (EC 2.1.1.45) |
| 45,22 | A→G | T217T (ACT→ACC) | 6.7% | Thymidylate synthase (EC 2.1.1.45) |
| 45,385 | A→G | C162C (TGT→TGC) | 6.1% | Thymidylate synthase (EC 2.1.1.45) |

| | | | | | | |
|---|---|---|---|---|---|---|
| 45,547 | T→C | V108V (GTA→GTG) | | 6.6% | | Thymidylate synthase (EC 2.1.1.45) |
| 45,55 | T→C | P107P (CCA→CCG) | | 6.4% | | Thymidylate synthase (EC 2.1.1.45) |
| 45,664 | G→A | L69L (CTC→CTT) | | 6.4% | | Thymidylate synthase (EC 2.1.1.45) |
| 45,736 | G→T | V45V (GTC→GTA) | | 5.2% | | Thymidylate synthase (EC 2.1.1.45) |
| 45,742 | A→C | P43P (CCT→CCG) | | 5.1% | | Thymidylate synthase (EC 2.1.1.45) |
| 45,748 | T→C | G41G (GGA→GGG) | | 5.4% | | Thymidylate synthase (EC 2.1.1.45) |
| 46,475 | G→A | A356A (GCC→GCT) | | 5.3% | | Ribonucleotide reductase of class Ia (aerobic), beta subunit (EC 1.17.4.1) |
| 46,513 | A→G | L344L (TTA→CTA) | | 5.1% | | Ribonucleotide reductase of class Ia (aerobic), beta subunit (EC 1.17.4.1) |
| 46,55 | G→A | P331P (CCC→CCT) | | 6.1% | | Ribonucleotide reductase of class Ia (aerobic), beta subunit (EC 1.17.4.1) |
| 46,559 | A→G | T328T (ACT→ACC) | | 6.2% | | Ribonucleotide reductase of class Ia (aerobic), beta subunit (EC 1.17.4.1) |
| 46,574 | A→G | H323H (CAT→CAC) | | 6.1% | | Ribonucleotide reductase of class Ia (aerobic), beta subunit (EC 1.17.4.1) |
| 46,595 | G→A | I316I (ATC→ATT) | | 6.3% | | Ribonucleotide reductase of class Ia (aerobic), beta subunit (EC 1.17.4.1) |
| 46,61 | A→G | N311N (AAT→AAC) | | 6.1% | | Ribonucleotide reductase of class Ia (aerobic), beta subunit (EC 1.17.4.1) |
| 46,631 | A→G | Y304Y (TAT→TAC) | | 5.4% | | Ribonucleotide reductase of class Ia (aerobic), beta subunit (EC 1.17.4.1) |
| 53,589 | T→A | F55Y (TTT→TAT) | | | 26.8% | Phage protein |
| 56,577 | G→T | L81F (TTG→TTT) | 5.6% | | | Phage protein |
| 57,321 | A→G | E10G (GAA→GGA) | 17.1% | | | hypothetical protein |
| 57,97 | T→A | V226V (GTT→GTA) | 67.6% | | | hypothetical protein |
| 57,973 | A→T | T227T (ACA→ACT) | 71.3% | | | hypothetical protein |
| 57,976 | A→G | Q228Q (CAA→CAG) | 68.9% | | | hypothetical protein |
| 57,982 | G→A | Q230Q (CAG→CAA) | 73.1% | | | hypothetical protein |
| 57,988 | G→A | K232K (AAG→AAA) | 73.7% | | | hypothetical protein |
| 58,009 | T→C | N239N (AAT→AAC) | 71.4% | | | hypothetical protein |
| 58,081 | C→T | Y263Y (TAC→TAT) | 82.5% | | | hypothetical protein |
| 58,093 | T→G | P267P (CCT→CCG) | 84.5% | | | hypothetical protein |
| 58,102 | T→C | T270T (ACT→ACC) | 85.7% | | | hypothetical protein |
| 58,114 | T→A | V274V (GTT→GTA) | 84.3% | | | hypothetical protein |
| 58,137 | C→T | A282V (GCT→GTT) | 83.8% | | | hypothetical protein |
| 58,18 | T→C | P296P (CCT→CCC) | 88.2% | | | hypothetical protein |
| 58,213 | G→T | G307G (GGG→GGT) | 85.4% | | | hypothetical protein |
| 58,225 | G→A | E311E (GAG→GAA) | 82.8% | | | hypothetical protein |
| 58,228 | A→G | E312E (GAA→GAG) | 83.4% | | | hypothetical protein |
| 58,246 | T→C | D318D (GAT→GAC) | 80.5% | | | hypothetical protein |
| 58,255 | A→G | K321K (AAA→AAG) | 75.1% | | | hypothetical protein |
| 58,258 | C→T | G322G (GGC→GGT) | 74.2% | | | hypothetical protein |
| 58,265 | A→G | I325V (ATT→GTT) | 71.7% | | | hypothetical protein |
| 58,266 | T→C | I325T (ATT→ACT) | 74.5% | | | hypothetical protein |
| 58,27 | T→C | A326A (GCT→GCC) | 73.4% | | | hypothetical protein |
| 58,285 | A→G | E331E (GAA→GAG) | 73.9% | | | hypothetical protein |
| 58,291 | C→T | T333T (ACC→ACT) | 67.3% | | | hypothetical protein |
| 58,294 | T→C | I334I (ATT→ATC) | 68.1% | | | hypothetical protein |
| 58,297 | C→T | S335S (AGC→AGT) | 67.2% | | | hypothetical protein |
| 58,301 | T→C | L337L (TTA→CTA) | 71.1% | | | hypothetical protein |
| 58,315 | G→T | E341D (GAG→GAT) | 75.4% | | | hypothetical protein |
| 58,321 | T→G | G343G (GGT→GGG) | 76.9% | | | hypothetical protein |
| 58,327 | T→C | Y345Y (TAT→TAC) | 77.6% | | | hypothetical protein |

| | | | | |
|---|---|---|---|---|
| 58,328 | C→T | L346L (CTA→TTA) | 76.4% | hypothetical protein |
| 58,372 | T→G | G360G (GGT→GGG) | 84.2% | hypothetical protein |
| 58,393 | G→A | E367E (GAG→GAA) | 81.3% | hypothetical protein |
| 58,396 | G→A | K368K (AAG→AAA) | 81.3% | hypothetical protein |
| 58,399 | G→A | E369E (GAG→GAA) | 82.1% | hypothetical protein |
| 58,405 | C→T | R371R (CGC→CGT) | 80.2% | hypothetical protein |
| 58,411 | T→C | A373A (GCT→GCC) | 82.4% | hypothetical protein |
| 58,417 | T→A | S375S (TCT→TCA) | 80.4% | hypothetical protein |
| 58,42 | G→A | E376E (GAG→GAA) | 81.5% | hypothetical protein |
| 58,459 | T→C | D389D (GAT→GAC) | 79.1% | hypothetical protein |
| 58,471 | C→T | N393N (AAC→AAT) | 75.4% | hypothetical protein |
| 58,477 | A→T | R395R (CGA→CGT) | 70.5% | hypothetical protein |
| 58,486 | A→T | A398A (GCA→GCT) | 66.5% | hypothetical protein |
| 58,487 | T→C | L399L (TTA→CTA) | 66.9% | hypothetical protein |
| 58,495 | C→A | A401A (GCC→GCA) | 66.3% | hypothetical protein |
| 58,507 | C→T | N405N (AAC→AAT) | 62.5% | hypothetical protein |
| 58,513 | G→A | K407K (AAG→AAA) | 60.7% | hypothetical protein |
| 58,519 | G→T | E409D (GAG→GAT) | 56.1% | hypothetical protein |
| 58,522 | C→A | I410I (ATC→ATA) | 56.3% | hypothetical protein |
| 58,525 | Δ1 bp | coding (1233/2937 nt) | 55.9% | hypothetical protein |
| 58,528:1 | +T | coding (1236/2937 nt) | 57.8% | hypothetical protein |
| 58,531 | A→T | V413V (GTA→GTT) | 58.6% | hypothetical protein |
| 58,541 | G→A | V417I (GTT→ATT) | 70.4% | hypothetical protein |
| 58,549 | T→C | L419L (CTT→CTC) | 71.7% | hypothetical protein |
| 58,555 | A→G | K421K (AAA→AAG) | 70.6% | hypothetical protein |
| 58,556 | T→C | L422L (TTA→CTA) | 70.1% | hypothetical protein |
| 58,567 | T→C | N425N (AAT→AAC) | 76.9% | hypothetical protein |
| 58,571 | C→T | L427L (CTA→TTA) | 75.9% | hypothetical protein |
| 58,576 | A→G | E428E (GAA→GAG) | 76.4% | hypothetical protein |
| 58,594 | G→A | K434K (AAG→AAA) | 80.2% | hypothetical protein |
| 58,6 | C→T | N436N (AAC→AAT) | 82.2% | hypothetical protein |
| 58,606 | A→G | V438V (GTA→GTG) | 82.3% | hypothetical protein |
| 58,639 | A→G | A449A (GCA→GCG) | 83.4% | hypothetical protein |
| 58,66 | C→T | T456T (ACC→ACT) | 84.7% | hypothetical protein |
| 58,663 | G→A | T457T (ACG→ACA) | 84.5% | hypothetical protein |
| 58,666 | A→G | A458A (GCA→GCG) | 84.9% | hypothetical protein |
| 58,727 | A→C | R479R (AGA→CGA) | 85.3% | hypothetical protein |
| 58,738 | G→A | V482V (GTG→GTA) | 85.0% | hypothetical protein |
| 58,744 | A→C | S484S (TCA→TCC) | 84.9% | hypothetical protein |
| 58,75 | C→T | A486A (GCC→GCT) | 84.1% | hypothetical protein |
| 58,765 | T→A | A491A (GCT→GCA) | 86.2% | hypothetical protein |
| 58,855 | T→C | F521F (TTT→TTC) | 88.9% | hypothetical protein |
| 58,915 | T→C | N541N (AAT→AAC) | 82.6% | hypothetical protein |
| 58,93 | T→C | A546A (GCT→GCC) | 81.3% | hypothetical protein |
| 58,942 | G→A | K550K (AAG→AAA) | 83.8% | hypothetical protein |
| 58,951 | C→T | R553R (CGC→CGT) | 85.2% | hypothetical protein |
| 58,954 | G→A | E554E (GAG→GAA) | 86.1% | hypothetical protein |
| 58,959 | T→C | I556T (ATT→ACT) | 86.0% | hypothetical protein |
| 58,972 | T→G | R560R (CGT→CGG) | 87.3% | hypothetical protein |
| 58,996 | C→A | I568I (ATC→ATA) | 88.2% | hypothetical protein |
| 59,003 | C→T | L571L (CTA→TTA) | 88.2% | hypothetical protein |
| 59,008 | C→T | S572S (TCC→TCT) | 92.4% | hypothetical protein |
| 59,014 | T→A | G574G (GGT→GGA) | 91.9% | hypothetical protein |

| | | | | | |
|---|---|---|---|---|---|
| 59,062 | C→T | F590F (TTC→TTT) | 87.4% | | hypothetical protein |
| 59,071 | A→G | P593P (CCA→CCG) | 87.9% | | hypothetical protein |
| 59,125 | G→A | E611E (GAG→GAA) | 85.7% | | hypothetical protein |
| 59,194 | A→G | L634L (CTA→CTG) | 85.8% | | hypothetical protein |
| 59,212 | G→T | P640P (CCG→CCT) | 86.0% | | hypothetical protein |
| 59,233 | C→T | G647G (GGC→GGT) | 88.1% | | hypothetical protein |
| 59,24 | C→T | L650L (CTA→TTA) | 86.9% | | hypothetical protein |
| 59,254 | T→G | T654T (ACT→ACG) | 86.3% | | hypothetical protein |
| 59,284 | C→T | N664N (AAC→AAT) | 84.2% | | hypothetical protein |
| 59,287 | C→T | S665S (TCC→TCT) | 83.3% | | hypothetical protein |
| 59,29 | T→G | T666T (ACT→ACG) | 82.1% | | hypothetical protein |
| 59,293 | A→T | S667S (TCA→TCT) | 82.0% | | hypothetical protein |
| 59,338 | A→G | L682L (CTA→CTG) | 81.9% | | hypothetical protein |
| 59,389 | T→A | G699G (GGT→GGA) | 80.6% | | hypothetical protein |
| 59,44 | T→C | N716N (AAT→AAC) | 79.7% | | hypothetical protein |
| 59,461 | C→T | F723F (TTC→TTT) | 82.5% | | hypothetical protein |
| 59,485 | C→T | G731G (GGC→GGT) | 84.3% | | hypothetical protein |
| 59,489 | A→C | K733Q (AAG→CAG) | 81.9% | | hypothetical protein |
| 59,494 | C→T | N734N (AAC→AAT) | 80.1% | | hypothetical protein |
| 59,497 | T→C | I735I (ATT→ATC) | 80.3% | | hypothetical protein |
| 59,533 | C→T | S747S (TCC→TCT) | 79.0% | | hypothetical protein |
| 59,548 | G→A | Q752Q (CAG→CAA) | 79.6% | | hypothetical protein |
| 59,554 | C→T | N754N (AAC→AAT) | 80.6% | | hypothetical protein |
| 59,566 | T→C | G758G (GGT→GGC) | 84.1% | | hypothetical protein |
| 59,593 | C→T | Y767Y (TAC→TAT) | 84.4% | 28.5% | hypothetical protein |
| 59,602 | C→T | R770R (CGC→CGT) | 82.1% | 26.3% | hypothetical protein |
| 59,605 | G→A | R771R (AGG→AGA) | 81.9% | 27.8% | hypothetical protein |
| 59,62 | G→A | R776R (AGG→AGA) | 81.7% | 28.7% | hypothetical protein |
| 59,623 | C→T | N777N (AAC→AAT) | 82.3% | 27.7% | hypothetical protein |
| 59,683 | T→C | S797S (AGT→AGC) | 79.8% | 28.3% | hypothetical protein |
| 59,71 | T→A | A806A (GCT→GCA) | 77.8% | 26.4% | hypothetical protein |
| 59,713 | T→G | G807G (GGT→GGG) | 78.0% | 27.3% | hypothetical protein |
| 59,719 | G→A | E809E (GAG→GAA) | 75.9% | 25.5% | hypothetical protein |
| 59,722 | C→T | Y810Y (TAC→TAT) | 76.5% | 26.7% | hypothetical protein |
| 59,724 | C→T | T811I (ACA→ATA) | 76.5% | 27.1% | hypothetical protein |
| 59,725 | A→G | T811T (ACA→ACG) | 78.1% | 28.4% | hypothetical protein |
| 59,761 | G→T | P823P (CCG→CCT) | 74.3% | 29.1% | hypothetical protein |
| 59,791 | G→A | A833A (GCG→GCA) | 75.4% | 33.2% | hypothetical protein |
| 59,81 | A→C | I840L (ATT→CTT) | 79.1% | 38.2% | hypothetical protein |
| 59,878 | A→G | K862K (AAA→AAG) | 78.0% | 29.5% | hypothetical protein |
| 59,887 | A→T | P865P (CCA→CCT) | 75.6% | 27.4% | hypothetical protein |
| 59,899 | T→G | A869A (GCT→GCG) | 76.5% | 25.9% | hypothetical protein |
| 59,908 | T→C | F872F (TTT→TTC) | 76.7% | 26.1% | hypothetical protein |
| 59,914 | G→A | K874K (AAG→AAA) | 77.7% | 27.7% | hypothetical protein |
| 59,929 | C→T | F879F (TTC→TTT) | 79.2% | 27.0% | hypothetical protein |
| 59,938 | C→T | D882D (GAC→GAT) | 79.7% | 29.3% | hypothetical protein |
| 59,956 | T→C | N888N (AAT→AAC) | 76.8% | 26.2% | hypothetical protein |
| 59,959 | A→C | V889V (GTA→GTC) | 77.2% | 26.6% | hypothetical protein |
| 59,962 | C→T | T890T (ACC→ACT) | 76.5% | 26.5% | hypothetical protein |
| 59,965 | T→C | I891I (ATT→ATC) | 76.8% | 27.3% | hypothetical protein |
| 59,986 | A→G | L898L (CTA→CTG) | 79.4% | 30.2% | hypothetical protein |
| 59,998 | A→G | K902K (AAA→AAG) | 78.7% | 30.1% | hypothetical protein |
| 60,004 | T→C | D904D (GAT→GAC) | 78.9% | 30.9% | hypothetical protein |

| | | | | | | |
|---|---|---|---|---|---|---|
| 60,022 | C→T | G910G (GGC→GGT) | 77.1% | 31.4% | | hypothetical protein |
| 60,079 | A→G | K929K (AAA→AAG) | 77.1% | | | hypothetical protein |
| 60,091 | G→T | T933T (ACG→ACT) | 75.5% | | | hypothetical protein |
| 60,094 | G→A | E934E (GAG→GAA) | 76.6% | | | hypothetical protein |
| 60,1 | C→T | I936I (ATC→ATT) | 73.9% | | | hypothetical protein |
| 60,118 | A→C | T942T (ACA→ACC) | 74.7% | | | hypothetical protein |
| 60,121 | C→T | Y943Y (TAC→TAT) | 74.0% | | | hypothetical protein |
| 60,157 | C→T | F955F (TTC→TTT) | 74.8% | | | hypothetical protein |
| 60,172 | A→G | E960E (GAA→GAG) | 75.6% | | | hypothetical protein |
| 60,184 | C→T | H964H (CAC→CAT) | 75.0% | | | hypothetical protein |
| 60,197 | T→G | S969A (TCA→GCA) | 76.8% | | | hypothetical protein |
| | | *979E (TAA→GAA) | | | | hypothetical protein |
| 60,227 | T→G | N4K (AAT→AAG) | 78.7% | | | Phage protein |
| 60,254 | T→C | F13F (TTT→TTC) | 79.0% | | | Phage protein |
| 60,266 | A→G | E17E (GAA→GAG) | 81.4% | | | Phage protein |
| 60,284 | T→C | Y23Y (TAT→TAC) | 81.2% | | | Phage protein |
| 60,401 | G→A | E62E (GAG→GAA) | 67.0% | | | Phage protein |
| 60,413 | A→C | V66V (GTA→GTC) | 64.2% | | | Phage protein |
| 60,426 | T→G | S71A (TCT→GCT) | 61.2% | | | Phage protein |
| 60,44 | A→G | K75K (AAA→AAG) | 62.5% | | | Phage protein |
| 60,446 | A→G | S77S (TCA→TCG) | 61.9% | | | Phage protein |
| 60,468 | T→G | intergenic (+19/-53) | 63.1% | | | Phage protein |
| 60,502 | Δ1 bp | intergenic (+53/-19) | 57.2% | | | Phage protein |
| 60,503 | Δ1 bp | intergenic (+54/-18) | 56.9% | | | Phage protein |
| 60,535 | C→T | R5R (CGC→CGT) | 56.8% | | | Phage protein |
| 60,547 | C→T | H9H (CAC→CAT) | 58.6% | | | Phage protein |
| 60,58 | A→G | K20K (AAA→AAG) | 60.3% | | | Phage protein |
| 60,646 | C→T | S42S (TCC→TCT) | 54.8% | | | Phage protein |
| 60,664 | G→A | R48R (AGG→AGA) | 57.1% | | | Phage protein |
| 60,685 | C→T | D55D (GAC→GAT) | 58.6% | | | Phage protein |
| 60,691 | G→A | Q57Q (CAG→CAA) | 56.3% | | | Phage protein |
| 65,741 | T→G | intergenic (+24/-36) | | | 10.7% | Phage protein |
| 65,742 | C→A | intergenic (+25/-35) | | | 10.4% | Phage protein |
| 68,710:1 | +C | coding (370/891 nt) | | | 24.2% | Phage DNA primase/helicase |
| 68,714 | Δ1 bp | coding (374/891 nt) | | | 23.5% | Phage DNA primase/helicase |
| 68,838 | G→A | P166P (CCG→CCA) | | | 28.2% | Phage DNA primase/helicase |
| 68,868 | A→G | P176P (CCA→CCG) | | | 29.8% | Phage DNA primase/helicase |
| 69,006 | T→C | A222A (GCT→GCC) | | | 26.9% | Phage DNA primase/helicase |
| 69,009 | G→T | G223G (GGG→GGT) | | | 27.5% | Phage DNA primase/helicase |
| 69,075 | T→C | A245A (GCT→GCC) | | | 25.1% | Phage DNA primase/helicase |
| 69,123 | C→T | F261F (TTC→TTT) | | | 23.0% | Phage DNA primase/helicase |
| 69,126 | C→T | L262L (CTC→CTT) | | | 23.1% | Phage DNA primase/helicase |
| 69,138 | C→T | G266G (GGC→GGT) | | | 22.6% | Phage DNA primase/helicase |
| 69,144 | T→C | D268D (GAT→GAC) | | | 22.2% | Phage DNA primase/helicase |
| 69,147 | A→G | P269P (CCA→CCG) | | | 22.0% | Phage DNA primase/helicase |
| 69,153 | T→C | D271D (GAT→GAC) | | | 23.6% | Phage DNA primase/helicase |
| 69,159 | C→T | N273N (AAC→AAT) | | | 22.0% | Phage DNA primase/helicase |
| 69,165 | T→A | D275E (GAT→GAA) | | | 21.8% | Phage DNA primase/helicase |
| 69,168 | A→G | E276E (GAA→GAG) | | | 22.2% | Phage DNA primase/helicase |
| 69,176 | T→C | M279T (ATG→ACG) | | | 22.9% | Phage DNA primase/helicase |
| 69,401 | A→G | L6L (TTA→TTG) | | | 27.8% | DNA polymerase, phage-associated |
| 69,416 | T→C | L11L (CTT→CTC) | | | 26.3% | DNA polymerase, phage-associated |
| 69,425 | C→T | H14H (CAC→CAT) | | | 38.0% | DNA polymerase, phage-associated |

| | | | | | |
|---|---|---|---|---|---|
| 69,434 | T→C | I17I (ATT→ATC) | | 37.1% | DNA polymerase, phage-associated |
| 69,44 | A→T | T19T (ACA→ACT) | | 38.3% | DNA polymerase, phage-associated |
| 69,443 | A→G | P20P (CCA→CCG) | | 36.6% | DNA polymerase, phage-associated |
| 69,458 | T→C | D25D (GAT→GAC) | | 38.9% | DNA polymerase, phage-associated |
| 69,485 | A→G | L34L (CTA→CTG) | | 40.3% | DNA polymerase, phage-associated |
| 69,494 | T→C | A37A (GCT→GCC) | | 41.6% | DNA polymerase, phage-associated |
| 69,512 | C→T | F43F (TTC→TTT) | | 41.5% | DNA polymerase, phage-associated |
| 69,515 | A→G | A44A (GCA→GCG) | | 41.2% | DNA polymerase, phage-associated |
| 69,536 | C→T | D51D (GAC→GAT) | | 41.9% | DNA polymerase, phage-associated |
| 69,542 | T→C | T53T (ACT→ACC) | | 42.5% | DNA polymerase, phage-associated |
| 69,554 | G→A | V57V (GTG→GTA) | | 43.8% | DNA polymerase, phage-associated |
| 69,59 | C→T | I69I (ATC→ATT) | | 42.4% | DNA polymerase, phage-associated |
| 69,614 | G→A | K77K (AAG→AAA) | | 40.6% | DNA polymerase, phage-associated |
| 69,725 | C→T | D114D (GAC→GAT) | | 39.8% | DNA polymerase, phage-associated |
| 69,731 | G→T | A116A (GCG→GCT) | | 40.5% | DNA polymerase, phage-associated |
| 69,773 | T→C | P130P (CCT→CCC) | | 40.8% | DNA polymerase, phage-associated |
| 69,776 | A→C | V131V (GTA→GTC) | | 39.7% | DNA polymerase, phage-associated |
| 69,83 | C→T | G149G (GGC→GGT) | | 42.5% | DNA polymerase, phage-associated |
| 69,839 | G→T | M152I (ATG→ATT) | | 42.8% | DNA polymerase, phage-associated |
| 69,89 | A→G | E169E (GAA→GAG) | | 41.7% | DNA polymerase, phage-associated |
| 70,037 | T→C | T218T (ACT→ACC) | 8.4% | 45.4% | DNA polymerase, phage-associated |
| 70,049 | T→C | H222H (CAT→CAC) | 9.6% | 47.1% | DNA polymerase, phage-associated |
| 70,064 | G→A | E227E (GAG→GAA) | 8.0% | 48.1% | DNA polymerase, phage-associated |
| 70,118 | C→T | G245G (GGC→GGT) | 6.1% | 40.8% | DNA polymerase, phage-associated |
| 70,13 | T→C | F249F (TTT→TTC) | 6.8% | 39.4% | DNA polymerase, phage-associated |
| 70,16 | C→T | C259C (TGC→TGT) | 5.3% | 39.1% | DNA polymerase, phage-associated |
| 70,193 | A→T | T270T (ACA→ACT) | 6.6% | 38.5% | DNA polymerase, phage-associated |
| 70,202 | A→G | L273L (TTA→TTG) | 6.7% | 37.5% | DNA polymerase, phage-associated |
| 70,226 | T→A | P281P (CCT→CCA) | 5.5% | 35.1% | DNA polymerase, phage-associated |
| 70,229 | C→T | Y282Y (TAC→TAT) | | 34.6% | DNA polymerase, phage-associated |
| 70,235 | A→G | A284A (GCA→GCG) | 5.6% | 37.7% | DNA polymerase, phage-associated |
| 70,25 | T→C | A289A (GCT→GCC) | 5.3% | 38.2% | DNA polymerase, phage-associated |
| 70,319 | T→C | Y312Y (TAT→TAC) | | 42.1% | DNA polymerase, phage-associated |
| 70,326 | C→T | L315L (CTA→TTA) | | 37.8% | DNA polymerase, phage-associated |
| 70,328 | A→G | L315L (CTA→CTG) | | 38.5% | DNA polymerase, phage-associated |
| 70,34 | T→C | C319C (TGT→TGC) | | 39.5% | DNA polymerase, phage-associated |
| 70,346 | T→C | F321F (TTT→TTC) | | 33.7% | DNA polymerase, phage-associated |
| 70,349 | C→G | L322L (CTC→CTG) | | 33.4% | DNA polymerase, phage-associated |
| 70,352 | G→A | Q323Q (CAG→CAA) | | 32.0% | DNA polymerase, phage-associated |
| 70,37 | T→A | G329G (GGT→GGA) | | 29.5% | DNA polymerase, phage-associated |
| 70,379 | T→C | I332I (ATT→ATC) | | 26.5% | DNA polymerase, phage-associated |
| 70,394 | A→G | L337L (TTA→TTG) | | 24.3% | DNA polymerase, phage-associated |
| 70,409 | C→T | Y342Y (TAC→TAT) | | 21.8% | DNA polymerase, phage-associated |
| 70,415 | A→G | L344L (TTA→TTG) | | 20.5% | DNA polymerase, phage-associated |
| 70,425 | C→T | L348L (CTG→TTG) | | 21.3% | DNA polymerase, phage-associated |
| 70,436 | T→C | A351A (GCT→GCC) | | 21.2% | DNA polymerase, phage-associated |
| 70,445 | G→A | K354K (AAG→AAA) | | 22.4% | DNA polymerase, phage-associated |
| 70,451 | T→C | Y356Y (TAT→TAC) | | 24.5% | DNA polymerase, phage-associated |
| 70,466 | C→T | V361V (GTC→GTT) | | 24.0% | DNA polymerase, phage-associated |
| 70,467 | G→A | V362I (GTT→ATT) | | 23.4% | DNA polymerase, phage-associated |
| 70,468 | T→A | V362D (GTT→GAT) | | 24.4% | DNA polymerase, phage-associated |
| 70,469 | T→A | V362V (GTT→GTA) | | 23.5% | DNA polymerase, phage-associated |
| 70,479 | A→C | K366Q (AAA→CAA) | | 27.8% | DNA polymerase, phage-associated |

| | | | | | |
|---|---|---|---|---|---|
| 70,49 | C→T | N369N (AAC→AAT) | | 30.0% | DNA polymerase, phage-associated |
| 70,496 | G→A | A371A (GCG→GCA) | | 30.9% | DNA polymerase, phage-associated |
| 70,505 | A→G | P374P (CCA→CCG) | | 33.5% | DNA polymerase, phage-associated |
| 70,544 | T→C | Y387Y (TAT→TAC) | | 37.8% | DNA polymerase, phage-associated |
| 70,556 | C→T | T391T (ACC→ACT) | | 38.7% | DNA polymerase, phage-associated |
| 70,569 | T→C | L396L (TTG→CTG) | | 38.8% | DNA polymerase, phage-associated |
| 70,61 | G→A | L409L (CTG→CTA) | 17.9% | 41.8% | DNA polymerase, phage-associated |
| 70,703 | T→A | L440L (CTT→CTA) | 25.7% | 40.7% | DNA polymerase, phage-associated |
| 70,745 | C→T | H454H (CAC→CAT) | 24.0% | 38.4% | DNA polymerase, phage-associated |
| 70,751 | C→T | H456H (CAC→CAT) | 25.6% | 38.9% | DNA polymerase, phage-associated |
| 70,772 | A→G | L463L (CTA→CTG) | 24.1% | 37.9% | DNA polymerase, phage-associated |
| 70,808 | T→C | P475P (CCT→CCC) | 20.6% | 39.4% | DNA polymerase, phage-associated |
| 70,85 | A→T | P489P (CCA→CCT) | 18.6% | 42.0% | DNA polymerase, phage-associated |
| 70,863 | G→A | V494I (GTC→ATC) | 17.1% | 42.0% | DNA polymerase, phage-associated |
| 70,868 | A→G | A495A (GCA→GCG) | 18.3% | 43.9% | DNA polymerase, phage-associated |
| 70,906 | C→T | A508V (GCT→GTT) | 22.2% | 43.0% | DNA polymerase, phage-associated |
| 70,934 | A→G | Q517Q (CAA→CAG) | 22.8% | 42.5% | DNA polymerase, phage-associated |
| 70,961 | T→C | P526P (CCT→CCC) | 21.2% | 39.1% | DNA polymerase, phage-associated |
| 70,973 | T→A | P530P (CCT→CCA) | 20.7% | 39.9% | DNA polymerase, phage-associated |
| 70,985 | T→C | S534S (TCT→TCC) | 19.9% | 39.8% | DNA polymerase, phage-associated |
| 71,003 | A→G | V540V (GTA→GTG) | 20.7% | 40.3% | DNA polymerase, phage-associated |
| 71,054 | A→G | L557L (CTA→CTG) | 19.5% | 37.3% | DNA polymerase, phage-associated |
| 71,063 | A→T | A560A (GCA→GCT) | 19.3% | 36.6% | DNA polymerase, phage-associated |
| 71,117 | A→C | A578A (GCA→GCC) | 19.5% | 32.5% | DNA polymerase, phage-associated |
| 71,132 | A→G | E583E (GAA→GAG) | 15.9% | 22.2% | DNA polymerase, phage-associated |
| 71,135 | A→T | A584A (GCA→GCT) | 15.7% | 22.0% | DNA polymerase, phage-associated |
| 71,136 | T→C | L585L (TTG→CTG) | 16.3% | 22.6% | DNA polymerase, phage-associated |
| 71,138 | G→T | L585F (TTG→TTT) | 15.2% | 22.1% | DNA polymerase, phage-associated |
| 71,141 | C→G | L586L (CTC→CTG) | 15.3% | 21.7% | DNA polymerase, phage-associated |
| 71,147 | A→G | Q588Q (CAA→CAG) | 17.1% | 23.9% | DNA polymerase, phage-associated |
| 71,15 | A→T | A589A (GCA→GCT) | 15.8% | 23.1% | DNA polymerase, phage-associated |
| 71,153 | C→T | A590A (GCC→GCT) | 15.2% | 21.6% | DNA polymerase, phage-associated |
| 71,156 | A→G | K591K (AAA→AAG) | 15.1% | 22.7% | DNA polymerase, phage-associated |
| 71,159 | T→A | T592T (ACT→ACA) | 15.5% | 21.9% | DNA polymerase, phage-associated |
| 71,162 | G→A | G593G (GGG→GGA) | 15.3% | 21.8% | DNA polymerase, phage-associated |
| 71,189 | A→G | A602A (GCA→GCG) | 16.8% | 27.2% | DNA polymerase, phage-associated |
| 71,195 | T→G | A604A (GCT→GCG) | 13.6% | 23.0% | DNA polymerase, phage-associated |
| 71,198 | A→G | K605K (AAA→AAG) | 14.1% | 23.5% | DNA polymerase, phage-associated |
| 71,201 | G→C | E606D (GAG→GAC) | 13.7% | 22.9% | DNA polymerase, phage-associated |
| 71,204 | C→T | Y607Y (TAC→TAT) | 13.1% | 21.9% | DNA polymerase, phage-associated |
| 71,207 | T→C | I608I (ATT→ATC) | 12.7% | 21.8% | DNA polymerase, phage-associated |
| 71,231 | G→A | P616P (CCG→CCA) | 15.3% | 28.5% | DNA polymerase, phage-associated |
| 71,291 | C→T | Y636Y (TAC→TAT) | 24.4% | 37.8% | DNA polymerase, phage-associated |
| 71,294 | C→T | S637S (AGC→AGT) | 24.0% | 39.1% | DNA polymerase, phage-associated |
| 71,297 | C→T | H638H (CAC→CAT) | 24.2% | 38.5% | DNA polymerase, phage-associated |
| 71,324 | C→T | N647N (AAC→AAT) | 25.5% | 40.9% | DNA polymerase, phage-associated |
| 71,339 | T→C | D652D (GAT→GAC) | 26.3% | 44.3% | DNA polymerase, phage-associated |
| 71,354 | G→T | G657G (GGG→GGT) | 28.5% | 43.8% | DNA polymerase, phage-associated |
| 71,528 | C→T | I715I (ATC→ATT) | 26.9% | 42.1% | DNA polymerase, phage-associated |
| 71,546 | C→T | I721I (ATC→ATT) | 26.8% | 43.2% | DNA polymerase, phage-associated |
| 71,555 | T→C | R724R (CGT→CGC) | 27.2% | 43.9% | DNA polymerase, phage-associated |
| 71,57 | C→T | D729D (GAC→GAT) | 26.8% | 45.3% | DNA polymerase, phage-associated |
| 71,642 | C→T | Y753Y (TAC→TAT) | 29.4% | 45.9% | DNA polymerase, phage-associated |

| | | | | | |
|---|---|---|---|---|---|
| 71,708 | C→T | R775R (CGC→CGT) | 28.6% | 40.4% | DNA polymerase, phage-associated |
| 71,738 | G→A | E785E (GAG→GAA) | 28.4% | 43.5% | DNA polymerase, phage-associated |
| 71,749 | G→A | R789K (AGG→AAG) | 25.7% | 43.1% | DNA polymerase, phage-associated |
| 71,754 | T→C | L791L (TTA→CTA) | 26.7% | 44.8% | DNA polymerase, phage-associated |
| 71,792 | G→A | K803K (AAG→AAA) | 21.9% | 44.2% | DNA polymerase, phage-associated |
| 71,801 | C→T | D806D (GAC→GAT) | 22.7% | 44.8% | DNA polymerase, phage-associated |
| 71,807 | G→A | K808K (AAG→AAA) | 21.7% | 44.6% | DNA polymerase, phage-associated |
| 71,817 | A→G | I812V (ATA→GTA) | 21.2% | 44.7% | DNA polymerase, phage-associated |
| 71,862 | T→C | N3N (AAT→AAC) | 21.6% | 50.9% | Phage protein |
| 71,892 | G→A | A13A (GCG→GCA) | 20.2% | 49.9% | Phage protein |
| 72,036 | G→A | K61K (AAG→AAA) | 22.7% | 53.1% | Phage protein |
| 72,099 | G→A | Q82Q (CAG→CAA) | 21.2% | 52.9% | Phage protein |
| 72,174 | G→A | L107L (TTG→TTA) | 21.9% | 53.9% | Phage protein |
| 72,431 | C→T | H28H (CAC→CAT) | 65.4% | 52.9% | Phage DNA helicase |
| 72,446 | C→T | T33T (ACC→ACT) | 66.2% | 53.5% | Phage DNA helicase |
| 72,528 | T→C | L61L (TTA→CTA) | 73.9% | 53.0% | Phage DNA helicase |
| 72,543 | A→G | I66V (ATA→GTA) | 71.5% | 51.3% | Phage DNA helicase |
| 72,581 | G→T | P78P (CCG→CCT) | 70.9% | 49.1% | Phage DNA helicase |
| 72,65 | T→C | D101D (GAT→GAC) | 73.5% | 52.4% | Phage DNA helicase |
| 72,68 | G→A | G111G (GGG→GGA) | 72.6% | 51.1% | Phage DNA helicase |
| 72,701 | A→C | A118A (GCA→GCC) | 72.5% | 38.9% | Phage DNA helicase |
| 72,704 | T→C | L119L (CTT→CTC) | 71.8% | 38.5% | Phage DNA helicase |
| 72,809 | G→A | P154P (CCG→CCA) | 73.5% | 35.3% | Phage DNA helicase |
| 72,812 | C→T | G155G (GGC→GGT) | 73.1% | 35.0% | Phage DNA helicase |
| 72,83 | A→G | K161K (AAA→AAG) | 71.8% | 34.2% | Phage DNA helicase |
| 72,857 | T→C | V170V (GTT→GTC) | 72.2% | 34.4% | Phage DNA helicase |
| 72,977 | C→A | S210S (TCC→TCA) | 72.5% | 30.8% | Phage DNA helicase |
| 72,983 | G→T | A212A (GCG→GCT) | 74.2% | 31.8% | Phage DNA helicase |
| 73,004 | T→C | S219S (TCT→TCC) | 73.4% | 30.6% | Phage DNA helicase |
| 73,007 | A→T | G220G (GGA→GGT) | 72.1% | 30.3% | Phage DNA helicase |
| 73,011 | T→C | L222L (TTA→CTA) | 74.0% | 31.2% | Phage DNA helicase |
| 73,019 | T→A | R224R (CGT→CGA) | 75.2% | 34.1% | Phage DNA helicase |
| 73,029 | C→T | L228L (CTA→TTA) | 76.7% | 34.7% | Phage DNA helicase |
| 73,067 | C→A | I240I (ATC→ATA) | 76.0% | 37.5% | Phage DNA helicase |
| 73,106 | T→C | I253I (ATT→ATC) | 77.4% | 38.6% | Phage DNA helicase |
| 73,115 | C→T | Y256Y (TAC→TAT) | 75.3% | 39.5% | Phage DNA helicase |
| 73,142 | T→C | N265N (AAT→AAC) | 76.1% | 37.3% | Phage DNA helicase |
| 73,151 | G→A | V268V (GTG→GTA) | 74.9% | 37.0% | Phage DNA helicase |
| 73,241 | G→A | G298G (GGG→GGA) | 71.8% | 29.8% | Phage DNA helicase |
| 73,253 | T→C | L302L (CTT→CTC) | 72.0% | 27.9% | Phage DNA helicase |
| 73,285 | T→C | I313T (ATA→ACA) | 72.4% | 26.3% | Phage DNA helicase |
| 73,298 | T→A | A317A (GCT→GCA) | 69.2% | 20.9% | Phage DNA helicase |
| 73,301 | T→A | L318L (CTT→CTA) | 69.5% | 21.5% | Phage DNA helicase |
| 73,302 | G→T | A319S (GCA→TCA) | 69.2% | 20.8% | Phage DNA helicase |
| 73,304 | A→T | A319A (GCA→GCT) | 71.7% | 21.9% | Phage DNA helicase |
| 73,306 | A→T | Q320L (CAG→CTG) | 70.8% | 21.1% | Phage DNA helicase |
| 73,31 | T→A | R321R (CGT→CGA) | 73.1% | 22.4% | Phage DNA helicase |
| 73,313 | T→A | G322G (GGT→GGA) | 73.5% | 23.4% | Phage DNA helicase |
| 73,322 | A→C | T325T (ACA→ACC) | 77.1% | 26.0% | Phage DNA helicase |
| 73,328 | A→G | E327E (GAA→GAG) | 77.2% | 27.3% | Phage DNA helicase |
| 73,334 | A→T | I329I (ATA→ATT) | 74.6% | 25.1% | Phage DNA helicase |
| 73,337 | A→C | G330G (GGA→GGC) | 75.3% | 25.9% | Phage DNA helicase |
| 73,394 | G→T | P349P (CCG→CCT) | 100% | 30.7% | Phage DNA helicase |

| | | | | | |
|---|---|---|---|---|---|
| 73,4 | G→A | V351V (GTG→GTA) | 100% | 30.1% | Phage DNA helicase |
| 73,418 | C→T | S357S (AGC→AGT) | 100% | 34.0% | Phage DNA helicase |
| 73,427 | T→C | S360S (TCT→TCC) | 100% | 35.4% | Phage DNA helicase |
| 73,43 | G→A | E361E (GAG→GAA) | 100% | 34.5% | Phage DNA helicase |
| 73,466 | T→G | I373M (ATT→ATG) | 100% | 35.9% | Phage DNA helicase |
| 73,475 | T→C | L376L (CTT→CTC) | 100% | 36.3% | Phage DNA helicase |
| 73,511 | T→A | G388G (GGT→GGA) | 100% | 34.5% | Phage DNA helicase |
| 73,643 | T→C | T432T (ACT→ACC) | 100% | 31.6% | Phage DNA helicase |
| 73,649 | T→C | T434T (ACT→ACC) | 100% | 30.5% | Phage DNA helicase |
| 73,652 | G→T | P435P (CCG→CCT) | 100% | 29.4% | Phage DNA helicase |
| 73,696 | T→C | L450S (TTA→TCA) | 100% | 31.7% | Phage DNA helicase |
| 73,716 | C→T | intergenic (+16/-205) | 100% | 32.2% | Phage DNA helicase |
| 73,73 | +GAGA | intergenic (+30/-191) | 93.6% | 31.8% | Phage DNA helicase |
| 73,766 | T→G | intergenic (+66/-155) | 100% | 37.8% | Phage DNA helicase |
| 73,838 | C→T | intergenic (+138/-83) | 100% | 53.4% | Phage DNA helicase |
| 73,965 | G→A | S15S (TCG→TCA) | 100% | 52.6% | Phage protein |
| 73,977 | C→T | N19N (AAC→AAT) | 100% | 53.2% | Phage protein |
| 74,016 | T→C | N32N (AAT→AAC) | 100% | 55.1% | Phage protein |
| 74,1 | A→G | I60M (ATA→ATG) | 100% | 57.2% | Phage protein |
| 74,169 | G→A | S83S (TCG→TCA) | 100% | 50.1% | Phage protein |
| 74,269 | C→T | G24G (GGC→GGT) | 100% | 51.0% | gp32-liked protein |
| 74,34 | A→G | N48S (AAC→AGC) | 100% | 49.4% | gp32-liked protein |
| 74,494 | C→T | A99A (GCC→GCT) | 100% | 52.8% | gp32-liked protein |
| 74,503 | T→C | I102I (ATT→ATC) | 100% | 52.4% | gp32-liked protein |
| 74,582 | T→C | L129L (TTG→CTG) | 100% | 50.3% | gp32-liked protein |
| 74,586 | A→G | N130S (AAC→AGC) | 100% | 49.5% | gp32-liked protein |
| 74,632 | C→T | G145G (GGC→GGT) | 100% | 50.3% | gp32-liked protein |
| 74,668 | T→C | I157I (ATT→ATC) | 100% | 49.4% | gp32-liked protein |
| 74,734 | G→A | A179A (GCG→GCA) | 100% | 51.4% | gp32-liked protein |
| 74,749 | T→C | I184I (ATT→ATC) | 100% | 50.8% | gp32-liked protein |
| 74,767 | C→T | D190D (GAC→GAT) | 100% | 50.0% | gp32-liked protein |
| 74,794 | G→A | E199E (GAG→GAA) | 100% | ? | gp32-liked protein |
| 74,859 | A→T | Q221L (CAA→CTA) | 100% | 43.6% | gp32-liked protein |
| 74,872 | A→G | E225E (GAA→GAG) | 100% | 42.7% | gp32-liked protein |
| 74,911 | C→T | N238N (AAC→AAT) | 100% | 42.0% | gp32-liked protein |
| 74,92 | T→A | A241A (GCT→GCA) | 100% | 42.2% | gp32-liked protein |
| 74,942 | C→T | Q249* (CAA→TAA) | | 40.6% | gp32-liked protein |
| 74,942 | 2 bp→TC | coding (745-746/774 nt) | 100% | | gp32-liked protein |
| 74,943 | A→C | Q249P (CAA→CCA) | | 42.0% | gp32-liked protein |
| 74,965 | T→C | D256D (GAT→GAC) | 100% | 42.8% | gp32-liked protein |
| 75,006 | T→C | intergenic (+35/-2) | | 45.3% | Phage-associated recombinase |
| 75,006 | 2 bp→CA | intergenic (+35/-1) | 100% | | Phage-associated recombinase |
| 75,007 | T→A | intergenic (+36/-1) | | 45.4% | Phage-associated recombinase |
| 75,04 | T→C | I11I (ATT→ATC) | 100% | 42.9% | Phage-associated recombinase |
| 75,043 | G→A | K12K (AAG→AAA) | 100% | 41.9% | Phage-associated recombinase |
| 75,046 | G→A | L13L (CTG→CTA) | 100% | 42.6% | Phage-associated recombinase |
| 75,049 | C→A | G14G (GGC→GGA) | 100% | 41.8% | Phage-associated recombinase |
| 75,059 | A→G | I18V (ATA→GTT) | 100% | 42.5% | Phage-associated recombinase |
| 75,061 | A→T | I18V (ATA→GTT) | 100% | 42.0% | Phage-associated recombinase |
| 75,064 | C→A | P19P (CCC→CCA) | 100% | 40.2% | Phage-associated recombinase |
| 75,067 | A→G | K20K (AAA→AAG) | 89.4% | 36.8% | Phage-associated recombinase |
| 75,068:1 | +G | coding (61/978 nt) | 9.0% | 5.3% | Phage-associated recombinase |
| 75,14 | T→C | L45L (TTG→CTT) | 100% | 69.5% | Phage-associated recombinase |

| | | | | | |
|---|---|---|---|---|---|
| 75,142 | G→T | L45L (TTG→CTT) | 100% | 67.7% | Phage-associated recombinase |
| 75,149 | A→G | I48V (ATT→GTT) | | 66.5% | Phage-associated recombinase |
| 75,149 | 2 bp→GC | coding (142-143/978 nt) | 100% | | Phage-associated recombinase |
| 75,15 | T→C | I48T (ATT→ACT) | | 66.7% | Phage-associated recombinase |
| 75,161 | T→A | L52I (TTA→ATA) | 94.5% | 61.9% | Phage-associated recombinase |
| 75,164 | T→C | F53L (TTT→CTT) | 100% | 62.7% | Phage-associated recombinase |
| 75,17 | C→G | H55D (CAT→GAT) | 92.8% | 54.8% | Phage-associated recombinase |
| 75,171 | A→T | H55L (CAT→CTT) | 93.2% | 54.4% | Phage-associated recombinase |
| 75,173 | T→G | S56A (TCC→GCC) | 91.5% | 52.9% | Phage-associated recombinase |
| 75,187 | G→A | S60S (TCG→TCA) | 93.1% | 51.0% | Phage-associated recombinase |
| 75,194 | G→A | V63I (GTA→ATA) | 89.9% | 46.6% | Phage-associated recombinase |
| 75,202 | T→G | L65L (CTT→CTG) | 82.9% | 27.5% | Phage-associated recombinase |
| 75,205 | A→T | L66L (CTA→CTT) | 81.1% | 26.1% | Phage-associated recombinase |
| 75,209 | A→C | K68Q (AAA→CAA) | 81.0% | 27.2% | Phage-associated recombinase |
| 75,211 | A→G | K68K (AAA→AAG) | 81.9% | 27.0% | Phage-associated recombinase |
| 75,214 | T→C | F69F (TTT→TTC) | 80.3% | 27.3% | Phage-associated recombinase |
| 75,22 | T→A | S71S (TCT→TCA) | 63.8% | 12.7% | Phage-associated recombinase |
| 75,226 | A→T | L73L (CTA→CTT) | 63.8% | 12.7% | Phage-associated recombinase |
| 75,229 | T→C | D74D (GAT→GAC) | 62.5% | 11.7% | Phage-associated recombinase |
| 75,233 | A→C | I76L (ATA→CTA) | 65.2% | 11.9% | Phage-associated recombinase |
| 75,234 | T→C | I76T (ATA→ACA) | 65.7% | 11.2% | Phage-associated recombinase |
| 75,238 | A→C | G77G (GGA→GGC) | 60.9% | 11.2% | Phage-associated recombinase |
| 75,24 | T→A | M78K (ATG→AAG) | 59.7% | 11.0% | Phage-associated recombinase |
| 75,241 | G→A | M78I (ATG→ATA) | 62.1% | 10.9% | Phage-associated recombinase |
| 75,242 | G→A | V79I (GTA→ATA) | 59.7% | 10.7% | Phage-associated recombinase |
| 75,244 | A→C | V79V (GTA→GTC) | 59.7% | 11.1% | Phage-associated recombinase |
| 75,25 | A→T | T81T (ACA→ACT) | 83.7% | 31.2% | Phage-associated recombinase |
| 75,253 | T→A | G82G (GGT→GGA) | 84.4% | 31.0% | Phage-associated recombinase |
| 75,263 | T→A | L86I (TTA→ATA) | 88.3% | 38.3% | Phage-associated recombinase |
| 75,265 | A→G | L86L (TTA→TTG) | 88.6% | 39.2% | Phage-associated recombinase |
| 75,266 | A→T | I87L (ATA→TTA) | 87.3% | 38.6% | Phage-associated recombinase |
| 75,289 | G→A | L94L (CTG→CTA) | 100% | 52.7% | Phage-associated recombinase |
| 75,292 | C→T | Y95Y (TAC→TAT) | 100% | 53.1% | Phage-associated recombinase |
| 75,295 | C→T | H96H (CAC→CAT) | 100% | 53.4% | Phage-associated recombinase |
| 75,301 | G→A | A98A (GCG→GCA) | 100% | 59.8% | Phage-associated recombinase |
| 75,316 | A→G | K103K (AAA→AAG) | 100% | 65.7% | Phage-associated recombinase |
| 75,322 | T→G | T105T (ACT→ACG) | 100% | 65.3% | Phage-associated recombinase |
| 75,325 | T→C | S106S (AGT→AGC) | 100% | 68.6% | Phage-associated recombinase |
| 75,328 | G→A | G107G (GGG→GGA) | 100% | 68.5% | Phage-associated recombinase |
| 75,331 | G→A | K108K (AAG→AAA) | 100% | 68.5% | Phage-associated recombinase |
| 75,391 | G→A | E128E (GAG→GAA) | 100% | 75.8% | Phage-associated recombinase |
| 75,394 | T→C | I129I (ATT→ATC) | 100% | 74.9% | Phage-associated recombinase |
| 75,4 | A→G | K131K (AAA→AAG) | | 74.6% | Phage-associated recombinase |
| 75,4 | 2 bp→GT | coding (393-394/978 nt) | 100% | | Phage-associated recombinase |
| 75,401 | C→T | P132S (CCA→TCA) | | 74.1% | Phage-associated recombinase |
| 75,403 | A→T | P132P (CCA→CCT) | 94.5% | 74.6% | Phage-associated recombinase |
| 75,415 | T→A | P136P (CCT→CCA) | | 76.7% | Phage-associated recombinase |
| 75,415 | 2 bp→AT | coding (408-409/978 nt) | 100% | | Phage-associated recombinase |
| 75,416 | G→T | A137S (GCT→TCT) | | 76.7% | Phage-associated recombinase |
| 75,433 | C→T | C142C (TGC→TGT) | 100% | 75.7% | Phage-associated recombinase |
| 75,448 | G→C | R147R (CGG→CGC) | 100% | 77.6% | Phage-associated recombinase |
| 75,463 | A→G | P152P (CCA→CCG) | 100% | 79.5% | Phage-associated recombinase |
| 75,496 | C→T | Y163Y (TAC→TAT) | 94.3% | 81.2% | Phage-associated recombinase |

| | | | | | |
|---|---|---|---|---|---|
| 75,532 | T→C | H175H (CAT→CAC) | 94.8% | 79.9% | Phage-associated recombinase |
| 75,559 | G→A | G184G (GGG→GGA) | 94.7% | 80.1% | Phage-associated recombinase |
| 75,565 | T→C | T186T (ACT→ACC) | 100% | 80.6% | Phage-associated recombinase |
| 75,574 | T→C | L189L (CTT→CTC) | 100% | 80.1% | Phage-associated recombinase |
| 75,644 | A→G | I213V (ATT→GTT) | 100% | 77.9% | Phage-associated recombinase |
| 75,72 | C→T | A238V (GCA→GTA) | | 77.7% | Phage-associated recombinase |
| 75,72 | 2 bp→TT | coding (713-714/978 nt) | 100% | | Phage-associated recombinase |
| 75,721 | A→T | A238A (GCA→GCT) | | 78.9% | Phage-associated recombinase |
| 75,763 | C→T | Y252Y (TAC→TAT) | 94.6% | 78.8% | Phage-associated recombinase |
| 75,995 | A→G | K10K (AAA→AAG) | 100% | 83.7% | Phage recombination related exonuclease (EC 3.1.11.-) |
| 76,034 | C→T | I23I (ATC→ATT) | 100% | 82.1% | Phage recombination related exonuclease (EC 3.1.11.-) |
| 76,07 | A→T | G35G (GGA→GGT) | | 76.6% | Phage recombination related exonuclease (EC 3.1.11.-) |
| 76,07 | 2 bp→TA | coding (105-106/1839 nt) | 100% | | Phage recombination related exonuclease (EC 3.1.11.-) |
| 76,071 | G→A | G36S (GGC→AGC) | | 78.0% | Phage recombination related exonuclease (EC 3.1.11.-) |
| 76,103 | A→T | T46T (ACA→ACT) | 100% | 71.5% | Phage recombination related exonuclease (EC 3.1.11.-) |
| 76,109 | C→T | I48I (ATC→ATT) | 100% | 64.5% | Phage recombination related exonuclease (EC 3.1.11.-) |
| 76,112 | G→A | E49E (GAG→GAA) | 100% | 65.5% | Phage recombination related exonuclease (EC 3.1.11.-) |
| 76,115 | G→A | E50E (GAG→GAA) | | 65.6% | Phage recombination related exonuclease (EC 3.1.11.-) |
| 76,115 | 2 bp→AC | coding (150-151/1839 nt) | 100% | | Phage recombination related exonuclease (EC 3.1.11.-) |
| 76,116 | T→C | L51L (TTG→CTG) | | 67.2% | Phage recombination related exonuclease (EC 3.1.11.-) |
| 76,118 | G→C | L51F (TTG→TTC) | 100% | 66.3% | Phage recombination related exonuclease (EC 3.1.11.-) |
| 76,127 | C→T | N54N (AAC→AAT) | 100% | 65.0% | Phage recombination related exonuclease (EC 3.1.11.-) |
| 76,136 | T→A | S57S (TCT→TCA) | 100% | 63.5% | Phage recombination related exonuclease (EC 3.1.11.-) |
| 76,139 | A→C | R58R (CGA→CGC) | 100% | 63.3% | Phage recombination related exonuclease (EC 3.1.11.-) |
| 76,145 | C→T | I60I (ATC→ATT) | 100% | 62.6% | Phage recombination related exonuclease (EC 3.1.11.-) |
| 76,151 | G→A | K62K (AAG→AAA) | 100% | 59.6% | Phage recombination related exonuclease (EC 3.1.11.-) |
| 76,157 | A→T | A64A (GCA→GCT) | | 52.8% | Phage recombination related exonuclease (EC 3.1.11.-) |
| 76,157 | 2 bp→TC | coding (192-193/1839 nt) | 100% | | Phage recombination related exonuclease (EC 3.1.11.-) |
| 76,158 | T→C | L65L (TTG→CTG) | | 52.2% | Phage recombination related exonuclease (EC 3.1.11.-) |
| 76,17 | A→T | N69Y (AAT→TAT) | 94.1% | 41.7% | Phage recombination related exonuclease (EC 3.1.11.-) |
| 76,171 | A→C | N69T (AAT→ACT) | 93.8% | 41.5% | Phage recombination related exonuclease (EC 3.1.11.-) |
| 76,173 | G→T | A70S (GCC→TCC) | 93.6% | 42.5% | Phage recombination related exonuclease (EC 3.1.11.-) |

| | | | | | |
|---|---|---|---|---|---|
| 76,175 | C→T | A70A (GCC→GCT) | 93.9% | 43.0% | Phage recombination related exonuclease (EC 3.1.11.-) |
| 76,178 | T→G | P71P (CCT→CCG) | 92.9% | 41.7% | Phage recombination related exonuclease (EC 3.1.11.-) |
| 76,181 | G→A | K72K (AAG→AAA) | 100% | | Phage recombination related exonuclease (EC 3.1.11.-) |
| 76,187 | A→G | E74E (GAA→GAG) | 100% | 45.2% | Phage recombination related exonuclease (EC 3.1.11.-) |
| 76,19 | C→T | Y75Y (TAC→TAT) | 100% | 52.6% | Phage recombination related exonuclease (EC 3.1.11.-) |
| 76,205 | C→T | Y80Y (TAC→TAT) | 94.9% | 54.9% | Phage recombination related exonuclease (EC 3.1.11.-) |
| 76,208 | T→C | F81F (TTT→TTC) | 100% | 54.4% | Phage recombination related exonuclease (EC 3.1.11.-) |
| 76,211 | A→G | S82S (TCA→TCG) | 94.2% | 53.9% | Phage recombination related exonuclease (EC 3.1.11.-) |
| 76,22 | T→A | D85E (GAT→GAA) | 94.7% | 60.2% | Phage recombination related exonuclease (EC 3.1.11.-) |
| 76,232 | G→A | E89E (GAG→GAA) | 100% | 60.8% | Phage recombination related exonuclease (EC 3.1.11.-) |
| 76,244 | T→A | V93V (GTT→GTA) | 100% | 65.6% | Phage recombination related exonuclease (EC 3.1.11.-) |
| 76,247 | C→T | V94V (GTC→GTT) | 100% | 68.1% | Phage recombination related exonuclease (EC 3.1.11.-) |
| 76,268 | G→T | T101T (ACG→ACT) | | 70.9% | Phage recombination related exonuclease (EC 3.1.11.-) |
| 76,268 | 2 bp→TC | coding (303-304/1839 nt) | 100% | | Phage recombination related exonuclease (EC 3.1.11.-) |
| 76,269 | T→C | L102L (TTG→CTG) | | 71.2% | Phage recombination related exonuclease (EC 3.1.11.-) |
| 76,271 | G→T | L102F (TTG→TTT) | 100% | 71.3% | Phage recombination related exonuclease (EC 3.1.11.-) |
| 76,283 | A→G | G106G (GGA→GGG) | 100% | 74.7% | Phage recombination related exonuclease (EC 3.1.11.-) |
| 76,301 | T→C | H112H (CAT→CAC) | 100% | 75.3% | Phage recombination related exonuclease (EC 3.1.11.-) |
| 76,319 | T→C | Y118Y (TAT→TAC) | 100% | 77.2% | Phage recombination related exonuclease (EC 3.1.11.-) |
| 76,352 | T→C | F129F (TTT→TTC) | 100% | 75.4% | Phage recombination related exonuclease (EC 3.1.11.-) |
| 76,385 | G→A | V140V (GTG→GTA) | 100% | 74.5% | Phage recombination related exonuclease (EC 3.1.11.-) |
| 76,4 | T→C | D145D (GAT→GAC) | 100% | 76.0% | Phage recombination related exonuclease (EC 3.1.11.-) |
| 76,446 | C→T | L161L (CTG→TTG) | 100% | 75.2% | Phage recombination related exonuclease (EC 3.1.11.-) |
| 76,46 | G→A | E165E (GAG→GAA) | 100% | 75.5% | Phage recombination related exonuclease (EC 3.1.11.-) |
| 76,463 | A→G | Q166Q (CAA→CAG) | 100% | 77.0% | Phage recombination related exonuclease (EC 3.1.11.-) |
| 76,478 | G→A | S171S (TCG→TCA) | 100% | 76.3% | Phage recombination related exonuclease (EC 3.1.11.-) |
| 76,485 | G→A | V174I (GTT→ATT) | 100% | 76.1% | Phage recombination related exonuclease (EC 3.1.11.-) |
| 76,493 | T→A | A176A (GCT→GCA) | 100% | 77.0% | Phage recombination related exonuclease (EC 3.1.11.-) |

| | | | | | | |
|---|---|---|---|---|---|---|
| 76,505 | G→A | E180E (GAG→GAA) | | 100% | 78.7% | Phage recombination related exonuclease (EC 3.1.11.-) |
| 76,527 | C→T | L188L (CTG→TTG) | | 100% | 79.2% | Phage recombination related exonuclease (EC 3.1.11.-) |
| 76,538 | G→A | Q191Q (CAG→CAA) | | 100% | 77.6% | Phage recombination related exonuclease (EC 3.1.11.-) |
| 76,563 | A→G | N200D (AAT→GAT) | | 100% | 80.8% | Phage recombination related exonuclease (EC 3.1.11.-) |
| 76,568 | G→A | G201G (GGG→GGA) | | 100% | 80.4% | Phage recombination related exonuclease (EC 3.1.11.-) |
| 76,571 | A→G | K202K (AAA→AAG) | | 100% | 80.0% | Phage recombination related exonuclease (EC 3.1.11.-) |
| 76,578 | C→T | L205L (CTA→TTA) | | 100% | 80.6% | Phage recombination related exonuclease (EC 3.1.11.-) |
| 76,61 | G→A | P215P (CCG→CCA) | | 100% | 81.3% | Phage recombination related exonuclease (EC 3.1.11.-) |
| 76,7 | G→A | K245K (AAG→AAA) | | 100% | 81.0% | Phage recombination related exonuclease (EC 3.1.11.-) |
| 76,748 | A→G | E261E (GAA→GAG) | | 100% | 83.3% | Phage recombination related exonuclease (EC 3.1.11.-) |
| 76,763 | G→A | L266L (TTG→TTA) | | 100% | 79.8% | Phage recombination related exonuclease (EC 3.1.11.-) |
| 76,775 | C→T | T270T (ACC→ACT) | | 100% | 79.2% | Phage recombination related exonuclease (EC 3.1.11.-) |
| 76,836 | A→G | S291G (AGC→GGC) | | 100% | 81.3% | Phage recombination related exonuclease (EC 3.1.11.-) |
| 76,892 | C→T | C309C (TGC→TGT) | | 100% | 80.9% | Phage recombination related exonuclease (EC 3.1.11.-) |
| 76,898 | T→C | T311T (ACT→ACC) | | 100% | 79.4% | Phage recombination related exonuclease (EC 3.1.11.-) |
| 76,901 | C→T | C312C (TGC→TGT) | | 100% | 78.6% | Phage recombination related exonuclease (EC 3.1.11.-) |
| 76,928 | T→G | A321A (GCT→GCG) | | 100% | 78.2% | Phage recombination related exonuclease (EC 3.1.11.-) |
| 76,954 | T→C | I330T (ATA→ACA) | | 100% | 80.3% | Phage recombination related exonuclease (EC 3.1.11.-) |
| 76,999 | G→A | R345K (AGG→AAG) | | 100% | 81.2% | Phage recombination related exonuclease (EC 3.1.11.-) |
| 77,039 | G→A | A358A (GCG→GCA) | 100% | 100% | 80.6% | Phage recombination related exonuclease (EC 3.1.11.-) |
| 77,054 | G→A | K363K (AAG→AAA) | 100% | 100% | 79.9% | Phage recombination related exonuclease (EC 3.1.11.-) |
| 77,069 | A→G | K368K (AAA→AAG) | 100% | 100% | 80.1% | Phage recombination related exonuclease (EC 3.1.11.-) |
| 77,075 | G→A | V370V (GTG→GTA) | 100% | 100% | 79.7% | Phage recombination related exonuclease (EC 3.1.11.-) |
| 77,09 | G→A | E375E (GAG→GAA) | 100% | 100% | 77.9% | Phage recombination related exonuclease (EC 3.1.11.-) |
| 77,111 | C→T | D382D (GAC→GAT) | | | 77.4% | Phage recombination related exonuclease (EC 3.1.11.-) |
| 77,111 | 2 bp→TA | coding (1146-1147/1839 nt) | 100% | 100% | | Phage recombination related exonuclease (EC 3.1.11.-) |
| 77,112 | G→A | A383T (GCT→ACT) | | | 77.0% | Phage recombination related exonuclease (EC 3.1.11.-) |
| 77,173 | C→A | A403E (GCA→GAA) | 100% | 100% | 77.4% | Phage recombination related exonuclease (EC 3.1.11.-) |

| | | | | | | |
|---|---|---|---|---|---|---|
| 77,18 | T→C | A405A (GCT→GCC) | 100% | 100% | 76.9% | Phage recombination related exonuclease (EC 3.1.11.-) |
| 77,186 | A→C | G407G (GGA→GGC) | 100% | 100% | 76.5% | Phage recombination related exonuclease (EC 3.1.11.-) |
| 77,202 | G→A | V413I (GTT→ATT) | 100% | 100% | 74.5% | Phage recombination related exonuclease (EC 3.1.11.-) |
| 77,228 | C→T | V421V (GTC→GTT) | 100% | 100% | 77.3% | Phage recombination related exonuclease (EC 3.1.11.-) |
| 77,255 | A→G | A430A (GCA→GCG) | 100% | 100% | 77.8% | Phage recombination related exonuclease (EC 3.1.11.-) |
| 77,258 | G→A | K431K (AAG→AAA) | 100% | 100% | 76.5% | Phage recombination related exonuclease (EC 3.1.11.-) |
| 77,279 | A→T | A438A (GCA→GCT) | 100% | 100% | 79.4% | Phage recombination related exonuclease (EC 3.1.11.-) |
| 77,282 | A→G | E439E (GAA→GAG) | 100% | 100% | 80.2% | Phage recombination related exonuclease (EC 3.1.11.-) |
| 77,292 | A→T | T443S (ACT→TCT) | 100% | 100% | 78.8% | Phage recombination related exonuclease (EC 3.1.11.-) |
| 77,312 | T→G | L449L (CTT→CTG) | 100% | 100% | 74.8% | Phage recombination related exonuclease (EC 3.1.11.-) |
| 77,322 | G→T | A453S (GCC→TCA) | 100% | 100% | 72.0% | Phage recombination related exonuclease (EC 3.1.11.-) |
| 77,324 | C→A | A453S (GCC→TCA) | 100% | 100% | 73.7% | Phage recombination related exonuclease (EC 3.1.11.-) |
| 77,333 | A→T | V456V (GTA→GTT) | 100% | 100% | 74.3% | Phage recombination related exonuclease (EC 3.1.11.-) |
| 77,351 | A→G | L462L (CTA→CTG) | 100% | 100% | 71.5% | Phage recombination related exonuclease (EC 3.1.11.-) |
| 77,354 | T→C | I463I (ATT→ATC) | 100% | 100% | 71.3% | Phage recombination related exonuclease (EC 3.1.11.-) |
| 77,357 | T→C | A464A (GCT→GCC) | 100% | 100% | 71.2% | Phage recombination related exonuclease (EC 3.1.11.-) |
| 77,366 | G→A | K467K (AAG→AAA) | 100% | 100% | | Phage recombination related exonuclease (EC 3.1.11.-) |
| 77,369 | C→T | N468N (AAC→AAT) | 100% | 100% | 67.3% | Phage recombination related exonuclease (EC 3.1.11.-) |
| 77,372 | A→G | L469L (CTA→CTG) | 100% | 100% | 68.7% | Phage recombination related exonuclease (EC 3.1.11.-) |
| 77,378 | A→G | G471G (GGA→GGG) | 100% | 100% | 71.2% | Phage recombination related exonuclease (EC 3.1.11.-) |
| 77,396 | C→T | S477S (AGC→AGT) | 100% | 100% | 68.8% | Phage recombination related exonuclease (EC 3.1.11.-) |
| 77,408 | C→T | F481F (TTC→TTT) | 100% | 100% | 69.1% | Phage recombination related exonuclease (EC 3.1.11.-) |
| 77,414 | A→G | E483E (GAA→GAG) | | | 70.2% | Phage recombination related exonuclease (EC 3.1.11.-) |
| 77,414 | 2 bp→GC | coding (1449-1450/1839 nt) | 100% | 100% | | Phage recombination related exonuclease (EC 3.1.11.-) |
| 77,415 | T→C | L484L (TTA→CTA) | | | 68.3% | Phage recombination related exonuclease (EC 3.1.11.-) |
| 77,417 | A→T | L484F (TTA→TTT) | 100% | 100% | 68.4% | Phage recombination related exonuclease (EC 3.1.11.-) |
| 77,42 | T→C | I485I (ATT→ATC) | 100% | 100% | 68.5% | Phage recombination related exonuclease (EC 3.1.11.-) |
| 77,429 | C→T | Y488Y (TAC→TAT) | 86.4% | 100% | 71.6% | Phage recombination related exonuclease (EC 3.1.11.-) |

| | | | | | | |
|---|---|---|---|---|---|---|
| 77,447 | G→C | G494G (GGG→GGC) | 100% | 100% | 76.5% | Phage recombination related exonuclease (EC 3.1.11.-) |
| 77,522 | C→T | T519T (ACC→ACT) | 100% | 100% | 80.3% | Phage recombination related exonuclease (EC 3.1.11.-) |
| 77,54 | T→C | S525S (TCT→TCC) | 100% | 100% | 80.9% | Phage recombination related exonuclease (EC 3.1.11.-) |
| 77,561 | T→C | I532I (ATT→ATC) | 100% | 100% | 81.6% | Phage recombination related exonuclease (EC 3.1.11.-) |
| 77,588 | A→G | R541R (CGA→CGG) | 100% | 100% | 81.5% | Phage recombination related exonuclease (EC 3.1.11.-) |
| 77,594 | A→G | L543L (CTA→CTG) | | | 81.8% | Phage recombination related exonuclease (EC 3.1.11.-) |
| 77,594 | 2 bp→GT | coding (1629-1630/1839 nt) | 100% | 100% | | Phage recombination related exonuclease (EC 3.1.11.-) |
| 77,595 | C→T | L544L (CTA→TTA) | | | 80.2% | Phage recombination related exonuclease (EC 3.1.11.-) |
| 77,663 | G→A | T566T (ACG→ACA) | 100% | 100% | 93.0% | Phage recombination related exonuclease (EC 3.1.11.-) |
| 77,688 | T→C | L575L (TTA→CTA) | 100% | 100% | 93.8% | Phage recombination related exonuclease (EC 3.1.11.-) |
| 77,696 | C→T | N577N (AAC→AAT) | 100% | 100% | 93.7% | Phage recombination related exonuclease (EC 3.1.11.-) |
| 77,732 | C→T | H589H (CAC→CAT) | 100% | 100% | 94.7% | Phage recombination related exonuclease (EC 3.1.11.-) |
| 77,75 | G→A | L595L (TTG→TTA) | 100% | 100% | 92.9% | Phage recombination related exonuclease (EC 3.1.11.-) |
| 77,753 | T→C | A596A (GCT→GCC) | 100% | 100% | 93.8% | Phage recombination related exonuclease (EC 3.1.11.-) |
| 77,768 | C→T | V601V (GTC→GTT) | 100% | 100% | 92.3% | Phage recombination related exonuclease (EC 3.1.11.-) |
| 77,771 | G→A | K602K (AAG→AAA) | 100% | 100% | 100% | Phage recombination related exonuclease (EC 3.1.11.-) |
| 77,795 | T→C | Y610Y (TAT→TAC) | 100% | 100% | 90.0% | Phage recombination related exonuclease (EC 3.1.11.-) |
| 77,801 | G→A | E612E (GAG→GAA) | 100% | 100% | 89.5% | Phage recombination related exonuclease (EC 3.1.11.-) |
| 77,813 | T→C | A2A (GCT→GCC) | 94.2% | 100% | 89.8% | Type II restriction endonuclease |
| 77,828 | A→G | E7E (GAA→GAG) | 89.2% | 100% | 83.3% | Type II restriction endonuclease |
| 77,834 | A→T | G9G (GGA→GGT) | 88.6% | 100% | 83.4% | Type II restriction endonuclease |
| 77,837 | A→G | K10K (AAA→AAG) | 89.3% | | 83.4% | Type II restriction endonuclease |
| 77,837 | 2 bp→GC | coding (30-31/483 nt) | | 100% | | Type II restriction endonuclease |
| 77,838 | A→C | R11R (AGA→CGA) | 88.6% | | 83.1% | Type II restriction endonuclease |
| 77,84 | A→T | R11S (AGA→AGT) | 87.0% | 100% | 82.1% | Type II restriction endonuclease |
| 77,842 | C→G | A12G (GCT→GGT) | 88.6% | 100% | 82.5% | Type II restriction endonuclease |
| 77,846 | G→A | E13E (GAG→GAA) | 88.0% | 100% | 81.5% | Type II restriction endonuclease |
| 77,852 | A→G | Q15Q (CAA→CAG) | 91.3% | 100% | 84.2% | Type II restriction endonuclease |
| 77,867 | T→A | L20L (CTT→CTA) | 86.4% | | 78.5% | Type II restriction endonuclease |
| 77,867 | 2 bp→AC | coding (60-61/483 nt) | | 100% | | Type II restriction endonuclease |
| 77,868 | A→C | R21R (AGA→CGA) | 87.3% | | 77.6% | Type II restriction endonuclease |
| 77,87 | A→T | R21S (AGA→AGT) | 87.8% | 100% | 76.6% | Type II restriction endonuclease |
| 77,876 | A→T | R23R (CGA→CGT) | 87.3% | 100% | 78.2% | Type II restriction endonuclease |
| 77,88 | A→G | K25E (AAG→GAG) | 87.8% | | 78.1% | Type II restriction endonuclease |
| 77,88 | 3 bp→GGT | coding (73-75/483 nt) | | 100% | | Type II restriction endonuclease |
| 77,881 | A→G | K25R (AAG→AGG) | 86.8% | | 77.9% | Type II restriction endonuclease |
| 77,882 | G→T | K25N (AAG→AAT) | 87.3% | | 76.9% | Type II restriction endonuclease |

| | | | | | | |
|---|---|---|---|---|---|---|
| 77,885 | A→T | L26L (CTA→CTT) | 87.6% | 100% | 78.0% | Type II restriction endonuclease |
| 77,894 | G→A | E29E (GAG→GAA) | 88.4% | | 78.1% | Type II restriction endonuclease |
| 77,894 | 2 bp→AC | coding (87-88/483 nt) | | 100% | | Type II restriction endonuclease |
| 77,895 | A→C | R30R (AGA→CGA) | 88.3% | | 78.6% | Type II restriction endonuclease |
| 77,897 | A→T | R30S (AGA→AGT) | 88.5% | 100% | 78.6% | Type II restriction endonuclease |
| 77,9 | G→T | V31V (GTG→GTT) | 88.2% | 100% | 77.4% | Type II restriction endonuclease |
| 77,934 | C→T | L43L (CTG→TTG) | 100% | 100% | 84.0% | Type II restriction endonuclease |
| 77,939 | A→G | K44K (AAA→AAG) | 100% | 100% | 85.4% | Type II restriction endonuclease |
| 77,965 | C→G | T53R (ACA→AGA) | 88.1% | 93.9% | 77.6% | Type II restriction endonuclease |
| 77,966 | A→C | T53T (ACA→ACC) | 88.5% | 100% | 78.4% | Type II restriction endonuclease |
| 77,969 | C→A | G54G (GGC→GGA) | 88.2% | 93.8% | 77.8% | Type II restriction endonuclease |
| 77,97 | A→C | K55Q (AAA→CAA) | 89.1% | 94.3% | 78.0% | Type II restriction endonuclease |
| 77,972 | A→C | K55N (AAA→AAC) | 89.1% | 94.0% | 78.1% | Type II restriction endonuclease |
| 77,975 | C→T | I56I (ATC→ATT) | 89.6% | 100% | 79.9% | Type II restriction endonuclease |
| 77,981 | G→A | K58K (AAG→AAA) | 90.4% | 100% | 81.7% | Type II restriction endonuclease |
| 77,987 | T→C | C60C (TGT→TGC) | 91.3% | 100% | 82.6% | Type II restriction endonuclease |
| 77,99 | T→C | F61F (TTT→TTC) | 90.5% | 100% | 81.1% | Type II restriction endonuclease |
| 77,993 | G→A | E62E (GAG→GAA) | 90.4% | 100% | 80.8% | Type II restriction endonuclease |
| 77,996 | A→T | V63V (GTA→GTT) | 92.3% | 100% | 80.8% | Type II restriction endonuclease |
| 77,999 | A→G | K64K (AAA→AAG) | 100% | | 82.3% | Type II restriction endonuclease |
| 77,999 | 4 bp→GTGG | coding (192-195/483 nt) | | 100% | | Type II restriction endonuclease |
| 78 | C→T | H65Y (CAT→TAT) | 91.6% | | 82.3% | Type II restriction endonuclease |
| 78,001 | A→G | H65R (CAT→CGT) | 100% | | 82.5% | Type II restriction endonuclease |
| 78,002 | T→G | H65Q (CAT→CAG) | 90.2% | | 82.2% | Type II restriction endonuclease |
| 78,016 | G→A | S70N (AGT→AAT) | 93.4% | 100% | 83.8% | Type II restriction endonuclease |
| 78,089 | T→C | G94G (GGT→GGC) | 100% | 100% | 87.3% | Type II restriction endonuclease |
| 78,107 | G→A | K100K (AAG→AAA) | 100% | 100% | 89.6% | Type II restriction endonuclease |
| 78,248 | G→A | E147E (GAG→GAA) | 100% | 100% | 93.9% | Type II restriction endonuclease |
| 78,364 | A→C | V25V (GTA→GTC) | 100% | 100% | 94.3% | Phage endonuclease |
| 78,367 | C→T | D26D (GAC→GAT) | 100% | 100% | 94.5% | Phage endonuclease |
| 78,388 | A→C | R33R (CGA→CGC) | 100% | 100% | 92.3% | Phage endonuclease |
| 78,4 | T→C | N37N (AAT→AAC) | 100% | 100% | 91.9% | Phage endonuclease |
| 78,409 | G→A | K40K (AAG→AAA) | 100% | 100% | 100% | Phage endonuclease |
| 78,442 | T→C | I51I (ATT→ATC) | 94.3% | 100% | 87.1% | Phage endonuclease |
| 78,445 | A→G | Q52Q (CAA→CAG) | 94.3% | 100% | 86.5% | Phage endonuclease |
| 78,449 | T→C | L54L (TTA→CTA) | 94.3% | 100% | 87.2% | Phage endonuclease |
| 78,46 | C→T | S57S (TCC→TCT) | 88.2% | 100% | 79.2% | Phage endonuclease |
| 78,463 | C→T | Y58Y (TAC→TAT) | 86.7% | 100% | 80.0% | Phage endonuclease |
| 78,466 | C→T | S59S (TCC→TCT) | 88.1% | 100% | 80.5% | Phage endonuclease |
| 78,469 | A→T | A60A (GCA→GCT) | 87.9% | 100% | 81.3% | Phage endonuclease |
| 78,475 | T→C | T62T (ACT→ACC) | 88.3% | 100% | 80.7% | Phage endonuclease |
| 78,478 | G→A | T63T (ACG→ACA) | 87.7% | 100% | 81.2% | Phage endonuclease |
| 78,481 | T→A | I64I (ATT→ATA) | 87.0% | 100% | 81.5% | Phage endonuclease |
| 78,485 | C→T | L66L (CTA→TTA) | 89.9% | 100% | 81.8% | Phage endonuclease |
| 78,49 | T→G | G67G (GGT→GGG) | 90.5% | 100% | 82.4% | Phage endonuclease |
| 78,496 | G→A | K69K (AAG→AAA) | 88.6% | 100% | 79.6% | Phage endonuclease |
| 78,502 | A→G | K71K (AAA→AAG) | 87.7% | 100% | 79.8% | Phage endonuclease |
| 78,507 | T→C | V73A (GTA→GCA) | 88.7% | 100% | 77.3% | Phage endonuclease |
| 78,511 | T→C | F74F (TTT→TTC) | | 100% | 56.1% | Phage endonuclease |
| 78,514 | T→C | R75R (CGT→CGC) | | 91.2% | 58.3% | Phage endonuclease |
| 78,514 | 2 bp→CT | coding (225-226/876 nt) | 100% | | | Phage endonuclease |
| 78,515 | C→T | L76L (CTA→TTA) | | 90.6% | 57.6% | Phage endonuclease |
| 78,52 | A→G | E77E (GAA→GAG) | 100% | 100% | 61.9% | Phage endonuclease |

| | | | | | | |
|---|---|---|---|---|---|---|
| 78,523 | T→C | H78H (CAT→CAC) | 100% | 88.7% | 60.1% | Phage endonuclease |
| 78,526 | A→T | L79L (CTA→CTT) | 100% | 89.7% | 58.5% | Phage endonuclease |
| 78,529 | A→T | P80P (CCA→CCT) | 100% | 85.5% | 62.4% | Phage endonuclease |
| 78,539 | A→G | S84G (AGT→GGG) | 100% | 100% | 68.3% | Phage endonuclease |
| 78,541 | T→G | S84G (AGT→GGG) | 100% | 92.3% | 70.1% | Phage endonuclease |
| 78,547 | T→A | R86R (CGT→CGA) | 100% | 85.1% | 55.0% | Phage endonuclease |
| 78,553 | A→G | E88E (GAA→GAG) | 100% | 82.1% | 51.8% | Phage endonuclease |
| 78,556 | G→A | K89K (AAG→AAA) | 100% | 82.0% | 52.4% | Phage endonuclease |
| 78,559 | T→C | Y90Y (TAT→TAC) | 100% | 82.2% | 50.6% | Phage endonuclease |
| 78,56 | G→T | A91S (GCA→TCA) | 81.9% | 81.9% | 50.5% | Phage endonuclease |
| 78,565 | A→G | Q92Q (CAA→CAG) | 100% | 91.1% | 65.9% | Phage endonuclease |
| 78,571 | A→G | T94T (ACA→ACG) | 100% | 92.6% | 68.8% | Phage endonuclease |
| 78,577 | A→G | E96E (GAA→GAG) | 100% | 92.2% | 68.5% | Phage endonuclease |
| 78,586 | A→G | A99A (GCA→GCG) | | 93.5% | 70.6% | Phage endonuclease |
| 78,586 | 2 bp→GT | coding (297-298/876 nt) | 100% | | | Phage endonuclease |
| 78,587 | C→T | L100L (CTG→TTG) | | 92.6% | 68.7% | Phage endonuclease |
| 78,589 | G→A | L100L (CTG→CTA) | 100% | 93.4% | 69.4% | Phage endonuclease |
| 78,595 | A→G | E102E (GAA→GAG) | 100% | 100% | 79.9% | Phage endonuclease |
| 78,604 | T→C | F105F (TTT→TTC) | 100% | 100% | 82.9% | Phage endonuclease |
| 78,61 | T→C | Y107Y (TAT→TAC) | 100% | 100% | 84.6% | Phage endonuclease |
| 78,628 | G→A | E113E (GAG→GAA) | 100% | 100% | 88.0% | Phage endonuclease |
| 78,631 | G→A | L114L (TTG→TTA) | 100% | 100% | 88.4% | Phage endonuclease |
| 78,664 | T→G | R125R (CGT→CGG) | 100% | 100% | 91.9% | Phage endonuclease |
| 78,706 | C→G | L139L (CTC→CTG) | 100% | 100% | 90.8% | Phage endonuclease |
| 78,709 | C→T | I140I (ATC→ATT) | 100% | 100% | 90.8% | Phage endonuclease |
| 78,734 | T→C | L149M (TTA→CTG) | 100% | 100% | 92.7% | Phage endonuclease |
| 78,736 | A→G | L149M (TTA→CTG) | 100% | 100% | 92.2% | Phage endonuclease |
| 78,739 | C→T | I150I (ATC→ATT) | 100% | 100% | 92.7% | Phage endonuclease |
| 78,745 | A→T | T152T (ACA→ACT) | 100% | 100% | 93.5% | Phage endonuclease |
| 78,767 | C→T | L160L (CTA→TTA) | 100% | 100% | 93.2% | Phage endonuclease |
| 78,772 | A→T | T161T (ACA→ACT) | 100% | 100% | 93.3% | Phage endonuclease |
| 78,781 | T→A | V164V (GTT→GTA) | 94.9% | 100% | 93.0% | Phage endonuclease |
| 78,787 | T→C | R166R (CGT→CGC) | 100% | 100% | 94.6% | Phage endonuclease |
| 78,79 | T→C | F167F (TTT→TTC) | 100% | 100% | 94.9% | Phage endonuclease |
| 78,851 | A→G | N188D (AAT→GAT) | 100% | 100% | 91.3% | Phage endonuclease |
| 78,871 | C→T | S194S (TCC→TCT) | 100% | 100% | 92.1% | Phage endonuclease |
| 78,874 | G→A | L195L (CTG→CTA) | 100% | 100% | 91.3% | Phage endonuclease |
| 78,889 | C→T | G200G (GGC→GGT) | 100% | 100% | 91.5% | Phage endonuclease |
| 78,916 | T→C | V209V (GTT→GTC) | 100% | 100% | 91.7% | Phage endonuclease |
| 78,919 | A→G | E210E (GAA→GAG) | 100% | 100% | 91.8% | Phage endonuclease |
| 78,973 | G→T | L228L (CTG→CTT) | 100% | 100% | 90.2% | Phage endonuclease |
| 78,991 | C→T | L234L (CTC→CTT) | 100% | 100% | 90.8% | Phage endonuclease |
| 79,051 | C→T | Y254Y (TAC→TAT) | 100% | 100% | 89.2% | Phage endonuclease |
| 79,117 | C→T | D276D (GAC→GAT) | 100% | 100% | 88.4% | Phage endonuclease |
| 79,176 | G→A | K5K (AAG→AAA) | | | 88.4% | Deoxyuridine 5'-triphosphate nucleotidohydrolase (EC 3.6.1.23) |
| 79,176 | 2 bp→AC | coding (15-16/447 nt) | 100% | 100% | | Deoxyuridine 5'-triphosphate nucleotidohydrolase (EC 3.6.1.23) |
| 79,177 | T→C | L6L (TTA→CTA) | | | 90.4% | Deoxyuridine 5'-triphosphate nucleotidohydrolase (EC 3.6.1.23) |
| 79,182 | C→T | T7T (ACC→ACT) | 100% | 100% | 89.9% | Deoxyuridine 5'-triphosphate nucleotidohydrolase (EC 3.6.1.23) |

| | | | | | | | |
|---|---|---|---|---|---|---|---|
| 79,242 | G→A | A27A (GCG→GCA) | 100% | 100% | | 90.9% | Deoxyuridine 5'-triphosphate nucleotidohydrolase (EC 3.6.1.23) |
| 79,296 | Δ1 bp | coding (135/447 nt) | 100% | 100% | | 84.9% | Deoxyuridine 5'-triphosphate nucleotidohydrolase (EC 3.6.1.23) |
| 79,300:1 | +A | coding (139/447 nt) | 93.7% | 100% | | 83.3% | Deoxyuridine 5'-triphosphate nucleotidohydrolase (EC 3.6.1.23) |
| 79,335 | T→A | R58R (CGT→CGA) | 100% | 100% | | 86.4% | Deoxyuridine 5'-triphosphate nucleotidohydrolase (EC 3.6.1.23) |
| 79,353 | G→A | V64V (GTG→GTA) | 100% | 100% | 72.2% | 87.6% | Deoxyuridine 5'-triphosphate nucleotidohydrolase (EC 3.6.1.23) |
| 79,359 | G→T | P66P (CCG→CCT) | 100% | 100% | 67.9% | 86.4% | Deoxyuridine 5'-triphosphate nucleotidohydrolase (EC 3.6.1.23) |
| 79,362 | T→C | R67R (CGT→CGC) | 100% | 100% | 67.6% | 86.4% | Deoxyuridine 5'-triphosphate nucleotidohydrolase (EC 3.6.1.23) |
| 79,365 | T→C | S68S (AGT→AGC) | 100% | 100% | 68.1% | 86.5% | Deoxyuridine 5'-triphosphate nucleotidohydrolase (EC 3.6.1.23) |
| 79,368 | C→T | S69S (TCC→TCT) | 100% | 100% | 65.2% | 86.3% | Deoxyuridine 5'-triphosphate nucleotidohydrolase (EC 3.6.1.23) |
| 79,392 | T→C | I77I (ATT→ATC) | 100% | 100% | 71.0% | 88.3% | Deoxyuridine 5'-triphosphate nucleotidohydrolase (EC 3.6.1.23) |
| 79,407 | A→G | G82G (GGA→GGG) | 100% | 100% | 65.0% | 88.4% | Deoxyuridine 5'-triphosphate nucleotidohydrolase (EC 3.6.1.23) |
| 79,413 | T→C | I84I (ATT→ATC) | 100% | 100% | 62.6% | 88.8% | Deoxyuridine 5'-triphosphate nucleotidohydrolase (EC 3.6.1.23) |
| 79,431 | A→T | G90G (GGA→GGT) | 100% | 100% | 57.0% | 87.2% | Deoxyuridine 5'-triphosphate nucleotidohydrolase (EC 3.6.1.23) |
| 79,449 | T→C | L96L (CTT→CTC) | 100% | 100% | 56.0% | 86.7% | Deoxyuridine 5'-triphosphate nucleotidohydrolase (EC 3.6.1.23) |
| 79,455 | C→T | N98N (AAC→AAT) | 100% | 100% | 52.5% | 84.0% | Deoxyuridine 5'-triphosphate nucleotidohydrolase (EC 3.6.1.23) |
| 79,458 | T→C | Y99Y (TAT→TAC) | 100% | 100% | 54.2% | 85.4% | Deoxyuridine 5'-triphosphate nucleotidohydrolase (EC 3.6.1.23) |
| 79,47 | T→G | I103M (ATT→ATG) | 100% | 100% | 50.7% | 82.2% | Deoxyuridine 5'-triphosphate nucleotidohydrolase (EC 3.6.1.23) |
| 79,482 | A→G | E107E (GAA→GAG) | 100% | 100% | 48.0% | 80.7% | Deoxyuridine 5'-triphosphate nucleotidohydrolase (EC 3.6.1.23) |
| 79,5 | T→C | C113C (TGT→TGC) | 100% | 100% | 41.0% | 76.6% | Deoxyuridine 5'-triphosphate nucleotidohydrolase (EC 3.6.1.23) |
| 79,504 | C→T | L115M (CTA→TTG) | 100% | 100% | 36.6% | 76.0% | Deoxyuridine 5'-triphosphate nucleotidohydrolase (EC 3.6.1.23) |
| 79,506 | A→G | L115M (CTA→TTG) | 100% | 100% | 37.4% | 76.0% | Deoxyuridine 5'-triphosphate nucleotidohydrolase (EC 3.6.1.23) |
| 79,524 | T→C | Y121Y (TAT→TAC) | 100% | 100% | 28.9% | 73.7% | Deoxyuridine 5'-triphosphate nucleotidohydrolase (EC 3.6.1.23) |
| 79,530 | T→A | T123T (ACT→ACA) | 100% | 100% | 28.8% | 70.5% | Deoxyuridine 5'-triphosphate nucleotidohydrolase (EC 3.6.1.23) |
| 79,545 | C→T | I128I (ATC→ATT) | 100% | 100% | 21.3% | 60.4% | Deoxyuridine 5'-triphosphate nucleotidohydrolase (EC 3.6.1.23) |
| 79,551 | C→T | D130D (GAC→GAT) | 100% | 100% | 20.2% | 57.0% | Deoxyuridine 5'-triphosphate nucleotidohydrolase (EC 3.6.1.23) |
| 79,559 | A→G | E133G (GAG→GGG) | 100% | 100% | 13.8% | 59.2% | Deoxyuridine 5'-triphosphate nucleotidohydrolase (EC 3.6.1.23) |
| 79,563 | G→A | E134E (GAG→GAA) | 100% | 100% | 14.1% | 57.1% | Deoxyuridine 5'-triphosphate nucleotidohydrolase (EC 3.6.1.23) |
| 79,567 | A→G | N136D (AAT→GAT) | 100% | 100% | 12.8% | 53.4% | Deoxyuridine 5'-triphosphate nucleotidohydrolase (EC 3.6.1.23) |

| | | | | | | | |
|---|---|---|---|---|---|---|---|
| 79,575 | G→T | G138G (GGG→GGT) | 100% | 100% | 11.0% | 53.8% | Deoxyuridine 5'-triphosphate nucleotidohydrolase (EC 3.6.1.23) |
| 79,584 | A→G | G141G (GGA→GGG) | 100% | 100% | 8.7% | 40.8% | Deoxyuridine 5'-triphosphate nucleotidohydrolase (EC 3.6.1.23) |
| 79,596 | A→G | S145S (TCA→TCG) | 100% | 100% | 5.8% | 33.1% | Deoxyuridine 5'-triphosphate nucleotidohydrolase (EC 3.6.1.23) |
| 79,601 | G→A | S147N (AGC→AAC) | ? | ? | | 28.9% | Deoxyuridine 5'-triphosphate nucleotidohydrolase (EC 3.6.1.23) |
| 79,602 | C→T | S147S (AGC→AGT) | 100% | ? | | 27.3% | Deoxyuridine 5'-triphosphate nucleotidohydrolase (EC 3.6.1.23) |
| 80,453 | G→A | H1192Y (CAT→TAT) | ? | ? | 100% | 100% | Phage long tail fiber pb1 |
| 80,495 | T→C | N1178D (AAC→GAC) | ? | ? | 100% | 100% | Phage long tail fiber pb1 |
| 80,557 | G→A | A1157V (GCC→GTC) | ? | ? | 100% | 100% | Phage long tail fiber pb1 |
| 84,368 | T→A | V27V (GTA→GTT) | 83.7% | 88.0% | 13.3% | 38.6% | Tail fiber p132 |
| 84,374 | C→T | M25I (ATG→ATA) | 77.5% | 100% | 12.4% | 35.4% | Tail fiber p132 |
| 84,376 | 2 bp→AG | coding (72-73/423 nt) | | 100% | | | Tail fiber p132 |
| 84,377 | A→G | F24F (TTT→TTC) | 79.8% | | 12.7% | 32.4% | Tail fiber p132 |
| 84,380 | T→C | Q23Q (CAA→CAG) | 78.3% | 100% | 12.8% | 33.4% | Tail fiber p132 |
| 84,386 | A→C | L21L (CTT→CTG) | 72.8% | 89.7% | 12.9% | 29.5% | Tail fiber p132 |
| 84,388 | G→T | L21I (CTT→ATT) | 72.8% | 89.6% | 12.9% | 29.1% | Tail fiber p132 |
| 84,398 | A→T | P17P (CCT→CCA) | 87.6% | 100% | 18.5% | 33.2% | Tail fiber p132 |
| 84,404 | G→T | N15K (AAC→AAA) | 82.6% | 92.7% | 17.6% | 32.1% | Tail fiber p132 |
| 84,415 | C→T | V12I (GTA→ATA) | 83.1% | 100% | 20.9% | 31.9% | Tail fiber p132 |
| 84,422 | A→G | D9D (GAT→GAC) | 84.8% | 100% | 19.9% | 32.6% | Tail fiber p132 |
| 84,425 | A→T | I8I (ATT→ATA) | 84.5% | 100% | 18.7% | 30.7% | Tail fiber p132 |
| 84,440 | T→A | T3T (ACA→ACT) | 83.2% | 100% | 22.1% | 31.4% | Tail fiber p132 |
| 84,454 | Δ1 bp | coding (2102/2103 nt) | 100% | 100% | | | Tail fiber pb4 |
| 84,468 | A→G | Y696Y (TAT→TAC) | 85.0% | 100% | 23.7% | 33.3% | Tail fiber pb4 |
| 84,496 | T→C | K687R (AAA→AGA) | 69.6% | 87.0% | 12.5% | 22.2% | Tail fiber pb4 |
| 85,149 | G→A | N469N (AAC→AAT) | 23.8% | | 7.9% | 9.2% | Tail fiber pb4 |
| 85,155 | T→A | T467T (ACA→ACT) | 22.8% | 79.0% | 11.7% | 13.1% | Tail fiber pb4 |
| 85,158 | A→T | G466G (GGT→GGA) | 21.3% | 100% | 11.9% | 12.6% | Tail fiber pb4 |
| 85,164 | T→A | V464V (GTA→GTT) | 18.5% | 82.7% | 11.8% | 12.8% | Tail fiber pb4 |
| 85,170 | C→T | G462G (GGG→GGA) | 19.7% | 100% | 10.8% | 12.7% | Tail fiber pb4 |
| 85,182 | C→A | R458S (AGG→AGT) | 20.4% | 100% | 10.0% | 12.8% | Tail fiber pb4 |
| 85,184 | T→G | R458R (AGG→CGG) | 18.6% | 100% | 10.4% | 12.5% | Tail fiber pb4 |
| 85,187 | A→G | L457L (TTA→CTA) | 18.5% | | 10.5% | 13.3% | Tail fiber pb4 |
| 85,187 | 4 bp→GCAC | coding (1366-1369/2103 nt) | | 100% | | | Tail fiber pb4 |
| 85,188 | T→C | T456T (ACA→ACG) | 19.0% | | 10.7% | 12.5% | Tail fiber pb4 |
| 85,189 | G→A | T456I (ACA→ATA) | 18.6% | | 9.5% | 11.8% | Tail fiber pb4 |
| 85,190 | T→C | T456A (ACA→GCA) | 21.3% | | 10.2% | 12.5% | Tail fiber pb4 |
| 85,194 | T→G | G454G (GGA→GGC) | 20.0% | 100% | 11.8% | 12.7% | Tail fiber pb4 |
| 85,203 | A→G | R451R (CGT→CGC) | 18.9% | 100% | 12.1% | 13.4% | Tail fiber pb4 |
| 85,206 | A→T | I450I (ATT→ATA) | 17.3% | 100% | 10.2% | 13.9% | Tail fiber pb4 |
| 85,209 | G→C | Y449* (TAC→TAG) | 18.2% | | 11.0% | 13.4% | Tail fiber pb4 |
| 85,209 | 2 bp→CC | coding (1346-1347/2103 nt) | | 100% | | | Tail fiber pb4 |
| 85,210 | T→C | Y449C (TAC→TGC) | 18.1% | | 10.7% | 13.6% | Tail fiber pb4 |
| 85,215 | T→A | T447T (ACA→ACT) | 51.5% | 100% | 23.5% | 29.4% | Tail fiber pb4 |
| 85,224 | G→A | G444G (GGC→GGT) | 63.2% | 100% | 23.1% | 31.3% | Tail fiber pb4 |
| 85,227 | A→T | I443I (ATT→ATA) | 63.4% | 100% | 22.5% | 30.7% | Tail fiber pb4 |
| 85,233 | C→T | L441L (CTG→CTA) | 69.3% | 100% | 26.0% | 35.8% | Tail fiber pb4 |
| 85,236 | T→C | K440K (AAA→AAG) | 75.9% | 100% | 26.8% | 36.5% | Tail fiber pb4 |

| | | | | | | | |
|---|---|---|---|---|---|---|---|
| 85,242 | T→C | K438K (AAA→AAG) | 81.1% | 100% | 31.7% | 44.8% | Tail fiber pb4 |
| 85,250 | T→A | T436S (ACT→TCT) | 86.2% | 100% | 35.7% | 48.3% | Tail fiber pb4 |
| 85,253 | C→T | A435T (GCT→ACT) | 86.2% | 100% | 34.7% | 48.4% | Tail fiber pb4 |
| 85,263 | A→T | G431G (GGT→GGA) | 87.6% | 100% | 38.6% | 56.2% | Tail fiber pb4 |
| 85,282 | C→T | R425K (AGA→AAA) | 90.2% | 100% | 45.6% | 62.2% | Tail fiber pb4 |
| 85,288 | C→T | G423D (GGT→GAT) | 89.2% | 100% | 44.9% | 60.2% | Tail fiber pb4 |
| 85,290 | A→C | G422G (GGT→GGG) | 100% | 100% | | | Tail fiber pb4 |
| 85,302 | G→A | G418G (GGC→GGT) | 88.9% | 100% | 45.6% | 63.2% | Tail fiber pb4 |
| 85,314 | A→G | I414I (ATT→ATC) | 88.6% | 100% | 49.9% | 66.6% | Tail fiber pb4 |
| 85,320 | T→A | P412P (CCA→CCT) | 87.3% | 100% | 50.4% | 66.6% | Tail fiber pb4 |
| 85,332 | A→G | G408G (GGT→GGC) | 87.3% | 100% | 50.9% | 70.6% | Tail fiber pb4 |
| 85,335 | G→A | T407T (ACC→ACT) | 87.7% | 100% | 51.8% | 70.3% | Tail fiber pb4 |
| 85,340 | A→G | L406L (TTA→CTA) | 88.3% | 100% | 54.0% | 72.6% | Tail fiber pb4 |
| 85,344 | A→G | T404T (ACT→ACC) | 87.4% | 100% | 52.9% | 71.8% | Tail fiber pb4 |
| 85,350 | G→T | V402V (GTC→GTA) | 88.2% | 100% | 55.6% | 74.3% | Tail fiber pb4 |
| 85,357 | C→T | S400N (AGC→AAC) | 87.9% | 100% | 59.4% | 74.0% | Tail fiber pb4 |
| 85,365 | G→A | N397N (AAC→AAT) | 88.7% | 100% | 59.5% | 72.6% | Tail fiber pb4 |
| 85,368 | A→G | I396I (ATT→ATC) | 87.6% | 100% | 60.2% | 72.9% | Tail fiber pb4 |
| 85,383 | T→A | V391V (GTA→GTT) | 88.4% | 100% | 62.8% | 73.4% | Tail fiber pb4 |
| 85,389 | T→A | G389G (GGA→GGT) | 88.1% | 100% | 62.4% | 74.6% | Tail fiber pb4 |
| 85,416 | A→C | P380P (CCT→CCG) | 92.7% | 100% | 63.7% | 76.4% | Tail fiber pb4 |
| 85,419 | G→A | D379D (GAC→GAT) | 92.9% | 100% | 64.0% | 76.8% | Tail fiber pb4 |
| 85,425 | T→A | S377S (TCA→TCT) | 91.7% | 100% | 63.8% | 76.0% | Tail fiber pb4 |
| 85,434 | G→T | A374A (GCC→GCA) | 91.7% | 100% | 67.1% | 76.6% | Tail fiber pb4 |
| 85,444 | C→T | G371E (GGA→GAA) | 90.7% | 100% | 70.5% | 78.8% | Tail fiber pb4 |
| 85,446 | C→A | A370A (GCG→GCT) | 89.7% | 100% | 70.0% | 78.7% | Tail fiber pb4 |
| 85,481 | C→T | A359T (GCT→ACT) | 91.4% | 100% | 67.3% | 77.3% | Tail fiber pb4 |
| 85,488 | T→G | I356I (ATA→ATC) | 88.1% | 100% | 63.7% | 72.1% | Tail fiber pb4 |
| 85,491 | C→T | L355L (CTG→CTA) | 88.5% | 100% | 62.9% | 71.8% | Tail fiber pb4 |
| 85,493 | G→A | L355L (CTG→TTG) | 88.3% | | 62.2% | 71.4% | Tail fiber pb4 |
| 85,493 | 2 bp→AA | coding (1062-1063/2103 nt) | | 100% | | | Tail fiber pb4 |
| 85,494 | G→A | F354F (TTC→TTT) | 88.5% | | 62.2% | 71.2% | Tail fiber pb4 |
| 85,497 | A→T | T353T (ACT→ACA) | 86.8% | 100% | 59.9% | 70.0% | Tail fiber pb4 |
| 85,503 | T→C | K351K (AAA→AAG) | 87.8% | 100% | 63.4% | 72.4% | Tail fiber pb4 |
| 85,509 | A→C | T349T (ACT→ACG) | 86.4% | 100% | 58.9% | 72.0% | Tail fiber pb4 |
| 85,512 | A→G | S348S (TCT→TCC) | 87.0% | 100% | 58.4% | 73.0% | Tail fiber pb4 |
| 85,515 | A→G | G347G (GGT→GGC) | 86.9% | 100% | 59.4% | 74.1% | Tail fiber pb4 |
| 85,524 | G→A | I344I (ATC→ATT) | 86.5% | 100% | 56.9% | 73.0% | Tail fiber pb4 |
| 85,545 | G→A | Y337Y (TAC→TAT) | 85.5% | 100% | 51.5% | 69.3% | Tail fiber pb4 |
| 85,554 | C→T | E334E (GAG→GAA) | 82.2% | 100% | 44.9% | 63.7% | Tail fiber pb4 |
| 85,557 | A→T | I333I (ATT→ATA) | 82.4% | 100% | 43.3% | 63.0% | Tail fiber pb4 |
| 85,572 | T→A | V328V (GTA→GTT) | 80.3% | 100% | 35.3% | 53.3% | Tail fiber pb4 |
| 85,587 | T→A | L323L (CTA→CTT) | 71.2% | 100% | 21.0% | 35.9% | Tail fiber pb4 |
| 85,590 | T→A | P322P (CCA→CCT) | 74.8% | 100% | 20.8% | 36.5% | Tail fiber pb4 |
| 85,593 | A→G | T321T (ACT→ACC) | 71.1% | 100% | 20.8% | 36.5% | Tail fiber pb4 |
| 85,602 | C→A | K318N (AAG→AAT) | 69.3% | 100% | 20.4% | 31.4% | Tail fiber pb4 |
| 85,617 | T→C | Q313Q (CAA→CAG) | 58.0% | 100% | | 12.7% | Tail fiber pb4 |
| 85,659 | G→C | A299A (GCC→GCG) | 49.9% | 100% | 19.0% | 34.8% | Tail fiber pb4 |
| 85,662 | A→T | I298I (ATT→ATA) | 51.6% | 100% | 19.0% | 35.0% | Tail fiber pb4 |
| 85,668 | C→G | S296S (TCG→TCC) | 68.3% | 100% | 25.7% | 42.2% | Tail fiber pb4 |
| 85,671 | C→G | V295V (GTG→GTC) | 70.0% | 100% | 25.6% | 43.5% | Tail fiber pb4 |
| 85,689 | T→A | V289V (GTA→GTT) | 82.0% | 100% | 29.7% | 44.6% | Tail fiber pb4 |

| | | | | | | | |
|---|---|---|---|---|---|---|---|
| 85,691 | C→T | V289I (GTA→ATA) | 81.4% | 100% | 27.7% | 42.8% | Tail fiber pb4 |
| 85,698 | T→C | P286P (CCA→CCG) | 85.4% | 100% | 28.5% | 49.2% | Tail fiber pb4 |
| 85,704 | T→C | S284S (TCA→TCG) | 86.4% | 100% | 33.7% | 52.1% | Tail fiber pb4 |
| 85,713 | T→A | T281T (ACA→ACT) | 87.2% | 100% | 37.0% | 55.7% | Tail fiber pb4 |
| 85,719 | T→C | S279S (TCA→TCG) | 86.5% | 100% | 35.1% | 56.9% | Tail fiber pb4 |
| 85,721 | A→C | S279A (TCA→GCA) | 86.6% | 100% | 34.9% | 56.4% | Tail fiber pb4 |
| 85,725 | A→T | A277A (GCT→GCA) | 91.3% | 100% | 35.2% | 58.7% | Tail fiber pb4 |
| 85,734 | C→T | V274V (GTG→GTA) | 89.4% | 100% | 42.4% | 62.0% | Tail fiber pb4 |
| 85,749 | T→A | A269A (GCA→GCT) | 90.8% | 100% | 50.6% | 66.8% | Tail fiber pb4 |
| 85,770 | A→T | A262A (GCT→GCA) | 90.7% | 100% | 52.8% | 65.4% | Tail fiber pb4 |
| 85,776 | C→T | E260E (GAG→GAA) | 91.1% | 100% | 50.8% | 63.1% | Tail fiber pb4 |
| 85,781 | C→T | A259T (GCA→ACA) | 90.1% | | 51.3% | 61.4% | Tail fiber pb4 |
| 85,781 | 2 bp→TG | coding (774-775/2103 nt) | | 100% | | | Tail fiber pb4 |
| 85,782 | T→G | S258S (TCA→TCC) | 90.7% | | 52.1% | 61.8% | Tail fiber pb4 |
| 85,794 | G→A | S254S (TCC→TCT) | 88.1% | 94.1% | 40.5% | 52.0% | Tail fiber pb4 |
| 85,797 | T→A | L253L (CTA→CTT) | 87.2% | 94.7% | 40.0% | 52.8% | Tail fiber pb4 |
| 85,799 | G→C | L253V (CTA→GTA) | 87.7% | 94.9% | 40.7% | 53.1% | Tail fiber pb4 |
| 85,800 | A→G | V252V (GTT→GTC) | 88.6% | 100% | 40.4% | 54.5% | Tail fiber pb4 |
| 85,803 | G→A | F251F (TTC→TTT) | 84.4% | 100% | 42.0% | 55.1% | Tail fiber pb4 |
| 85,809 | A→T | R249R (CGT→CGA) | 86.6% | 100% | 39.5% | 55.3% | Tail fiber pb4 |
| 85,815 | G→A | N247N (AAC→AAT) | 86.7% | 100% | 42.7% | 57.6% | Tail fiber pb4 |
| 85,821 | A→T | G245G (GGT→GGA) | 83.2% | 100% | 41.1% | 57.8% | Tail fiber pb4 |
| 85,824 | T→A | A244A (GCA→GCT) | 82.7% | 100% | 41.6% | 56.8% | Tail fiber pb4 |
| 85,826 | C→A | A244S (GCA→TCA) | 82.1% | | 42.9% | 55.8% | Tail fiber pb4 |
| 85,826 | 2 bp→AT | coding (729-730/2103 nt) | | 100% | | | Tail fiber pb4 |
| 85,827 | A→T | G243G (GGT→GGA) | 82.1% | | 42.6% | 55.8% | Tail fiber pb4 |
| 85,839 | A→C | D239E (GAT→GAG) | 75.0% | 100% | 38.2% | 49.6% | Tail fiber pb4 |
| 85,845 | T→C | Q237Q (CAA→CAG) | 71.8% | 100% | 37.3% | 43.8% | Tail fiber pb4 |
| 85,848 | C→T | V236V (GTG→GTA) | 68.5% | 100% | 34.7% | 43.2% | Tail fiber pb4 |
| 85,854 | T→A | L234L (CTA→CTT) | 63.4% | 100% | 22.8% | 38.2% | Tail fiber pb4 |
| 85,857 | A→G | D233D (GAT→GAC) | 63.7% | 100% | 23.1% | 36.6% | Tail fiber pb4 |
| 85,863 | G→T | R231R (CGC→CGA) | 48.6% | 100% | 14.2% | 20.2% | Tail fiber pb4 |
| 85,865 | G→T | R231S (CGC→AGC) | 48.5% | | 13.9% | 19.8% | Tail fiber pb4 |
| 85,865 | 2 bp→TC | coding (690-691/2103 nt) | ? | 100% | | | Tail fiber pb4 |
| 85,866 | T→C | E230E (GAA→GAG) | 50.2% | | 14.6% | 20.1% | Tail fiber pb4 |
| 85,875 | G→A | D227D (GAC→GAT) | 34.4% | 100% | 10.9% | 10.0% | Tail fiber pb4 |
| 85,878 | T→C | A226A (GCA→GCG) | 33.4% | 100% | 10.9% | 9.5% | Tail fiber pb4 |
| 86,445 | C→T | R37R (AGG→AGA) | 17.3% | | | | Tail fiber pb4 |
| 86,448 | G→A | G36G (GGC→GGT) | 17.2% | | | | Tail fiber pb4 |
| 86,451 | T→A | I35I (ATA→ATT) | 19.9% | | | | Tail fiber pb4 |
| 86,463 | A→G | D31D (GAT→GAC) | 21.8% | | | | Tail fiber pb4 |
| 86,468 | A→T | S30T (TCT→ACT) | 22.4% | | | | Tail fiber pb4 |
| 86,469 | G→A | Y29Y (TAC→TAT) | 23.0% | | | | Tail fiber pb4 |
| 86,472 | G→T | I28I (ATC→ATA) | 24.8% | | | | Tail fiber pb4 |
| 86,478 | A→G | H26H (CAT→CAC) | 32.5% | | | | Tail fiber pb4 |
| 86,486 | T→C | I24V (ATC→GTC) | 35.7% | | | | Tail fiber pb4 |
| 86,490 | C→T | A22A (GCG→GCA) | 32.6% | | | | Tail fiber pb4 |
| 86,495 | A→T | L21M (TTG→ATG) | 33.3% | | | | Tail fiber pb4 |
| 86,496 | C→T | T20T (ACG→ACA) | 33.0% | | | | Tail fiber pb4 |
| 86,508 | T→C | L16L (TTA→TTG) | 28.9% | | | | Tail fiber pb4 |
| 86,510 | A→T | L16I (TTA→ATA) | 27.7% | | | | Tail fiber pb4 |

| | | | | | | | |
|---|---|---|---|---|---|---|---|
| 86,511 | G→T | V15V (GTC→GTA) | 25.5% | | | | Tail fiber pb4 |
| 86,513 | C→T | V15I (GTC→ATC) | 31.4% | | | | Tail fiber pb4 |
| 86,514 | G→A | S14S (AGC→AGT) | 25.5% | | | | Tail fiber pb4 |
| 86,522 | G→A | L12L (CTA→TTA) | 27.6% | | | | Tail fiber pb4 |
| 86,532 | A→G | A8A (GCT→GCC) | 31.4% | | | | Tail fiber pb4 |
| 86,547 | T→C | S3S (TCA→TCG) | 31.4% | | | | Tail fiber pb4 |
| 87,125 | G→A | L761L (CTA→TTA) | 36.1% | | 7.7% | 8.0% | Tail fiber pb3 |
| 87,125 | 2 bp→AC | coding (2280-2281/2850 nt) | | 100% | | | Tail fiber pb3 |
| 87,126 | T→C | E760E (GAA→GAG) | 40.3% | | 10.1% | 9.0% | Tail fiber pb3 |
| 87,129 | C→A | G759G (GGG→GGT) | 36.6% | 100% | 8.1% | 8.4% | Tail fiber pb3 |
| 87,138 | A→T | G756G (GGT→GGA) | 48.9% | 100% | 14.1% | 16.7% | Tail fiber pb3 |
| 87,146 | C→A | A754S (GCT→TCT) | 46.3% | 100% | 14.1% | 17.1% | Tail fiber pb3 |
| 87,150 | T→A | I752I (ATA→ATT) | 51.2% | 100% | 14.1% | 17.4% | Tail fiber pb3 |
| 87,152 | T→C | I752V (ATA→GTA) | 48.8% | | 14.2% | 16.3% | Tail fiber pb3 |
| 87,152 | 2 bp→CT | coding (2253-2254/2850 nt) | ? | 100% | | | Tail fiber pb3 |
| 87,153 | C→T | M751I (ATG→ATA) | 48.6% | | 15.2% | 17.8% | Tail fiber pb3 |
| 87,155 | T→A | M751L (ATG→TTG) | 50.1% | 100% | 14.1% | 16.8% | Tail fiber pb3 |
| 87,162 | A→G | P748P (CCT→CCC) | 53.7% | 100% | 15.6% | 17.5% | Tail fiber pb3 |
| 87,165 | A→T | T747T (ACT→ACA) | 46.0% | 100% | 13.9% | 16.4% | Tail fiber pb3 |
| 87,168 | T→A | P746P (CCA→CCT) | 47.8% | 100% | 15.4% | 15.8% | Tail fiber pb3 |
| 87,171 | T→A | T745T (ACA→ACT) | 47.6% | 100% | 14.6% | 17.9% | Tail fiber pb3 |
| 87,183 | G→A | D741D (GAC→GAT) | 60.6% | 100% | 19.7% | 28.4% | Tail fiber pb3 |
| 87,195 | T→A | L737L (CTA→CTT) | 68.3% | 100% | 20.7% | 31.5% | Tail fiber pb3 |
| 87,204 | A→G | N734N (AAT→AAC) | 75.3% | 100% | 22.9% | 33.6% | Tail fiber pb3 |
| 87,216 | A→T | P730P (CCT→CCA) | 79.8% | 100% | 23.7% | 33.1% | Tail fiber pb3 |
| 87,219 | A→T | I729I (ATT→ATA) | 79.4% | 100% | 20.9% | 32.3% | Tail fiber pb3 |
| 87,223 | C→T | G728D (GGT→GAT) | 81.4% | 100% | 20.9% | 32.4% | Tail fiber pb3 |
| 87,243 | C→T | Q721Q (CAG→CAA) | 81.1% | 100% | 14.4% | 32.8% | Tail fiber pb3 |
| 87,246 | T→A | E720D (GAA→GAT) | 76.9% | 100% | 15.5% | 33.4% | Tail fiber pb3 |
| 87,249 | G→A | S719S (TCC→TCT) | 77.0% | 100% | 13.7% | 31.3% | Tail fiber pb3 |
| 87,252 | A→G | N718N (AAT→AAC) | 77.9% | 100% | 17.1% | 31.4% | Tail fiber pb3 |
| 87,255 | A→G | I717I (ATT→ATC) | 74.0% | 100% | 17.0% | 30.4% | Tail fiber pb3 |
| 87,273 | G→A | Y711Y (TAC→TAT) | 49.5% | 100% | | 7.7% | Tail fiber pb3 |
| 87,276 | C→T | E710E (GAG→GAA) | 51.2% | 100% | | 7.4% | Tail fiber pb3 |
| 87,279 | T→C | Q709Q (CAA→CAG) | 52.6% | 100% | | 7.9% | Tail fiber pb3 |
| 87,360 | T→G | E682D (GAA→GAC) | 24.5% | 79.3% | 7.1% | 10.8% | Tail fiber pb3 |
| 87,369 | G→A | F679F (TTC→TTT) | 36.1% | 100% | 8.4% | 14.0% | Tail fiber pb3 |
| 87,372 | A→T | A678A (GCT→GCA) | 36.6% | 77.5% | 8.5% | 13.9% | Tail fiber pb3 |
| 87,380 | G→C | P676A (CCT→GCT) | 34.9% | 85.1% | 8.2% | 12.5% | Tail fiber pb3 |
| 87,393 | T→A | I671I (ATA→ATT) | 40.5% | 100% | 8.4% | 13.6% | Tail fiber pb3 |
| 87,399 | T→A | I669I (ATA→ATT) | 38.4% | 81.6% | 7.1% | 13.2% | Tail fiber pb3 |
| 87,402 | A→G | F668F (TTT→TTC) | 38.6% | 100% | 9.0% | 13.1% | Tail fiber pb3 |
| 87,407 | T→G | K667Q (AAA→CAA) | 38.5% | 81.7% | 8.4% | 12.9% | Tail fiber pb3 |
| 87,416 | C→A | V664L (GTA→TTA) | 37.5% | 81.5% | 8.6% | 12.7% | Tail fiber pb3 |
| 87,420 | A→G | F662F (TTT→TTC) | 37.3% | 78.8% | 8.3% | 12.3% | Tail fiber pb3 |
| 87,423 | A→T | S661R (AGT→AGA) | 39.6% | 82.1% | 8.8% | 11.8% | Tail fiber pb3 |
| 87,424 | C→G | S661T (AGT→ACT) | 35.7% | 81.5% | 8.2% | 11.9% | Tail fiber pb3 |
| 87,425 | T→A | S661C (AGT→TGT) | 35.3% | 81.7% | 8.0% | 11.9% | Tail fiber pb3 |
| 87,426 | T→G | L660F (TTA→TTC) | 35.6% | 81.7% | 8.1% | 12.2% | Tail fiber pb3 |
| 87,428 | A→G | L660L (TTA→CTA) | 33.4% | | 7.4% | 12.8% | Tail fiber pb3 |
| 87,428 | 2 bp→GG | coding (1977-1978/2850 nt) | | 100% | | | Tail fiber pb3 |

| | | | | | | | |
|---|---|---|---|---|---|---|---|
| 87,429 | A→G | T659T (ACT→ACC) | 35.8% | | 7.9% | 12.0% | Tail fiber pb3 |
| 87,432 | A→T | R658R (CGT→CGA) | 33.3% | 81.5% | 9.3% | 12.0% | Tail fiber pb3 |
| 87,434 | G→T | R658S (CGT→AGT) | 34.0% | 81.5% | 8.5% | 12.1% | Tail fiber pb3 |
| 87,435 | C→A | S657S (TCG→TCT) | 34.1% | 81.5% | 8.5% | 12.1% | Tail fiber pb3 |
| 87,438 | G→A | Y656Y (TAC→TAT) | 32.9% | 79.1% | 7.8% | 11.7% | Tail fiber pb3 |
| 87,462 | A→G | D648D (GAT→GAC) | 62.2% | 100% | 27.5% | 33.6% | Tail fiber pb3 |
| 87,468 | G→A | Y646Y (TAC→TAT) | 61.5% | 89.8% | 26.5% | 33.4% | Tail fiber pb3 |
| 87,469 | T→A | Y646F (TAC→TTC) | 61.8% | 91.1% | 27.7% | 35.7% | Tail fiber pb3 |
| 87,474 | C→T | R644R (AGG→AGA) | 62.5% | 92.2% | 25.8% | 35.6% | Tail fiber pb3 |
| 87,483 | G→A | Y641Y (TAC→TAT) | 50.9% | 100% | 27.2% | 33.4% | Tail fiber pb3 |
| 87,489 | A→G | N639N (AAT→AAC) | 55.9% | 100% | 23.2% | 33.8% | Tail fiber pb3 |
| 87,495 | A→T | I637I (ATT→ATA) | 55.5% | 100% | 23.4% | 33.2% | Tail fiber pb3 |
| 87,501 | A→T | A635A (GCT→GCA) | 40.2% | 100% | 21.5% | 25.7% | Tail fiber pb3 |
| 87,513 | C→T | Q631Q (CAG→CAA) | 30.0% | 100% | 16.3% | 22.2% | Tail fiber pb3 |
| 87,516 | G→T | L630L (CTC→CTA) | 30.5% | 91.2% | 15.8% | 21.8% | Tail fiber pb3 |
| 87,518 | G→A | L630F (CTC→TTC) | 30.0% | 100% | 15.2% | 21.5% | Tail fiber pb3 |
| 87,522 | C→T | K628K (AAG→AAA) | | 100% | | | Tail fiber pb3 |
| 87,525 | G→A | D627D (GAC→GAT) | 27.5% | 100% | 17.1% | 22.7% | Tail fiber pb3 |
| 87,528 | A→T | L626L (CTT→CTA) | 28.0% | 91.5% | 17.1% | 22.9% | Tail fiber pb3 |
| 87,530 | G→A | L626F (CTT→TTT) | 29.6% | 100% | 17.0% | 21.8% | Tail fiber pb3 |
| 87,532 | C→T | G625D (GGT→GAT) | 29.5% | | 17.0% | 20.9% | Tail fiber pb3 |
| 87,532 | 2 bp→TT | coding (1873-1874/2850 nt) | | 100% | | | Tail fiber pb3 |
| 87,533 | C→T | G625S (GGT→AGT) | 29.3% | | 17.6% | 21.6% | Tail fiber pb3 |
| 87,540 | C→T | Q622Q (CAG→CAA) | 46.5% | 100% | 22.1% | 33.0% | Tail fiber pb3 |
| 87,550 | A→T | F619Y (TTT→TAT) | 47.4% | 100% | 27.0% | 36.0% | Tail fiber pb3 |
| 87,552 | C→T | K618K (AAG→AAA) | 49.3% | 100% | 26.2% | 35.8% | Tail fiber pb3 |
| 87,555 | C→A | S617S (TCG→TCT) | 51.9% | 100% | 28.9% | 35.1% | Tail fiber pb3 |
| 87,564 | G→A | F614F (TTC→TTT) | 60.3% | 100% | 32.8% | 39.5% | Tail fiber pb3 |
| 87,567 | C→G | T613T (ACG→ACC) | 60.2% | 100% | 32.7% | 40.3% | Tail fiber pb3 |
| 87,570 | G→A | I612I (ATC→ATT) | 61.4% | 100% | 31.7% | 40.7% | Tail fiber pb3 |
| 87,579 | T→A | T609T (ACA→ACT) | 74.4% | 100% | 33.4% | 45.2% | Tail fiber pb3 |
| 87,591 | T→A | L605L (CTA→CTT) | 83.6% | 100% | 35.4% | 43.6% | Tail fiber pb3 |
| 87,594 | T→A | A604A (GCA→GCT) | 82.2% | 100% | 36.3% | 43.9% | Tail fiber pb3 |
| 87,597 | A→T | P603P (CCT→CCA) | 79.6% | 100% | 36.2% | 43.1% | Tail fiber pb3 |
| 87,600 | A→G | D602D (GAT→GAC) | 80.9% | 100% | 35.1% | 44.0% | Tail fiber pb3 |
| 87,608 | A→T | L600I (TTA→ATA) | 83.3% | | 36.6% | 45.9% | Tail fiber pb3 |
| 87,608 | 2 bp→TG | coding (1797-1798/2850 nt) | | 100% | | | Tail fiber pb3 |
| 87,609 | C→G | S599S (TCG→TCC) | 83.1% | | 36.6% | 46.0% | Tail fiber pb3 |
| 87,618 | T→A | V596V (GTA→GTT) | 83.3% | 100% | 42.3% | 53.9% | Tail fiber pb3 |
| 87,627 | A→G | F593F (TTT→TTC) | 84.2% | 100% | 44.9% | 57.8% | Tail fiber pb3 |
| 87,651 | A→T | S585S (TCT→TCA) | 86.3% | 100% | 44.3% | 59.5% | Tail fiber pb3 |
| 87,656 | A→G | L584L (TTA→CTA) | 86.4% | | 46.6% | 61.7% | Tail fiber pb3 |
| 87,656 | 2 bp→GC | coding (1749-1750/2850 nt) | | 100% | | | Tail fiber pb3 |
| 87,657 | G→C | D583E (GAC→GAG) | 85.7% | | 45.8% | 60.4% | Tail fiber pb3 |
| 87,663 | G→A | D581D (GAC→GAT) | 86.1% | 100% | 44.3% | 60.2% | Tail fiber pb3 |
| 87,675 | G→A | D577D (GAC→GAT) | 83.7% | 100% | 39.8% | 51.7% | Tail fiber pb3 |
| 87,678 | T→C | L576L (CTA→CTG) | 83.8% | 100% | 39.9% | 50.9% | Tail fiber pb3 |
| 87,685 | C→T | G574D (GGT→GAT) | 71.4% | | 34.2% | 44.1% | Tail fiber pb3 |
| 87,685 | 3 bp→TTA | coding (1719-1721/2850 nt) | | 100% | | | Tail fiber pb3 |
| 87,686 | C→T | G574S (GGT→AGT) | 70.2% | | 34.2% | 44.6% | Tail fiber pb3 |
| 87,687 | T→A | I573I (ATA→ATT) | 70.0% | | 34.7% | 43.7% | Tail fiber pb3 |

| | | | | | | | |
|---|---|---|---|---|---|---|---|
| 87,690 | T→C | K572K (AAA→AAG) | 72.0% | 100% | 33.8% | 45.2% | Tail fiber pb3 |
| 87,692 | T→C | K572E (AAA→GAA) | 72.0% | 100% | 34.4% | 44.8% | Tail fiber pb3 |
| 87,702 | T→A | T568T (ACA→ACT) | 66.4% | | 36.4% | 45.3% | Tail fiber pb3 |
| 87,702 | 2 bp→AT | coding (1703-1704/2850 nt) | | 100% | | | Tail fiber pb3 |
| 87,703 | G→T | T568K (ACA→AAA) | 64.5% | | 36.3% | 45.1% | Tail fiber pb3 |
| 87,710 | G→A | H566Y (CAT→TAT) | 68.5% | 100% | 36.4% | 44.7% | Tail fiber pb3 |
| 87,720 | T→A | T562T (ACA→ACT) | 43.6% | 100% | 21.8% | 32.2% | Tail fiber pb3 |
| 87,723 | T→A | I561I (ATA→ATT) | 43.7% | 100% | 21.9% | 31.7% | Tail fiber pb3 |
| 87,726 | T→C | R560R (AGA→AGG) | 43.7% | 100% | 21.8% | 32.3% | Tail fiber pb3 |
| 87,732 | C→T | E558E (GAG→GAA) | 27.2% | 100% | 13.0% | 27.1% | Tail fiber pb3 |
| 87,734 | C→G | E558Q (GAG→CAG) | 27.6% | 100% | 12.7% | 27.7% | Tail fiber pb3 |
| 87,738 | T→A | S556S (TCA→TCT) | 28.6% | 100% | 12.4% | 28.2% | Tail fiber pb3 |
| 87,741 | C→T | L555L (CTG→CTA) | 28.3% | 100% | 14.0% | 26.9% | Tail fiber pb3 |
| 87,743 | G→A | L555L (CTG→TTG) | 28.1% | 100% | 10.6% | 27.6% | Tail fiber pb3 |
| 87,747 | A→G | N553N (AAT→AAC) | 30.8% | 100% | 14.6% | 29.1% | Tail fiber pb3 |
| 87,750 | T→A | I552I (ATA→ATT) | 34.3% | 100% | 15.9% | 29.7% | Tail fiber pb3 |
| 88,548 | G→A | N286N (AAC→AAT) | | 100% | | 11.9% | Tail fiber pb3 |
| 88,551 | A→G | N285H (AAT→CAC) | | 100% | | 11.2% | Tail fiber pb3 |
| 88,553 | T→G | N285H (AAT→CAC) | | 100% | 7.4% | 11.6% | Tail fiber pb3 |
| 88,557 | C→T | E283E (GAG→GAA) | | 100% | | 11.0% | Tail fiber pb3 |
| 88,560 | A→C | T282T (ACT→ACG) | | 100% | 9.2% | 10.8% | Tail fiber pb3 |
| 88,566 | T→C | A280A (GCA→GCG) | 24.9% | 100% | 18.7% | 31.3% | Tail fiber pb3 |
| 88,578 | G→T | I276I (ATC→ATA) | 36.9% | 100% | 33.0% | 43.0% | Tail fiber pb3 |
| 88,581 | A→T | G275G (GGT→GGA) | 37.9% | 100% | 31.5% | 43.0% | Tail fiber pb3 |
| 88,584 | A→T | P274P (CCT→CCA) | 37.2% | 100% | 31.4% | 41.1% | Tail fiber pb3 |
| 88,587 | A→T | I273I (ATT→ATA) | 38.8% | 100% | 32.2% | 40.3% | Tail fiber pb3 |
| 88,596 | A→T | V270V (GTT→GTA) | 54.1% | 100% | 42.2% | 51.5% | Tail fiber pb3 |
| 88,599 | T→A | G269G (GGA→GGT) | 56.0% | 100% | 41.3% | 52.8% | Tail fiber pb3 |
| 88,605 | T→A | V267V (GTA→GTT) | 53.1% | 100% | 43.6% | 52.9% | Tail fiber pb3 |
| 88,608 | A→T | V266V (GTT→GTA) | 53.2% | 100% | 44.0% | 53.6% | Tail fiber pb3 |
| 88,611 | A→T | P265P (CCT→CCA) | 52.8% | 100% | 42.5% | 53.8% | Tail fiber pb3 |
| 88,618 | A→T | F263Y (TTT→TAT) | 55.4% | 100% | 39.5% | 51.5% | Tail fiber pb3 |
| 88,620 | T→C | K262K (AAA→AAG) | 55.1% | 100% | 41.2% | 51.1% | Tail fiber pb3 |
| 88,629 | T→A | L259F (TTA→TTT) | 64.4% | 100% | 40.7% | 52.5% | Tail fiber pb3 |
| 88,631 | A→G | L259L (TTA→CTA) | 65.3% | 100% | 40.1% | 52.2% | Tail fiber pb3 |
| 88,638 | G→A | D256D (GAC→GAT) | 67.7% | 100% | 38.8% | 54.0% | Tail fiber pb3 |
| 88,643 | A→T | L255I (TTA→ATA) | 68.7% | 100% | 39.8% | 56.1% | Tail fiber pb3 |
| 88,647 | A→T | V253V (GTT→GTA) | 70.1% | 100% | 39.1% | 56.7% | Tail fiber pb3 |
| 88,659 | T→A | V249V (GTA→GTT) | 74.8% | 100% | 36.7% | 58.0% | Tail fiber pb3 |
| 88,686 | G→A | Y240Y (TAC→TAT) | 70.1% | 100% | 32.6% | 53.5% | Tail fiber pb3 |
| 88,695 | T→A | S237S (TCA→TCT) | 67.7% | 100% | 28.3% | 52.6% | Tail fiber pb3 |
| 88,701 | A→T | G235G (GGT→GGA) | 61.6% | 100% | 24.7% | 45.6% | Tail fiber pb3 |
| 88,707 | G→T | L233L (CTC→CTA) | 51.3% | 100% | 17.3% | 32.6% | Tail fiber pb3 |
| 88,710 | T→C | K232K (AAA→AAG) | 50.8% | 100% | 18.4% | 32.6% | Tail fiber pb3 |
| 88,713 | C→T | K231K (AAG→AAA) | | 100% | | | Tail fiber pb3 |
| 88,719 | G→A | S229S (TCC→TCT) | 38.3% | 100% | 12.8% | 29.2% | Tail fiber pb3 |
| 88,724 | 2 bp→TT | coding (681-682/2850 nt) | | 100% | | | Tail fiber pb3 |
| 88,728 | T→C | K226K (AAA→AAG) | | 100% | | | Tail fiber pb3 |
| 88,731 | A→G | Y225Y (TAT→TAC) | | 100% | | | Tail fiber pb3 |
| 88,734 | G→T | R224R (CGC→AGA) | | 100% | | | Tail fiber pb3 |
| 88,736 | G→T | R224R (CGC→AGA) | | 100% | | | Tail fiber pb3 |
| 88,743 | T→C | K221Q (AAA→CAG) | | 100% | | | Tail fiber pb3 |

| | | | | | | | |
|---|---|---|---|---|---|---|---|
| 88,745 | T→G | K221Q (AAA→CAG) | | 100% | | | Tail fiber pb3 |
| 88,749 | T→C | Q219Q (CAA→CAG) | | 100% | | | Tail fiber pb3 |
| 88,758 | A→T | A216A (GCT→GCA) | | 100% | | | Tail fiber pb3 |
| 88,763 | 2 bp→GA | coding (642-643/2850 nt) | | 100% | | | Tail fiber pb3 |
| 88,769 | T→A | T213S (ACT→TCT) | | 75.0% | | | Tail fiber pb3 |
| 88,771 | G→A | T212I (ACC→ATC) | | 69.6% | | | Tail fiber pb3 |
| 88,784 | A→C | S208A (TCT→GCT) | | 71.5% | | | Tail fiber pb3 |
| 88,785 | G→A | H207H (CAC→CAT) | | 71.6% | | | Tail fiber pb3 |
| 88,800 | G→A | D202D (GAC→GAT) | | 71.3% | | | Tail fiber pb3 |
| 89,867 | A→T | I50I (ATT→ATA) | 17.6% | ? | | | Tail fiber pb9 |
| 89,870 | T→A | G49G (GGA→GGT) | 17.5% | ? | | | Tail fiber pb9 |
| 89,890 | T→C | I43V (ATA→GTA) | 22.1% | ? | | 5.3% | Tail fiber pb9 |
| 89,891 | T→C | K42K (AAA→AAG) | 21.6% | ? | | | Tail fiber pb9 |
| 89,897 | C→T | E40E (GAG→GAA) | 22.0% | ? | | | Tail fiber pb9 |
| 89,900 | T→G | K39N (AAA→AAC) | 22.4% | ? | | 5.6% | Tail fiber pb9 |
| 89,903 | G→A | V38V (GTC→GTT) | 20.2% | ? | | 5.1% | Tail fiber pb9 |
| 89,915 | A→T | P34P (CCT→CCA) | 20.0% | ? | | 5.7% | Tail fiber pb9 |
| 89,990 | G→A | N9N (AAC→AAT) | 60.4% | 100% | 28.4% | 34.7% | Tail fiber pb9 |
| 90,002 | A→G | D5D (GAT→GAC) | 71.4% | 100% | 42.4% | 53.7% | Tail fiber pb9 |
| 90,010 | G→A | L3L (CTA→TTA) | 77.0% | 100% | 43.9% | 57.9% | Tail fiber pb9 |
| 90,058 | G→A | S17S (TCC→TCT) | 90.8% | 100% | 47.3% | 68.9% | hypothetical protein/Phage tail fiber |
| 90,107 | A→T | M1K (ATG→AAG) †‡ | 81.1% | 100% | 37.3% | 51.2% | hypothetical protein/Phage tail fiber |
| 90,108 | T→C | M1M (ATG→GTG) †‡ | 75.6% | 93.1% | 37.4% | 49.6% | hypothetical protein/Phage tail fiber |
| 90,123 | A→G | intergenic (-15/+3) | 74.2% | 100% | 31.6% | 42.0% | hypothetical protein/Phage tail fiber |
| 90,125 | A→T | intergenic (-17/+1) | 73.5% | 94.4% | 32.1% | 41.4% | hypothetical protein/Phage tail fiber |
| 90,302 | G→C | Q1178E (CAG→GAG) | 32.7% | 100% | 5.9% | 21.0% | Pore-forming tail protein pb2 |
| 90,336 | A→G | T1166T (ACT→ACC) | 79.5% | 100% | 35.3% | 52.6% | Pore-forming tail protein pb2 |
| 90,341 | T→C | I1165V (ATT→GTT) | 78.4% | 100% | 34.0% | 55.1% | Pore-forming tail protein pb2 |
| 90,345 | C→T | E1163E (GAG→GAA) | 77.5% | 100% | 34.7% | 55.1% | Pore-forming tail protein pb2 |
| 90,348 | A→T | T1162T (ACT→ACA) | 76.1% | 100% | 34.2% | 56.1% | Pore-forming tail protein pb2 |
| 90,354 | G→A | H1160H (CAC→CAT) | 72.2% | 100% | 34.5% | 56.8% | Pore-forming tail protein pb2 |
| 90,375 | T→A | V1153V (GTA→GTT) | 64.1% | 100% | 36.2% | 52.8% | Pore-forming tail protein pb2 |
| 90,378 | T→C | G1152G (GGA→GGG) | 62.5% | 100% | 39.1% | 54.5% | Pore-forming tail protein pb2 |
| 90,384 | A→G | Y1150Y (TAT→TAC) | 63.6% | 100% | 32.6% | 54.0% | Pore-forming tail protein pb2 |
| 90,39 | C→A | M1148I (ATG→ATT) | 58.0% | | 31.3% | 51.6% | Pore-forming tail protein pb2 |
| 90,39 | 2 bp→AT | coding (3443-3444/3708 nt) | | 100% | | | Pore-forming tail protein pb2 |
| 90,391 | A→T | M1148K (ATG→AAG) | 58.1% | | 31.0% | 52.3% | Pore-forming tail protein pb2 |
| 90,396 | G→A | G1146G (GGC→GGT) | 55.2% | 100% | 31.5% | 48.7% | Pore-forming tail protein pb2 |
| 90,399 | T→C | E1145E (GAA→GAG) | 56.8% | 100% | 32.2% | 49.6% | Pore-forming tail protein pb2 |
| 90,402 | C→A | A1144A (GCG→GCT) | 54.6% | 100% | 32.1% | 49.8% | Pore-forming tail protein pb2 |
| 90,408 | T→A | P1142P (CCA→CCT) | 56.2% | 100% | 33.0% | 50.6% | Pore-forming tail protein pb2 |
| 90,414 | A→G | F1140F (TTT→TTC) | 46.3% | 100% | 30.8% | 48.7% | Pore-forming tail protein pb2 |
| 90,417 | G→A | S1139S (TCC→TCT) | 46.8% | 100% | 29.9% | 47.0% | Pore-forming tail protein pb2 |
| 90,427 | T→C | N1136S (AAT→AGT) | 39.3% | | 20.9% | 37.5% | Pore-forming tail protein pb2 |
| 90,427 | 3 bp→CCG | coding (3405-3407/3708 nt) | | 100% | | | Pore-forming tail protein pb2 |
| 90,428 | T→C | N1136D (AAT→GAT) | 39.3% | | 20.9% | 37.8% | Pore-forming tail protein pb2 |
| 90,429 | A→G | G1135G (GGT→GGC) | 41.1% | | 20.6% | 36.8% | Pore-forming tail protein pb2 |
| 90,432 | G→T | I1134I (ATC→ATA) | 43.4% | 92.7% | 20.9% | 38.0% | Pore-forming tail protein pb2 |
| 90,435 | A→G | G1133G (GGT→GGC) | 43.6% | 100% | 20.5% | 38.1% | Pore-forming tail protein pb2 |
| 90,45 | T→G | L1128L (CTA→CTC) | 21.2% | 89.4% | 15.3% | 23.7% | Pore-forming tail protein pb2 |
| 90,452 | G→T | L1128I (CTA→ATA) | 17.0% | 85.8% | 14.0% | 21.3% | Pore-forming tail protein pb2 |

| | | | | | | | |
|---|---|---|---|---|---|---|---|
| 90,456 | G→A | S1126S (TCC→TCT) | 16.3% | 84.4% | 13.9% | 20.5% | Pore-forming tail protein pb2 |
| 90,537 | T→C | A1099A (GCA→GCG) | 48.7% | 100% | 19.2% | 40.2% | Pore-forming tail protein pb2 |
| 90,546 | G→A | D1096D (GAC→GAT) | 63.2% | 100% | 24.7% | 49.3% | Pore-forming tail protein pb2 |
| 90,552 | G→A | I1094I (ATC→ATT) | 63.9% | 100% | 24.2% | 47.7% | Pore-forming tail protein pb2 |
| 90,556 | G→C | S1093C (TCT→TGT) | 64.0% | 91.7% | 27.4% | 49.1% | Pore-forming tail protein pb2 |
| 90,557 | A→T | S1093T (TCT→ACT) | 64.4% | 92.3% | 27.1% | 49.2% | Pore-forming tail protein pb2 |
| 90,558 | A→T | S1092S (TCT→TCA) | 63.9% | 91.8% | 26.9% | 48.0% | Pore-forming tail protein pb2 |
| 90,57 | T→A | A1088A (GCA→GCT) | 74.3% | 100% | 43.0% | 60.9% | Pore-forming tail protein pb2 |
| 90,597 | C→G | A1079A (GCG→GCC) | 85.3% | 100% | 58.8% | 73.7% | Pore-forming tail protein pb2 |
| 90,624 | T→C | L1070L (CTA→CTG) | 91.9% | 100% | 63.4% | 77.8% | Pore-forming tail protein pb2 |
| 90,626 | G→A | L1070L (CTA→TTA) | 90.6% | 100% | 63.5% | 77.2% | Pore-forming tail protein pb2 |
| 90,645 | T→C | P1063P (CCA→CCG) | 90.3% | 100% | 64.7% | 79.2% | Pore-forming tail protein pb2 |
| 90,66 | T→C | A1058A (GCA→GCG) | 93.3% | 100% | 65.4% | 79.7% | Pore-forming tail protein pb2 |
| 90,678 | T→C | A1052A (GCA→GCG) | 93.5% | 100% | 65.5% | 78.8% | Pore-forming tail protein pb2 |
| 90,681 | T→A | T1051T (ACA→ACT) | 93.3% | 100% | 65.3% | 78.2% | Pore-forming tail protein pb2 |
| 90,684 | C→T | Q1050Q (CAG→CAA) | 91.2% | 100% | 62.3% | 75.8% | Pore-forming tail protein pb2 |
| 90,687 | A→G | I1049I (ATT→ATC) | 91.4% | 100% | 63.1% | 75.9% | Pore-forming tail protein pb2 |
| 90,693 | G→A | I1047I (ATC→ATT) | 91.2% | 100% | 64.5% | 77.0% | Pore-forming tail protein pb2 |
| 90,699 | T→C | K1045K (AAA→AAG) | 92.0% | 100% | 68.7% | 78.8% | Pore-forming tail protein pb2 |
| 90,711 | A→G | D1041D (GAT→GAC) | 92.5% | 100% | 67.1% | 77.7% | Pore-forming tail protein pb2 |
| 90,717 | C→T | Q1039Q (CAG→CAA) | 93.1% | 100% | 64.6% | 75.5% | Pore-forming tail protein pb2 |
| 90,723 | T→C | K1037K (AAA→AAG) | 92.7% | 100% | 69.5% | 75.9% | Pore-forming tail protein pb2 |
| 90,743 | A→G | L1031L (TTA→CTA) | 91.1% | 90.0% | 61.2% | 70.5% | Pore-forming tail protein pb2 |
| 90,744 | T→C | K1030K (AAA→AAG) | 92.8% | 91.2% | 61.9% | 70.8% | Pore-forming tail protein pb2 |
| 90,747 | T→C | K1029K (AAA→AAG) | 92.4% | 91.3% | 60.8% | 69.5% | Pore-forming tail protein pb2 |
| 90,752 | G→A | L1028L (CTG→TTG) | 93.6% | 90.8% | 60.6% | 70.1% | Pore-forming tail protein pb2 |
| 90,753 | C→T | K1027K (AAG→AAA) | 93.3% | 91.4% | 61.0% | 70.6% | Pore-forming tail protein pb2 |
| 90,768 | C→T | E1022E (GAG→GAA) | 91.2% | 90.4% | 55.0% | 64.9% | Pore-forming tail protein pb2 |
| 90,771 | A→T | S1021S (TCT→TCA) | 90.2% | 89.5% | 54.8% | 64.8% | Pore-forming tail protein pb2 |
| 90,777 | G→A | G1019G (GGC→GGT) | 89.9% | 88.5% | 51.5% | 63.0% | Pore-forming tail protein pb2 |
| 90,786 | C→T | K1016K (AAG→AAA) | 87.9% | 86.7% | 50.0% | 61.1% | Pore-forming tail protein pb2 |
| 90,792 | T→C | E1014E (GAA→GAG) | 84.9% | 85.1% | 52.6% | 59.0% | Pore-forming tail protein pb2 |
| 90,795 | T→C | A1013A (GCA→GCG) | 83.7% | 84.4% | 50.1% | 55.8% | Pore-forming tail protein pb2 |
| 90,801 | A→G | I1011I (ATT→ATC) | 82.4% | 81.8% | 48.3% | 54.0% | Pore-forming tail protein pb2 |
| 90,807 | Δ1 bp | coding (3027/3708 nt) | 75.8% | 75.0% | 40.9% | 44.4% | Pore-forming tail protein pb2 |
| 90,809:1 | +C | coding (3025/3708 nt) | 75.8% | 74.7% | 40.5% | 44.3% | Pore-forming tail protein pb2 |
| 90,816 | C→A | A1006A (GCG→GCT) | 79.8% | 75.8% | 41.1% | 44.2% | Pore-forming tail protein pb2 |
| 90,825 | T→C | Q1003Q (CAA→CAG) | 75.3% | 100% | 26.9% | 33.7% | Pore-forming tail protein pb2 |
| 90,831 | G→A | S1001S (AGC→AGT) | 71.9% | 100% | 25.9% | 32.6% | Pore-forming tail protein pb2 |
| 90,834 | G→A | T1000T (ACC→ACT) | 70.9% | 100% | 27.2% | 33.7% | Pore-forming tail protein pb2 |
| 90,843 | T→C | Q997Q (CAA→CAG) | 70.5% | 100% | 22.9% | 30.8% | Pore-forming tail protein pb2 |
| 90,855 | G→A | A993A (GCC→GCT) | 57.4% | 86.0% | 12.7% | 16.8% | Pore-forming tail protein pb2 |
| 90,857 | C→A | A993S (GCC→TCC) | 56.7% | 84.8% | 14.9% | 17.4% | Pore-forming tail protein pb2 |
| 90,864 | C→T | Q990Q (CAG→CAA) | 52.8% | 100% | 11.0% | 15.8% | Pore-forming tail protein pb2 |
| 90,87 | A→G | G988G (GGT→GGC) | 54.7% | 85.7% | 12.4% | 18.3% | Pore-forming tail protein pb2 |
| 90,884 | T→A | M984L (ATG→TTG) | 63.8% | 91.8% | 18.0% | 34.0% | Pore-forming tail protein pb2 |
| 90,888 | C→G | T982T (ACG→ACC) | 63.3% | 91.4% | 16.7% | 32.9% | Pore-forming tail protein pb2 |
| 90,899 | G→A | L979L (CTA→TTA) | 71.9% | 93.7% | 23.6% | 42.7% | Pore-forming tail protein pb2 |
| 90,9 | T→G | S978S (TCA→TCC) | 72.0% | 100% | 25.1% | 43.6% | Pore-forming tail protein pb2 |
| 90,909 | T→A | S975S (TCA→TCT) | 77.1% | 100% | 37.9% | 59.0% | Pore-forming tail protein pb2 |
| 90,915 | T→C | Q973Q (CAA→CAG) | 78.6% | 100% | 38.3% | 61.6% | Pore-forming tail protein pb2 |
| 90,926 | A→C | S970A (TCT→GCT) | 81.2% | 100% | 45.1% | 67.2% | Pore-forming tail protein pb2 |
| 90,936 | A→G | N966N (AAT→AAC) | 85.7% | 100% | 45.8% | 66.6% | Pore-forming tail protein pb2 |

| | | | | | | | |
|---|---|---|---|---|---|---|---|
| 90,945 | G→A | S963S (AGC→AGT) | 87.5% | 100% | 47.1% | 69.0% | Pore-forming tail protein pb2 |
| 90,953 | A→C | S961A (TCT→GCT) | 90.4% | | 53.2% | 72.3% | Pore-forming tail protein pb2 |
| 90,953 | 2 bp→CC | coding (2880-2881/3708 nt) | | 100% | | | Pore-forming tail protein pb2 |
| 90,954 | T→C | V960V (GTA→GTG) | 90.8% | | 52.2% | 72.8% | Pore-forming tail protein pb2 |
| 90,96 | A→G | T958T (ACT→ACC) | 91.1% | 100% | 56.2% | 75.9% | Pore-forming tail protein pb2 |
| 90,969 | G→A | S955S (TCC→TCT) | 91.5% | 100% | 61.2% | 77.4% | Pore-forming tail protein pb2 |
| 90,978 | C→T | E952E (GAG→GAA) | 89.8% | 100% | 62.8% | 79.2% | Pore-forming tail protein pb2 |
| 90,987 | T→C | K949K (AAA→AAG) | 93.0% | 100% | 67.0% | 82.2% | Pore-forming tail protein pb2 |
| 91,032 | A→T | A934A (GCT→GCA) | 90.7% | 100% | 63.0% | 75.7% | Pore-forming tail protein pb2 |
| 91,044 | A→G | D930D (GAT→GAC) | 91.6% | 100% | 60.7% | 72.8% | Pore-forming tail protein pb2 |
| 91,05 | T→C | G928G (GGA→GGG) | 92.3% | 100% | 60.0% | 71.3% | Pore-forming tail protein pb2 |
| 91,074 | A→G | Y920Y (TAT→TAC) | 81.4% | 100% | 53.6% | 63.3% | Pore-forming tail protein pb2 |
| 91,077 | T→C | V919V (GTA→GTG) | 81.3% | 100% | 53.7% | 63.7% | Pore-forming tail protein pb2 |
| 91,086 | T→C | V916V (GTA→GTG) | 76.3% | 100% | 43.9% | 55.8% | Pore-forming tail protein pb2 |
| 91,095 | C→T | G913G (GGG→GGA) | 74.9% | 100% | 39.7% | 49.4% | Pore-forming tail protein pb2 |
| 91,371 | G→T | A821A (GCC→GCA) | 33.9% | 58.4% | | | Pore-forming tail protein pb2 |
| 91,377 | A→C | T819T (ACT→ACG) | 33.7% | 58.4% | | | Pore-forming tail protein pb2 |
| 91,379 | T→C | T819A (ACT→GCT) | 37.5% | 54.1% | | | Pore-forming tail protein pb2 |
| 91,41 | A→G | D808D (GAT→GAC) | 75.1% | 100% | 27.8% | 42.1% | Pore-forming tail protein pb2 |
| 91,413 | A→C | A807A (GCT→GCG) | 74.1% | 100% | 28.7% | 42.2% | Pore-forming tail protein pb2 |
| 91,425 | G→A | Y803Y (TAC→TAT) | 86.1% | 100% | 35.1% | 50.3% | Pore-forming tail protein pb2 |
| 91,446 | T→C | E796E (GAA→GAG) | 82.3% | 100% | 34.5% | 53.0% | Pore-forming tail protein pb2 |
| 91,449 | A→T | V795V (GTT→GTA) | 79.9% | 100% | 34.8% | 52.7% | Pore-forming tail protein pb2 |
| 91,452 | G→A | T794T (ACC→ACT) | 82.9% | 100% | 34.1% | 53.4% | Pore-forming tail protein pb2 |
| 91,455 | C→T | L793L (TTG→TTA) | 79.4% | 100% | 32.8% | 54.0% | Pore-forming tail protein pb2 |
| 91,457 | A→G | L793L (TTG→CTG) | 80.7% | 100% | 33.7% | 54.5% | Pore-forming tail protein pb2 |
| 91,461 | A→T | L791L (CTT→CTA) | 75.4% | 100% | 32.5% | 54.2% | Pore-forming tail protein pb2 |
| 91,464 | A→T | N790K (AAT→AAA) | 76.9% | 100% | 32.8% | 52.7% | Pore-forming tail protein pb2 |
| 91,469 | A→G | L789L (TTA→CTA) | 76.2% | 100% | 34.1% | 53.3% | Pore-forming tail protein pb2 |
| 91,473 | T→A | A787A (GCA→GCT) | 79.4% | 100% | 34.0% | 52.8% | Pore-forming tail protein pb2 |
| 91,478 | G→A | L786L (CTA→TTA) | 77.0% | | 32.7% | 53.0% | Pore-forming tail protein pb2 |
| 91,478 | 2 bp→AT | coding (2355-2356/3708 nt) | | 100% | | | Pore-forming tail protein pb2 |
| 91,479 | A→T | R785R (CGT→CGA) | 79.5% | | 34.1% | 53.9% | Pore-forming tail protein pb2 |
| 91,481 | G→T | R785S (CGT→AGT) | 79.2% | 100% | 33.6% | 53.1% | Pore-forming tail protein pb2 |
| 91,485 | C→T | Q783Q (CAG→CAA) | 79.5% | 100% | 32.7% | 53.5% | Pore-forming tail protein pb2 |
| 91,492 | T→G | D781A (GAT→GCT) | 80.1% | | 32.0% | 54.8% | Pore-forming tail protein pb2 |
| 91,492 | 3 bp→GTA | coding (2340-2342/3708 nt) | | 100% | | | Pore-forming tail protein pb2 |
| 91,493 | C→T | D781N (GAT→AAT) | 77.9% | | 31.3% | 54.2% | Pore-forming tail protein pb2 |
| 91,494 | T→A | T780T (ACA→ACT) | 78.7% | | 30.9% | 55.1% | Pore-forming tail protein pb2 |
| 91,508 | T→C | N776D (AAC→GAC) | 86.9% | 100% | 46.9% | 65.3% | Pore-forming tail protein pb2 |
| 91,512 | T→C | L774L (TTA→TTG) | 88.7% | 100% | 48.3% | 66.8% | Pore-forming tail protein pb2 |
| 91,514 | A→G | L774L (TTA→CTA) | 87.6% | 100% | 52.0% | 66.9% | Pore-forming tail protein pb2 |
| 91,539 | C→A | L765L (CTG→CTT) | 94.3% | 100% | | 73.7% | Pore-forming tail protein pb2 |
| 91,542 | G→A | I764I (ATC→ATT) | 94.1% | 100% | | 72.7% | Pore-forming tail protein pb2 |
| 91,575 | A→G | A753A (GCT→GCC) | 100% | 100% | | 80.7% | Pore-forming tail protein pb2 |
| 91,6 | T→C | K745R (AAA→AGA) | 100% | 100% | | 83.9% | Pore-forming tail protein pb2 |
| 91,654 | C→T | R727Q (CGG→CAG) | 100% | 87.0% | | 86.5% | Pore-forming tail protein pb2 |
| 91,757 | T→G | K693Q (AAA→CAA) | | 84.8% | | 80.8% | Pore-forming tail protein pb2 |
| 91,757 | 2 bp→GG | coding (2076-2077/3708 nt) | 100% | | | | Pore-forming tail protein pb2 |
| 91,758 | A→G | N692N (AAT→AAC) | | 85.3% | | 81.0% | Pore-forming tail protein pb2 |
| 91,827 | G→A | Y669Y (TAC→TAT) | 100% | 80.4% | | 83.3% | Pore-forming tail protein pb2 |

| | | | | | | |
|---|---|---|---|---|---|---|
| 91,845 | G→A | S663S (TCC→TCT) | 100% | 79.0% | 78.1% | Pore-forming tail protein pb2 |
| 91,852 | A→G | I661T (ATT→ACT) | 100% | 77.8% | 76.7% | Pore-forming tail protein pb2 |
| 91,866 | T→C | L656L (CTA→CTG) | 100% | 69.4% | 69.3% | Pore-forming tail protein pb2 |
| 91,869 | G→T | T655T (ACC→ACA) | 100% | 68.9% | 69.8% | Pore-forming tail protein pb2 |
| 91,872 | A→G | N654N (AAT→AAC) | 100% | 69.2% | 71.0% | Pore-forming tail protein pb2 |
| 91,876 | A→T | F653Y (TTT→TAT) | 100% | 66.6% | 69.0% | Pore-forming tail protein pb2 |
| 91,878 | C→G | G652G (GGG→GGC) | 100% | 67.1% | 68.4% | Pore-forming tail protein pb2 |
| 91,881 | T→C | L651M (TTA→CTG) | 100% | 67.4% | 67.9% | Pore-forming tail protein pb2 |
| 91,883 | A→G | L651M (TTA→CTG) | 100% | 67.6% | 67.6% | Pore-forming tail protein pb2 |
| 91,89 | A→G | N648N (AAT→AAC) | 100% | 66.6% | 67.5% | Pore-forming tail protein pb2 |
| 91,902 | T→C | E644E (GAA→GAG) | 100% | 66.0% | 64.9% | Pore-forming tail protein pb2 |
| 91,923 | T→G | G637G (GGA→GGC) | 100% | 52.0% | 44.9% | Pore-forming tail protein pb2 |
| 91,927 | G→A | A636V (GCA→GTA) | | 42.8% | 31.3% | Pore-forming tail protein pb2 |
| 91,927 | 2 bp→AA | coding (1906-1907/3708 nt) | 100% | | | Pore-forming tail protein pb2 |
| 91,928 | C→A | A636S (GCA→TCA) | | 42.9% | 31.6% | Pore-forming tail protein pb2 |
| 91,932 | G→T | L634L (CTC→CTA) | 100% | 44.6% | 31.9% | Pore-forming tail protein pb2 |
| 91,943 | C→T | V631I (GTA→ATA) | ? | 16.6% | 10.5% | Pore-forming tail protein pb2 |
| 91,944 | A→G | N630N (AAT→AAC) | ? | 17.7% | 10.4% | Pore-forming tail protein pb2 |
| 91,95 | A→T | I628I (ATT→ATA) | ? | 18.0% | 11.2% | Pore-forming tail protein pb2 |
| 91,953 | T→A | V627V (GTA→GTT) | ? | 19.4% | 10.7% | Pore-forming tail protein pb2 |
| 91,954 | A→G | V627A (GTA→GCA) | ? | 18.7% | 11.1% | Pore-forming tail protein pb2 |
| 91,956 | C→A | G626G (GGG→GGT) | ? | 19.3% | 10.9% | Pore-forming tail protein pb2 |
| 91,957 | C→T | G626E (GGG→GAG) | ? | 18.8% | 10.7% | Pore-forming tail protein pb2 |
| 91,96 | T→G | E625A (GAA→GCA) | ? | 19.3% | 11.0% | Pore-forming tail protein pb2 |
| 91,962 | A→T | F624L (TTT→TTA) | ? | 21.2% | 11.0% | Pore-forming tail protein pb2 |
| 91,967 | C→T | A623T (GCT→ACT) | ? | 19.2% | 11.5% | Pore-forming tail protein pb2 |
| 91,968 | T→A | A622A (GCA→GCT) | ? | 19.8% | 11.3% | Pore-forming tail protein pb2 |
| 91,974 | G→A | T620T (ACC→ACT) | ? | 19.6% | 12.7% | Pore-forming tail protein pb2 |
| 92,049 | C→T | Q595Q (CAG→CAA) | 100% | 71.6% | 63.7% | Pore-forming tail protein pb2 |
| 92,064 | T→C | Q590Q (CAA→CAG) | 100% | 79.5% | 72.1% | Pore-forming tail protein pb2 |
| 92,091 | A→T | T581T (ACT→ACA) | 100% | 81.9% | 73.0% | Pore-forming tail protein pb2 |
| 92,094 | A→G | S580S (TCT→TCC) | 100% | 82.0% | 73.6% | Pore-forming tail protein pb2 |
| 92,103 | A→G | D577D (GAT→GAC) | 100% | 83.9% | 75.2% | Pore-forming tail protein pb2 |
| 92,106 | G→A | Y576Y (TAC→TAT) | 100% | 82.7% | 75.1% | Pore-forming tail protein pb2 |
| 92,116 | T→C | D573G (GAT→GGT) | 100% | 83.7% | 77.3% | Pore-forming tail protein pb2 |
| 92,133 | C→T | R567R (AGG→AGA) | 100% | 84.2% | 79.6% | Pore-forming tail protein pb2 |
| 92,136 | G→C | H566Q (CAC→CAG) | 100% | 84.1% | 80.2% | Pore-forming tail protein pb2 |
| 92,154 | T→C | Q560Q (CAA→CAG) | 100% | 82.8% | 77.8% | Pore-forming tail protein pb2 |
| 92,157 | G→A | F559F (TTC→TTT) | 100% | 81.3% | 76.0% | Pore-forming tail protein pb2 |
| 92,16 | T→C | E558E (GAA→GAG) | 100% | 81.6% | 76.7% | Pore-forming tail protein pb2 |
| 92,184 | G→A | A550A (GCC→GCT) | 100% | 78.7% | 75.8% | Pore-forming tail protein pb2 |
| 92,235 | G→A | D533D (GAC→GAT) | 100% | 64.7% | 63.0% | Pore-forming tail protein pb2 |
| 92,238 | A→T | D532E (GAT→GAA) | 100% | 65.0% | 61.8% | Pore-forming tail protein pb2 |
| 92,241 | A→T | V531V (GTT→GTA) | 100% | 65.0% | 62.4% | Pore-forming tail protein pb2 |
| 92,247 | T→C | K529K (AAA→AAG) | 100% | 60.5% | 58.1% | Pore-forming tail protein pb2 |
| 92,25 | G→A | S528G (AGC→GGT) | 100% | 58.5% | 56.1% | Pore-forming tail protein pb2 |
| 92,252 | T→C | S528G (AGC→GGT) | 100% | 59.7% | 56.4% | Pore-forming tail protein pb2 |
| 92,256 | C→T | R526R (AGG→AGA) | 100% | 57.5% | 57.0% | Pore-forming tail protein pb2 |
| 92,259 | T→G | A525T (GCA→ACC) | 100% | 58.1% | 56.9% | Pore-forming tail protein pb2 |
| 92,261 | C→T | A525T (GCA→ACC) | 100% | 57.2% | 56.7% | Pore-forming tail protein pb2 |
| 92,264 | T→C | I524V (ATA→GTA) | | 58.7% | 58.0% | Pore-forming tail protein pb2 |
| 92,264 | 2 bp→CT | coding (1569-1570/3708 nt) | 100% | | | Pore-forming tail protein pb2 |

| | | | | | | |
|---|---|---|---|---|---|---|
| 92,265 | C→T | Q523Q (CAG→CAA) | | 58.1% | 58.2% | Pore-forming tail protein pb2 |
| 92,268 | C→T | L522L (CTG→CTA) | 100% | 58.7% | 59.2% | Pore-forming tail protein pb2 |
| 92,271 | C→A | G521G (GGG→GGT) | 100% | 59.0% | 58.9% | Pore-forming tail protein pb2 |
| 92,28 | T→C | A518A (GCA→GCG) | 100% | 60.3% | 60.5% | Pore-forming tail protein pb2 |
| 92,283 | G→A | T517T (ACC→ACT) | 100% | 60.7% | 59.0% | Pore-forming tail protein pb2 |
| 92,289 | C→T | E515E (GAG→GAA) | 100% | 61.3% | 58.6% | Pore-forming tail protein pb2 |
| 92,318 | A→C | S506A (TCT→GCT) | 61.6% | 34.1% | 35.2% | Pore-forming tail protein pb2 |
| 92,466 | A→C | L456L (CTT→CTG) | 100% | 54.8% | 61.8% | Pore-forming tail protein pb2 |
| 92,469 | G→A | S455S (TCC→TCT) | 100% | 55.4% | 62.3% | Pore-forming tail protein pb2 |
| 92,472 | T→C | E454E (GAA→GAG) | 100% | 56.5% | 62.6% | Pore-forming tail protein pb2 |
| 92,475 | C→T | R453R (AGG→AGA) | | 55.0% | 62.5% | Pore-forming tail protein pb2 |
| 92,475 | 2 bp→TT | coding (1358-1359/3708 nt) | 100% | | | Pore-forming tail protein pb2 |
| 92,476 | C→T | R453K (AGG→AAG) | | 54.5% | 62.7% | Pore-forming tail protein pb2 |
| 92,487 | T→C | L449L (CTA→CTG) | 100% | 65.9% | | Pore-forming tail protein pb2 |
| 92,502 | C→T | G444G (GGG→GGA) | 100% | 72.3% | | Pore-forming tail protein pb2 |
| 92,52 | C→T | L438L (TTG→TTA) | 100% | 73.9% | | Pore-forming tail protein pb2 |
| 92,553 | A→G | D427D (GAT→GAC) | 100% | 78.8% | | Pore-forming tail protein pb2 |
| 92,563 | T→C | K424R (AAA→AGA) | 100% | 79.0% | | Pore-forming tail protein pb2 |
| 92,573 | G→T | Q421K (CAA→AAA) | 100% | | | Pore-forming tail protein pb2 |
| 92,603 | C→T | V411I (GTA→ATA) | 100% | 77.5% | | Pore-forming tail protein pb2 |
| 92,616 | A→T | T406T (ACT→ACA) | 100% | 77.9% | | Pore-forming tail protein pb2 |
| 92,652 | A→C | G394G (GGT→GGG) | 100% | 76.7% | | Pore-forming tail protein pb2 |
| 92,661 | C→T | K391K (AAG→AAA) | 100% | 75.3% | | Pore-forming tail protein pb2 |
| 92,673 | T→C | Q387Q (CAA→CAG) | 100% | 70.0% | | Pore-forming tail protein pb2 |
| 92,676 | T→C | Q386Q (CAA→CAG) | 100% | 71.1% | | Pore-forming tail protein pb2 |
| 92,679 | T→A | V385V (GTA→GTT) | 100% | 70.4% | | Pore-forming tail protein pb2 |
| 92,680:1 | +T | coding (1154/3708 nt) | 100% | 67.3% | | Pore-forming tail protein pb2 |
| 92,683 | Δ1 bp | coding (1151/3708 nt) | 100% | 67.9% | | Pore-forming tail protein pb2 |
| 92,688 | T→C | L382L (CTA→CTG) | 100% | 72.0% | | Pore-forming tail protein pb2 |
| 92,703 | G→A | N377N (AAC→AAT) | 100% | 70.1% | | Pore-forming tail protein pb2 |
| 92,715 | C→T | Q373Q (CAG→CAA) | 93.1% | 63.9% | | Pore-forming tail protein pb2 |
| 92,724 | C→T | P370P (CCG→CCA) | 88.3% | 58.7% | | Pore-forming tail protein pb2 |
| 92,73 | A→T | A368A (GCT→GCA) | 88.3% | 52.9% | | Pore-forming tail protein pb2 |
| 92,738 | A→G | L366L (TTA→CTA) | 82.4% | 43.2% | | Pore-forming tail protein pb2 |
| 92,739 | A→T | V365V (GTT→GTA) | 81.6% | 42.8% | | Pore-forming tail protein pb2 |
| 92,75 | A→G | L362L (TTA→CTA) | 79.3% | 31.4% | | Pore-forming tail protein pb2 |
| 92,754 | A→G | G360G (GGT→GGC) | 75.8% | 28.0% | | Pore-forming tail protein pb2 |
| 92,763 | T→C | K357K (AAA→AAG) | 59.7% | 13.2% | | Pore-forming tail protein pb2 |
| 92,766 | G→A | T356T (ACC→ACT) | 58.6% | 10.3% | | Pore-forming tail protein pb2 |
| 92,775 | T→A | A353A (GCA→GCT) | 52.6% | | | Pore-forming tail protein pb2 |
| 92,778 | T→A | I352I (ATA→ATT) | 47.3% | | | Pore-forming tail protein pb2 |
| 92,781 | A→T | V351V (GTT→GTA) | 47.5% | | | Pore-forming tail protein pb2 |
| 92,784 | A→C | A350A (GCT→GCG) | 47.4% | | | Pore-forming tail protein pb2 |
| 92,817 | A→G | T339T (ACT→ACC) | 70.0% | 14.0% | | Pore-forming tail protein pb2 |
| 92,831 | T→C | I335V (ATT→GTT) | | 27.5% | | Pore-forming tail protein pb2 |
| 92,831 | 2 bp→CA | coding (1002-1003/3708 nt) | 100% | | | Pore-forming tail protein pb2 |
| 92,832 | G→A | N334N (AAC→AAT) | | 27.4% | | Pore-forming tail protein pb2 |
| 92,838 | A→T | T332T (ACT→ACA) | 100% | 37.1% | | Pore-forming tail protein pb2 |
| 92,847 | A→T | T329T (ACT→ACA) | 100% | 45.1% | | Pore-forming tail protein pb2 |
| 92,853 | A→T | A327A (GCT→GCA) | 100% | 52.9% | | Pore-forming tail protein pb2 |
| 92,864 | G→A | L324L (CTA→TTA) | 100% | 59.6% | | Pore-forming tail protein pb2 |
| 92,877 | G→A | D319D (GAC→GAT) | 100% | 63.1% | | Pore-forming tail protein pb2 |

| | | | | | | |
|---|---|---|---|---|---|---|
| 92,883 | A→G | Y317Y (TAT→TAC) | 100% | 68.9% | | Pore-forming tail protein pb2 |
| 92,889 | G→A | D315D (GAC→GAT) | 100% | 69.8% | | Pore-forming tail protein pb2 |
| 92,901 | A→G | I311I (ATT→ATC) | 100% | 72.5% | | Pore-forming tail protein pb2 |
| 92,904 | A→G | T310T (ACT→ACC) | 100% | 73.0% | | Pore-forming tail protein pb2 |
| 92,907 | A→G | V309V (GTT→GTC) | 100% | 73.4% | | Pore-forming tail protein pb2 |
| 92,913 | A→C | L307L (CTT→CTG) | 100% | 76.9% | | Pore-forming tail protein pb2 |
| 92,916 | T→C | E306E (GAA→GAG) | 100% | 76.0% | | Pore-forming tail protein pb2 |
| 92,924 | A→G | L304L (TTG→CTG) | 100% | 77.9% | | Pore-forming tail protein pb2 |
| 92,943 | T→A | S297S (TCA→TCT) | 100% | 79.0% | | Pore-forming tail protein pb2 |
| 92,949 | A→G | G295G (GGT→GGC) | 100% | 79.4% | | Pore-forming tail protein pb2 |
| 92,988 | A→G | G282G (GGT→GGC) | 100% | 77.9% | | Pore-forming tail protein pb2 |
| 93,003 | A→C | A277A (GCT→GCG) | 100% | 75.0% | | Pore-forming tail protein pb2 |
| 93,02 | G→A | L272L (CTA→TTA) | 100% | 74.1% | | Pore-forming tail protein pb2 |
| 93,042 | A→G | A264A (GCT→GCC) | 100% | 73.1% | | Pore-forming tail protein pb2 |
| 93,06 | G→A | S258S (TCC→TCT) | 100% | 68.7% | | Pore-forming tail protein pb2 |
| 93,063 | G→T | A257A (GCC→GCA) | 100% | 68.8% | | Pore-forming tail protein pb2 |
| 93,066 | A→T | S256S (TCT→TCA) | 100% | 69.9% | | Pore-forming tail protein pb2 |
| 93,069 | G→A | S255S (TCC→TCT) | 100% | 69.5% | | Pore-forming tail protein pb2 |
| 93,075 | T→C | Q253Q (CAA→CAG) | 100% | 69.0% | | Pore-forming tail protein pb2 |
| 93,078 | A→T | R252R (CGT→AGA) | 100% | 68.4% | | Pore-forming tail protein pb2 |
| 93,08 | G→T | R252R (CGT→AGA) | 100% | 68.3% | | Pore-forming tail protein pb2 |
| 93,084 | A→T | A250A (GCT→GCA) | 100% | 70.2% | | Pore-forming tail protein pb2 |
| 93,09 | T→C | E248E (GAA→GAG) | 100% | 71.0% | | Pore-forming tail protein pb2 |
| 93,099 | G→A | I245I (ATC→ATT) | 100% | 65.5% | | Pore-forming tail protein pb2 |
| 93,102 | T→A | A244A (GCA→GCT) | 100% | 66.5% | | Pore-forming tail protein pb2 |
| 93,108 | G→T | G242G (GGC→GGA) | 100% | 65.7% | | Pore-forming tail protein pb2 |
| 93,114 | T→A | A240A (GCA→GCT) | 100% | 65.4% | | Pore-forming tail protein pb2 |
| 93,117 | C→T | E239E (GAG→GAA) | 100% | 65.7% | | Pore-forming tail protein pb2 |
| 93,14 | A→T | S232T (TCC→ACC) | 100% | 51.6% | | Pore-forming tail protein pb2 |
| 93,26 | A→G | F192L (TTT→CTT) | 100% | 24.6% | | Pore-forming tail protein pb2 |
| 93,264 | A→T | A190A (GCT→GCA) | 100% | 25.0% | | Pore-forming tail protein pb2 |
| 93,27 | Δ1 bp | coding (564/3708 nt) | | 26.1% | | Pore-forming tail protein pb2 |
| 93,27 | 2 bp→C | coding (563-564/3708 n t) | 100% | | | Pore-forming tail protein pb2 |
| 93,271 | A→C | L188* (TTA→TGA) | | 26.6% | | Pore-forming tail protein pb2 |
| 93,276:1 | +A | coding (558/3708 nt) | 100% | 26.0% | | Pore-forming tail protein pb2 |
| 93,281 | A→G | L185L (TTA→CTA) | 100% | 26.3% | | Pore-forming tail protein pb2 |
| 93,284 | C→T | G184S (GGT→AGT) | 100% | 25.7% | | Pore-forming tail protein pb2 |
| 93,293 | C→T | V181I (GTT→ATT) | | 35.6% | | Pore-forming tail protein pb2 |
| 93,293 | 2 bp→TC | coding (540-541/3708 n t) | 100% | | | Pore-forming tail protein pb2 |
| 93,294 | T→C | K180K (AAA→AAG) | | 36.9% | | Pore-forming tail protein pb2 |
| 93,306 | A→C | A176A (GCT→GCG) | 100% | 52.8% | | Pore-forming tail protein pb2 |
| 93,33 | A→T | A168A (GCT→GCA) | 100% | 63.4% | | Pore-forming tail protein pb2 |
| 93,404 | T→C | T144A (ACT→GCT) | 100% | 80.7% | | Pore-forming tail protein pb2 |
| 93,411 | G→A | G141G (GGC→GGT) | 100% | 79.3% | | Pore-forming tail protein pb2 |
| 93,417 | A→T | V139V (GTT→GTA) | | 79.2% | 71.6% | Pore-forming tail protein pb2 |
| 93,417 | 2 bp→TG | coding (416-417/3708 n t) | 100% | | | Pore-forming tail protein pb2 |
| 93,418 | A→G | V139A (GTT→GCT) | | 80.8% | 71.9% | Pore-forming tail protein pb2 |
| 93,426 | G→A | T136T (ACC→ACT) | 100% | 79.8% | 70.9% | Pore-forming tail protein pb2 |
| 93,429 | G→A | D135D (GAC→GAT) | 100% | 79.9% | 70.6% | Pore-forming tail protein pb2 |
| 93,435 | C→T | L133L (TTG→TTA) | 100% | 82.5% | 70.0% | Pore-forming tail protein pb2 |
| 93,444 | T→C | Q130Q (CAA→CAG) | 100% | 82.9% | 68.7% | Pore-forming tail protein pb2 |

| | | | | | | |
|---|---|---|---|---|---|---|
| 93,447 | A→G | V129V (GTT→GTC) | 100% | 82.6% | 65.5% | Pore-forming tail protein pb2 |
| 93,45 | C→T | K128K (AAG→AAA) | 100% | 82.1% | 65.0% | Pore-forming tail protein pb2 |
| 93,453 | C→T | E127E (GAG→GAA) | 100% | 83.1% | 65.5% | Pore-forming tail protein pb2 |
| 93,485 | C→T | A117T (GCA→ACA) | | 79.7% | 68.1% | Pore-forming tail protein pb2 |
| 93,485 | 2 bp→TT | coding (348-349/3708 nt) | 100% | | | Pore-forming tail protein pb2 |
| 93,486 | C→T | K116K (AAG→AAA) | | 80.2% | 68.5% | Pore-forming tail protein pb2 |
| 93,517 | G→A | A106V (GCT→GTT) | | 79.4% | 68.6% | Pore-forming tail protein pb2 |
| 93,517 | 2 bp→AT | coding (316-317/3708 nt) | 100% | | | Pore-forming tail protein pb2 |
| 93,518 | C→T | A106T (GCT→ACT) | | 79.2% | 68.0% | Pore-forming tail protein pb2 |
| 93,528 | G→A | D102D (GAC→GAT) | 100% | 80.0% | 66.6% | Pore-forming tail protein pb2 |
| 93,546 | T→C | Q96Q (CAA→CAG) | 100% | 79.1% | 70.7% | Pore-forming tail protein pb2 |
| 93,555 | C→T | L93L (CTG→CTA) | 100% | 78.6% | 73.4% | Pore-forming tail protein pb2 |
| 93,567 | C→T | K89K (AAG→AAA) | 100% | 77.0% | 71.1% | Pore-forming tail protein pb2 |
| 93,58 | C→T | G85D (GGT→GAT) | 100% | 78.4% | 71.8% | Pore-forming tail protein pb2 |
| 93,584 | C→T | D84N (GAT→AAT) | 100% | 78.6% | 69.9% | Pore-forming tail protein pb2 |
| 93,594 | A→G | T80T (ACT→ACC) | 100% | 77.9% | 68.3% | Pore-forming tail protein pb2 |
| 93,602 | T→C | S78G (AGT→GGT) | 100% | 75.7% | 61.3% | Pore-forming tail protein pb2 |
| 93,604 | A→G | L77P (CTT→CCT) | | 28.6% | 61.6% | Pore-forming tail protein pb2 |
| 93,604 | 2 bp→GC | coding (229-230/3708 nt) | 100% | | | Pore-forming tail protein pb2 |
| 93,605 | G→C | L77V (CTT→GTT) | | 75.7% | 60.9% | Pore-forming tail protein pb2 |
| 93,609 | G→A | G75G (GGC→GGT) | 100% | 75.7% | 61.6% | Pore-forming tail protein pb2 |
| 93,627 | A→T | N69K (AAT→AAA) | 100% | | | Pore-forming tail protein pb2 |
| 93,63 | T→A | T68T (ACA→ACT) | 100% | 74.4% | 61.6% | Pore-forming tail protein pb2 |
| 93,651 | A→G | D61D (GAT→GAC) | 100% | 78.8% | 67.2% | Pore-forming tail protein pb2 |
| 93,672 | G→A | Y54Y (TAC→TAT) | 100% | 77.2% | 66.2% | Pore-forming tail protein pb2 |
| 93,675 | T→A | L53L (CTA→CTT) | 100% | 77.5% | 66.6% | Pore-forming tail protein pb2 |
| 93,695 | T→C | S47G (AGT→GGT) | 100% | 78.8% | 66.6% | Pore-forming tail protein pb2 |
| 93,698 | G→A | L46L (CTA→TTA) | 100% | 77.8% | 65.7% | Pore-forming tail protein pb2 |
| 93,717 | A→G | S39S (TCT→TCC) | 100% | 81.6% | 69.1% | Pore-forming tail protein pb2 |
| 93,726 | G→A | A36A (GCC→GCT) | 100% | 82.9% | 71.4% | Pore-forming tail protein pb2 |
| 93,732 | G→A | N34N (AAC→AAT) | 100% | 82.6% | 71.5% | Pore-forming tail protein pb2 |
| 93,74 | G→A | L32L (CTA→TTA) | 100% | 82.0% | 74.1% | Pore-forming tail protein pb2 |
| 93,798 | G→A | D12D (GAC→GAT) | 100% | 84.4% | 74.1% | Pore-forming tail protein pb2 |
| 93,806 | G→A | L10L (CTA→TTA) | 100% | 83.9% | 72.8% | Pore-forming tail protein pb2 |
| 93,835 | T→A | intergenic (-2/+83) | 100% | 84.9% | 71.6% | Phage tail fiber |
| 93,858 | Δ1 bp | intergenic (-25/+60) | 91.2% | 77.7% | 67.3% | Phage tail fiber |
| 93,868 | T→A | intergenic (-35/+50) | 100% | 85.4% | 75.7% | Phage tail fiber |
| 93,903 | Δ1 bp | intergenic (-70/+15) | 100% | 100% | | Phage tail fiber |
| 93,921 | T→C | R122R (CGA→CGG) | 100% | 82.6% | 75.6% | Tail fiber p138 |
| 93,927 | G→T | R120R (CGC→CGA) | 100% | 82.8% | 75.0% | Tail fiber p138 |
| 93,934 | C→T | R118K (AGA→AAA) | 100% | 79.8% | 69.6% | Tail fiber p138 |
| 93,936 | A→G | A117A (GCT→GCC) | 100% | 80.3% | 70.7% | Tail fiber p138 |
| 93,957 | A→T | I110I (ATT→ATA) | 100% | 79.7% | 68.4% | Tail fiber p138 |
| 93,96 | C→T | R109R (AGG→AGA) | 100% | 81.3% | 69.0% | Tail fiber p138 |
| 93,963 | T→A | G108G (GGA→GGT) | 100% | 80.5% | 68.1% | Tail fiber p138 |
| 94,005 | A→G | D94D (GAT→GAC) | 100% | 83.9% | 70.8% | Tail fiber p138 |
| 94,056 | T→C | E77E (GAA→GAG) | 100% | 85.2% | 67.6% | Tail fiber p138 |
| 94,068 | T→G | T73T (ACA→ACC) | 100% | 83.5% | 67.5% | Tail fiber p138 |
| 94,086 | A→G | F67F (TTT→TTC) | 100% | 83.6% | 67.1% | Tail fiber p138 |
| 94,098 | C→T | L63L (TTG→TTA) | 100% | 82.1% | 69.6% | Tail fiber p138 |
| 94,107 | C→T | K60K (AAG→AAA) | 100% | 81.1% | 67.4% | Tail fiber p138 |

| | | | | | | |
|---|---|---|---|---|---|---|
| 94,131 | G→A | F52F (TTC→TTT) | 100% | 81.9% | 69.8% | Tail fiber p138 |
| 94,191 | A→G | I32I (ATT→ATC) | 100% | 78.9% | 71.8% | Tail fiber p138 |
| 94,203 | G→A | D28D (GAC→GAT) | 100% | 79.2% | 71.6% | Tail fiber p138 |
| 94,233 | G→A | D18D (GAC→GAT) | 100% | 78.5% | 69.3% | Tail fiber p138 |
| 94,236 | G→A | P17P (CCC→CCT) | 100% | 78.7% | 71.3% | Tail fiber p138 |
| 94,239 | C→T | E16E (GAG→GAA) | 100% | 77.9% | 68.1% | Tail fiber p138 |
| 94,269 | G→A | Y6Y (TAC→TAT) | 100% | 79.9% | 75.1% | Tail fiber p138 |
| 94,317 | T→G | intergenic (-31/+31) | 100% | 80.0% | 71.8% | Phage protein |
| 94,441 | G→A | N104N (AAC→AAT) | 100% | 77.2% | 67.5% | Tail fiber p139 |
| 94,471 | A→G | P94P (CCT→CCC) | 100% | 77.5% | 66.2% | Tail fiber p139 |
| 94,489 | A→G | I88I (ATT→ATC) | 100% | 78.6% | 62.3% | Tail fiber p139 |
| 94,492 | C→T | L87L (CTG→CTA) | 100% | 78.2% | 60.7% | Tail fiber p139 |
| 94,519 | G→A | T78T (ACC→ACT) | 100% | 80.3% | 61.9% | Tail fiber p139 |
| 94,543 | A→C | A70A (GCT→GCG) | 100% | 79.9% | 54.7% | Tail fiber p139 |
| 94,558 | T→A | A65A (GCA→GCT) | 100% | 76.4% | 49.9% | Tail fiber p139 |
| 94,567 | G→A | F62F (TTC→TTT) | 100% | 73.9% | 50.5% | Tail fiber p139 |
| 94,576 | A→G | D59D (GAT→GAC) | 100% | 74.8% | 47.6% | Tail fiber p139 |
| 94,588 | C→T | Q55Q (CAG→CAA) | 100% | 70.3% | 43.2% | Tail fiber p139 |
| 94,594 | C→T | L53L (TTG→CTA) | 100% | 66.7% | 37.1% | Tail fiber p139 |
| 94,596 | A→G | L53L (TTG→CTA) | 100% | 66.7% | 39.0% | Tail fiber p139 |
| 94,603 | A→G | N50N (AAT→AAC) | 100% | 66.2% | 36.7% | Tail fiber p139 |
| 94,612 | C→T | E47E (GAG→GAA) | 100% | 62.2% | 37.9% | Tail fiber p139 |
| 94,627 | T→A | S42S (TCA→TCT) | 100% | 58.2% | 32.2% | Tail fiber p139 |
| 94,635 | T→C | I40V (ATC→GTC) | 100% | 58.3% | 42.1% | Tail fiber p139 |
| 94,642 | C→T | K37K (AAG→AAA) | 100% | 52.5% | 41.4% | Tail fiber p139 |
| 94,645 | C→A | S36S (TCG→TCT) | 100% | 51.9% | 40.2% | Tail fiber p139 |
| 94,648 | G→A | T35T (ACC→ACT) | 100% | 52.9% | 44.0% | Tail fiber p139 |
| 94,663 | A→G | Y30Y (TAT→TAC) | 100% | 52.4% | 43.0% | Tail fiber p139 |
| 94,677 | A→G | L26L (TTA→CTA) | 100% | 48.4% | 40.7% | Tail fiber p139 |
| 94,687 | A→T | P22P (CCT→CCA) | 100% | 46.7% | 38.5% | Tail fiber p139 |
| 94,702 | T→G | S17S (TCA→TCC) | 100% | 47.2% | 41.2% | Tail fiber p139 |
| 94,717 | A→G | R12R (CGT→CGC) | 100% | 50.4% | 40.7% | Tail fiber p139 |
| 94,72 | G→A | T11T (ACC→ACT) | 100% | 49.4% | 40.7% | Tail fiber p139 |
| 94,726 | G→A | L9L (CTC→CTT) | 100% | 51.3% | 40.5% | Tail fiber p139 |
| 94,754 | Δ1 bp | coding (895/897 nt) | 89.1% | 45.2% | 33.8% | Tail fiber p139 |
| 94,758 | T→A | R297S (AGA→AGT) | 93.4% | 47.7% | 32.4% | Tail fiber p140 |
| 94,759 | C→A | R297I (AGA→ATA) | 92.8% | 47.5% | 32.3% | Tail fiber p140 |
| 94,773 | G→A | N292N (AAC→AAT) | 78.6% | 29.7% | 13.4% | Tail fiber p140 |
| 94,775 | T→C | N292D (AAC→GAC) | 79.0% | 30.7% | 13.6% | Tail fiber p140 |
| 94,782:1 | +C | coding (867/897 nt) | 71.4% | 28.2% | 13.7% | Tail fiber p140 |
| 94,785 | A→T | P288P (CCT→CCA) | 78.7% | 29.7% | 15.7% | Tail fiber p140 |
| 94,788 | A→G | D287D (GAT→GAC) | | 30.7% | 16.1% | Tail fiber p140 |
| 94,795 | T→G | D285A (GAT→GCT) | 78.7% | 30.9% | 15.2% | Tail fiber p140 |
| 94,796 | C→A | D285Y (GAT→TAT) | 78.6% | 33.0% | 16.4% | Tail fiber p140 |
| 94,806 | A→T | I281I (ATT→ATA) | 78.6% | 31.5% | 14.9% | Tail fiber p140 |
| 94,809 | T→A | V280I (GTA→ATT) | 100% | 31.7% | 14.7% | Tail fiber p140 |
| 94,811 | C→T | V280I (GTA→ATT) | 100% | 32.1% | 14.8% | Tail fiber p140 |
| 94,815 | A→C | Y278* (TAT→TAG) | | 33.5% | 15.0% | Tail fiber p140 |
| 94,815 | 2 bp→CC | coding (833-834/897 nt) | 100% | | | Tail fiber p140 |
| 94,816 | T→C | Y278C (TAT→TGT) | | 33.1% | 15.0% | Tail fiber p140 |
| 94,821 | T→A | V276V (GTA→GTT) | 100% | 31.0% | 15.3% | Tail fiber p140 |
| 94,824 | C→T | Q275K (CAG→AAA) | 100% | 29.9% | 14.3% | Tail fiber p140 |
| 94,826 | G→T | Q275K (CAG→AAA) | 100% | 30.7% | 14.3% | Tail fiber p140 |

| | | | | | | |
|---|---|---|---|---|---|---|
| 94,832 | A→C | L273V (TTG→GTG) | 100% | 30.1% | 13.0% | Tail fiber p140 |
| 94,842 | T→A | K269N (AAA→AAT) | 85.3% | 29.8% | 13.0% | Tail fiber p140 |
| 94,844 | T→C | K269E (AAA→GAA) | 100% | 29.7% | 12.7% | Tail fiber p140 |
| 94,856 | A→T | S265T (TCA→ACA) | 82.7% | 22.4% | 10.5% | Tail fiber p140 |
| 94,986 | G→C | S221S (TCC→TCG) | 88.3% | | | Tail fiber p140 |
| 95,004 | G→T | V215V (GTC→GTA) | 91.9% | 5.9% | | Tail fiber p140 |
| 95,015 | G→T | R212R (CGA→AGA) | 100% | 5.4% | | Tail fiber p140 |
| 95,023 | G→T | T209N (ACT→AAT) | 92.3% | 5.8% | | Tail fiber p140 |
| 95,03 | C→T | V207I (GTT→ATT) | 100% | 5.4% | | Tail fiber p140 |
| 95,046 | T→A | I201V (ATA→GTT) | 100% | | | Tail fiber p140 |
| 95,048 | T→C | I201V (ATA→GTT) | 100% | | | Tail fiber p140 |
| 95,052 | 2 bp→CT | coding (596-597/897 nt) | 100% | | | Tail fiber p140 |
| 95,07 | A→G | C193C (TGT→TGC) | 100% | | | Tail fiber p140 |
| 95,073 | T→C | Q192Q (CAA→CAG) | 100% | | | Tail fiber p140 |
| 95,076 | T→C | Q191Q (CAA→CAG) | 100% | | | Tail fiber p140 |
| 95,205 | A→G | S148S (TCT→TCC) | 100% | | | Tail fiber p140 |
| 95,214 | T→C | G145G (GGA→GGG) | 100% | 18.6% | 11.0% | Tail fiber p140 |
| 95,217 | A→T | S144S (TCT→TCA) | 100% | 18.4% | 12.8% | Tail fiber p140 |
| 95,235 | T→A | L138L (CTA→CTT) | 100% | 36.0% | 18.8% | Tail fiber p140 |
| 95,238 | G→A | I137I (ATC→ATT) | 100% | 36.0% | 19.8% | Tail fiber p140 |
| 95,241 | C→T | P136P (CCG→CCA) | 100% | 36.4% | 18.8% | Tail fiber p140 |
| 95,248 | C→T | S134N (AGT→AAT) | 100% | 36.1% | 23.3% | Tail fiber p140 |
| 95,258 | G→A | L131L (CTA→TTA) | 100% | 37.0% | 22.2% | Tail fiber p140 |
| 95,268 | A→T | V127V (GTT→GTA) | 100% | 36.7% | 23.0% | Tail fiber p140 |
| 95,274 | T→A | S125S (TCA→TCT) | 100% | 38.1% | 21.7% | Tail fiber p140 |
| 95,283 | G→A | C122C (TGC→TGT) | 100% | 37.9% | 16.9% | Tail fiber p140 |
| 95,292 | G→A | F119F (TTC→TTT) | 100% | 34.7% | 15.1% | Tail fiber p140 |
| 95,298 | A→T | I117V (ATT→GTA) | 100% | 36.0% | 16.1% | Tail fiber p140 |
| 95,3 | T→C | I117V (ATT→GTA) | 100% | 36.4% | 16.0% | Tail fiber p140 |
| 95,305 | C→T | S115N (AGC→AAC) | 100% | 36.4% | 15.0% | Tail fiber p140 |
| 95,307 | A→G | N114N (AAT→AAC) | 100% | 38.4% | 16.9% | Tail fiber p140 |
| 95,312 | T→C | N113D (AAT→GAT) | | 37.8% | 17.2% | Tail fiber p140 |
| 95,312 | 2 bp→CT | coding (336-337/897 nt) | 100% | | | Tail fiber p140 |
| 95,313 | A→T | H112Q (CAT→CAA) | | 38.1% | 16.4% | Tail fiber p140 |
| 95,315 | G→T | H112N (CAT→AAT) | 100% | 38.4% | 16.8% | Tail fiber p140 |
| 95,319 | G→T | I110I (ATC→ATA) | 100% | 38.2% | 16.7% | Tail fiber p140 |
| 95,321 | T→C | I110V (ATC→GTC) | | 40.0% | 17.1% | Tail fiber p140 |
| 95,321 | 2 bp→CT | coding (327-328/897 nt) | 100% | | | Tail fiber p140 |
| 95,322 | A→T | I109I (ATT→ATA) | | 38.2% | 16.1% | Tail fiber p140 |
| 95,328 | C→G | M107I (ATG→ATC) | 100% | 38.3% | 15.2% | Tail fiber p140 |
| 95,334 | G→A | F105F (TTC→TTT) | 100% | 37.8% | 16.2% | Tail fiber p140 |
| 95,423 | A→C | S76A (TCT→GCT) | 63.7% | | | Tail fiber p140 |
| 95,433 | C→T | T72T (ACG→ACA) | 100% | | | Tail fiber p140 |
| 95,437 | G→T | T71N (ACT→AAT) | 100% | | | Tail fiber p140 |
| 95,442 | G→A | F69F (TTC→TTT) | 100% | | | Tail fiber p140 |
| 95,454 | C→T | L65L (TTG→TTA) | 100% | 6.2% | | Tail fiber p140 |
| 95,46 | A→T | I63I (ATT→ATA) | 100% | 5.8% | | Tail fiber p140 |
| 95,466 | T→A | S61S (TCA→TCT) | 100% | 6.5% | | Tail fiber p140 |
| 95,473 | T→G | D59A (GAT→GCT) | | 5.6% | | Tail fiber p140 |
| 95,473 | 2 bp→GT | coding (175-176/897 nt) | 100% | | | Tail fiber p140 |
| 95,484 | A→T | I55I (ATT→ATA) | 100% | 6.0% | | Tail fiber p140 |
| 95,571 | A→T | S26S (TCT→TCA) | 55.9% | | | Tail fiber p140 |
| 95,574 | G→T | L25L (CTC→CTA) | 55.5% | | | Tail fiber p140 |

| | | | | | | |
|---|---|---|---|---|---|---|
| 95,583 | G→A | F22F (TTC→TTT) | 61.7% | | | Tail fiber p140 |
| 95,59 | T→A | Y20F (TAT→TTT) | 61.8% | | | Tail fiber p140 |
| 95,592 | C→T | G19G (GGG→GGA) | 68.4% | | | Tail fiber p140 |
| 95,593 | C→G | G19A (GGG→GCG) | 68.4% | | | Tail fiber p140 |
| 95,613 | A→T | V12V (GTT→GTA) | 84.1% | | | Tail fiber p140 |
| 95,615 | C→T | V12I (GTT→ATT) | 78.4% | | | Tail fiber p140 |
| 95,616 | T→A | I11I (ATA→ATT) | 77.7% | | | Tail fiber p140 |
| 95,618 | T→C | I11V (ATA→GTA) | 100% | | | Tail fiber p140 |
| 95,622 | A→T | S9S (TCT→TCA) | 100% | | | Tail fiber p140 |
| 95,634 | G→T | L5L (CTC→CTA) | 71.5% | | | Tail fiber p140 |
| 95,644 | A→T | F2Y (TTT→TAT) | 71.5% | | | Tail fiber p140 |
| 96,001 | C→T | E348E (GAG→GAA) | | 6.7% | | Phage major tail protein pb6 |
| 96,007 | T→A | G346G (GGA→GGT) | | 7.1% | | Phage major tail protein pb6 |
| 96,012 | A→T | S345T (TCT→ACT) | 79.1% | 7.2% | | Phage major tail protein pb6 |
| 96,018 | G→A | L343L (CTG→TTG) | | 7.4% | | Phage major tail protein pb6 |
| 96,019 | C→A | E342D (GAG→GAT) | 79.0% | 8.7% | | Phage major tail protein pb6 |
| 96,024 | A→T | S341T (TCA→ACA) | 80.0% | 8.1% | | Phage major tail protein pb6 |
| 96,025 | T→A | P340P (CCA→CCT) | 75.6% | 8.5% | | Phage major tail protein pb6 |
| 96,04 | T→C | E335E (GAA→GAG) | 100% | 13.6% | 8.0% | Phage major tail protein pb6 |
| 96,055 | T→A | L330L (CTA→CTT) | 100% | 26.8% | 14.0% | Phage major tail protein pb6 |
| 96,058 | C→T | V329V (GTG→GTA) | 100% | 25.9% | 15.0% | Phage major tail protein pb6 |
| 96,067 | C→A | T326T (ACG→ACT) | 100% | 32.9% | 19.5% | Phage major tail protein pb6 |
| 96,086 | C→T | S320N (AGC→AAC) | 100% | 36.0% | 18.3% | Phage major tail protein pb6 |
| 96,091 | A→G | H318H (CAT→CAC) | 100% | 38.4% | 14.9% | Phage major tail protein pb6 |
| 96,094 | T→C | A317A (GCA→GCG) | 100% | 38.6% | 14.0% | Phage major tail protein pb6 |
| 96,1 | C→T | K315K (AAG→AAA) | 100% | 34.9% | 12.8% | Phage major tail protein pb6 |
| 96,106 | T→A | V313V (GTA→GTT) | 86.3% | 33.2% | 9.2% | Phage major tail protein pb6 |
| 96,111 | G→A | L312L (CTA→TTA) | 80.7% | 32.1% | 7.7% | Phage major tail protein pb6 |
| 96,112 | A→C | V311V (GTT→GTG) | 80.8% | 32.9% | 9.1% | Phage major tail protein pb6 |
| 96,118 | G→T | A309A (GCC→GCA) | 80.7% | 31.2% | 8.0% | Phage major tail protein pb6 |
| 96,121 | A→C | P308P (CCT→CCG) | 80.8% | 31.1% | 8.3% | Phage major tail protein pb6 |
| 96,124 | A→T | R307R (CGT→CGA) | 82.8% | 30.8% | 8.0% | Phage major tail protein pb6 |
| 96,127 | C→T | G306G (GGG→GGA) | 80.8% | 31.2% | 8.2% | Phage major tail protein pb6 |
| 96,128 | C→T | G306E (GGG→GAG) | 80.8% | 31.4% | 7.8% | Phage major tail protein pb6 |
| 96,13 | C→A | E305D (GAG→GAT) | 80.7% | 31.8% | 8.4% | Phage major tail protein pb6 |
| 96,134 | G→T | T304N (ACT→AAT) | 80.7% | 31.2% | 8.1% | Phage major tail protein pb6 |
| 96,135 | T→C | T304A (ACT→GCT) | 81.8% | 31.7% | 9.3% | Phage major tail protein pb6 |
| 96,142 | T→A | G301G (GGA→GGT) | 78.1% | 30.1% | 8.6% | Phage major tail protein pb6 |
| 96,145 | C→A | G300G (GGG→GGT) | 77.5% | 29.8% | 8.0% | Phage major tail protein pb6 |
| 96,148 | T→C | L299L (CTA→CTG) | 79.9% | 29.2% | 9.1% | Phage major tail protein pb6 |
| 96,15 | G→A | L299L (CTA→TTA) | 78.1% | 28.0% | 8.2% | Phage major tail protein pb6 |
| 96,151 | T→A | I298I (ATA→ATT) | 77.9% | 29.8% | 10.1% | Phage major tail protein pb6 |
| 96,154 | T→A | L297L (CTA→CTT) | 100% | 27.5% | 8.8% | Phage major tail protein pb6 |
| 96,157 | G→T | A296A (GCC→GCA) | 86.3% | 28.7% | 8.6% | Phage major tail protein pb6 |
| 96,163 | C→T | E294E (GAG→GAA) | 100% | 50.2% | 20.1% | Phage major tail protein pb6 |
| 96,187 | A→G | T286T (ACT→ACC) | 100% | 61.3% | 32.6% | Phage major tail protein pb6 |
| 96,205 | A→G | Y280Y (TAT→TAC) | 100% | 58.1% | 35.1% | Phage major tail protein pb6 |
| 96,208 | T→C | L279L (CTA→CTG) | 100% | 58.4% | 34.8% | Phage major tail protein pb6 |
| 96,211 | C→T | E278E (GAG→GAA) | 100% | 57.0% | 33.6% | Phage major tail protein pb6 |
| 96,22 | G→A | G275G (GGC→GGT) | 100% | 60.1% | 35.7% | Phage major tail protein pb6 |
| 96,228 | A→C | S273A (TCT→GCT) | 100% | 60.4% | 34.9% | Phage major tail protein pb6 |
| 96,235 | A→G | N270N (AAT→AAC) | 100% | 61.5% | 35.1% | Phage major tail protein pb6 |
| 96,238 | C→A | L269F (TTG→TTT) | 94.8% | 60.6% | 33.8% | Phage major tail protein pb6 |

| | | | | | | |
|---|---|---|---|---|---|---|
| 96,24 | A→G | L269L (TTG→CTG) | 100% | 59.9% | 34.3% | Phage major tail protein pb6 |
| 96,241 | A→G | Y268Y (TAT→TAC) | 93.5% | 59.9% | 36.2% | Phage major tail protein pb6 |
| 96,244 | A→T | A267A (GCT→GCA) | 94.5% | 60.5% | 35.2% | Phage major tail protein pb6 |
| 96,247 | A→T | T266T (ACT→ACA) | 94.9% | 60.7% | 34.8% | Phage major tail protein pb6 |
| 96,252 | A→G | L265L (TTA→CTA) | | 60.2% | 32.9% | Phage major tail protein pb6 |
| 96,252 | 2 bp→GA | coding (792-793/1392 nt) | 100% | | | Phage major tail protein pb6 |
| 96,253 | G→A | S264S (TCC→TCT) | | 59.4% | 32.2% | Phage major tail protein pb6 |
| 96,259 | T→A | T262T (ACA→ACT) | 94.6% | 65.1% | 41.0% | Phage major tail protein pb6 |
| 96,289 | A→G | I252I (ATT→ATC) | 100% | 69.4% | 48.9% | Phage major tail protein pb6 |
| 96,301 | A→T | V248V (GTT→GTA) | 100% | 62.7% | 44.7% | Phage major tail protein pb6 |
| 96,307 | G→T | S246S (TCC→TCA) | 100% | 59.2% | 44.1% | Phage major tail protein pb6 |
| 96,313 | T→A | V244V (GTA→GTT) | 100% | 49.1% | 39.1% | Phage major tail protein pb6 |
| 96,315 | C→T | V244I (GTA→ATA) | | 48.5% | 38.9% | Phage major tail protein pb6 |
| 96,315 | 2 bp→TA | coding (729-730/1392 nt) | 100% | | | Phage major tail protein pb6 |
| 96,316 | G→A | N243N (AAC→AAT) | | 47.9% | 39.2% | Phage major tail protein pb6 |
| 96,319 | A→C | P242P (CCT→CCG) | 100% | 48.9% | 39.7% | Phage major tail protein pb6 |
| 96,322 | T→G | T241T (ACA→ACC) | 100% | 51.7% | 41.6% | Phage major tail protein pb6 |
| 96,327 | A→G | L240L (TTA→CTA) | 100% | 49.8% | 41.5% | Phage major tail protein pb6 |
| 96,331 | G→C | T238T (ACC→ACG) | 100% | 49.6% | 40.2% | Phage major tail protein pb6 |
| 96,334 | A→G | I237I (ATT→ATC) | 100% | 48.7% | 41.3% | Phage major tail protein pb6 |
| 96,337 | A→G | N236N (AAT→AAC) | 100% | 51.4% | 41.6% | Phage major tail protein pb6 |
| 96,34 | A→G | N235N (AAT→AAC) | 100% | 50.5% | 42.7% | Phage major tail protein pb6 |
| 96,364 | C→A | T227T (ACG→ACT) | 100% | 59.9% | 45.6% | Phage major tail protein pb6 |
| 96,37 | T→A | P225P (CCA→CCT) | 100% | 58.5% | 39.3% | Phage major tail protein pb6 |
| 96,373 | T→A | I224I (ATA→ATT) | 100% | 57.6% | 38.9% | Phage major tail protein pb6 |
| 96,382 | A→T | S221A (TCT→GCA) | 100% | 53.6% | 38.5% | Phage major tail protein pb6 |
| 96,384 | A→C | S221A (TCT→GCA) | 100% | 53.4% | 38.1% | Phage major tail protein pb6 |
| 96,39 | G→C | H219D (CAT→GAT) | | 43.2% | 27.0% | Phage major tail protein pb6 |
| 96,39 | 2 bp→CA | coding (654-655/1392 nt) | 100% | | | Phage major tail protein pb6 |
| 96,391 | G→A | T218T (ACC→ACT) | | 42.9% | 26.7% | Phage major tail protein pb6 |
| 96,393 | T→A | T218S (ACC→TCC) | 100% | 43.0% | 26.0% | Phage major tail protein pb6 |
| 96,4 | G→A | N215D (AAC→GAT) | 100% | 41.3% | 29.6% | Phage major tail protein pb6 |
| 96,402 | T→C | N215D (AAC→GAT) | 100% | 42.4% | 31.8% | Phage major tail protein pb6 |
| 96,409 | C→T | K212K (AAG→AAA) | 100% | 40.5% | 29.1% | Phage major tail protein pb6 |
| 96,412 | C→T | L211L (TTG→CTA) | 100% | 40.4% | 26.5% | Phage major tail protein pb6 |
| 96,414 | A→G | L211L (TTG→CTA) | 100% | 40.4% | 26.8% | Phage major tail protein pb6 |
| 96,418 | C→A | T209T (ACG→ACT) | 100% | 40.7% | 27.3% | Phage major tail protein pb6 |
| 96,423 | G→A | L208L (CTA→TTA) | 90.2% | 39.4% | 26.0% | Phage major tail protein pb6 |
| 96,424 | T→C | K207K (AAA→AAG) | 100% | 40.9% | 28.6% | Phage major tail protein pb6 |
| 96,427 | A→G | N206N (AAT→AAC) | 100% | 39.0% | 28.0% | Phage major tail protein pb6 |
| 96,433 | G→A | I204I (ATC→ATT) | 100% | 48.9% | 33.5% | Phage major tail protein pb6 |
| 96,436 | A→G | Y203Y (TAT→TAC) | 100% | 51.3% | 35.0% | Phage major tail protein pb6 |
| 96,448 | G→A | L199L (CTC→CTT) | 100% | 59.3% | 39.5% | Phage major tail protein pb6 |
| 96,45 | G→T | L199I (CTC→ATC) | 94.6% | 58.2% | 40.0% | Phage major tail protein pb6 |
| 96,46 | A→G | T195T (ACT→ACC) | 93.0% | 62.5% | 47.6% | Phage major tail protein pb6 |
| 96,48 | T→A | I189L (ATA→TTA) | 93.5% | 58.8% | 46.1% | Phage major tail protein pb6 |
| 96,482 | T→G | E188A (GAA→GCA) | 94.2% | 59.4% | 45.3% | Phage major tail protein pb6 |
| 96,487 | C→T | P186P (CCG→CCA) | 94.4% | 57.7% | 44.8% | Phage major tail protein pb6 |
| 96,49 | G→A | D185D (GAC→GAT) | 94.1% | 57.6% | 44.3% | Phage major tail protein pb6 |
| 96,499 | C→T | Q182Q (CAG→CAA) | 100% | 54.4% | 39.2% | Phage major tail protein pb6 |
| 96,502 | T→G | R181S (AGA→AGC) | 100% | 55.2% | 39.9% | Phage major tail protein pb6 |

| | | | | | | |
|---|---|---|---|---|---|---|
| 96,508 | A→T | L179L (CTT→CTA) | 100% | 51.2% | 41.3% | Phage major tail protein pb6 |
| 96,51 | G→A | L179F (CTT→TTT) | | 51.5% | 40.0% | Phage major tail protein pb6 |
| 96,51 | 2 bp→AC | coding (534-535/1392 nt) | 100% | | | Phage major tail protein pb6 |
| 96,511 | A→C | P178P (CCT→CCG) | | 52.1% | 40.5% | Phage major tail protein pb6 |
| 96,517 | T→G | L176L (CTA→CTC) | 100% | 51.0% | 40.5% | Phage major tail protein pb6 |
| 96,523 | G→A | N174N (AAC→AAT) | 77.9% | 50.0% | 41.8% | Phage major tail protein pb6 |
| 96,541 | G→A | T168T (ACC→ACT) | 75.7% | 48.0% | 40.7% | Phage major tail protein pb6 |
| 96,544 | C→T | V167V (GTG→GTA) | 74.9% | 47.5% | 38.6% | Phage major tail protein pb6 |
| 96,553 | T→A | I164I (ATA→ATT) | 71.0% | 44.6% | 32.4% | Phage major tail protein pb6 |
| 96,562 | G→A | I161I (ATC→ATT) | 65.6% | 41.0% | 28.4% | Phage major tail protein pb6 |
| 96,574 | T→A | V157V (GTA→GTT) | 62.1% | 33.3% | 19.3% | Phage major tail protein pb6 |
| 96,586 | A→G | N153N (AAT→AAC) | 49.3% | 22.0% | 12.4% | Phage major tail protein pb6 |
| 96,589 | A→G | I152I (ATT→ATC) | 48.4% | 21.1% | 12.7% | Phage major tail protein pb6 |
| 96,598 | G→A | S149S (TCC→TCT) | 37.9% | 13.1% | | Phage major tail protein pb6 |
| 96,601 | A→G | D148D (GAT→GAC) | 38.6% | 13.6% | | Phage major tail protein pb6 |
| 96,607 | A→G | Y146Y (TAT→TAC) | 28.4% | | | Phage major tail protein pb6 |
| 96,61 | G→A | S145S (AGC→AGT) | 21.2% | | | Phage major tail protein pb6 |
| 96,619 | C→A | K142N (AAG→AAT) | 6.4% | | | Phage major tail protein pb6 |
| 96,637 | T→G | I136I (ATA→ATC) | 7.7% | | | Phage major tail protein pb6 |
| 96,664 | G→A | A127A (GCC→GCT) | 22.1% | 11.4% | | Phage major tail protein pb6 |
| 96,666 | C→A | A127S (GCC→TCC) | 22.6% | 11.3% | | Phage major tail protein pb6 |
| 96,667 | A→G | N126N (AAT→AAC) | 22.6% | 11.8% | | Phage major tail protein pb6 |
| 96,679 | A→G | N122N (AAT→AAC) | 43.3% | 34.4% | 26.7% | Phage major tail protein pb6 |
| 96,688 | A→G | F119F (TTT→TTC) | 51.7% | 40.3% | 32.0% | Phage major tail protein pb6 |
| 96,696 | C→T | A117T (GCT→ACT) | 55.7% | 47.7% | 33.7% | Phage major tail protein pb6 |
| 96,712 | G→T | G111G (GGC→GGA) | 62.2% | 59.1% | 44.0% | Phage major tail protein pb6 |
| 96,721 | A→T | G108G (GGT→GGA) | 68.4% | 60.5% | 46.5% | Phage major tail protein pb6 |
| 96,724 | T→A | A107A (GCA→GCT) | 68.7% | 60.7% | 44.6% | Phage major tail protein pb6 |
| 96,725 | G→T | A107E (GCA→GAA) | 69.3% | 59.8% | 44.5% | Phage major tail protein pb6 |
| 96,73 | G→A | N105N (AAC→AAT) | 74.9% | 64.9% | 52.1% | Phage major tail protein pb6 |
| 96,736 | G→A | P103P (CCC→CCT) | 74.3% | 64.3% | 54.3% | Phage major tail protein pb6 |
| 96,738 | G→C | P103A (CCC→GCC) | 74.6% | 63.7% | 54.5% | Phage major tail protein pb6 |
| 96,742 | G→A | G101G (GGC→GGT) | 71.4% | 62.7% | 56.0% | Phage major tail protein pb6 |
| 96,769 | G→A | Y92Y (TAC→TAT) | 80.2% | 76.2% | 67.8% | Phage major tail protein pb6 |
| 96,775 | G→A | P90P (CCC→CCT) | 79.8% | 75.9% | 67.0% | Phage major tail protein pb6 |
| 96,784 | C→T | Q87Q (CAG→CAA) | 78.1% | 75.5% | 64.2% | Phage major tail protein pb6 |
| 96,79 | A→G | N85N (AAT→AAC) | 78.2% | 75.8% | 63.9% | Phage major tail protein pb6 |
| 96,791 | T→C | N85S (AAT→AGT) | 77.9% | 76.1% | 64.5% | Phage major tail protein pb6 |
| 96,795 | A→T | S84T (TCT→ACT) | 77.5% | 74.6% | 63.2% | Phage major tail protein pb6 |
| 96,796 | G→A | T83T (ACC→ACT) | 77.8% | 75.0% | 64.6% | Phage major tail protein pb6 |
| 96,817 | A→G | I76I (ATT→ATC) | 77.5% | 74.4% | 66.4% | Phage major tail protein pb6 |
| 96,823 | A→T | T74T (ACT→ACA) | 75.8% | 72.0% | 61.1% | Phage major tail protein pb6 |
| 96,826 | A→G | S73S (TCT→TCC) | 75.8% | 72.5% | 60.7% | Phage major tail protein pb6 |
| 96,829 | A→G | F72F (TTT→TTC) | 74.8% | 72.4% | 60.5% | Phage major tail protein pb6 |
| 96,832 | A→G | S71S (AGT→AGC) | 75.1% | 73.5% | 61.8% | Phage major tail protein pb6 |
| 96,847 | A→G | N66N (AAT→AAC) | 76.9% | 74.8% | 67.2% | Phage major tail protein pb6 |
| 96,883 | T→A | P54P (CCA→CCT) | 78.1% | 73.7% | 64.9% | Phage major tail protein pb6 |
| 96,889 | T→C | P52P (CCA→CCG) | 76.5% | 74.2% | 65.6% | Phage major tail protein pb6 |
| 96,895 | C→A | A50A (GCG→GCT) | 78.5% | 73.0% | 66.3% | Phage major tail protein pb6 |
| 96,901 | A→G | N48N (AAT→AAC) | 77.3% | 73.2% | 65.2% | Phage major tail protein pb6 |
| 96,916 | C→T | T43T (ACG→ACA) | 79.3% | 73.3% | 69.5% | Phage major tail protein pb6 |
| 96,921 | T→A | T42S (ACA→TCA) | 79.9% | 74.6% | 69.8% | Phage major tail protein pb6 |

| | | | | | | |
|---|---|---|---|---|---|---|
| 96,984 | T→C | N21D (AAT→GAT) | 76.1% | 44.5% | 70.1% | Phage major tail protein pb6 |
| 96,985 | A→G | H20H (CAT→CAC) | 76.6% | 45.2% | 70.2% | Phage major tail protein pb6 |
| 97,122 | C→T | E146E (GAG→GAA) | 80.7% | 46.9% | 64.6% | Phage protein |
| 97,198 | A→G | L121P (CTA→CCA) | 76.4% | 51.9% | 63.2% | Phage protein |
| 97,269 | G→A | D97D (GAC→GAT) | 79.4% | | | Phage protein |
| 97,275 | C→T | V95V (GTG→GTA) | 81.1% | | | Phage protein |
| 104,91 | G→A | L266L (CTC→CTT) | | 31.5% | | Phage terminase, large subunit |
| 104,913 | G→A | D265D (GAC→GAT) | | 31.3% | | Phage terminase, large subunit |
| 104,925 | C→A | E261D (GAG→GAT) | | 28.2% | | Phage terminase, large subunit |
| 104,928 | G→A | I260I (ATC→ATT) | | 27.9% | | Phage terminase, large subunit |
| 104,937 | G→A | F257F (TTC→TTT) | | 30.1% | | Phage terminase, large subunit |
| 104,955 | A→G | G251G (GGT→GGC) | | 28.0% | | Phage terminase, large subunit |
| 104,967 | G→A | S247S (TCC→TCT) | | 26.0% | | Phage terminase, large subunit |
| 104,982 | A→G | Y242Y (TAT→TAC) | | 22.1% | | Phage terminase, large subunit |
| 104,988 | T→C | Q240Q (CAA→CAG) | | 22.5% | | Phage terminase, large subunit |
| 104,997 | A→G | Y237Y (TAT→TAC) | | 22.3% | | Phage terminase, large subunit |
| 105,007 | T→C | N234S (AAT→AGT) | | 20.6% | | Phage terminase, large subunit |
| 105,012 | A→T | T232T (ACT→ACA) | | 18.8% | | Phage terminase, large subunit |
| 105,021 | C→T | A229A (GCG→GCA) | | 17.9% | | Phage terminase, large subunit |
| 105,03 | G→A | I226I (ATC→ATT) | | 17.2% | | Phage terminase, large subunit |
| 105,036 | G→A | N224N (AAC→AAT) | | 17.3% | | Phage terminase, large subunit |
| 105,041 | G→A | L223L (CTG→TTG) | | 19.4% | | Phage terminase, large subunit |
| 105,045 | A→G | A221A (GCT→GCC) | | 19.1% | | Phage terminase, large subunit |
| 105,066 | C→T | T214T (ACG→ACA) | | 22.2% | | Phage terminase, large subunit |
| 105,088 | C→T | G207D (GGC→GAC) | | 15.7% | | Phage terminase, large subunit |
| 105,089 | C→T | G207S (GGC→AGC) | | 16.3% | | Phage terminase, large subunit |
| 105,093 | T→C | L205L (TTA→TTG) | | 15.5% | | Phage terminase, large subunit |
| 105,099 | T→A | E203D (GAA→GAT) | | 15.7% | | Phage terminase, large subunit |
| 105,102 | G→A | N202N (AAC→AAT) | | 15.2% | | Phage terminase, large subunit |
| 105,104 | T→C | N202D (AAC→GAC) | | 15.7% | | Phage terminase, large subunit |
| 105,108 | C→T | G200G (GGG→GGA) | | 15.5% | | Phage terminase, large subunit |
| 105,111 | T→G | K199N (AAA→AAC) | | 15.9% | | Phage terminase, large subunit |
| 105,113 | T→A | K199* (AAA→TAA) | | 16.3% | | Phage terminase, large subunit |
| 105,114 | C→G | E198D (GAG→GAC) | | 16.1% | | Phage terminase, large subunit |
| 105,115 | T→G | E198A (GAG→GCG) | | 16.3% | | Phage terminase, large subunit |
| 105,117 | A→G | Y197Y (TAT→TAC) | | 15.0% | | Phage terminase, large subunit |
| 105,12 | G→A | F196F (TTC→TTT) | | 14.8% | | Phage terminase, large subunit |
| 105,141 | A→T | G189G (GGT→GGA) | | 15.9% | | Phage terminase, large subunit |
| 105,15 | A→G | T186T (ACT→ACC) | | 14.4% | | Phage terminase, large subunit |
| 105,156 | G→A | I184I (ATC→ATT) | | 12.4% | | Phage terminase, large subunit |
| 105,168 | T→C | K180K (AAA→AAG) | | 13.0% | | Phage terminase, large subunit |
| 105,174 | G→A | N178N (AAC→AAT) | | 9.9% | | Phage terminase, large subunit |
| 105,177 | T→A | P177P (CCA→CCT) | | 10.9% | | Phage terminase, large subunit |
| 105,18 | C→T | K176K (AAG→AAA) | | 9.6% | | Phage terminase, large subunit |
| 105,198 | A→C | L170L (CTT→CTG) | | 8.9% | | Phage terminase, large subunit |
| 105,206 | T→C | I168V (ATT→GTT) | | 6.5% | | Phage terminase, large subunit |
| 105,207 | G→C | D167E (GAC→GAG) | | 6.1% | | Phage terminase, large subunit |
| 105,208 | T→C | D167G (GAC→GGC) | | 6.2% | | Phage terminase, large subunit |
| 105,209 | C→T | D167N (GAC→AAC) | | 5.4% | | Phage terminase, large subunit |
| 105,213 | T→G | A165A (GCA→GCC) | | 6.7% | | Phage terminase, large subunit |
| 105,216 | T→A | A164A (GCA→GCT) | | 6.6% | | Phage terminase, large subunit |
| 105,217 | G→T | A164E (GCA→GAA) | | 6.2% | | Phage terminase, large subunit |
| 105,222 | C→A | G162G (GGG→GGT) | | 6.2% | | Phage terminase, large subunit |

| | | | | |
|---|---|---|---|---|
| 105,225 | A→C | V161V (GTT→GTG) | 6.8% | Phage terminase, large subunit |
| 105,228 | G→A | D160D (GAC→GAT) | 5.6% | Phage terminase, large subunit |
| 105,231 | T→A | S159S (TCA→TCT) | 6.0% | Phage terminase, large subunit |
| 105,237 | C→T | A157A (GCG→GCA) | 5.4% | Phage terminase, large subunit |
| 105,24 | T→C | A156A (GCA→GCG) | 6.0% | Phage terminase, large subunit |
| 105,243 | T→C | E155E (GAA→GAG) | 5.9% | Phage terminase, large subunit |
| 105,246 | A→G | D154D (GAT→GAC) | 6.4% | Phage terminase, large subunit |
| 105,258 | G→A | F150F (TTC→TTT) | 8.7% | Phage terminase, large subunit |
| 105,267 | C→T | S147S (TCG→TCA) | 10.7% | Phage terminase, large subunit |
| 105,3 | A→G | S136S (TCT→TCC) | 13.8% | Phage terminase, large subunit |
| 105,306 | C→T | L134L (CTG→CTA) | 12.8% | Phage terminase, large subunit |
| 105,312 | G→A | F132F (TTC→TTT) | 11.3% | Phage terminase, large subunit |
| 105,315 | G→T | L131L (CTC→CTA) | 12.2% | Phage terminase, large subunit |
| 105,318 | C→A | S130S (TCG→TCT) | 12.4% | Phage terminase, large subunit |
| 105,321 | G→A | G129G (GGC→GGT) | 11.2% | Phage terminase, large subunit |
| 105,324 | G→A | N128N (AAC→AAT) | 11.4% | Phage terminase, large subunit |
| 105,327 | T→A | A127A (GCA→GCT) | 11.8% | Phage terminase, large subunit |
| 105,33 | C→T | L126L (CTG→CTA) | 11.4% | Phage terminase, large subunit |
| 105,332 | G→A | L126L (CTG→TTG) | 10.8% | Phage terminase, large subunit |
| 105,351 | G→A | A119A (GCC→GCT) | 17.8% | Phage terminase, large subunit |
| 105,366 | A→G | T114T (ACT→ACC) | 22.5% | Phage terminase, large subunit |
| 105,369 | C→T | Q113Q (CAG→CAA) | 22.6% | Phage terminase, large subunit |
| 105,374 | A→G | L112L (TTA→CTA) | 22.5% | Phage terminase, large subunit |
| 105,375 | T→G | G111G (GGA→GGC) | 22.1% | Phage terminase, large subunit |
| 105,378 | A→G | Y110Y (TAT→TAC) | 22.4% | Phage terminase, large subunit |
| 105,405 | T→A | S101S (TCA→TCT) | 24.0% | Phage terminase, large subunit |
| 105,411 | A→T | G99G (GGT→GGA) | 23.4% | Phage terminase, large subunit |
| 105,417 | G→A | N97N (AAC→AAT) | 21.6% | Phage terminase, large subunit |
| 105,42 | A→G | A96A (GCT→GCC) | 21.8% | Phage terminase, large subunit |
| 105,438 | T→A | A90A (GCA→GCT) | 25.1% | Phage terminase, large subunit |
| 105,444 | A→T | V88V (GTT→GTA) | 23.3% | Phage terminase, large subunit |
| 105,447 | A→T | L87L (CTT→CTA) | 23.6% | Phage terminase, large subunit |
| 105,459 | A→G | N83N (AAT→AAC) | 25.8% | Phage terminase, large subunit |
| 105,462 | G→A | P82P (CCC→CCT) | 24.4% | Phage terminase, large subunit |
| 105,470:1 | +C | coding (238/1317 nt) | 21.3% | Phage terminase, large subunit |
| 105,473 | G→C | L79V (CTT→GTT) | 21.6% | Phage terminase, large subunit |
| 105,477 | Δ1 bp | coding (231/1317 nt) | 21.7% | Phage terminase, large subunit |
| 105,483 | T→C | G75G (GGA→GGG) | 26.0% | Phage terminase, large subunit |
| 105,504 | A→G | S68S (TCT→TCC) | 27.6% | Phage terminase, large subunit |
| 105,51 | A→G | G66G (GGT→GGC) | 26.3% | Phage terminase, large subunit |
| 105,519 | A→G | R63R (CGT→CGC) | 29.8% | Phage terminase, large subunit |
| 105,534 | T→C | T58T (ACA→ACG) | 29.3% | Phage terminase, large subunit |
| 105,54 | G→A | F56F (TTC→TTT) | 28.4% | Phage terminase, large subunit |
| 105,725 | T→A | A155A (GCA→GCT) | 7.3% | Phage protein |
| 105,731 | C→G | S153S (TCG→TCC) | 6.9% | Phage protein |
| 105,752 | A→G | N146N (AAT→AAC) | 6.7% | Phage protein |
| 105,755 | C→T | Q145Q (CAG→CAA) | 5.7% | Phage protein |
| 105,761 | A→C | G143G (GGT→GGG) | 5.8% | Phage protein |
| 105,766 | A→C | S142A (TCC→GCC) | 5.4% | Phage protein |
| 105,767 | G→T | G141G (GGC→GGA) | 5.2% | Phage protein |
| 105,788 | A→G | I134I (ATT→ATC) | 5.4% | Phage protein |
| 105,791 | A→G | N133N (AAT→AAC) | 6.2% | Phage protein |

| | | | | |
|---|---|---|---|---|
| 105,794 | A→G | T132T (ACT→ACC) | 5.8% | Phage protein |
| 105,806 | C→T | P128P (CCG→CCA) | 5.5% | Phage protein |
| 105,838 | G→A | L118L (CTG→TTG) | 5.4% | Phage protein |
| 105,866 | C→T | K108K (AAG→AAA) | 6.4% | Phage protein |
| 105,872 | C→T | K106K (AAG→AAA) | 6.6% | Phage protein |
| 105,875 | G→A | H105H (CAC→CAT) | 6.5% | Phage protein |
| 105,878 | C→T | A104A (GCG→GCA) | 6.7% | Phage protein |
| 105,881 | C→T | K103K (AAG→AAA) | 6.3% | Phage protein |
| 105,899 | G→A | I97I (ATC→ATT) | 6.4% | Phage protein |
| 105,908 | G→A | D94D (GAC→GAT) | 6.6% | Phage protein |
| 105,923 | A→G | T89T (ACT→ACC) | 7.2% | Phage protein |
| 105,926 | T→C | E88E (GAA→GAG) | 7.6% | Phage protein |
| 105,932 | A→T | L86L (CTT→CTA) | 7.3% | Phage protein |
| 105,938 | T→C | E84E (GAA→GAG) | 7.4% | Phage protein |
| 105,944 | C→T | K82K (AAG→AAA) | 5.7% | Phage protein |
| 105,965 | A→T | L75L (CTT→CTA) | 5.9% | Phage protein |
| 105,971 | C→A | G73G (GGG→GGT) | 5.2% | Phage protein |
